# High temperature ligation enables *in vitro* assembly of eukaryotic genes

**DOI:** 10.64898/2026.10.09.758000

**Authors:** Noah Evan Robinson, Bryan Gerber, Weilin Zhang, Sixiang Wang, Daniel Graves, Hanqiao Zhang, Jiahe Yue, Gohta Aihara, Jerry Huang, Kaihang Wang

## Abstract

Our ability to efficiently construct synthetic DNA products is essential to studying and engineering biology. There is an abundance of widely adopted DNA assembly technologies suitable for generating synthetic constructs from diverse DNA inputs, but these techniques suffer from low specificity of connection during assembly, which decreases the accuracy, efficiency, and throughput of construction. Here we propose a broadly applicable paradigm for DNA assembly, High Temperature (HighT) assembly, that lowers misconnection rates by ligating complementary single strand DNA overhangs at reaction temperatures significantly higher than their predicted melting temperatures. By expanding the concept of melting temperature to demarcate smaller association fractions of DNA duplexes, we characterize HighT and demonstrate 40 percentage points increase in accuracy, and 4 orders of magnitude increase in yield of transformants when applying the technique to conventional assembly technologies. HighT is then applied to develop cloning pipelines for the development of efficient assembly-to-integration protocols, the construction of specific low-abundance isoforms of medically relevant proteins from human gDNA and oligos, and the de novo construction of a 10 kb, GC rich genomic region from the functionally extinct Northern White Rhinoceros created entirely in vitro.

## Main

Synthetic DNA is crucial to the contemporary study of biology as it enables us to precisely control the information encoded in engineered constructs to more systematically probe the correlation between genotype and phenotype^1,2^. Long synthetic DNA molecules are constructed by assembling smaller DNA fragments derived from chemically or enzymatically synthesized oligonucleotides (oligos), PCR, or sourced from pre-existing DNA templates such as bacterial plasmids, genomic DNA (gDNA), or complementary DNA (cDNA). Through a combination of these diverse inputs, we can cover the theoretical sequence space required for bioengineering of diverse systems through the efficient construction of DNA molecules.

Our ability to construct DNA polymers from any source is limited by the misconnection rate between DNA fragments during assembly — a problem compounded by an increasing number of assembly junctions^3^. We previously developed the novel DNA assembly technique Sidewinder to address this misconnection issue and enable the assembly of large numbers of fragments into complex DNA constructs^3^. Sidewinder assembly is directed by highly optimized barcode sequences not present in the final construct via a 3-Way Junction (3WJ) intermediate, enabling specificity measured as a 1 in 1,000,000 misconnection rate^3^.

Sidewinder improves specificity via the 3WJ, yet other DNA assembly techniques rely on the native 2-Way Junction (2WJ) to direct the association between molecules^3^. The 2WJ suffers from higher misconnection rates as the junction information which directs assembly is constrained to the sequence space of the final synthetic construct^3–11^, leading to unoptimized assemblies where exposed single strand DNA (ssDNA) overhangs can form secondary structures, mis-associate, and mis-ligate. To increase reaction specificity while operating within 2WJ sequence constraints, we observed that previous 2WJ techniques share a common feature where annealing and ligation reactions typically occur at room temperature or below the melting temperatures (T_m_) of the DNA overhangs participating in the assembly reaction^12–16^. Contrasting with this existing paradigm, we propose High Temperature (HighT) assembly, defined by ligation at temperatures significantly higher than the melting temperature of assembly overhangs to increase the stringency of 2WJ connection rate for larger and more robust assemblies with reduced incorrect byproducts.

In this study, we first propose a mechanism and demonstrate proof-of-principle for HighT assembly (**Fig. 1**). Then we validate that HighT ligations enable more robust 2WJ assembly by systematically optimizing 2WJ design and construction framework while using a range of input DNA including plasmids, gDNA, and oligo assemblies. Ultimately, the advancements to 2WJ assembly methods with HighT can result in lower misconnection rates, over 10,000× increase in the number of correct transformants, and enable direct PCR amplification from HighT reactions which can be used in subsequent HighT rounds for *in vitro* hierarchical assembly of larger constructs without the need for *in vivo* intermediates.

**Fig. 1.**
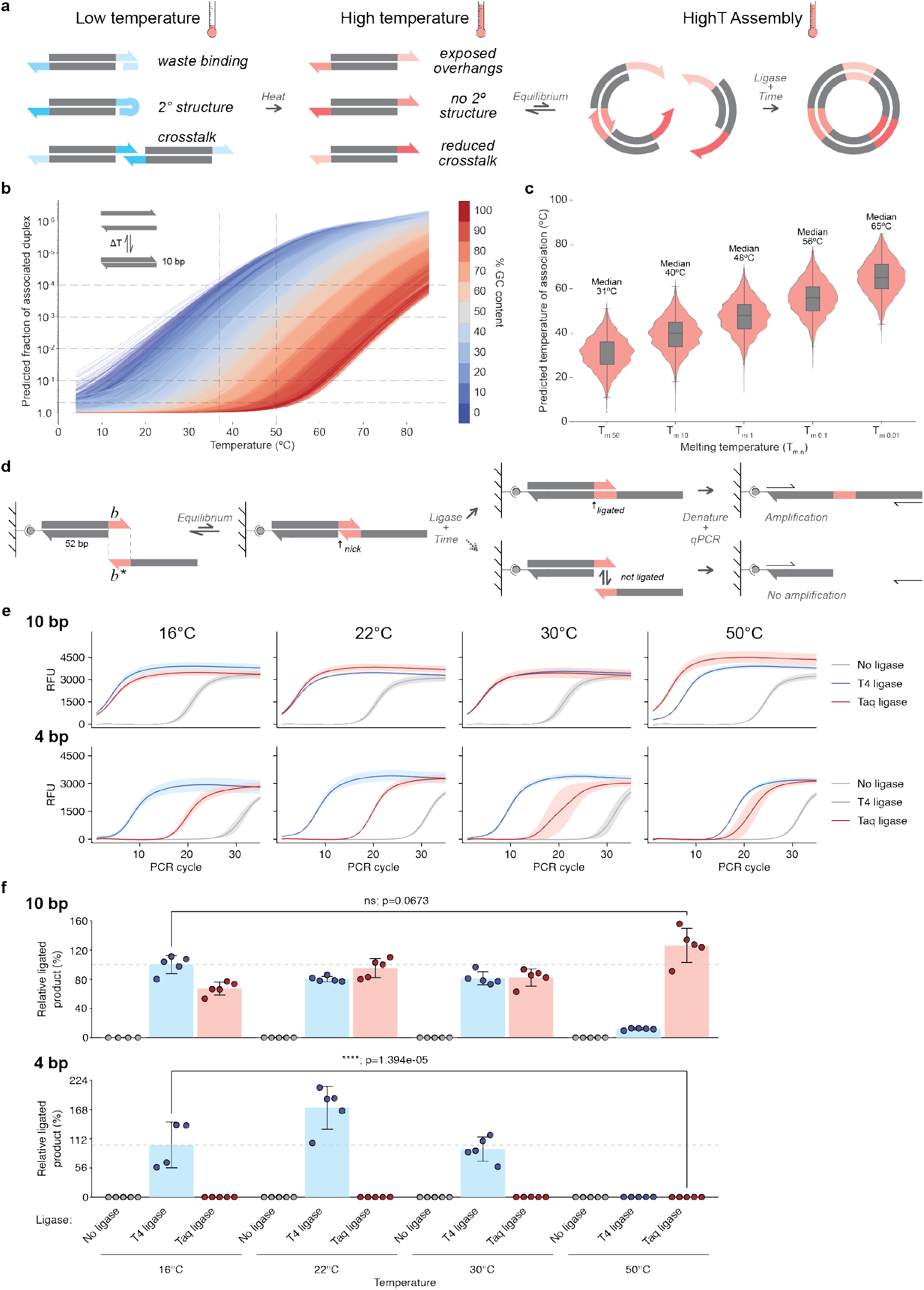
High temperatures increase specificity of DNA associations that are ligated over time. **a)** Schematic depicting the advantages offered by High Temperature (HighT) ligation in comparison to low temperature ligation in producing exposed overhangs for assembly. **b)** NUPACK simulated melt curves for all 10 bp duplexes with unpaired intramolecular secondary structures at 50°C. **c)** Violin plot and nested box and whisker plot depicting the distribution of melting temperatures in the form of T_m-n_ for the simulated data presented in **b**. The n demarcates the proportion of species bound in the duplex for a 10 nM solution. Summary statistics can be found in **Source data**. d) Schematic depicting the qPCR assay used to measure ligation activity of 4 bp and 10 bp overhangs under different ligase and temperature conditions. **e)** qPCR amplification curves depicting mean fluorescence ± SD over PCR cycles for the 4 bp and 10 bp overhang ligation reactions at 16°C, 22°C, 30°C, and 50°C with no ligase, T4 DNA Ligase, or Taq DNA Ligase. **f)** Bar plot depicting the mean ± SD of relative ligation product for 4 bp and 10 bp overhangs at 16°C, 22°C, 30°C, and 50°C with no ligase, T4 DNA Ligase, or Taq DNA Ligase. The starting quantity of T4 DNA Ligase at 16°C was defined as 100% and used to normalize all samples.

We then use these improvements to demonstrate rapid one-step multi-fragment assembly-to-integration protocols, a new paradigm for constructing specific low-abundance isoforms of eukaryotic genes from gDNA templates, and construction of a 10-kb, GC-rich linear dsDNA encoding the genomic region from the functionally extinct Northern White Rhinoceros created entirely *de novo* and *in vitro*.

## Results

### Mechanism of High Temperature assembly

While operating within the 2WJ framework that is shared by conventional DNA assembly technologies, there are limited approaches to optimization which do not place restrictions on, or result in scars in, the assembled product such as enhanced design of orthogonal overhang sequences in the given sequence space, improved fragment processing, and assembly conditions (enzymes, buffers, temperatures)^12,17–19^.

Of these factors, assembly temperature remains a nuanced variable that can influence the quality and success of the assembly^6,13^ especially for ligation-based assembly methods where the conventional consideration of temperature has been of the balance between enzyme activity and the stability of the interaction between the overhangs being ligated^12,15^. These two factors seemingly conflict as higher reaction temperatures increase ligase activity while destabilizing interactions between overhangs whereas lower reaction temperatures decrease ligase activity while stabilizing interactions between overhangs^15,20^. Prior literature demonstrations have generally decided in favor of lower temperature ligations, prioritizing overhang stability at the cost of decreased enzyme activity^12–16^. However, we propose high ligation temperatures significantly above the predicted T_m_ of the overhang will improve the accuracy and efficiency of 2WJ assembly methods.

To demonstrate high temperature ligation as a viable alternative, it must be shown that 1) DNA interactions can be more specific at higher temperatures; 2) single stranded DNA overhangs are able to associate, even if transiently, at temperatures above their predicted T_m_; and 3) single stranded DNA overhangs are able to be ligated at temperatures above their predicted T_m_.

Towards providing evidence for claim 1), it has been previously demonstrated that higher temperatures confer increased specificity in the association between DNA duplexes^21,22^ with resolution as sensitive as a single base mismatch at 50°C and above^23^. However, prior arts opt to implement low temperatures to stabilize intended overhang interactions^12–16^. This incidentally also stabilizes all unintended DNA interactions like secondary structure and mis-association between non-partnered fragments (**Fig. 1a, Extended data 1a**). Conversely, high ligation temperatures, in addition to increasing enzyme activity, will reduce overhang secondary structure and reduce instances of mis-association (**Fig. 1a, Extended data 1a**). These precedents and data provide evidence for claim 1) to show that DNA interactions can be more specific at higher temperatures.

We next show that single stranded DNA overhangs are able to associate, even if transiently, at temperatures above their predicted T_m_ utilizing the thermodynamic algorithm NUPACK^24,25^. We simulated melt curves of short DNA duplexes at a range of temperatures to predict the fraction of species associated at a given temperature for a given DNA sequence. The free energy calculations for secondary structures in NUPACK are calculated using nearest-neighbor empirical parameters utilizing a body of measurements over decades of research from multiple groups that provide the basis for rigorous analysis of interacting strands^26–28^. This enabled us to simulate a far greater number of interacting strands than would be feasible with empirical measurement alone. We simulated duplex association at discrete temperatures 4°C-85°C at a 10 nM concentration for all 10 bp DNA duplexes which are predicted by NUPACK to have a self-unpaired secondary structure at 50°C (**Methods**). This resulted in 803,866 valid strands to compose 401,933 unique melt curves that predict association between molecules in salt conditions comparable to a ligation reaction (**Fig. 1b**).

The resultant melt curves depict a broad range of temperatures for which 10 bp duplexes are stably associated, with melt curves generally clustering by the GC content of the duplex. This set of 10 bp duplexes have a median predicted T_m_ (temperature where 50% of species are associated) of 31°C. However, we propose to expand the concept T_m_ to describe other points in the melt curve using the notation T_m-n_ where n is the % of species associated in a duplex. For example, a T_m-10_ would demarcate the temperature where 10% of species are associated, T_m-1_ where 1% of species are associated, and so on. With this expended conceptualization of T_m_ this set of 10 bp duplexes has a median predicted T_m-10_ of 40°C, a median predicted T_m-1_ of 48°C, a median predicted T_m-0.1_ of 56°C, and a median predicted T_m-0.01_ of 65°C (**Fig. 1c**).

Although 17°C higher than the conventional T_m-50_ measurement, these data suggest that at 48°C half of these 10 bp overhangs sequences will have at least 1% of species associated. We do not propose that a T_m-1_ is intrinsically preferable to T_m-50_ but rather a ligation at T_m-1_, in so far as 1% association is sufficient for ligation to occur over time, will enable the reaction system to benefit from the improved specificity at higher temperatures and could yield overall better assembly results. Similarly, we do not propose that continually increasing temperature will continue to improve reaction fidelity since very high temperatures are universally destabilizing to both intended and unintended duplex formation (**Fig. 1b, Extended data 2**). Under such conditions, the differentiation between intended and unintended duplex afforded by HighT is weakened, such that there may be an optimal temperature where differentiation is highest that will depend on the particular interacting species and other reaction conditions.

This then leads to claim 3) to demonstrate that ligation can occur at temperatures significantly above the reaction T_m-50_. Prior studies have demonstrated that ligation significantly above T_m-50_ is experimentally feasible^14,29,30^. For example, Wang et al. showed that dsDNA with 4 bp overhangs can undergo efficient ligation at 45–70°C using Tth DNA ligase, despite having predicted T_m-50_ values below 10°C ^30^. Similarly, the demonstration of the 52-piece Golden Gate assembly yielded a more accurate assembly when conducted at a fixed 37°C temperature when compared to a 16°C-37°C cycling protocol^13,29^. While the specific T_m-n_ value for these particular overhangs at these temperatures will differ from one another, in each case, the fraction of DNA duplexes which are transiently associated may associate long enough for ligation to irreversibly connect the two molecules together. In the context of assembly, a ligation removes the two interacting species from the equilibrium of unpaired ⇌ paired, allowing the reaction to tend toward completion over time (**Fig. 1a**) by Le Chatelier’s principle^31^.

To demonstrate this empirically, we designed an experimental assay to show that single stranded DNA overhangs, at some frequency, can be ligated at temperatures above their predicted T_m-50_ which enables the reaction to tend towards completion over time. A pre-annealed dsDNA duplex with an exposed 4 or 10 bp overhang was bound to a streptavidin coated PCR tube. Then a reaction mixture containing ligase and free ssDNA oligo with the complementary overhang was added and incubated at 16°C, 22°C, 30°C, and 50°C (**Fig. 1d**). After ligation, the tubes were washed, rigorously denatured with 20 mM NaOH, and washed again such that the only products remaining in solution will be either ligated (112 bases) or un-ligated (62 bases) ssDNA oligo fixed to the solid surface (**Fig. 1d**). We then used qPCR with specific primers (**Fig. 1d**) for a precise measurement of the ligation product independent from an individual annealing event as only upon ligation is the valid qPCR amplification target generated.

We established ligation reaction conditions with no ligase (control), T4 DNA Ligase, and Taq DNA Ligase. For both the 4 bp and 10 bp overhang, T4 and Taq ligase produced a lower mean Cq value than the no ligase control at all four temperatures, indicating ligated product formation under all tested conditions. (**Fig. 1e, Extended data 1**). To compare the ligation yield across samples, we normalized all samples’ starting quantity (qPCR measurement of ligated product) to T4 DNA Ligase at 16°C, the manufacturer recommended ligation temperature, which we define here as an arbitrary 100% ligation efficiency.

For the 10 bp overhangs, we observe no significant difference in relative ligation efficiency between T4 ligase at 16°C and the HighT condition with Taq ligase at 50°C (two-sided Welch’s t-test on log₂-transformed starting quantities, p = 0.067). Importantly, ligation product was generated despite this duplex’s predicted T_m-50_ of 31°C whereas 50°C corresponds to the T_m-0.53_ indicating only 0.53% of species in association at equilibrium. This provides evidence showing that small fraction of associated overhangs could lead to efficient ligation under long incubation, supporting claim 3 (**Fig. 1f, Extended Data Figure 1b**).

For the 4 bp overhang, the T4 ligase conditions at 16°C, 22°C, and 30°C yield comparable relative ligation efficiencies to each other (**Fig. 1f**). At 50°C, T4 DNA ligase’s relative ligation efficiency decreases significantly. This may be due to the choice of ligase (T4 ligase and not High-T4 ligase) as we saw a decrease in T4 ligase activity at 50°C with 10 bp overhangs as well. Taq ligase also showed substantially lower ligation efficiency relative to T4 DNA Ligase at all temperature conditions but still showed some evidence of ligation as compared to the negative control.

These simulated and empirical results demonstrate the theoretical viability for HighT assembly. The conceptual differentiation of low-association T_m_ values allowed us to propose a mechanism where infrequent molecular associations at higher temperatures still occur often enough for successful ligation, resulting in increased reaction specificity via the globally destabilizing reaction conditions favoring association of perfectly matched overhangs.

### Optimization of 2WJ PCR-based assembly

Having conceptually and experimentally validated the feasibility of ligation at high temperatures, we designed a multi-piece USER assembly (**Extended Data 3**) that was assembled under different reaction conditions to test HighT improvements to multi-fragment DNA assembly (**Fig. 2**).

**Fig. 2.**
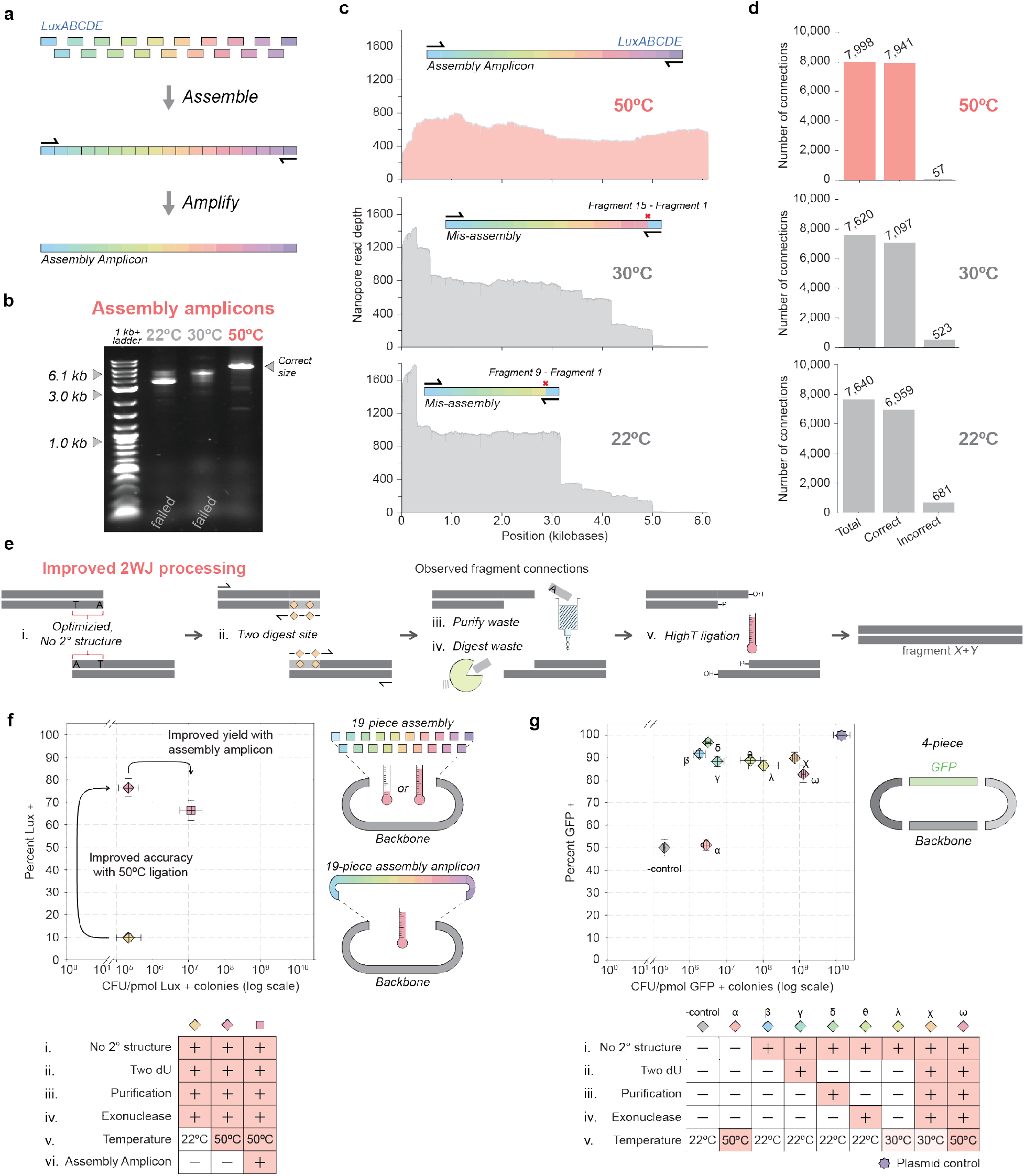
High temperature ligation enables improved accuracy and yield of DNA assembly. **a)** Schematic representation of a multi-fragment assembly and subsequent PCR amplification generating an Assembly Amplicon. **b)** DNA agarose gel depicting 1µL of PCR product comparing the Assembly Amplicon of the *LuxABCDE* cassette assembled at 22°C, 30°C, and 50°C. **c)** Coverage plot depicting the consensus sequence of the *LuxABCDE* Assembly Amplicon at the different assembly temperatures. **d)** Bar plots depicting analysis of all possible fragment connections in the *LuxABCDE* Assembly Amplicon Nanopore sequencing data, comparing the number of correct and incorrect connections at 22°C, 30°C and 50°C. **e)** Schematic depicting the improved 2WJ cloning mechanism with corresponding modifications to the protocol which improve the reaction accuracy and efficiency. **f)** Plot depicting the transformation accuracy (y axis) vs yield (x axis) in colony forming units per pmol plasmid backbone transformed for 3 biological replicates of the *Lux* cassette 20-piece at 22°C, 20-piece at 50°C, and 20-piece 50°C Assembly Amplicon. The combination of improvements **i-vi** are shown with red (+). **g)** Plot depicting the transformation accuracy (y axis) vs yield (x axis) in colony forming units per pmol plasmid backbone transformed for 3 biological replicates of the 4-piece GFP assembly. The combination of improvements **i-v** used for each condition **α** - **ω** are shown with red (+).

When conducting multi-fragment assembly, we had previously seen with Sidewinder that the increased specificity conferred by its 3WJ design enabled direct PCR from assembled product which allowed for specific isolation and amplification of full-length assembly product^3^. Owing to HighT’s higher 2WJ specificity conferred by increased ligation temperatures, we hypothesized that HighT may also enable a PCR to be conducted using the assembly reaction as template, resulting in an “Assembly Amplicon” (**Fig. 2a**). Due to PCR’s bias towards amplification of smaller products^32^ we believe other 2WJ protocols do not implement PCR of the assembled product since any mis-ligations resulting in truncated product may be preferentially amplified, preventing amplification of the full-length target product. However, the HighT improvements to 2WJ assembly may substantially reduce the presence of truncated products in the assembly reaction, enabling preferential amplification of full-length assembly product. PCR of an Assembly Amplicon would allow for entirely *in vitro* selection of the properly assembled final product, increasing the yield and enabling the possibility of *in vitro* hierarchical assembly with multiple rounds of assembly without the need for cellular intermediates.

Towards this end, we designed a 19-piece assembly that split the 6.1 kb *LuxABCDE* cassette into approximately equal length fragments and conducted ligations at 22°C, 30°C, and 50°C. Conducting a PCR after the ligation at each of the corresponding temperatures produced amplicons in each case, but only the 50°C protocol produced a strong band at the correct size corresponding to the Assembly Amplicon of the 6.1 kb target whereas the assemblies conducted at lower ligation temperatures produced amplicons of the incorrect size (**Fig. 2b**). Nanopore sequencing of the Assembly Amplicons allows insight into the identities of the final products. When aligned to the target 6.1kb *Lux* sequence, the 50°C Assembly Amplicon had consistent coverage spanning the entire *Lux* cassette with fragments in the correct order and correct orientation (**Fig. 2c**). In contrast, the 22°C and 30°C assemblies suffered from inconsistent coverage, limited full length final product, and precipitous drops in coverage consistent with mis-assemblies between fragments 1 & 9 at 22°C and mis-assemblies between fragments 2 & 12 and 1 & 15 at 30°C (**Fig. 2c**).

The sequencing data was then checked for all possible fragment connections to check for correctly or incorrectly ligated assemblies (**Methods**). The 22°C and 30°C Assembly Amplicons had a junction mis-connection rate of approximately 1 in 11 (91.1% accurate) and 1 in 15 (93.1% accurate) while 50°C had a reduced misconnection rate to approximately 1 in 140 (99.3% accurate) (**Fig. 2d**), thus both qualitatively and quantitively demonstrating that HighT assembly is feasible and confers greater reaction specificity.

The 19-piece *Lux* assembly was also assembled with a backbone and transformed to compare yield and accuracy both with and without HighT and Assembly Amplicon. When ligating at 22°C (no HighT), with an improved protocol for fragment processing (**Fig. 2e**, **Methods**), we produced a full 20-piece assembly with an average accuracy of 9.8% Lux+ colonies with a yield (defined as the normalized number of phenotypically correct clones) of 2.17 × 10^5^ Lux+ colonies per pmol of assembly transformed (**Fig. 2f**). With HighT ligation at 50°C we observed greatly improved accuracy to 76.4% Lux+ clones while maintaining a comparable yield of 2.14 × 10^5^ Lux+ colonies/pmol (**Fig. 2f**). When cloning the HighT Assembly Amplicon of the *Lux* cassette, we observed a slight decrease in proportion of Lux+ clones at 66.4% but gain nearly two orders of magnitude increased yield of 1.18 × 10^7^ Lux+ colonies/pmol (**Fig. 2f**). This demonstrates HighT conferring increased accuracy to multi-fragment assemblies and further validates the hypothesis that the HighT-enabled Assembly Amplicon and increases yield of assembled products.

The literature characterizations of USER generally demonstrate relatively small assemblies, usually around 5 DNA fragments assembled simultaneously with a decrease in accuracy and efficiency as the number of fragments increases^12,33^. In contrast, our HighT assembly protocol and the utilization of Assembly Amplicons enable larger multi-fragment assemblies to be conducted. These results were accomplished using HighT in addition to a series of other improvements which are broadly applicable to 2WJ techniques to aid in assembly (**Fig. 2f**). These improvements all work to optimize the overhangs generated for the assembly to be exposed mutually exclusive sequences free from secondary structure to improve association and ligation to the intended assembly partner (**Fig. 2e**). This includes: (**i**) optimizing overhang sequences to be free from secondary structures or cross reactivity, (**ii**) additional digestion sites for exposing overhangs, (**iii**) purification of digested fragments, (**iv**) enzymatic digestion of waste, in addition to (**v**) HighT ligation (**Fig. 2e**).

For a 4-piece USER assembly of a green fluorescent protein (GFP) plasmid (where assembly accuracy can be calculated by the proportion of the green-fluorescence (GFP+) phenotype (**Extended Data 3c**)), a published literature USER protocol with 22°C ligation^12^ resulted in an average accuracy of 50.0% GFP+ colonies with a yield of 2.01 × 10^5^ GFP+ colonies/pmol plasmid transformed. For a HighT ligation at 50°C, we observed a comparable accuracy of 51.2% GFP+ clones but observed an order of magnitude increase in yield to 2.79 × 10^6^ GFP+ colonies/pmol (**Fig. 2g α**). This assembly requires the presence of three of the four assembly fragments for cell survival likely conferring the increased colony count at higher temperatures for this construct.

We then redesigned our 4-piece assembly to ensure that the USER overhang sequences had reduced secondary structure using the NUPACK web application^24^ (**Methods**). We also used NUPACK to assess all exposed ssDNA overhangs participating in the reaction for crosstalk between non-partnered assembly fragments. With the redesigned no secondary structure (NSS) overhangs we were able to increase the average accuracy of our assembly from 50.0% to 91.8% and increase efficiency from 2.01 × 10^5^ GFP+ colonies/pmol to 1.82 × 10^6^ GFP+ colonies/pmol with ligation at 22°C (**Fig. 2g β**). We found that the NSS overhangs significantly enhanced the improvements to accuracy and yield afforded by the subsequent protocol modifications compared to when secondary structure is present (**Extended Data 3d**). This suggests that the optimization of overhang sequences is another essential factor to consider for successful 2WJ assembly.

The subsequent modifications to the protocol are all shown using our improved NSS overhangs, as each offers additional advantages over this new established baseline by ensuring more exposed ssDNA overhang participate in the assembly process. These modifications and their resultant improvements were: including two digestion sites (**Fig. 2g γ**, 88.3% fluorescent with 5.61 × 10^6^ GFP + colonies/pmol), purification of digested product (**Fig. 2g δ,** 96.7% fluorescence with 3.14 × 10^6^ GFP + colonies/pmol), exonuclease digestion of residual waste (**Fig. 2g θ**, 88.7% fluorescence with 4.47 × 10^7^ GFP + colonies/pmol), and lastly HighT assembly (**Fig. 2g λ**, 86.5% fluorescence with 1.03 × 10^8^ GFP + colonies/pmol).

We then tested an additional assembly condition that combines all modifications into a single reaction, which when conducted with 30°C ligation resulted in an accuracy of 89.9% and a yield of 7.41 × 10^8^ GFP + colonies/pmol (**Fig. 2g χ**) and HighT 50°C ligation resulted in an accuracy of 82.7% and an improved yield of 1.27 × 10^9^ GFP + colonies/pmol (**Fig. 2g ω**). Overall, the combination of these optimizations can result in a 40-percentage point increase in accuracy (**χ** 89.9% compared to -control 50.0%), while also resulting in nearly 10,000× increased yield of correct clones. These reaction conditions also enable us to approach the theoretical maximum accuracy and yield for this construct, as demonstrated by the transformation of the positive control (a nicked miniprep plasmid) (**Fig. 2g**).

### One-step assembly and integration

In addition to carrying DNA on episomal replicons, genomic integration of synthetic DNA is often deployed for improved construct stability^34^ and expression^35^. A variety of mechanisms have been used to integrate large pieces of DNA into genomes, including naturally occurring bacteriophage integrases to direct insertions, excisions, inversions and translocations of DNA^36,37^. We theorized that the high yield and low mis-assembly rate afforded by HighT could enable rapid DNA assembly, transformation, and integration of multipiece synthetic inserts into recipient genomes without episomal replicon intermediates. To accomplish this, we developed two independent integration schemes, our first integrates a circularized HighT construct, inserting DNA at a predefined target location, and the second integrates a linear HighT construct, replacing DNA at a predefined target location.

For circular integration, we used the HK022 recombinase to direct integration. This recombinase could theoretically work interchangeably with similar recombinases (lambda phage, phage 186, phi80, and P21)^38^. These systems rely upon three major components: a recombinase, an attP recombination site, and its compatible attB recombination site. Using these three components, we created a mechanism to integrate HighT synthetic DNA into a user defined location within recipient genomes (**Fig. 3a**). We designed our system such that successful integration at a defined location would result in the true gain of a transcriptionally active antibiotic marker.

**Fig 3.**
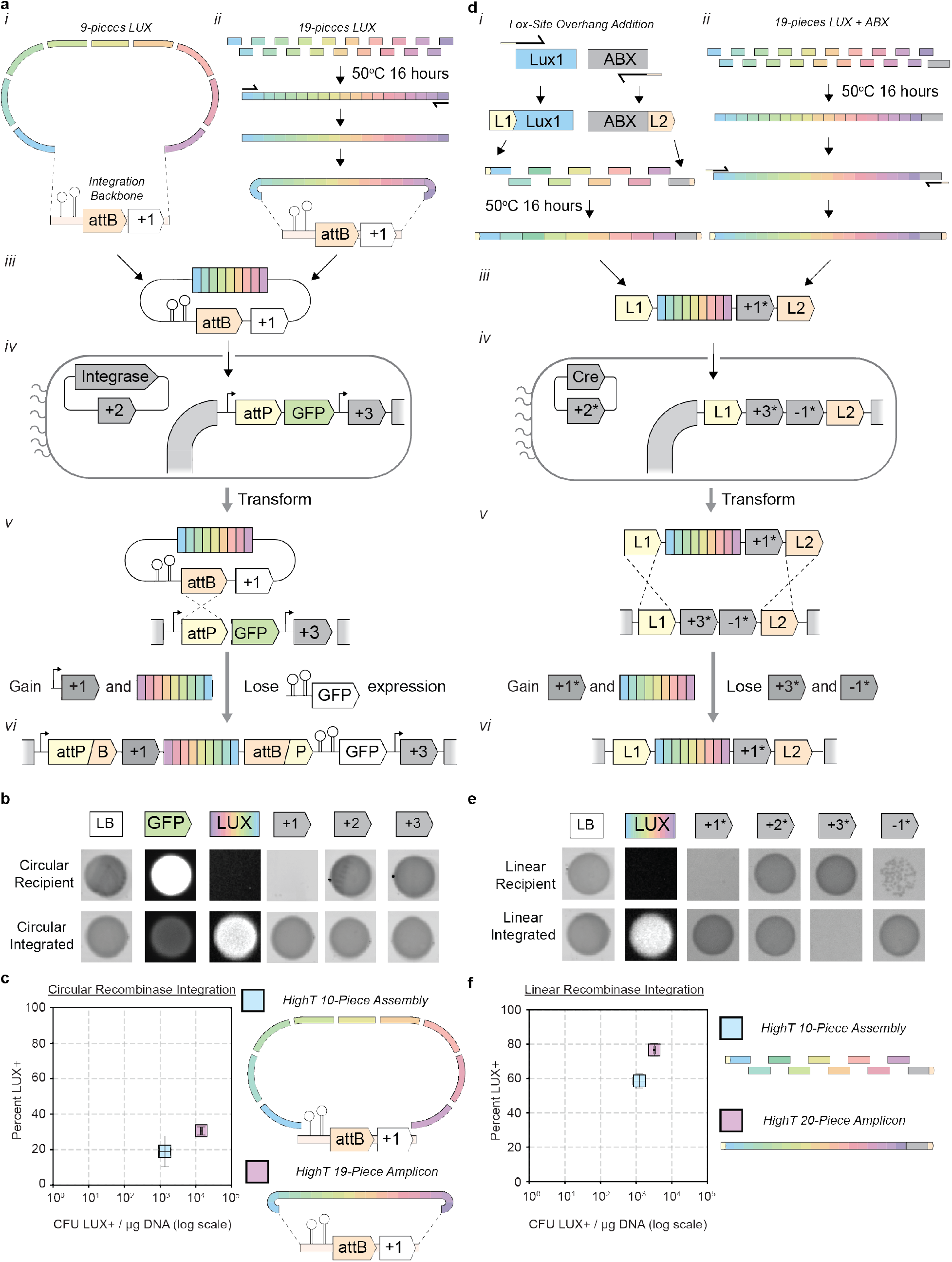
HighT assembly coupled with two independent genome integration schemes. **a)** Schematic depicting circular HighT assembly genomically integrated using a circular recombination system. *LuxABCDE* is divided into 9 fragments, which are HighT ligated to an integration backbone fragment containing two terminators, an attB sequence, and a transcriptionally inactive antibiotic resistance marker +1 (**i**). *LuxABCDE* is divided into 19 fragments, which are HighT ligated and amplified into an Assembly Amplicon that is subsequently assembled with the same integration backbone (**ii**). The final assembly product of both constructions (**iii**) are electroporated into recipient cells with a plasmid containing the attB integrase, antibiotic resistance marker +2 and a pre-engineered genome containing a promoter, attP site, transcriptionally active GFP, and antibiotic resistance marker +3 (**iv**). Through protein mediated DNA crossover (**v**) the recipient genome gains transcriptionally active antibiotic resistance marker +1, the cargo insert, and losses fluorescence expression in the post-integrated genome (**vi**). **b)** Phenotypic spot assay where pre-integration attBP circular recipient is compared to post-integration circular attBP cells on various growth media (+1 = kanamycin, +2 = carbenicillin, +3 = chloramphenicol). **c)** Plot depicting the transformation accuracy (y axis) vs yield (x axis) in colony forming units per pmol plasmid backbone transformed for 3 biological replicates of the 10-piece scheme (**i**) Average accuracy=19.1%, Standard deviation=10.7%, Average yield=1.37 × 10^3^ Lux + colonies / µg DNA, Standard deviation= 5.51 × 10^2^ Lux + colonies/ µg DNA) and the 19-piece Assembly Amplicon scheme (**ii**) (Average accuracy=30.6%, Standard deviation=2.8%, Average yield=1.43 × 10^4^ Lux + colonies/ µg DNA, Standard deviation=1.33× 10^3^ Lux + colonies/ µg DNA). **d)** Schematic depicting LoxP (L1) and Lox2272 (L2) recombination sites onto distal ends of integrated cargo to enable Cre recombinase mediated genome integration. **e)** Phenotypic spot assay where pre-integration LoxP linear recipient is compared to post-integration linear replaced LoxP cells on various growth media (+1* = chloramphenicol, +2* = tetracycline, +3* = spectinomycin, −1* = *sacB* sucrose sensitivity). **f)** Transformation results for LoxP **d***i* 10-piece assemblies (blue) relative to **d***ii* 20-piece Assembly Amplicon (pink). Plot depicting the transformation accuracy (y axis) vs yield (x axis) in colony forming units per pmol plasmid backbone transformed for 3 biological replicates of the 10-piece scheme (**i**) (Average accuracy=58.6%, Standard deviation=5.1%, Average yield=1.22× 10^3^ Lux + colonies/ µg DNA, Standard deviation=5.61× 10^2^ Lux + colonies/ µg DNA) and the 19-piece Assembly Amplicon scheme (**ii**) (Average accuracy=76.6%, Standard deviation=4.5%, Average yield=3.22× 10^3^ Lux + colonies/ µg DNA, Standard deviation=2.68× 10^2^ Lux + colonies/ µg DNA).

First, cargo DNA and a selection cassette DNA are assembled via HighT (**Fig. 3a i, ii**) to construct an integration plasmid (**Fig. 3a iii**) that is transformed into recipient cells (**Fig. 3a iv**). The recipient contains the recombinase protein and a pre-engineered user defined integration site containing a transcriptionally active fluorescent marker. Upon on target recombination (**Fig. 3a v**), a previously transcriptionally inactive antibiotic selection marker gains an upstream constitutive promoter to transcriptionally activate antibiotic resistance to enable selection (**Fig. 3a vi**). This transcriptional activation of the antibiotic marker is directly coupled to transcriptional inactivation of the fluorescent marker, which functions as an additional verification of correct integrants (**Fig. 3b)**.

Using this system, we demonstrated the ability to directly integrate a 10-piece HighT assembly containing the *Lux* operon to yield transformants of 19.06% Lux+ clones with 1.36 × 10^3^ Lux+ colonies/µg DNA transformed. (**Fig. 3c**). When implementing the HighT Assembly Amplicon, we successfully amplify a 19-piece assembly of the *Lux* cassette resulting in higher yield and accuracy with 30.62% Lux+ clones with 1.43 × 10^4^ Lux+ colonies/µg (**Fig. 3c**). This increase in accuracy and yield is achieved despite the 2x increase in number of assembly fragments. PCRs flanking the genomic location of the insertion confirmed all *Lux* positive cells had integrations at the correct location (**Extended Data 4a**).

We set out to design a parallel system to integrate linear dsDNA (**Fig. 3d**). Input DNA containing flanking recombination sites are derived from HighT ligations (**Fig. 3d i**) or from HighT Assembly Amplicons (**Fig. 3d ii**) and are transformed (**Fig. 3d iii**) into a cell with a Cre recombinase and pre-integrated recombination site (**Fig. 3d iv**). Integration of HighT assemblies is accomplished through two independent crossing over events between distally located recombination sites (L1 and L2), one at each end of the assembly (**Fig. 3d v**). We chose to leverage the well-studied Cre-LoxP integrase system, where one Cre recombinase protein can independently act on two different non-cross reactive orthogonal 34-bp recognition sites, LoxP (L1) and Lox2272 (L2)^39,40^, allowing separate recombination events to occur from the expression of a single recombinase protein.

The gain of an antibiotic marker in the assembly product is directly coupled to the replacement of the genomic region (**Fig. 3d vi**), resulting in a simultaneous loss of a genomically integrated negative selection marker^9^ for additional assurance of on-target integration (**Fig. 3e**). While all clones genotyped showed integration at the correct location (**Extended Data 4b)**, we still demonstrated the ability to use the negative selection marker to counterselection against potential failed integration events (**Extended Data 4c)**.

Using the linear replacement system, we demonstrate integration of both 10-piece HighT direct assembly (58.58% Lux+ with 1.22 × 10^3^ Lux+ colonies/µg DNA) and 20-piece HighT Assembly Amplicon (76.62% Lux+ with 3.18 × 10^3^ Lux+ colonies/µg DNA) (**Fig. 3f**). To demonstrate reproducibility of HighT integrations and demonstrate the ability of both parallel recombinase systems to control insert DNA orientation, we integrated 10-piece assemblies, Assembly Amplicons, and control amplicons into separate recipient cells with mirrored oriented landing sites (**Extended Data 5**).

The use of very high efficiency commercial electrocompetent cells can yield more colonies than in-house made cell which may be advantageous for larger and more complex Assembly Amplicons. To leverage this, we designed a mechanism that begins with first transforming HighT Assembly Amplicons ligated to plasmid backbones containing two separate origins of transfer (OriT) into commercial competent cells, then conjugating them to the linear LoxP integration strain (**Extended Data 6**). A system using additional positive and negative selection markers can be used to select for correct integrants over parent donor and recipient strains.

These data demonstrate that the improvements afforded by the improved 2WJ assembly protocols enable robust integration of assembly product directly into host genomes with no requirement for stable episomal replicon intermediates.

### Cloning eukaryotic coding sequences

HighT can be used to construct definitive DNA sequences to encode medically relevant protein isoforms as an alternative to PCR from cDNA (**Fig. 4a**). One single human gene can produce multiple isoforms through alternative splicing of mRNA with exon compositions differing between tissue type and disease state^41,42^. We selected three proteins whose NCBI references indicate alternative splicing patterns (**Supplement Table 3**): the MAPT gene which creates the tau protein implicated in Alzheimer’s disease^43^, the VEGFA protein which regulates blood vessel formation and whose upregulation is often found in growing tumors^44,45^, and the BRCA1 tumor suppressor gene whose mutation is linked to breast cancer^46^.

**Fig 4.**
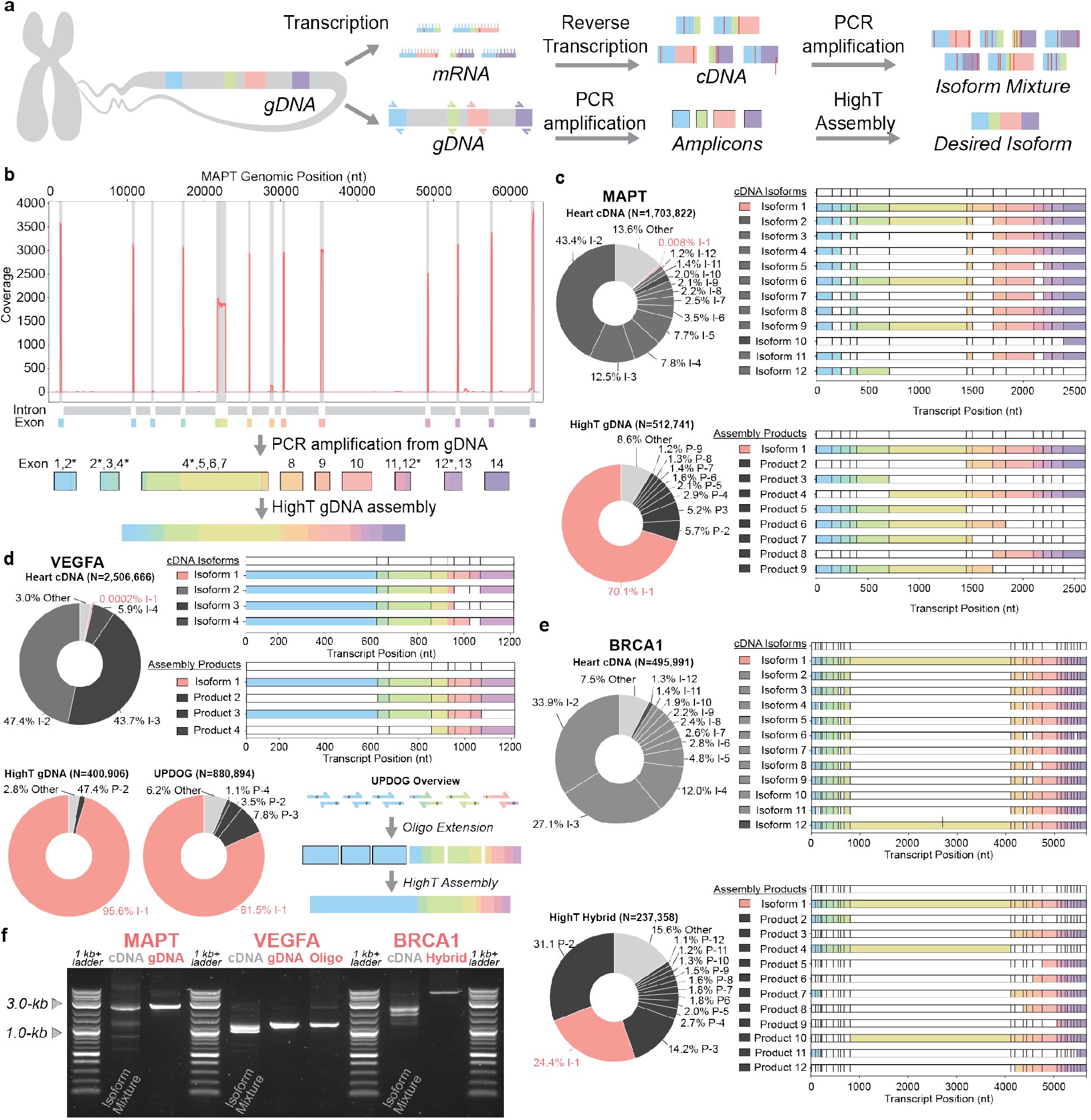
Low abundance human protein isoform variants are constructed with HighT. **a)** Schematic depicting isolation of isoform splicing variants from eukaryotic systems from error prone cDNA or amplified from low error gDNA and assembled with HighT. **b)** Coverage plot of cDNA MAPT amplicon reads aligned to MAPT genomic chromosomal location. **c)** PacBio sequencing to compare isoform variation in amplicons of MAPT generated from human heart cDNA or gDNA HighT Assembly Amplicons. Target isoform denoted in red, all isoforms or sequencing artifacts with >1% representation are given unique identification numbers. **d)** Comparison of isoform variation in amplicons of VEGFA generated from human heart cDNA compared to two different HighT assemblies, one gDNA HighT Assembly Amplicons and the second entirely from oligo based UPDOG & HighT Assembly Amplicon. **e)** Comparison of isoform variation in amplicons of BRCA1 generated from human heart cDNA or a hybrid gDNA amplicon and oligo based HighT assembly. **f)** DNA agarose gel depicting 50 ng of either cDNA amplicon or HighT Assembly Amplicons from **panels c-e** for MAPT, VEGFA, and BRCA1.

While dominant variants of small isoforms can be readily isolated from cDNA amplicons for cloning, low abundance isoforms or longer isoforms which are amplified less preferentially can be more difficult or impractical to isolate. We set out to illustrate that both template DNA in the form of human gDNA as well as synthetically produced oligos could be used in HighT cloning to construct definitive isoforms of human proteins regardless of the natural abundance and length (**Fig. 4a**).

In order to identify long mRNA isoforms from the medically relevant MAPT, VEGFA, and BRCA1 genes, we conducted PCR from cDNA derived from healthy human heart, brain, spleen, skeletal muscle, and testis tissue, with heart yielding most consistent amplification for all target genes. We mapped the heart cDNA reads onto to the genomic chromosomal reference sequence (sample MAPT depiction in **Fig. 4b**) and combined that information with NCBI mRNA reads (**Supplement Table 3**) to determine exon locations (or alternative splicing variant regions) and identify examples of long naturally occurring isoforms for each MAPT, VEGFA, and BRCA1.

For the identified long target isoform of MAPT, the heart cDNA amplicon had extremely low abundance with this isoform composing just 137 reads out of 1,703,822 (0.00804%). In contrast, we successfully constructed the same target isoform of 14 exons, with a 9-piece assembly from amplicons targeting MAPT exons in gDNA (taking advantage longer 5’ ends of amplification primers to encode for some small exons (**Fig. 4b**)), yielding >70% of reads assigned to this isoform by PacBio (**Fig. 4c**) and >95% by Nanopore (**Extended Data 7a**). VEGFA was also constructed using gDNA as template with a 4-piece assembly to compose 9 identified exons, yielding an assembly with >95% (PacBio, **Fig. 4d**) and >90% by Nanopore (**Extended Data 7b)**. The same target isoform in heart cDNA had 6 read out of 2,506,666 reads (0.000239%, **Source Data**).

While smaller exons can be easily incorporated in the 5’ end of the gDNA amplification primer, sometimes long stretches of multiple short exons cannot be encoded in amplification primers, making gDNA amplification impractical in these instances. To address this challenge while still leveraging HighT and the advancements to 2WJ assembly, we designed an oligo-based assembly method Uracil Primer Dimer Overhang Generation (UPDOG). In isolated PCR reactions, overlapping primers are extended to make short, synthetic 2WJ assembly fragments. Then by processing with our improved 2WJ protocol and assembly with HighT, UPDOG is able to generate an entirely synthetic product (**Fig. 4d**). VEGFA constructed entirely with UPDOG yields ∼80% correctly assembled constructs with PacBio and ∼70% with Nanopore, all with no gDNA template employed (**Fig. 4d**).

Combining both gDNA amplicons and UPDOG assembly in a hybrid manner can utilize the advantages of both techniques. Employing gDNA amplification for longer exons and synthetic UPDOG cloning for shorter exons enables a streamlined protocol for uniformly processing fragments to construct low abundance isoforms. To synthesize the target isoform of BRCA1, we designed a 13-fragment assembly to construct a 25-exon splicing BRCA1 variant. This hybrid assembly was able to yield 24% correct constructs as analyzed by PacBio sequencing (**Fig. 4e**) and over 75% correct analyzed by Nanopore (**Extended Data 7c**); no natural amplicons were found in the 495,991 heart cDNA read for this long isoform.

Relative to cDNA amplifications, HighT has qualitatively cleaner DNA products for the construction of all 3 target isoforms of MAPT, VEGFA, and BRCA1 while enabling control over exactly which isoform is derived (**Fig. 4f**). We additionally used the per-base accuracy of PacBio sequencing to measure the SNP rate in the cDNA amplicons and our HighT constructs (**Extended Data 8**) but the reported SNP frequency of cDNA^47–50^ and gDNA^51–51^ are below the per-molecule detection limit for PacBio sequencing.

### *In vitro* assembly of large and complex DNA

*De novo* construction of large DNA products remains a premium commodity that is not commonly utilized in routine research applications^53,54^. We believe this perceived barrier limits the scope of research which requires large synthetic DNA. *De novo* construction of DNA sequences has been historically limited by the accuracy and reliability of the assembly techniques used to stitch the chemically synthesized synthetic DNA oligos together^2^. Beyond this, construction of large DNA products usually requires an *in vivo* intermediate step where shorter DNA assemblies are conducted and subcloned into bacteria^17,55–57^ to physically isolate clones with correct constructs from a population with misassembled products (**Extended data 9**). The *in vivo* intermediates ultimately increase cost and time owing to limited throughput and difficulty of automation. Further, there are instances, particularly in pharmaceutical applications, where cell-free DNA is required due to concerns regarding byproducts such as endotoxins that persist after an *in vivo* subcloning step^58^. As a result, it would be advantageous for there to be an assembly technique or combination of techniques which were reliable enough to produce large DNA products (10 kb and beyond) *de novo* and entirely *in vitro*.

Towards this end, we combined our 3WJ assembly technique, Sidewinder, with our 2WJ HighT assembly for the high-fidelity construction of a complex, linear, 10 kb, dsDNA product. To produce 10 kb entirely *de novo* and cell-free, we first use Sidewinder to produce ten 1-kb DNA products starting from synthetic oligos composing Sidewinder fragments. The 3WJ assembly intermediates are then amplified with dU containing primers, simultaneously removing the 3WJ and incorporating a dU residue into the assembled products. These products are then processed and assembled with HighT ligation to a final size of 10 kb, optionally amplified into an Assembly Amplicon (**Fig. 5a**).

**Fig 5.**
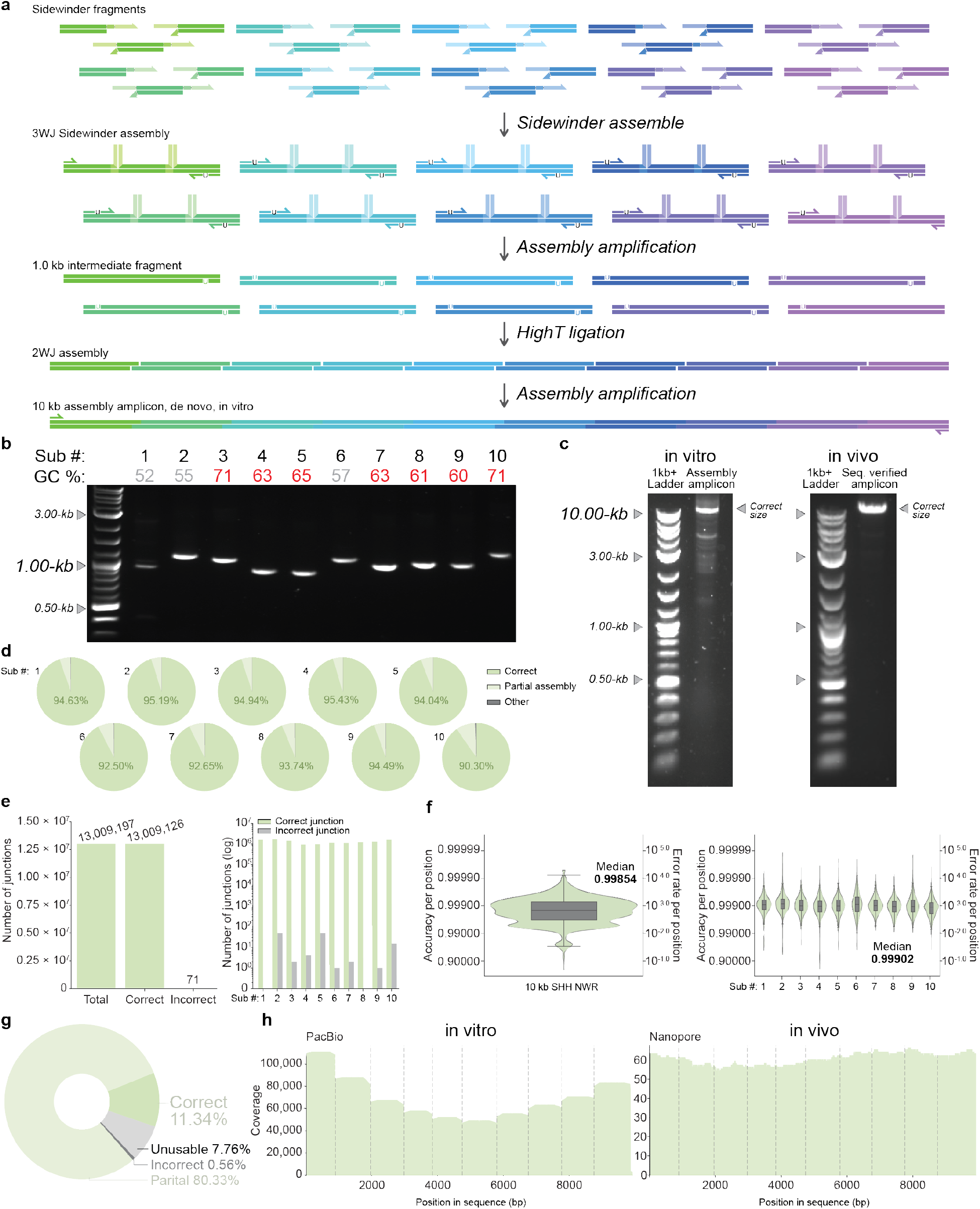
High fidelity assembly enables *in vitro* hierarchical assembly of large and complex linear DNA constructs. **a)** Schematic depicting the *de novo* assembly pipeline integrating Sidewinder with HighT 2WJ assembly for *in vitro* hierarchical assembly of large constructs. **b)** DNA agarose gel depicting 50 ng of the ten 1 kb Sidewinder assemblies post PCR amplification with dU incorporation. **c)** DNA agarose gel depicting 50 ng of the 10 kb HighT assembly post PCR amplification with *in vitro* hierarchical assembly (left) and a comparative PCR from the *in vivo* assembled confirmed SNP free template (right). **d)** Fragment level PacBio sequencing analysis each of the primary assembly step using Sidewinder. Reads are analyzed considering the connections between Sidewinder fragments composing the each of the 1 kb sub-assemblies with correct products (green), partially aligned products (subset of fragments in the correct order) (light green), and PCR or sequencing artifacts or incorrect assemblies (grey). **e)** Junction level PacBio sequencing analysis from unfiltered data counting all observed Sidewinder junctions for each of the ten 1 kb amplicons depicting total junctions, correctly ligated junctions, and incorrectly ligated junctions with the global count (left) and the count by sequence (right). **f)** Nucleotide level PacBio sequencing analysis as a violin plot and nested box and whisker plot depicting the distribution of per-base accuracies for each position for each of reads containing the full 10 kb assembly (left) and all reads aligned to each of the ten 1 kb segments. Summary statistics available in **Source Data**. **g)** Fragment level PacBio sequencing analysis the secondary assembly step using HighT. Reads are analyzed considering the connections between the ten HighT fragments with correct products (green), partially aligned products (subset of fragments in the correct order) (light green), and PCR or sequencing artifacts or incorrect assemblies (grey). **h)** Coverage plots of the 10 kb amplicons for the *in vitro* hierarchical assembly sequenced with PacBio (left) and the *in vivo* assembled confirmed SNP free template.

Our target for the *de novo* cell-free construction was the genomic sequence and upstream context for the *Sonic Hedgehog* (*SHH*) gene of the northern white rhinoceros. The northern white rhino is a functionally extinct species with only two females remaining^59^ and *SHH* plays an important role in embryonic development^60^. In addition to the large genomic gene size, it also is a very complex sequence for DNA construction as it has regions which are incredibly GC rich, with multiple 1 kb regions with over 60% GC content, 50 bp regions with as high as 94% GC content, as well as independent stretches of 11 and 15 consecutive guanine bases (**Fig. 5b, Extended data 10**).

Sidewinder was used to *de novo* construct each of the 1 kb intermediates and successfully produces a strong, target amplicon of the expected size for all 10 intermediates despite the high GC content (**Fig. 5b**). Due to the 10^-5^ to 10^-6^ misconnection rate of Sidewinder^3^, these *in vitro* intermediates can be directly processed for a second round of assembly according to our HighT protocol and subsequently amplified for a 10 kb Assembly Amplicon. The high GC content and large size of this assembly makes this a difficult amplification even when PCR amplifying from a perfect template^61^ yet despite these extreme conditions, with optimized PCR protocols using DMSO to overcome long GC stretches (**Methods**), the PCR of the final 10 kb assembly shows a strong target amplicon of the expected size (**Fig. 5c**).

To quantify the assembly accuracy, PacBio sequencing was done on the 10 kb final product for single molecule high-fidelity reads^62^. At the primary assembly step with Sidewinder, fragment level analysis (measuring that each Sidewinder fragment appears in the correct order) shows consistently high rates for correctly assembled constructs for each of the 10 sub assemblies (each fragment in the correct order) ranging from 90.3% correct (sub assembly 10) to as high as 95.43% correct (sub assembly 4) (**Fig. 5d**). The next major products seen are partial assemblies (a subset of fragments in the correct order), with PCR artifacts, sequencing artifacts, or incorrect assemblies making up the small proportion of remaining reads (**Fig. 5d**). To more specifically assess the connection rate at the 3WJ, we applied our junction analysis pipeline to identify all instances of ligated 3WJs that could result from either correct or incorrect ligation^3^. In total, out of 13,009,197 total observed junctions, we observed just 71 mis-ligated junctions for a misconnection rate of just 1 in 183,228 (**Fig. 5e**). For 2 of the ten 1 kb intermediates, we observed 0 mis-ligated junctions out of 2,794,355 connections while the remaining 8 sequences had junction accuracies between 99.9951% and 99.9999% (**Fig. 5e**). Characterizing the assembly at the nucleotide level to leverage the per-base accuracy of the PacBio sequencing, the median per-base accuracy for reads containing the full 10 kb assembly is 0.99854 whereas reads aligned to any of the 10 sub assemblies have a median accuracy of 0.99902 or approximately 1 SNP in 1,020 bases (**Fig. 5f**).

When characterizing the HighT assembly of the ten 1-kb fragments into the final 10-kb Assembly Amplicon, the PacBio sequencing shows 11.34% of product being the full length 10 kb Assembly Amplicon (**Fig. 5g, h**) with the majority of reads being partial assemblies (a subset of the assembly fragments connected in the correct order). These data conflict with the prominent band of the correct size on the agarose gel and the Nanopore sequencing data which observed 90.2% full length product (**Extended data 7d**).

We did not observe any SNP free clones in the PacBio dataset due to low coverage, unreliability in sequencing across homopolymeric runs (**Extended data 10**) and the observed PacBio sequencing error rate of ∼10^-3.5^ lowering the probability of accurately assigning each of the 10,000 bases in a single molecule even if it were SNP free (**Extended data 8**). Depending on the tolerance to potential SNPs in the final construct, the *in vitro* Sidewinder and HighT pipeline offers an avenue for more rapidly obtaining large synthetic DNA constructs for various applications. To obtain a nucleotide perfect construct, we cloned the 1 kb Sidewinder products *in vivo* and confirmed mutation-free sequences, verified with whole plasmid Nanopore sequencing and pair-end Sanger Sequencing for homopolymeric regions (**Extended data 10**). PCR amplicons from identified nucleotide perfect 1 kb intermediates were assembled with HighT into the final 10 kb target construct (**Fig. 5c**) similarly verified for nucleotide perfection.

## Discussion

With the discovery of Sidewinder as an accurate methodology for DNA assembly using the 3WJ^3^, we set out to characterize a series of improvements which could be implemented to prior arts based on the standard 2WJ connection between molecules. In this study, we conceptualized and demonstrated a novel paradigm for the assembly of DNA molecules with 2WJ, HighT assembly, that offers a conceptual advancement over conventional protocols through the demonstration of accurate and robust assemblies at reaction temperatures significantly above the T_m-50_ of ssDNA assembly overhangs.

We defined a mechanistic explanation for our hypothesis that expands on the concept of T_m_ to account for smaller fractions of association <50% where infrequent molecular interactions at higher temperatures still occur often and stably enough for successful ligation. By Le Chatelier’s principle, these infrequent ligations events cause the reaction to tend towards completion over time as species are irreversibly removed from the equilibrium between of associated ⇌ unassociated duplex. We demonstrated how this confers increased reaction specificity via the globally destabilizing reaction conditions favoring association of perfectly matched molecules.

The resultant HighT methodology, paired with additional improvements to the design and processing of assembly overhangs, greatly improves assembly accuracy and yield to enable more reliable construction of DNA molecules via the 2WJ. The fundamental principles of these advancements may be applied to the abundance of existing 2WJ assembly techniques, offering more reliable construction with pre-established workflows. We show multi-fragment assemblies with as high as 96.7% correctly assembled products and demonstrate particular advantage when HighT assembly is applied to PCR-based multi-fragment assemblies, breaking new barriers for USER assembly reaction sizes with up to 20 fragments assembled simultaneously, a 40-point increase in accuracy, and over four orders of magnitude increase in yield over prior arts.

Further, the greatly enhanced fidelity at the 2WJ increases assembly accuracy such that PCR may be conducted on the assembled product. This conceptual advancement of the “Assembly Amplicon” was leveraged in a variety of context to greatly enhance assembly yield, specifically for large multi-fragment assemblies. Assembly Amplicons also allow for rapid hierarchical assembly of DNA products entirely *in vitro* and cell free. The streamlined *in vitro* protocols afforded by the Assembly Amplicon can decrease costs, time to construction, enhance automation, and reduce potential concerns with regard to cellular endotoxins which may be present after *in vivo* cloning. The Assembly Amplicon is limited by the mutation frequency of the starting material but offers a high throughput avenue for workflows which are less sensitive to mutations.

Throughout these demonstrations, a variety of temperatures are utilized for HighT. In each case, HighT confers high accuracy and ease of assembly for multi-fragment products and does not necessarily need to be defined by a single elevated temperature. Conceptually at ultra-low temperatures, prefect matches and mismatches across assembly junctions universally associate strongly and at ultra-high temperatures, prefect matches and mismatches across assembly junctions universally associate poorly. In both cases, the relative stability or instability of the reverse complement hydrogen bonding pairs has limited differentiation between intended and unintended pairs across assembly junction. Thus, the optimal HighT temperature where there is the greatest differentiation across the assembly junction will depend necessarily on the length of overhangs, the overhang sequences being used, the choice of ligase, buffer conditions, DNA concentrations and other variables. In light of this, this work demonstrated 50°C as a reliable HighT temperature as a proof of principle. Future work will be dedicated to predicting and optimizing assembly-specific HighT conditions.

The advancements described with HighT assembly and Assembly Amplicons demonstrate new workflows that may ease the studying and engineering of biological systems. We show robust integration of assembly product directly into host bacterial genomes with no requirement for stable episomal replicon intermediates using two independent integration schemes. This can streamline genome engineering by saving on the time and costs associated with sequencing plasmid intermediates prior to genomic integration, thereby enabling more efficient protocol automation. We further demonstrate that reliable HighT assembly facilitates the generation of important eukaryotic genes and specific isoforms. Through the combined approaches of gDNA amplification and synthetic construction, we may more reliably isolate isoforms of interest via HighT construction when rare mRNA isoforms are impractical to isolate from cDNA amplification.

In addition to more reliably isolating the correct isoform, gDNA as source material offers additional advantages due to fundamental limitations of amplification from cDNA. The abundance of single nucleotide point mutations (SNPs) in cDNA results from the high error rate of transcription and subsequent reverse transcription processes. The error rate of transcription is reported to be 10^-4^-10^-6^ per bp^47,48,74^, reverse transcription to be 10^-4^-10^-5^ per bp^49,50^, and the error rate of eukaryotic genomic DNA replication with proofreading and error correction at 10^-9^-10^-10^ per bp^51,52^. As such, using patient derived genomic DNA as a low SNP-rate template for PCR amplification of specific exons which are subsequently combined via high fidelity HighT ligation offers a potential advantage in both aspects over conventional methods. Comparing the SNP rate our gDNA assemblies with the SNP rate of a control plasmid shows our error rate is equal to or lower than the detection rate of PacBio, indicating further investigation would be needed to determine the true mutation frequency when conducting assembly sourced from different DNA inputs.

Finally, HighT paired with the state-of-the-art assembly technique from oligos, Sidewinder, enables the *de novo* and cell-free construction of a complex 10 kb target sequence. The interfacing of the two techniques enables the construction of large, linear, dsDNA rapidly and entirely *in vitro.* The lack of *in vivo* cloning both in the intermediate and final construction steps opens new opportunities for use of large and complex linear dsDNA molecules. Depending on downstream applications, the error rate of input oligos may ultimately limit the final size of the *de novo* Assembly Amplicon so more work must be done towards reducing error of input DNA. The demonstration of the *SHH* gene of the northern white rhino is pertinent, as the species is functionally extinct and historically, many biological tools that have been crucial to conducting research and making therapeutics have been sourced from existing genetic components found from living organisms such as CRISPR/Cas9 technologies^63^, transposons^64^, and therapeutics like biologics^65^ to name a few. With the capacity for producing large genomic regions *de novo* we may expand our repertoire of sequences to include historic genes that are no longer accessible by PCR amplification and leverage these sequences to find new and useful tools and therapeutics. Additionally, improvements to AI design will expand our sequence space even further to include DNA sequences which have never existed, adding to the need for rapid *de novo* construction.

HighT offers a distinct set of advantages that can be applied to 2WJ assembly techniques that can be leveraged for the more efficient engineering of biological systems.

## Acknowledgements

K.W. acknowledges funding support from the Shurl and Kay Curci Foundation, NIH (1DP2GM140937) and Caltech Center for Environmental Microbial Interactions (CEMI).

## Author contributions

N.E.R., B.G., W.Z., M.W., D.G. and J.Y. performed DNA assembly experiments; H.Z., N.R., and G.A. developed informatic pipelines for the analysis of sequencing data with input from B.G. and K.W.; M.W., J.H., N.E.R. and B.G. performed data analysis. All of the authors discussed the results. N.E.R., B.G., M.W., and K.W. wrote the manuscript with input from all of the authors.

## Methods

### Simulated 10 bp overhang melt curves

From the set of all possible 10 bp overhangs (4^10^ = 1,048,576), a subset was chosen for simulated melt curves by removing sequences with strong secondary structures and homodimers. This was accomplished using NUPACK^24,25^ nupack-v4.0.1.12 with strand_conc=1e-8, material=“dna”, temp_c=50, sodium=0.150, magnesium=0.01, and max_complex_size=2. Each of the 1,048,576 sequences were screened individually using a test tube analysis job using the *tube* analysis function. The 10 base sequences which both sense and antisense strands had 90% unpaired bases at the defined parameters were accepted for further analysis. This accounts for intramolecular secondary structures such as hairpins and unintended homodimer duplexes which may interfere with intended duplex formation in the simulated melt curves.

The resultant list of 401,933 unique duplexes, composed of strand *a* and reverse complement strand *b*, were screened using a test tube analysis job using the *tube* analysis function with strand_conc=1e-8, material=“dna”, sodium=0.150, magnesium=0.01, and max_complex_size=2 for each whole number temperature 4°C to 85°C. Then the calculated concentration of Complex (a+b) was treated as the duplex concentration for each temperature point. Owing to the simulation being conducted over discrete temperatures, for a given value of T_m-n_ the first temperature which has the concentration of Complex (a+b) fall below the proportional *n* value is used as the T_m-n_. For example, T_m-50_ is determined by the first temperature where [Complex (a+b)] ≤ 0.5e-8.

### Oligo Purchasing

Amplification primers and assembly primers containing dU residues were ordered from Integrated DNA Technologies and Millipore-Sigma with standard desalt purification and at the lowest mass offered, usually 25 nmol. Oligos for generating the oligo-based assembly fragments were purchased from Millipore-Sigma with Standard DNA Synthesis for DNA Oligos in Tubes which has a maximum oligo length of 120 bases, ordered with standard desalt purification at 50 nmol. The 10 kb SHH Sidewinder oligos which were shipped in water at 50 µM. All other oligos were shipped dry. All oligos are listed in **Supplement Table 1**.

### qPCR ligation assay

The dsDNA holder complex was annealed in 50 µL 1X Binding and Wash buffer (5 mM Tris-HCl pH 7.4, 0.5 mM EDTA, 1 M NaCl) with 2 pmol biotinylated oligo and 1 pmol complement with initial denaturation of 95°C for 3 min followed by −0.1°C per second to 22°C. The streptavidin coated tubes (High Capacity Streptavidin Coated PCR Tubes, Biomat: PCRSTDF-SB75/100) were washed with 200 µL 1X Binding and Wash buffer and were incubated with the annealed dsDNA duplex for 1 hr at 22°C followed by two washes with 200 µL 1X Binding and Wash buffer.

Ligation mixtures were prepared with 1 µL ligase per 100µL reaction with 0.1 pmol (for 4 base experiment) or 0.4 pmol (for 10 base experiment) ssDNA oligo with either 1X T4 DNA ligase buffer with T4 ligase (New England Biolabs), 1X Hifi Taq ligase buffer with Taq DNA Ligase (New England Biolabs), or only 1X Hifi Taq ligase buffer (control). For the respective conditions, 100 µL was added to the coated tube and directly incubated for 16 hr at 16°C, 22°C, 30°C, or 50°C. After the incubation, the reaction mixture was removed and the tubes were immediately washed with 200 µL of 1X Binding and Wash buffer and then washed two times with 200 µL 20 mM NaOH with 5 min incubation after each addition, followed by a pH neutralization with 200 µL of 1X Binding and Wash buffer.

The qPCR was then conducted according to manufacturer’s instructions with 20 pmol each forward and reverse primer using the qPCR mix (SsoAdvanced Universal SYBR Green Supermix 2X, Biorad: 1725271) on the BioRad CFX Opus 96 Real-Time PCR System. PCR conditions were 98°C 3 min, 98°C 10 sec, 56°C 10 sec, 60°C 10 sec, return to step 2 35X, melt curve analysis 65°C to 95°C at 0.5°C increment at 5 sec per step, hold 4°C.

### Fragment Design

The 4-piece GFP assemblies in **Fig 2** were designed systematically as to reduce bias that may come from specifically chosen overhangs.

For the negative control, the first overhang selected was the first valid sequence of the form AN_x_WN_y_WN_z_T (where one W is an A and the other a T) with a T_m-50_ of 30°C at the end of the GFP coding sequence. This ensured that the overhang could be compatible with a two dU condition and that the entire GFP gene could be a single permissible fragment. The plasmid was then divided into four equal length fragments. The location for the next overhang was chosen to be the first valid overhang of the form AN_x_WN_y_WN_z_T with a T_m-50_ at 30°C after moving an equidistant 917 bases from the terminal T of the previous overhang. This was repeated until all 4 overhangs were chosen. Through this process, the Rep101-GFP junction has substantial crosstalk with itself and the GFP-KanR junction has a looped secondary structure as determined by the NUPACK web application^24,25^ at 30°C, 1µM.

For the no secondary structure (NSS) assembly, all overhangs were chosen to be of the form AN_x_WN_y_WN_z_T with a T_m-50_ of 30°C and were chosen to have reduced secondary structure and minimal cross reactivity (less than 13% concentration defect at 1 µM) at 30°C. The first junction was created after the coding sequence of GFP using pre-generated sequences^66^ because the junction flanking the GFP gene is not present in a coding sequence. Subsequent junctions were again chosen by moving an equidistance from the previous overhang and choosing the first valid junction that satisfied the NSS conditions. All secondary structure and cross reactivity of chosen overhangs were checked using NUPACK web application.

For the two dU condition, the position of the second dU was chosen which most evenly reduces the melting temperature of both resultant waste strands as to maximize the difference in T_m-50_ between the waste strands and the T_m-50_ of the intended assembly partner.

All primers used to amplify DNA fragments are listed in **Supplement Table 1**. All final constructs and templates can be found in FASTA format in **Supplement Table 2**.

### PCR amplification and purification

The polymerase used for amplification of input PCR fragments or final amplification of Assembly Amplicons are listed in **Extended Methods**. PCRs are conducted in 50µL reactions.

The backbone fragments for the GFP construct assemblies were amplified from 2.5 ng of a KanR, pSC101, SacB template. The GFP gene was amplified from 2.5ng of a CmR, p15a, mGL, SacB template. SacB was utilized in the templates such that sucrose selection could be used to select against any residual template in the final transformation. The backbone fragment for the Lux construct assemblies was amplified from 2.5 ng of a CmR, p15a, mGL, SacB template such that GFP could be used to identify, and sucrose selection could be used to select against, any residual template in the final transformation. The fragments of the Lux cassette were amplified from 2.5 ng of a TetR, BAC, *LuxABCDE* template. Sequences for the templates are listed in **Supplement Table 2**.

Post PCR amplification of assembly fragments, purification was done using a QIAquick PCR Purification Kit (Qiagen). Multiple 50uL PCR reactions can be passed simultaneously through the same purification column to increase the final concentration of the purified amplicon. Purified products are eluted in 10% elution buffer.

### Fragment processing

The conditions used to process and assemble each construct demonstrated in this paper are listed in **Extended Methods**.

All assembly fragments sourced from bacterial plasmid templates are treated with DpnI (New England Biolabs) for 1hr at 37°C with a 20 min heat inactivation at 80°C in a 50 µL reaction of 1x CutSmart using 1 µL DpnI per 50 µL of initial PCR reaction. This digestion occurs after PCR purification of the initial fragments and prior to assembly. Conditions which perform USER digestion prior to assembly had the USER enzyme mix added during this step. All USER assemblies were treated with 1µL USER enzyme mix (New England Biolabs) per 40pmol dU (or minimally 1µL if total moles less than 40 pmol) and digested for at least 1 hr at 37°C.

An additional purification was conducted after this digestion using either QIAquick PCR Purification Kit (Qiagen), Monarch DNA Gel Extraction Kit (New England Biolabs), or Quantum Prep PCR Kleen Spin Column (Biorad) (**Extended Methods**). Kleen column purification had the temperature of the sample raised to 50°C immediately after digestion prior to passing through the column and was the method used for the **Fig 2g δ**. Purified products were then measured by Qubit 1x dsDNA High Sensitivity Assay Kit, (Invitrogen, ThermoFisher Scientific) and stored at −20°C until assembly.

Oligos to compose Sidewinder fragments were shipped at 50 µM in water and processed based on this reported concentration. The amount of coding oligo from the stock resuspension required for a 2uM final concentration in 25uL would be individually phosphorylated in a 22.5uL reaction using 1uL of T4PNK (New England Biolabs) in 1xT4 ligase buffer at 37°C for 1hr, followed by an enzyme deactivation at 80°C for 10 minutes. The corresponding volume of stock barcode oligo needed for 1uM in 25uL was then added to the mix for a total volume of 25 µL.

Heteroduplexes were annealed in PCR tubes on a thermocycler with an initial denaturation of 98°C for 10 minutes, followed by a gradual decrease in temperature down to 25°C at −1°C per minute. Once fragments are annealed, they are kept at 4°C until use. Fragments were then purified via gel extraction from an 8% TBE-PAGE gel (Invitrogen, ThermoFisher Scientific) and run at 200v for 35 minutes, extracted via razor blade, and eluted through osmosis into the assembly buffer for the respective conditions^3^.

### Fragment assembly

Processed fragments were assembled under one of the following thermocycler conditions (**Extended Methods**)

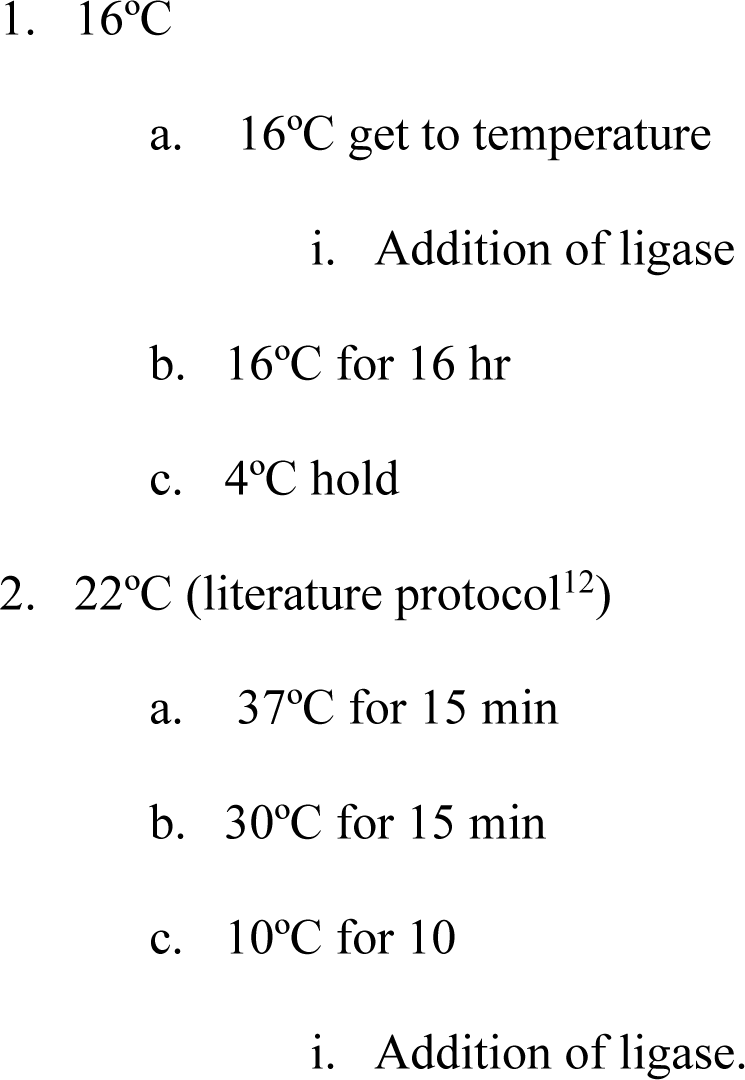
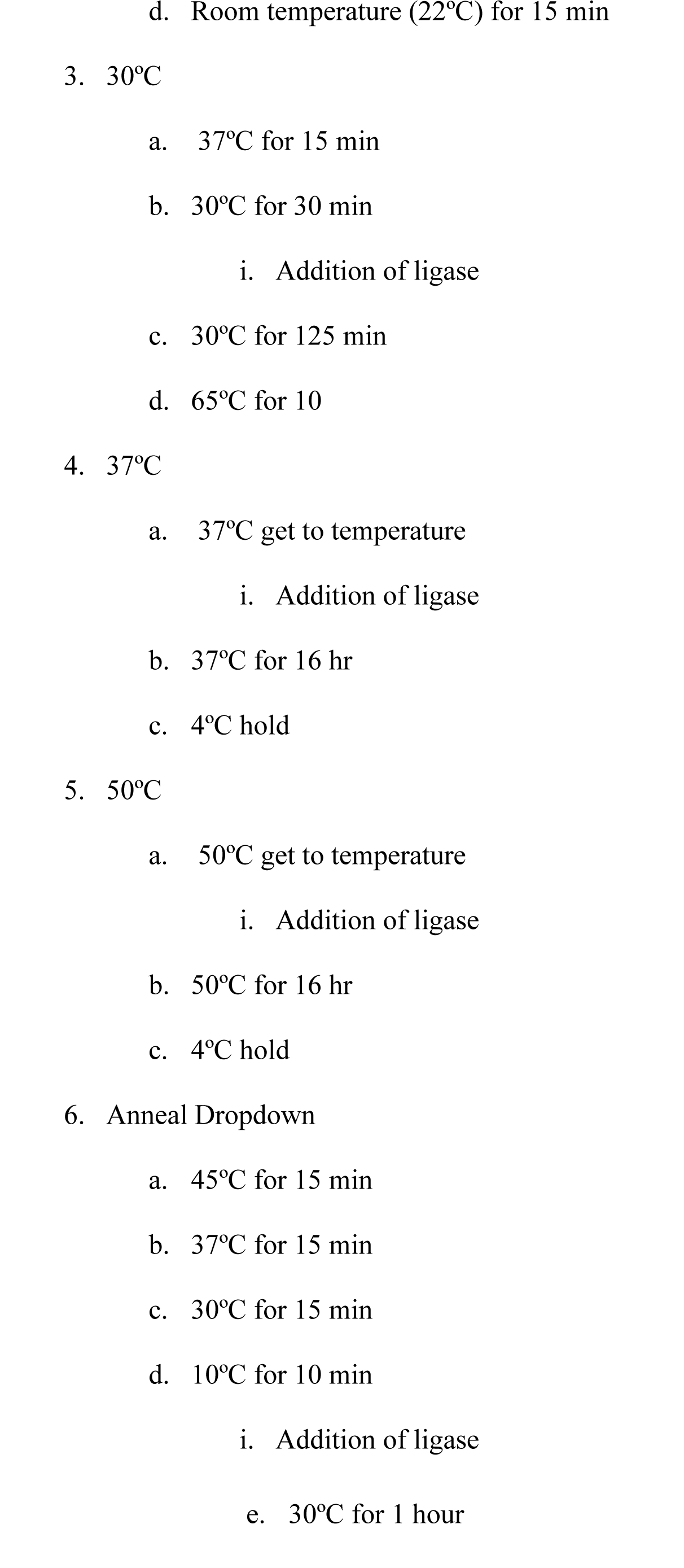

### Human cDNA and gDNA assemblies

Amplicon libraries were generated by select amplification of MAPT, VEGFA, and BRCA1 using designed primers (**Supplement Table 1**). PCR Ready Frist Strand cDNA from healthy human tissues were purchased from BioChain. The cDNA from human heart, brain, spleen, skeletal muscle, and testis tissue were tested, with heart providing most consistent expression with isoform diversity across all 3 gene targets.

Exon fragments for MAPT, VEGFA, and BRCA1 were amplified similarly but from BioChain Human Heart gDNA (**Supplement Table 1**).

### Sidewinder 1 kb sub-assemblies

The Sidewinder assemblies for 1 kb SHH sub-assemblies were conducted at between 1-3 nM in 50 µL 1x Hifi Taq Buffer (New England Biolabs). We utilized two thermocycler conditions for assembly of the Sidewinder 1 kb sub-assemblies. Assemblies were conducted in 50uL reactions in 1× HiFi Taq buffer (New England Biolabs). For sub-assemblies S1, S2, S3, S4, S5, S6, S9, and S10, we used a thermocycler protocol with 85°C for 5 minutes followed by the addition of 2uL of Taq ligase (New England Biolabs), then a slow decrease of −1°C per min down to 37°C, incubate at 37°C for 20 minutes, and repeat the thermocycling 85°C-37°C three additional times. For sub-assemblies S7, and S8, the “cycling” protocol of 85°C for 5 minutes, followed by the addition of 2uL of Taq ligase, then the reaction then cycles between 85°C for 1 minute and 50°C for 2 minutes for 100 cycles. These cycles are then followed by 50°C for 1 hr.

PCR is then conducted on the Sidewinder assemblies with 7.5% volume DMSO in 50µL in reactions of 25 µL RepliQa HiFi ToughMix (Quantabio), 1.5 µL 10µM primers, 3.75 µL molecular biology grade DMSO (Sigma-Aldrich), 17.25 µL H_2_O, and 1 µL Sidewinder assembly as template. PCR reactions are conducted using the manufacturers recommended protocol for 3-step PCR amplification or using a published “slowdown” protocol for amplifying GC rich sequences^67,68^.

### SHH 10 kb assemblies

For *in vitro* assembly, 1 kb Sidewinder sub-assemblies S3, S4, S7, S8, S9, and S10 were amplified from Sidewinder product amplicons, and S1, S2, S5, and S6 were amplified directly from Sidewinder product using RepliQa HiFi ToughMix (Quantabio) as described in PCR amplification and purification. Amplicons were purified using a QIAquick PCR Purification Kit (Qiagen), eluted in 40 µL of 10% Elution Buffer, and digested in a 50 µL reaction containing 1 µL USER enzyme (New England Biolabs), 5 µL 10× CutSmart buffer, and 5 µL H₂O. Reactions were incubated at 37°C for 1 hour followed by 80°C for 20 minutes. Fragments were then gel extracted using the Monarch DNA Gel Extraction Kit (New England Biolabs). Concentrations of each sub-assembly were measured by Qubit 1× dsDNA High Sensitivity Assay Kit (Invitrogen, ThermoFisher Scientific). Assemblies were set up with 0.4 pmol of each fragment in 50 µL 1x Hifi Taq Buffer (New England Biolabs) and assembled using the 50 °C protocol.

The resulting 10 kb SHH assembly product was amplified with PrimeSTAR GXL Polymerase (Takara Bio) at 7.5% DMSO. PCR reactions are conducted using the manufacturers recommended protocol for 3-step PCR amplification with 63°C annealing and 68°C extension for 10 min.

For *in vivo* assembly, all 1 kb Sidewinder sub-assemblies were amplified from Sidewinder product amplicons. Sub-assemblies were purified using QIAquick, digested with USER enzyme as described for the *in vitro* assembly, purified again with QIAquick, and quantified by Qubit. Each 1 kb fragment was assembled into the pSC101 backbone using the 50°C assembly condition and transformed described.

All *in vivo* assemblies were validated by Nanopore sequencing through the GENEWIZ service. Sub-assemblies S1, S2, S4, and S5, which contain homopolymers >8 bp, were also sent for Sanger sequencing to confirm these regions. Sequence-validated fragments were amplified from colony resuspension, purified, digested, gel extracted, and assembled with 0.4 pmol of each construct as described for the *in vitro* assembly. The final 10 kb SHH product was assembled into the BAC backbone, transformed, and sequenced to identify SNP-free clones.

### Sequencing Analysis

The coverage plots of 22°C, 30°C and 50°C assembly amplicon were generated sequencing product of standard PCR purification with Oxford Nanopore. Reads (FASTQ) are aligned to the 6.1 kb *LuxABCDE* casette reference sequence using minimap2 (v2.30) ^69,70^ with the map-ont preset, and alignments were streamed directly to SAMtools (v1.6)^71^ for coordinate sorting and indexing. Per-base coverage was calculated with SAMtools depth -aa to include zero-coverage positions and coverage values were parsed into contig-length–matched arrays. Coverage profiles were concatenated across contigs and plotted as read depth versus reference position.

The fragment connection analysis for the 22°C, 30°C and 50°C assembly amplicon was done by generating a list of all valid connections between all fragments accounting for 5’ and 3’ orientation and overhang supersession. Using these ∼650 base references, Oxford Nanopore reads (FASTQ) were aligned using minimap2 (-x map-ont, -k 15, -w 10, -N 150, --secondary=yes, -p 0.6, 8 threads) and alignments were filtered by requiring coverage of ≥0.75× the specific reference length (absolute floor 250 bp) and ≥0.82 identity (MAPQ not thresholded). To prevent strand-mirroring double counts, passing hits were deduplicated per read by collapsing alignments with the same junction assignment and similar read coordinates using 25 bp–binned query start/end intervals. Each retained hit was collapsed into one of three connection classes where the left (L) and right (R) end of two fragments are connected in sense-sense orientation (RL), sense-antisense orientation (RR*), or sense-antisense orientation (LL*). Raw counts can be found in (**Source data**).

Both Oxford Nanopore Sequencing and PacBio was done for amplicons from human cDNA, HighT isoform assemblies, and the northern white rhino SHH 10 kb final product.

Nanopore sequencing reads of cDNA PCR products were aligned to gDNA reference sequences using minimap2 and further processed with SAMtools to generate a coverage plot showing the exons to be selected. For PCR products of the assembly reaction of the selected exons, plasmidsaurus Premium sequencing service was used for analysis. FASTQ reads that are shorter than 10% of the desired isoform or that do not contain both PCR primers were discarded since those are likely sequencing artifacts. Remaining reads were aligned against the desired isoform sequence using the blastn-short program of BLASTN. Alignments with more than 70% of query coverage and more than 70% of percent identity were considered valid hits. Constructed isoforms were identified based on the order of the fragments in the alignment. Isoforms identified were present at >1% of filtered reads, with all reads <1% binned into Other.

For the PacBio reads, for SNP profiling: reads were aligned to reference sequences using the Smith-Waterman local alignment algorithm^72^ with a match score of 5, mismatch penalty of 4, gap open penalty of 10, and gap extend penalty of 0.5. Only reads with a mean Phred quality score of greater than or equal to Q39.5 were retained. At the base level, only positions with a Phred score of greater than or equal to Q40 were included to minimize sequencing errors. Reads were further filtered to require greater than or equal to 90% alignment identity and greater than or equal to 80% aligned fraction of the reference length. The SNP rate at each position was calculated as the total number of non-reference events (substitutions, deletions, and insertions) divided by total coverage at that position. Homopolymeric runs have the total insertions and deletions distributed evenly across the stretch of the run to account for discrepancies in assigning insertions and deletions to one of a particular position in a homopolymeric run.

Fragment-level analysis: read sequences were aligned to all fragment references using BLASTN. Reads returning no hits to any fragment were classified as unusable. A complete assembly was assigned when all fragments were detected in the correct order of the gene sequence; assemblies with correct connections between fragments but missing a subset of fragments with no misconnections were classified as partial. Reads containing fragments joined in an incorrect order were classified as “other” including PCR and sequencing artifacts or incorrect assemblies. Fragment hits were required to cover at least 80% of the fragment reference length.

Sidewinder Junction analysis: a junction was defined as 25 base pairs flanking both the 3’ and 5’ ends of each Sidewinder junction. For each sample, all possible junctions, including both correct ligations and potential misligations, were generated from the fragment references and aligned to the raw reads using BLASTN with sensitive parameters. Junction hits were filtered using query coverage (greater than or equal to 80%) and percent identity (greater than or equal to 80%) thresholds. For reads identified as potential misligations, they were counted as true misligations if a read contained a seamless junction the exact 6 base pairs on either side of the ligation point between non-partnered fragments.

Homopolymers in northern white rhino assemblies validated through pair end Sanger sequencing using GENEWIZ Sanger sequencing service.

### Agarose gel imaging

All gel images are 1-2% agarose gels stained with Sybr Safe (Invitrogen, ThermoFisher Scientific) run at 135V for 25 minutes in 0.5x TBE buffer with the 1 kb+ ladder (New England Biolabs). The 19-piece LuxABCDE assembly gel depicts 1 µL of final PCR loaded onto the gel. Sidewinder assembly gel depicts 50ng of sample loaded onto the gel. The 10 kb sample depicts 5 µL final PCR loaded onto gel.

### Transformation

Outside of attBP and LoxP tranformations (below section), all transformations were done using NEB 10-beta electrocompetent E. coli cells (New England Biolabs). Assemblies were processed as described. Cells were transformed using 2 µL of assembly product into 25 µL electrocompetent cells and transferred to an electroporation cuvette and pulsed at 2500V. Then 2 mL NEB 10-beta Stable Outgrowth Media (New England Biolabs) is added to the cuvette, and entire volume is transferred to a 15 mL tube for a 1-hour recovery at 37°C with shaking.

A dilution series is then established assuming 2 mL final volume by adding 22.2 µL of the n-1 to 200 µL 10% Luria-Bertani (LB) to make the nth dilution. All 200 µL are spread on an LB-agar plate using glass beads. Antibiotic concentrations and negative marker concentrations used in this study, unless specifically otherwise noted, are: Carbenicillin 100 µg/mL, Chloramphenicol 20 µg/mL, Kanamycin 50 µg/mL, Spectinomycin 60 µg/mL, Streptomycin 100 µg/mL, Tetracycline 10 µg/mL, Sucrose 7.5% w/v, Chlorinated Phenylalanine (p-chloro-DL-phenylalanine) 5 mM, Hygromycin 200 µg/mL, Apramycin 200 µg/mL. Plates are then incubated for 24-48 hours until all clones could be clearly distinguished.

### Strain engineering and electrocompetency

For LambdaRed recombination, Cre-LoxP recombination, or attBP recombination, electrocompetent cells were prepared similarly. Cells containing recombination machinery were grown overnight in 10 mL of LB plus respective antibiotics. OD was measured at 600 nm using NanoDrop One^C^. The next day cells were diluted to OD = 0.05 in 500-1000 mL of LB and antibiotics and incubated at 37C with 170 rpm shaking. Cultures were grown until OD = 0.2, where Arabinose was added to final 0.5% concentration to induce recombination machinery. At OD = 0.5, cells were placed on ice for 30 minutes. Cultures were spun down at 4C for 10 minutes at 4000 rpm and washed with pre-chilled water, then again with pre-chilled 15% glycerol. Cells were spun down and resuspended in 750 µL of 15% glycerol before flash frozen in liquid nitrogen and stored at 80C.

For the attBP and LoxP recipient cells, MDS42 ^73^ was used as the parental strain and LambdaRed recombination was first used to knockout RecA and the native HK022 attB sites by integrating a *chlor-sacB* selection cassette before patching out with a 120 bp primer to generate a scarless deletion (DNA in **Supplement Table 2**). A PCR amplicon of the attP or 2LoxP landing site was recombined in at the *dif* site. Strains were cured of the LambdaRed plasmid before having the pSC101-HK022 (AmpR) or pSC101-Cre (TetR) plasmids introduced. These final strains were made induced and electrocompetent as described above for integration experiments.

### Transformation and plating attBP and LoxP ligations

50-100 ng of purified ligations were transformed into electrocompetent cells and recovered for 4 hours before plating. attBP transformants were serially diluted on plates containing chloramphenicol, Kanamycin, and carbenicillin. LoxP transformants were plated on just chloramphenicol. Transformations were performed in triplicate, from independent ligations reactions. Cells were restreaked on same selection plates before genotyping (**Extended Data 4**).

### Phenotyping and imaging

For phenotypic spots, colonies were grown overnight, OD normalized to 0.2, then 5 µL of cells were spotted on agar plates containing selective media.

Imaging of colorimetric, luminescence and GFP was performed using BioRad ChemiDoc Imaging System with following settings: Chemiluminescent Blot 30s (white tray), Colorimetric Blot 0.5s (white tray), GFP on SYBR Green 0.5s (blue tray).

### Statistics

All reported assemblies in **Fig 2f, 2g, 3c, 3f** figures were done with three biological replicates. Each sample was an independent assembly reaction that was transformed a single time to get respective colony counts reported in **Source Data**. Data presentation is composed of the mean and standard deviation of the biological replicates.

## Data availability

The sequencing data generated in this study have been deposited in the NCBI Sequence Read Archive with the submission number PRJNA1464736. Source data are provided for this paper.

## Extended data

**Extended Data 1.**
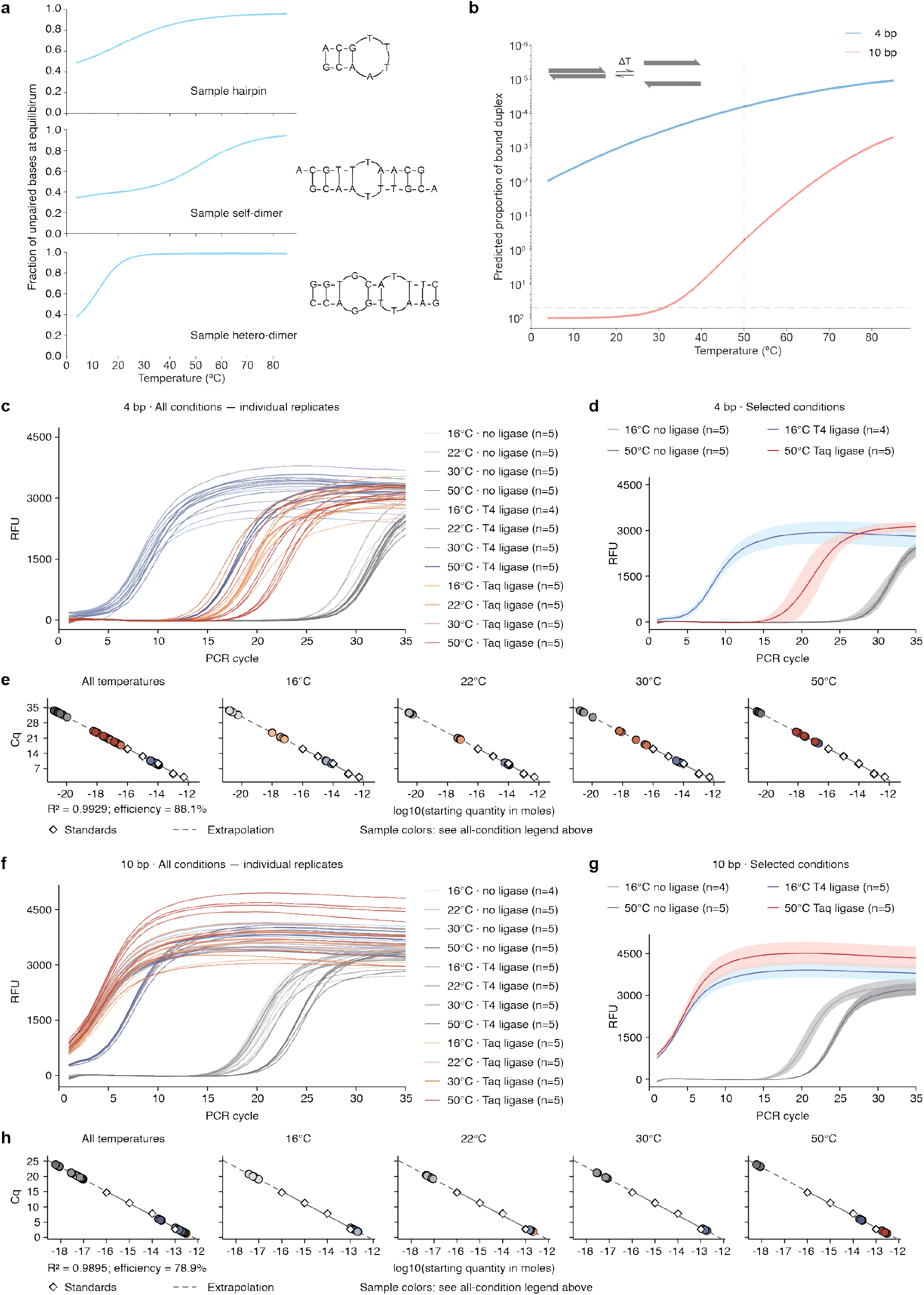
Raw data for qPCR showing effective ligation for overhangs above their T_m-50_. **a)** NUPACK simulated melt curves for DNA with secondary structures. **b)** NUPACK simulated melting curves for 4 bp and 10 bp overhangs used in the overhang ligation qPCR experiment at discrete temperatures 4°C to 85°C. **c)** qPCR amplification curves depicting changes in fluorescence over PCR cycles for all individual replicates of the 4 bp overhang ligation reactions at 16°C, 22°C, 30°C, and 50°C with no ligase, T4 DNA Ligase, or Taq DNA Ligase. **d)** qPCR amplification curves depicting mean fluorescence ± SD over PCR cycles for the 4 bp overhang ligation reactions at conditions 16°C with no ligase, 16°C with T4 DNA Ligase, 50°C with no ligase, and 50°C with Taq DNA Ligase. **e)** qPCR standard curves for 4 bp overhang ligation reactions, showing Cq as a function of log10 starting quantity across all temperatures and each temperature individually. Dashed lines indicate extrapolation beyond the standard range. **f)** qPCR amplification curves depicting changes in fluorescence over PCR cycles for all individual replicates of the 10 bp overhang ligation reactions at 16°C, 22°C, 30°C, and 50°C with no ligase, T4 DNA Ligase, or Taq DNA Ligase. **g)** qPCR amplification curves depicting mean fluorescence ± SD over PCR cycles for the 10 bp overhang ligation reactions at conditions 16°C with no ligase, 16°C with T4 DNA Ligase, 50°C with no ligase, and 50°C with Taq DNA Ligase. **h)** qPCR standard curves for 10 bp overhang ligation reactions, showing Cq as a function of log10 starting quantity across all temperatures and each temperature individually. Dashed lines indicate extrapolation beyond the standard range.

**Extended Data 2.**
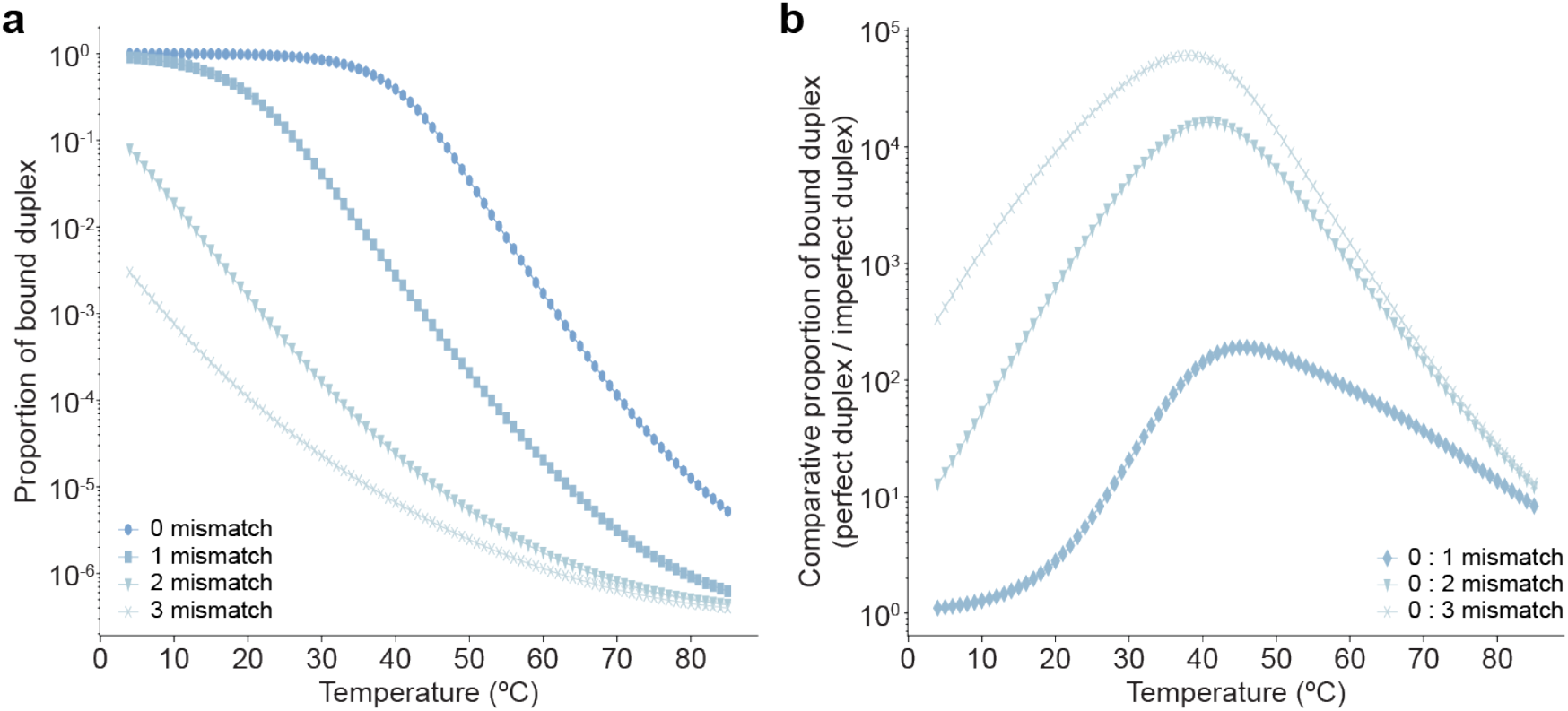
HighT has greatest differentiation at a sequence-dependent optimal temperature between ultra-low and ultra-high reaction temperatures. a) Plot depicting a NUPACK simulation of the proportion of species in the bound duplex for 10 bp ssDNA oligos over a range of temperatures for a perfect duplex (0 mismatch) and increasing number of mismatched bases. b) Plot depicting a NUPACK simulation of the ratio between the proportion of species in the bound duplex state for a perfect duplex (0 mismatch) divided by the proportion of species in the bound duplex state for the 1, 2, and 3 mismatched duplex over a range of temperatures for 10 bp ssDNA oligos.

**Extended Data 3.**
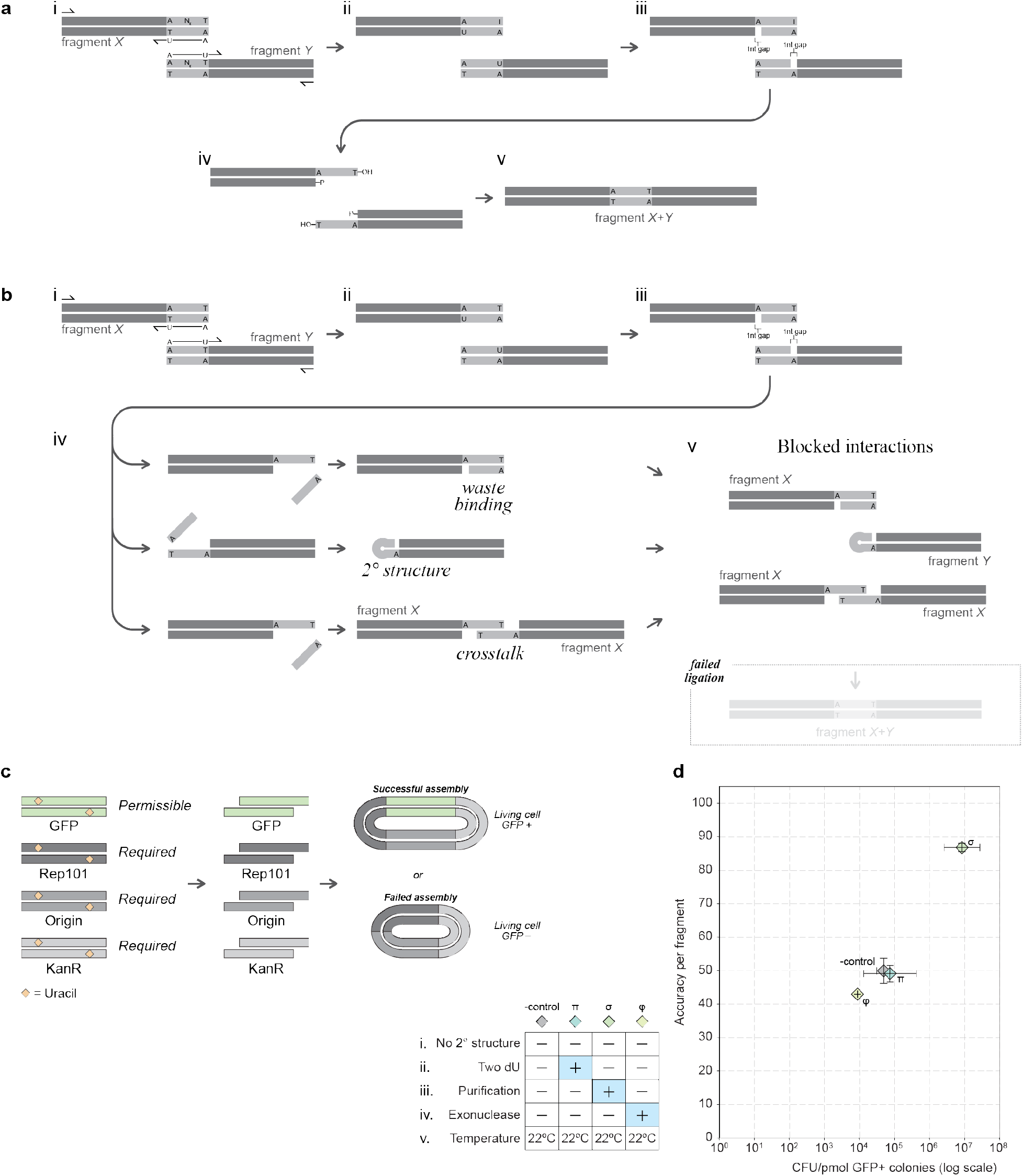
Improvements to USER protocol are substantially burdened by secondary structure and crosstalk in overhangs. **a)** USER assembly generates overhangs by amplifying a target sequence with a terminal sequence of the form AN_x_T with a dU in place of the dT in the primers. The dU is targeted with USER enzyme cocktail to generate the 2WJ overhang for assembly. **b)** Unoptimized protocols can result in various byproducts which decrease the overall fidelity of assembly. **c)** Schematic depicting the possible transformant genotypes after a 4-piece GFP assembly. The one permissible fragment encoding GFP does not correlate with cell survival while 3 required fragments encode for the backbone and are required for transformant survival on antibiotic selection. **d)** Plot depicting the transformation accuracy (y axis) vs yield (x axis) in colony forming units per pmol plasmid backbone transformed for the 4-piece GFP assembly conducted using overhangs not optimized for secondary structure at 22°C literature protocol under published conditions (grey, **-control**), second digest site (blue, **π**), purification (green, **σ**), and exonuclease digestion (yellow, **φ**).

**Extended Data 4.**
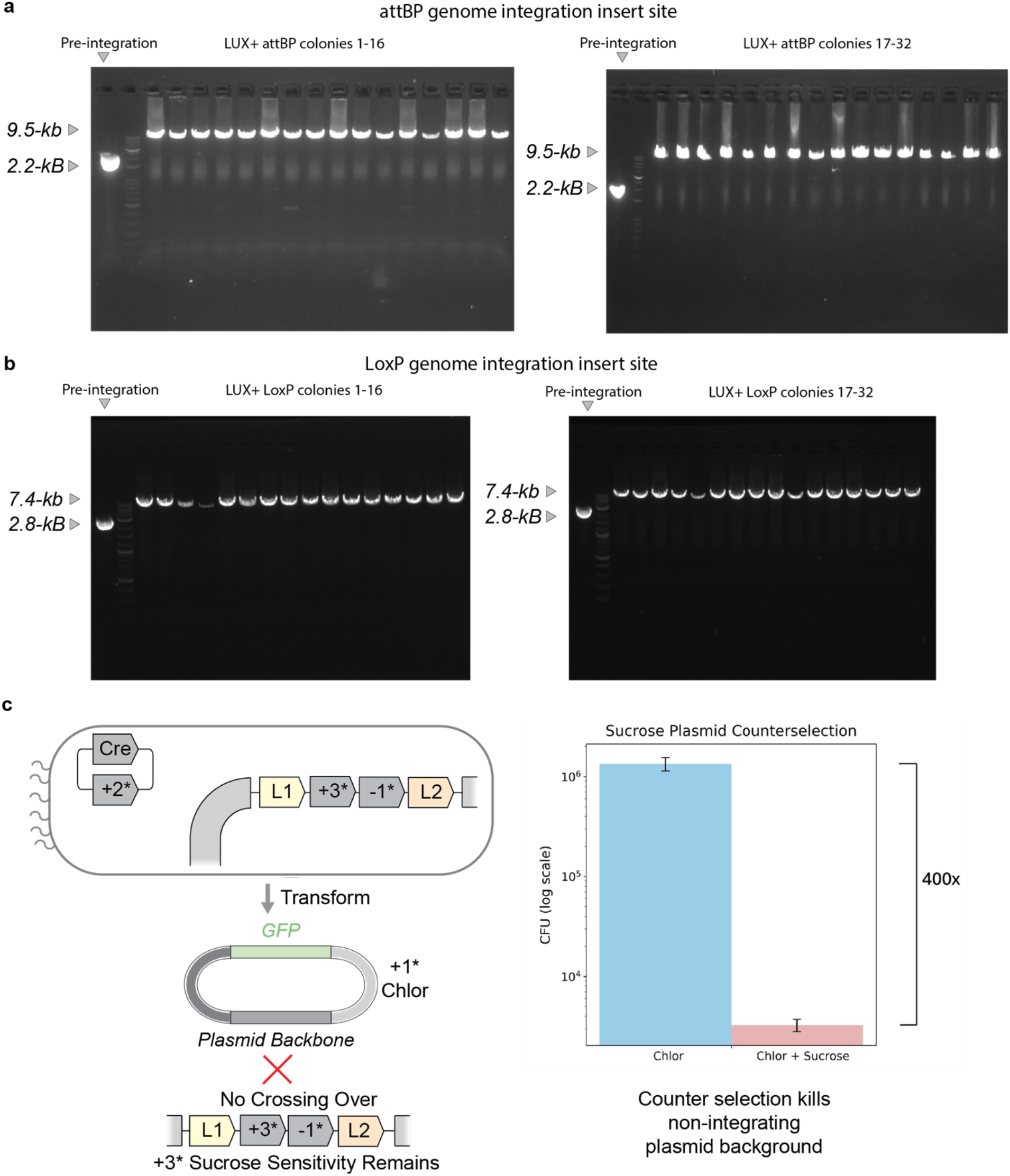
attBP and LoxP integrations occur at expected recipient genomic location. **a)** PCR check with primers binding to genomic region flanking expected insert site near *dif* location in genome. Size of genomic insert increases from pre-integration 2.2 kB to post-integration 9.5 kB for attBP integration when checking 32 *Lux*+ colonies. **b)** Same genomic site comparing LoxP pre-integration 2.8 kB amplicon with post-integration 7.4 B amplicon for 32 *Lux*+ colonies. **c)** Reduction in colonies surviving when selecting against non-integrating plasmids. GFP+ chlor plasmids are transformed into LoxP pre-integration recipient strains containing the *sacB* negative selection gene. As the recipient genomic *sacB* locus is not lost with plasmid transformation, sucrose counterselection can be used to reduce episomal transformants by ∼400x.

**Extended Data 5.**
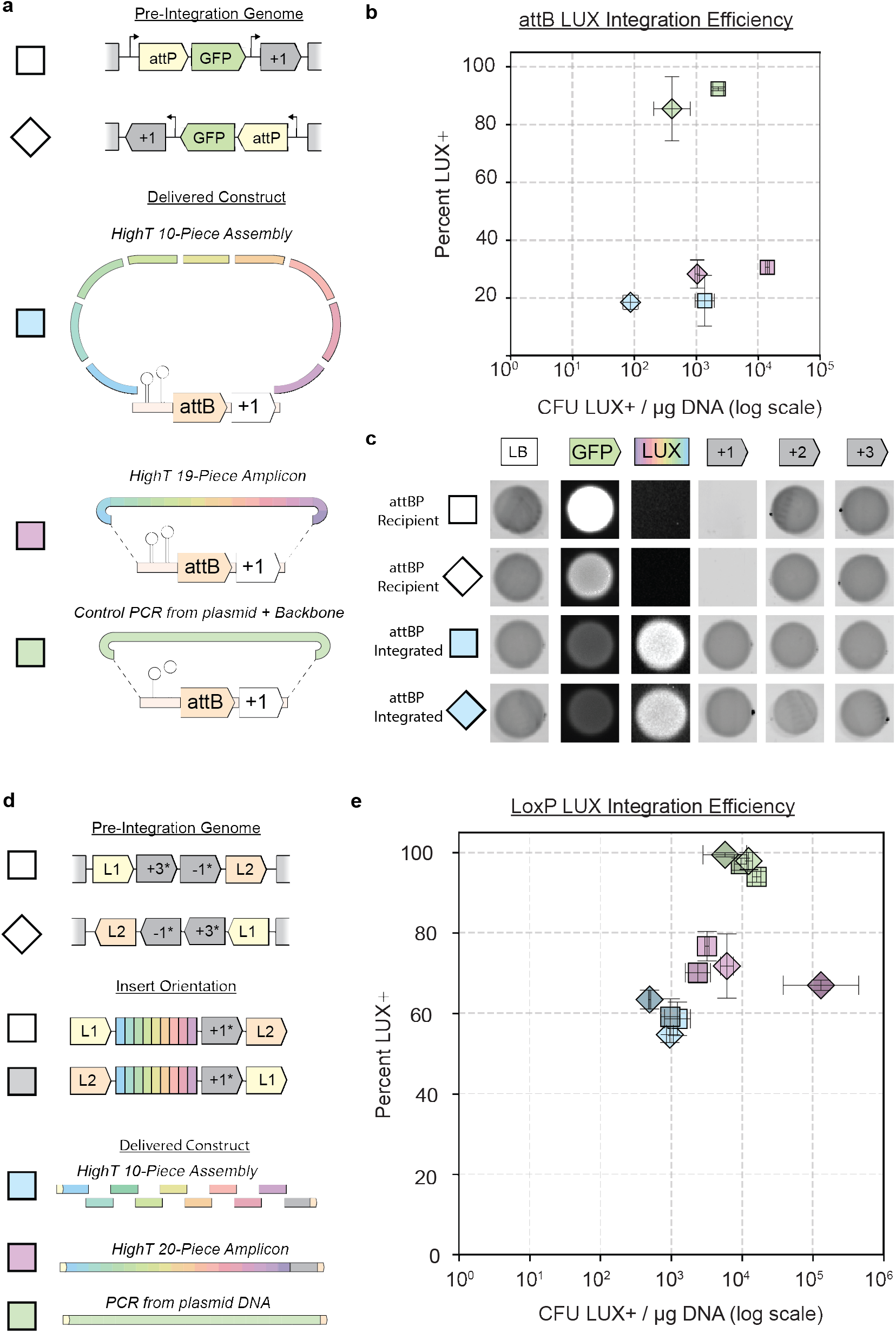
attBP and LoxP integration schemes can integrate with different orientations. **a)** Orientation of inserted genomic attP landing spot, in two different directions (square and triangle). Delivered constructs are either HighT 10-piece assemblies (blue), 19-piece assembly amplicon ligated to backbone (pink), or PCR product from sequence verified template ligated to backbone (green). **b)** Integration efficiencies of the three constructs integrated into two different recipient cell orientations. **c)** Phenotypic spots of pre-integration attBP Recipients and post-integration cells with recipient landing sight oriented in two different directions. Resistance gene: +1 = Kanamycin, +2 = Carbenicillin, +3 = Chloramphenicol. **d)** Cells with two different recipient LoxP/Lox2272 landing site orientations are transformed with products where the relative orientation of the distal LoxP = L1 and Lox2272 = L2 are swapped. Delivered constructs are either HighT 10-piece assemblies (blue), 20-piece assembly amplicon (pink), or PCR product from sequence verified template (green) flanked by L1 or L2 at distal ends. **e)** Integration efficiencies of the three different constructs sources, each tested independently with the two L1/L2 or L2/L1 orientations, and separately integrated in two different recipient cell orientations.

**Extended Data 6.**
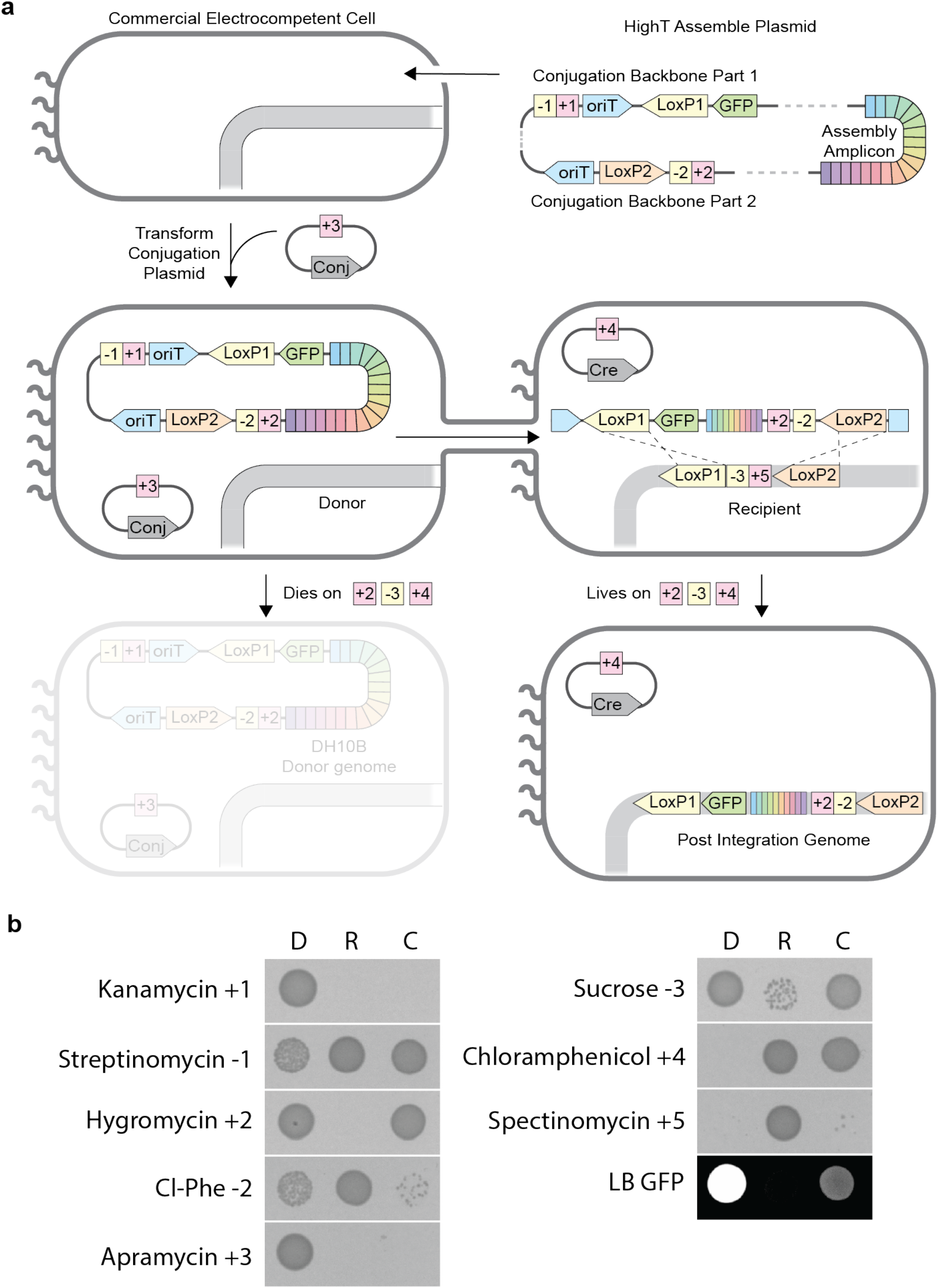
HighT assembled constructs can be conjugated to recipient cells. **a)** Assembly Amplicon is HighT assembled to two oriT containing conjugation backbones. This assembly is transformed into high efficiency commercial electrocompetent cells. A conjugation plasmid is then added to make this cell a Donor. Conjugation between the two oriT sites on the donor plasmid is performed and delivered into the same Recipient as in Fig. 2d. After integrating the conjugated DNA, the Chimeric result has a unique phenotype that enables positive and negative selection to kill off both parents, Donor and Recipient. **b)** Phenotypic results of 5 µL of OD_600_ = 0.2 cells from Donor conjugator (D), Recipient integrator (R), and Chimeric result (C) spotted on selective agar. Selection markers and concentrations: +1 kanamycin Resistance (100 µg/mL), −1 streptinomycin Resistance (300 µg/mL), +2 Hygromycin Resistance (200 µg/mL), −2 Cl-Phe Sensitivity (5 mM p-chloro-DL-phenylalanine), +3 Apramycin Resistance (200 µg/mL), −3 Sucrose Sensitivity (7.5% w/v), +4 Chloramphenicol Resistance (20 µg/mL), and +5 Spectinomycin Resistance (60 µg/mL). GFP was taken for cells grown on LB only plates.

**Extended Data 7.**
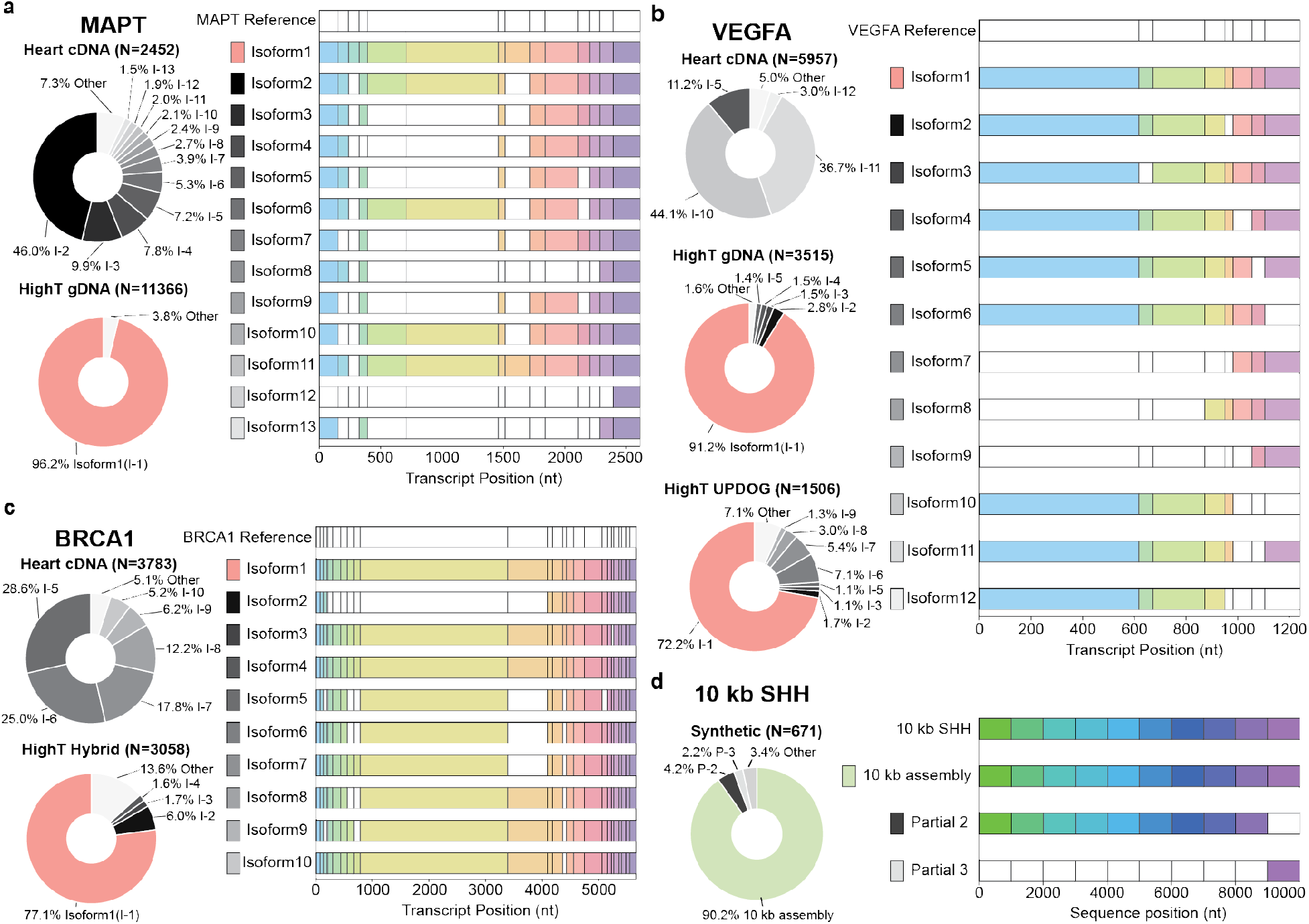
Nanopore sequencing of cDNA amplicons and HighT Assembly products shows high proportion of desired product through HighT assembly. Nanopore sequencing analysis depicting products represented at greater than 1% of the total reads and their corresponding exon or fragment pattern for a) MAPT, b) VEGFA c) BRCA1, and d) 10 kb northern white rhino Sonic Hedgehog (SHH).

**Extended Data 8.**
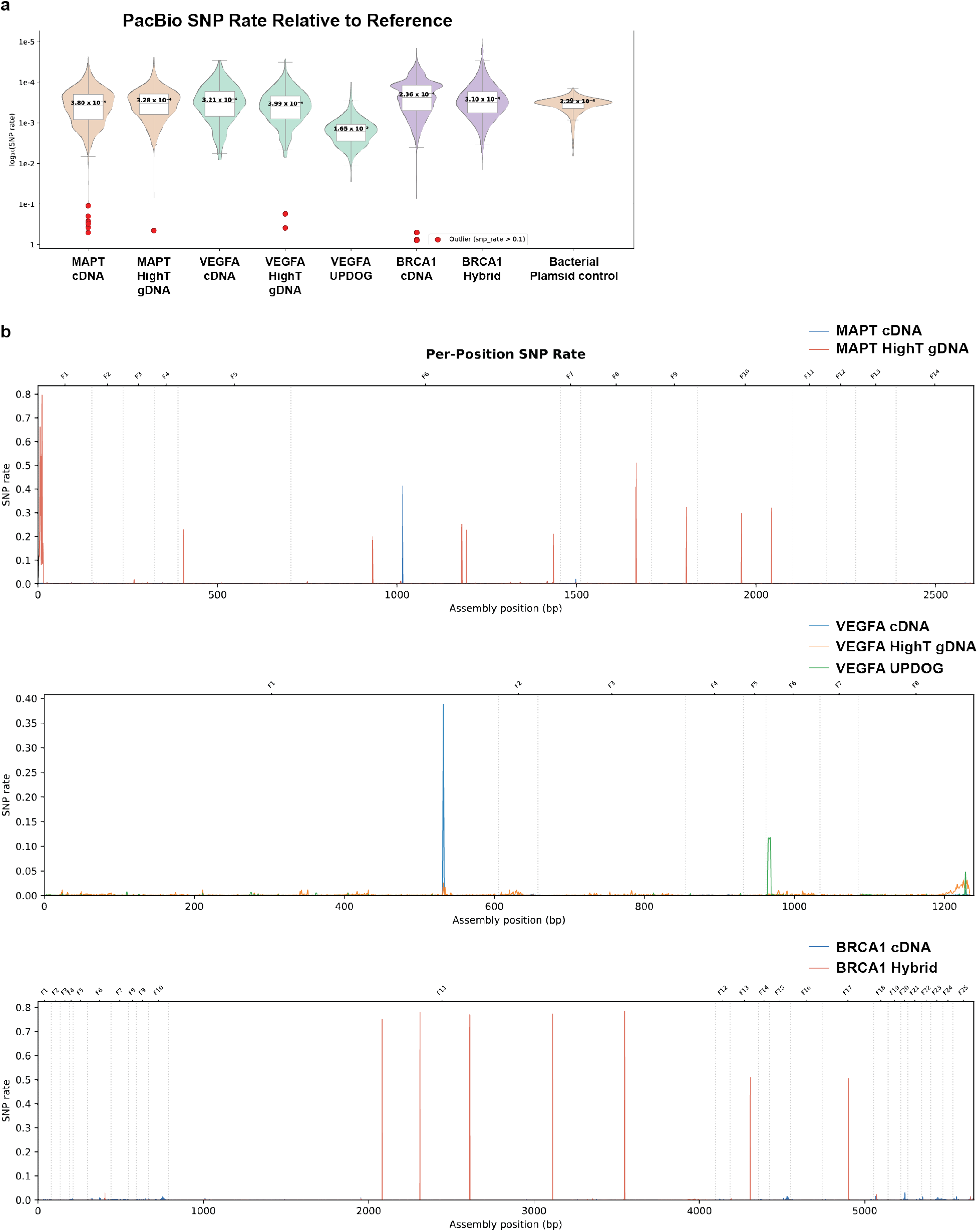
SNP profile of PacBio Reads. **a)** SNP rates for each assembly relative to reference. Accuracy at each nucleotide for every PacBio read was counted. Positions with rate >10% are considered allelic diversity, potentially due to naturally occurring variation in patient derived samples. Summary statistics are available in **Source data. b)** SNP rate per position for the three target genes showing location of specific SNPs.

**Extended Data 9.**
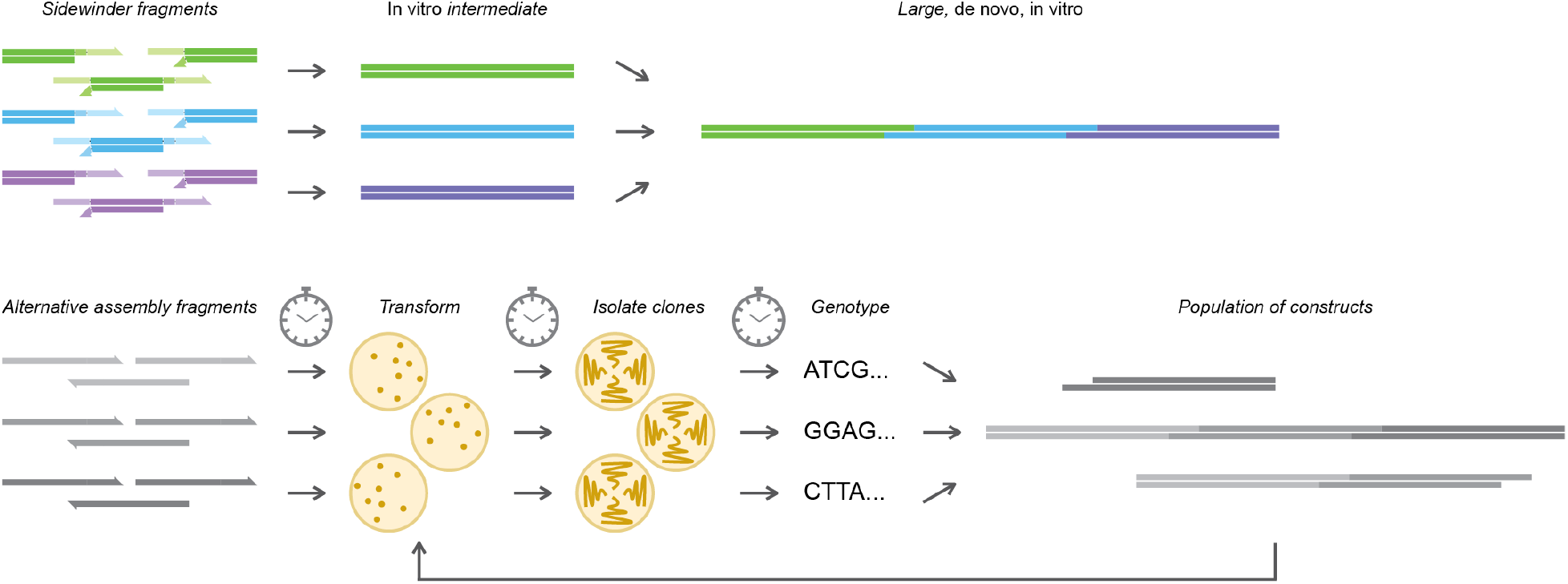
Increased assembly fidelity enables more rapid *in vitro* assembly protocols. Sidewinder enables a high fidelity for the proper connection that allows for highly pure *in vitro* intermediates to be generated and carried directly through for secondary rounds of hierarchical cloning. Alternative assembly techniques require an *in vivo* transformation in order to isolate correctly assembled clones prior to subsequent rounds of assembly.

**Extended Data 10.**
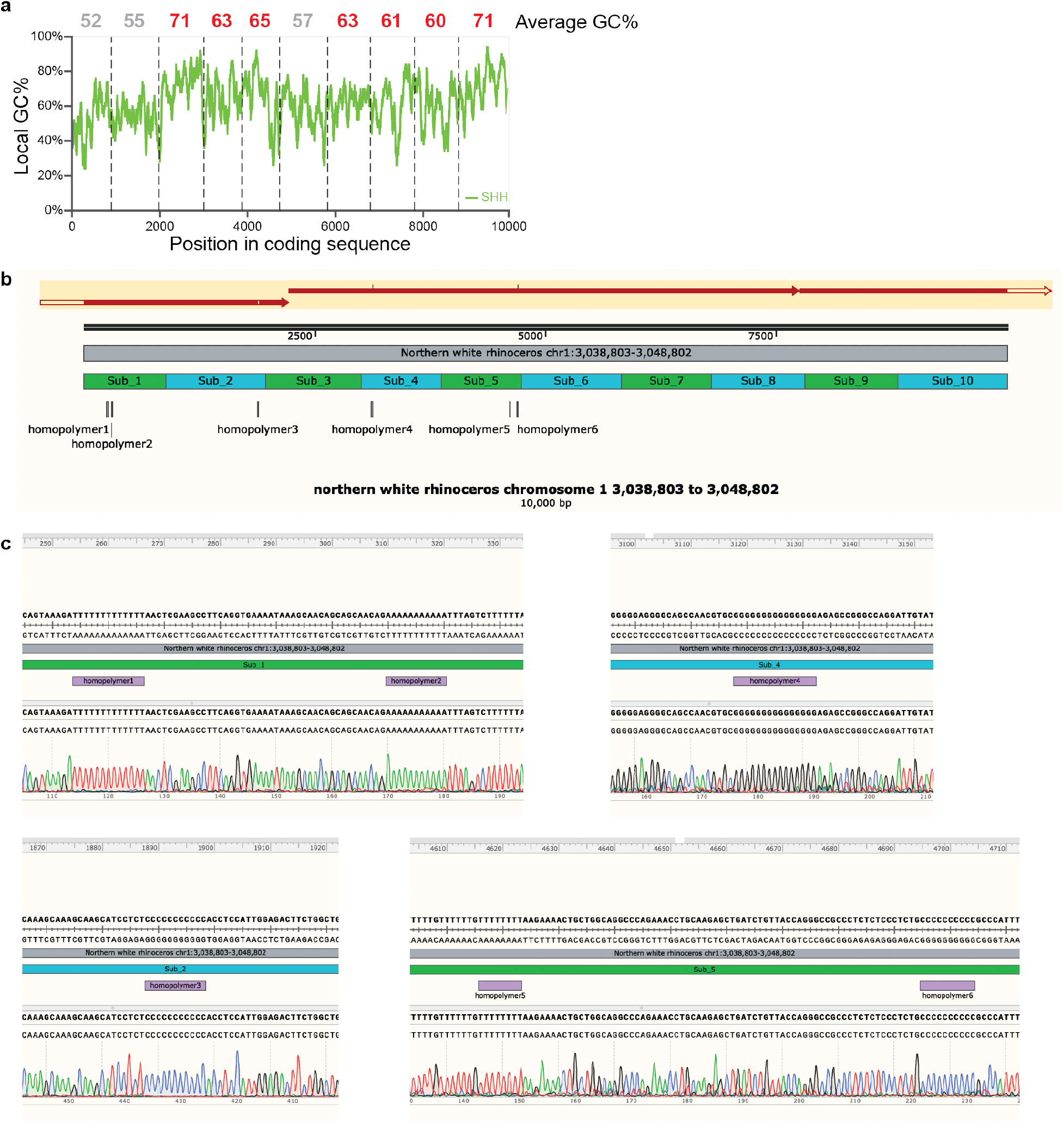
The complexity of the genomic sequence of the *Sonic Hedgehog* gene (SHH) in the northern white rhino adds difficulty in oligo synthesis, assembly, and sequencing. a) The average GC content of the 10 Sidewinder assemblies is listed above the region in the plot, and the dashed lines depict the boundaries of the 10, 1 kb Sidewinder assemblies. Multiple 1 kb regions have over 60% GC content, and there are 50 bp regions with as high as 94% GC content, as well as independent stretches of 11 and 15 consecutive guanine bases. b) Nanopore sequencing of the *in vivo* cloned 10 kb synthetic construct (red arrows) aligned to the Northern White Rhinoceros chromosome 1 shows alignment across the complete 10 kb stretch with individual SNPs aligned to homopolymer stretches. c) Sanger sequencing across the six homopolymer stretches in the synthetic construct to confirm mutation-free product in each of the 1 kb Sidewinder *in vivo* intermediates.

