## Supplementary Table 2 for "High temperature ligation enables *in vitro* assembly of eukaryotic genes"

Supplementary Table 2. FASTA Format sequences for sequences used and constructed.

>KanR_GFP_pSC101_Standard-USER (3666 bp)

gagttatacacagggctgggatctattctttttatctttttttattctttctttattctataaattataaccacttgaatataaacaaaaaaaacacacaaaggtctagcggaatttacagagggtctagcagaatttacaagttttccagcaaaggtctagcagaatttacagatacccacaactcaaaggaaaaggactagtaattatcattgactagcccatctcaattggtatagtgattaaaatcacctagaccaattgagatgtatgtctgaattagttgttttcaaagcaaatgaactagcgattagtcgctatgacttaacggagcatgaaaccaagctaattttatgctgtgtggcactactcaaccccacgattgaaaaccctacaaggaaagaacggacggtatcgttcacttataaccaatacgctcagatgatgaacatcagtagggaaaatgcttatggtgtattagctaaagcaaccagagagctgatgacgagaactgtggaaatcaggaatcctttggttaaaggctttgagattttccagtggacaaactatgccaagttctcaagcgaaaaattagaattagtttttagtgaagagatattgccttatcttttccagttaaaaaaattcataaaatataatctggaacatgttaagtcttttgaaaacaaatactctatgaggatttatgagtggttattaaaagaactaacacaaaagaaaactcacaaggcaaatatagagattagccttgatgaatttaagttcatgttaatgcttgaaaataactaccatgagtttaaaaggcttaaccaatgggttttgaaaccaataagtaaagatttaaacacttacagcaatatgaaattggtggttgataagcgaggccgcccgactgatacgttgattttccaagttgaactagatagacaaatggatctcgtaaccgaacttgagaacaaccagataaaaatgaatggtgacaaaataccaacaaccattacatcagattcctacctacataacggactaagaaaaacactacacgatgctttaactgcaaaaattcagctcaccagttttgaggcaaaatttttgagtgacatgcaaagtaagtatgatctcaatggttcgttctcatggctcacgcaaaaacaacgaaccacactagagaacatactggctaaatacggaaggatctgatgtgaTTGACAGCTAGCTCAGTCCTAGGTATAATGCTAGCaaagaggagaaaggatctatggctattacacgaagaagttttcttaaaggcgtagcaacgacaagtgccgcgtcagttatcggtccaagcttattagcgtcagcttcagcaaaagccgcgATGGTGTCGAAAGGTGAGGAGCTTTTCACGGGGGTGGTTCCAATCTTAGTCGAGCTTGATGGTGACGTGAATGGCCATAAGTTTAGCGTGCGCGGCGAGGGTGAGGGCGATGCGACGAACGGTAAACTTACTTTAAAGTTTATTTGTACTACTGGCAAATTACCGGTCCCGTGGCCGACCCTGGTCACTACGCTTGGGAATGGGGTAGCATGTTTTGCCCGTTACCCTGACCACATGAAACAGCACGATTTCTTTAAATCTGCCATGCCCGAAGGGTATGTTCAGGAGCGCACTATCAGTTTTAAAGACGACGGAACGTACAAAACTCGTGCTGAAGTAAAGTTCGAAGGTGATACCCTTGTGAACCGTATCGTCCTTAAAGGCATCGACTTCAAAGAGGACGGTAACATTTTAGGCCACAAATTAGAGTACAACTTCAATTCACATAAGGTTTATATCACAGCAGATAAGCAAAAAAATGGCATCAAGGCAAATTTTAAGACCCGCCACAACGTAGAAGATGGTGGAGTCCAGTTGGCGGACCACTACCAGCAAAATACACCGATTGGCGATGGTCCGGTTTTGCTTCCCGACAATCACTATCTTAGCCACCAGAGTAAACTGAGTAAGGACCCAAATGAAAAGCGCGACCACATGGTATTGAAAGAGCGCGTTACAGCAGCTGGAATTACACACGATATGGACGAACTTTATAAGTAAGGTACCCTCGAGtttgtttatttttctaaatacattcaaatatgtatccgctcatgagacaataaccctgataaatgcttcaataatattgaaaaaggaagagtATGAGCCATATTCAACGGGAAACGTCTTGCTCTAGGCCGCGATTAAATTCCAACATGGATGCTGATTTATATGGGTATAAATGGGCTCGCGATAATGTCGGGCAATCAGGTGCGACAATCTATCGATTGTATGGGAAGCCCGATGCGCCAGAGTTGTTTCTGAAACATGGCAAAGGTAGCGTTGCCAATGATGTTACAGATGAGATGGTCAGACTAAACTGGCTGACGGAATTTATGCCTCTTCCGACCATCAAGCATTTTATCCGTACTCCTGATGATGCATGGTTACTCACCACTGCGATCCCCGGGAAAACAGCATTCCAGGTATTAGAAGAATATCCTGATTCAGGTGAAAATATTGTTGATGCGCTGGCAGTGTTCCTGCGCCGGTTGCATTCGATTCCTGTTTGTAATTGTCCTTTTAACAGCGACCGCGTATTTCGTCTCGCTCAGGCGCAATCACGAATGAATAACGGTTTGGTTGATGCGAGTGATTTTGATGACGAGCGTAATGGCTGGCCTGTTGAACAAGTCTGGAAAGAAATGCATAAACTTTTGCCATTCTCACCGGATTCAGTCGTCACTCATGGTGATTTCTCACTTGATAACCTTATTTTTGACGAGGGGAAATTAATAGGTTGTATTGATGTTGGACGAGTCGGAATCGCAGACCGATACCAGGATCTTGCCATCCTATGGAACTGCCTCGGTGAGTTTTCTCCTTCATTACAGAAACGGCTTTTTCAAAAATATGGTATTGATAATCCTGATATGAATAAATTGCAGTTTCATTTGATGCTCGATGAGTTTTTCTAATTAATTAAACTAGTGAtgtctaacaattcgttcaagccgacggatatcgagctcgcttggactcctgttgatagatccagtaatgacctcagaactccatctggatttgttcagaacgctcggttgccgccgggcgttttttattggtgagaatccaagcactagggacagtaagacgggtaagcctgttgatgataccgctgccttactgggtgcattagccagtctgaatgacctgtcacgggataatccgaagtggtcagactggaaaatcagagggcaggaactgctgaacagcaaaaagtcagatagcaccacatagcagacccgccataaaacgccctgagaagcccgtgacgggcttttcttgtattatgggtagtttccttgcatgaatccataaaaggcgcctgtagtgccatttacccccattcactgccagagccgtgagcgcagcgaactgaatgtcacgaaaaagacagcgactcaggtgcctgatggtcggagacaaaaggaatattcagcgatttgcccgagcttgcgagggtgctacttaagcctttagggttttaaggtctgttttgtagaggagcaaacagcgtttgcgacatccttttgtaatactgcggaactgactaaagtagt

> KanR_GFP_pSC101_NSS-USER (3682 bp)

gagttatacacagggctgggatctattctttttatctttttttattctttctttattctataaattataaccacttgaatataaacaaaaaaaacacacaaaggtctagcggaatttacagagggtctagcagaatttacaagttttccagcaaaggtctagcagaatttacagatacccacaactcaaaggaaaaggactagtaattatcattgactagcccatctcaattggtatagtgattaaaatcacctagaccaattgagatgtatgtctgaattagttgttttcaaagcaaatgaactagcgattagtcgctatgacttaacggagcatgaaaccaagctaattttatgctgtgtggcactactcaaccccacgattgaaaaccctacaaggaaagaacggacggtatcgttcacttataaccaatacgctcagatgatgaacatcagtagggaaaatgcttatggtgtattagctaaagcaaccagagagctgatgacgagaactgtggaaatcaggaatcctttggttaaaggctttgagattttccagtggacaaactatgccaagttctcaagcgaaaaattagaattagtttttagtgaagagatattgccttatcttttccagttaaaaaaattcataaaatataatctggaacatgttaagtcttttgaaaacaaatactctatgaggatttatgagtggttattaaaagaactaacacaaaagaaaactcacaaggcaaatatagagattagccttgatgaatttaagttcatgttaatgcttgaaaataactaccatgagtttaaaaggcttaaccaatgggttttgaaaccaataagtaaagatttaaacacttacagcaatatgaaattggtggttgataagcgaggccgcccgactgatacgttgattttccaagttgaactagatagacaaatggatctcgtaaccgaacttgagaacaaccagataaaaatgaatggtgacaaaataccaacaaccattacatcagattcctacctacataacggactaagaaaaacactacacgatgctttaactgcaaaaattcagctcaccagttttgaggcaaaatttttgagtgacatgcaaagtaagtatgatctcaatggttcgttctcatggctcacgcaaaaacaacgaaccacactagagaacatactggctaaatacggaaggatctgaATCCATGCCTTTGACAGCTAGCTCAGTCCTAGGTATAATGCTAGCaaagaggagaaaggatctatggctattacacgaagaagttttcttaaaggcgtagcaacgacaagtgccgcgtcagttatcggtccaagcttattagcgtcagcttcagcaaaagccgcgATGGTGTCGAAAGGTGAGGAGCTTTTCACGGGGGTGGTTCCAATCTTAGTCGAGCTTGATGGTGACGTGAATGGCCATAAGTTTAGCGTGCGCGGCGAGGGTGAGGGCGATGCGACGAACGGTAAACTTACTTTAAAGTTTATTTGTACTACTGGCAAATTACCGGTCCCGTGGCCGACCCTGGTCACTACGCTTGGGAATGGGGTAGCATGTTTTGCCCGTTACCCTGACCACATGAAACAGCACGATTTCTTTAAATCTGCCATGCCCGAAGGGTATGTTCAGGAGCGCACTATCAGTTTTAAAGACGACGGAACGTACAAAACTCGTGCTGAAGTAAAGTTCGAAGGTGATACCCTTGTGAACCGTATCGTCCTTAAAGGCATCGACTTCAAAGAGGACGGTAACATTTTAGGCCACAAATTAGAGTACAACTTCAATTCACATAAGGTTTATATCACAGCAGATAAGCAAAAAAATGGCATCAAGGCAAATTTTAAGACCCGCCACAACGTAGAAGATGGTGGAGTCCAGTTGGCGGACCACTACCAGCAAAATACACCGATTGGCGATGGTCCGGTTTTGCTTCCCGACAATCACTATCTTAGCCACCAGAGTAAACTGAGTAAGGACCCAAATGAAAAGCGCGACCACATGGTATTGAAAGAGCGCGTTACAGCAGCTGGAATTACACACGATATGGACGAACTTTATAAGTAAATCCCATCTCTGGTACCCTCGAGtttgtttatttttctaaatacattcaaatatgtatccgctcatgagacaataaccctgataaatgcttcaataatattgaaaaaggaagagtATGAGCCATATTCAACGGGAAACGTCTTGCTCTAGGCCGCGATTAAATTCCAACATGGATGCTGATTTATATGGGTATAAATGGGCTCGCGATAATGTCGGGCAATCAGGTGCGACAATCTATCGATTGTATGGGAAGCCCGATGCGCCAGAGTTGTTTCTGAAACATGGCAAAGGTAGCGTTGCCAATGATGTTACAGATGAGATGGTCAGACTAAACTGGCTGACGGAATTTATGCCTCTTCCGACCATCAAGCATTTTATCCGTACTCCTGATGATGCATGGTTACTCACCACTGCGATCCCCGGGAAAACAGCATTCCAGGTATTAGAAGAATATCCTGATTCAGGTGAAAATATTGTTGATGCGCTGGCAGTGTTCCTGCGCCGGTTGCATTCGATTCCTGTTTGTAATTGTCCTTTTAACAGCGACCGCGTATTTCGTCTCGCTCAGGCGCAATCACGAATGAATAACGGTTTGGTTGATGCGAGTGATTTTGATGACGAGCGTAATGGCTGGCCTGTTGAACAAGTCTGGAAAGAAATGCATAAACTTTTGCCATTCTCACCGGATTCAGTCGTCACTCATGGTGATTTCTCACTTGATAACCTTATTTTTGACGAGGGGAAATTAATAGGTTGTATTGATGTTGGACGAGTCGGAATCGCAGACCGATACCAGGATCTTGCCATCCTATGGAACTGCCTCGGTGAGTTTTCTCCTTCATTACAGAAACGGCTTTTTCAAAAATATGGTATTGATAATCCTGATATGAATAAATTGCAGTTTCATTTGATGCTCGATGAGTTTTTCTAATTAATTAAACTAGTGAtgtctaacaattcgttcaagccgacggatatcgagctcgcttggactcctgttgatagatccagtaatgacctcagaactccatctggatttgttcagaacgctcggttgccgccgggcgttttttattggtgagaatccaagcactagggacagtaagacgggtaagcctgttgatgataccgctgccttactgggtgcattagccagtctgaatgacctgtcacgggataatccgaagtggtcagactggaaaatcagagggcaggaactgctgaacagcaaaaagtcagatagcaccacatagcagacccgccataaaacgccctgagaagcccgtgacgggcttttcttgtattatgggtagtttccttgcatgaatccataaaaggcgcctgtagtgccatttacccccattcactgccagagccgtgagcgcagcgaactgaatgtcacgaaaaagacagcgactcaggtgcctgatggtcggagacaaaaggaatattcagcgatttgcccgagcttgcgagggtgctacttaagcctttagggttttaaggtctgttttgtagaggagcaaacagcgtttgcgacatccttttgtaatactgcggaactgactaaagtagt

>LuxABCDE_p15a_CmR (7869 bp)

atatccctaggtataaacgcagaaaggcccacccgaaggtgagccagtgtgaCCTCCTCATCCATGCCTGAATTCGCGGCCGCTTCTAGAGGAAAtcataaaaaatttatTTGCTTtgtgagcggataacaatTATAATAGATTCAATTGTGAGCGGATAACAATTACTAGAGAAAGAGGAGAAATACTAGATGAAATTTGGAAACTTTTTGCTTACATACCAACCTCCCCAATTTTCTCAAACAGAGGTAATGAAACGTTTGGTTAAATTAGGTCGCATCTCTGAGGAGTGTGGTTTTGATACCGTATGGTTACTGGAGCATCATTTCACGGAGTTTGGTTTGCTTGGTAACCCTTATGTCGCTGCTGCATATTTACTTGGCGCGACTAAAAAATTGAATGTAGGAACTGCCGCTATTGTTCTTCCCACAGCCCATCCAGTACGCCAACTTGAAGATGTGAATTTATTGGATCAAATGTCAAAAGGACGATTTCGGTTTGGTATTTGCCGAGGGCTTTACAACAAGGACTTTCGCGTATTCGGCACAGATATGAATAACAGTCGCGCCTTAGCGGAATGCTGGTACGGGCTGATAAAGAATGGCATGACAGAGGGATATATGGAAGCTGATAATGAACATATCAAGTTCCATAAGGTAAAAGTAAACCCCGCGGCGTATAGCAGAGGTGGCGCACCGGTTTATGTGGTGGCTGAATCAGCTTCGACGACTGAGTGGGCTGCTCAATTTGGCCTACCGATGATATTAAGTTGGATTATAAATACTAACGAAAAGAAAGCACAACTTGAGCTTTATAATGAAGTGGCTCAAGAATATGGGCACGATATTCATAATATCGACCATTGCTTATCATATATAACATCTGTAGATCATGACTCAATTAAAGCGAAAGAGATTTGCCGGAAATTTCTGGGGCATTGGTATGATTCTTATGTGAATGCTACGACTATTTTTGATGATTCAGACCAAACAAGAGGTTATGATTTCAATAAAGGGCAGTGGCGTGACTTTGTATTAAAAGGACATAAAGATACTAATCGCCGTATTGATTACAGTTACGAAATCAATCCCGTGGGAACGCCGCAGGAATGTATTGACATAATTCAAAAAGACATTGATGCTACAGGAATATCAAATATTTGTTGTGGATTTGAAGCTAATGGAACAGTAGACGAAATTATTGCTTCCATGAAGCTCTTCCAGTCTGATGTCATGCCATTTCTTAAAGAAAAACAACGTTCGCTATTATATTAATAATACTAGAGAAAGAGGAGAAATACTAGATGAAATTTGGATTGTTCTTCCTTAACTTCATCAATTCAACAACTGTTCAAGAACAAAGTATAGTTCGCATGCAGGAAATAACGGAGTATGTTGATAAGTTGAATTTTGAACAGATTTTAGTGTATGAAAATCATTTTTCAGATAATGGTGTTGTCGGCGCTCCTCTGACTGTTTCTGGTTTTCTGCTCGGTTTAACAGAGAAAATTAAAATTGGTTCATTAAATCACATCATTACAACTCATCATCCTGTCCGCATAGCGGAGGAAGCTTGCTTATTGGATCAGTTAAGTGAAGGGAGATTTATTTTAGGGTTTAGTGATTGCGAAAAAAAAGATGAAATGCATTTTTTTAATCGCCCGGTTGAATATCAACAGCAACTATTTGAAGAGTGTTATGAAATCATTAACGATGCTTTAACAACAGGCTATTGTAATCCAGATAACGATTTTTATAGCTTCCCTAAAATATCTGTAAATCCCCATGCTTATACGCCAGGCGGACCTCGGAAATATGTAACAGCAACCAGTCATCATATTGTTGAGTGGGCGGCCAAAAAAGGTATTCCTCTCATCTTTAAGTGGGATGATTCTAATGATGTTAGATATGAATATGCTGAAAGATATAAAGCCGTTGCGGATAAATATGACGTTGACCTATCAGAGATAGACCATCAGTTAATGATATTAGTTAACTATAACGAAGATAGTAATAAAGCTAAACAAGAGACGCGTGCATTTATTAGTGATTATGTTCTTGAAATGCACCCTAATGAAAATTTCGAAAATAAACTTGAAGAAATAATTGCAGAAAACGCTGTCGGAAATTATACGGAGTGTATAACTGCGGCTAAGTTGGCAATTGAAAAGTGTGGTGCGAAAAGTGTATTGCTGTCCTTTGAACCAATGAATGATTTGATGAGCCAAAAAAATGTAATCAATATTGTTGATGATAATATTAAGAAGTACCACATGGAATATACCTAATAATACTAGAGAAAGAGGAGAAATACTAGATGACTAAAAAAATTTCATTCATTATTAACGGCCAGGTTGAAATCTTTCCCGAAAGTGATGATTTAGTGCAATCCATTAATTTTGGTGATAATAGTGTTTACCTGCCAATATTGAATGACTCTCATGTAAAAAACATTATTGATTGTAATGGAAATAACGAATTACGGTTGCATAACATTGTCAATTTTCTCTATACGGTAGGGCAAAGATGGAAAAATGAAGAATACTCAAGACGCAGGACATACATTCGTGACTTAAAAAAATATATGGGATATTCAGAAGAAATGGCTAAGCTAGAGGCCAATTGGATATCTATGATTTTATGTTCTAAAGGCGGCCTTTATGATGTTGTAGAAAATGAACTTGGTTCTCGCCATATCATGGATGAATGGCTACCTCAGGATGAAAGTTATGTTCGGGCTTTTCCGAAAGGTAAATCTGTACATCTGTTGGCAGGTAATGTTCCATTATCTGGGATCATGTCTATATTACGCGCAATTTTAACTAAGAATCAGTGTATTATAAAAACATCGTCAACCGATCCTTTTACCGCTAATGCATTAGCGTTAAGTTTTATTGATGTAGACCCTAATCATCCGATAACGCGCTCTTTATCTGTTATATATTGGCCCCACCAAGGTGATACATCACTCGCAAAAGAAATTATGCGACATGCGGATGTTATTGTCGCTTGGGGAGGGCCAGATGCGATTAATTGGGCGGTAGAGCATGCGCCATCTTATGCTGATGTGATTAAATTTGGTTCTAAAAAGAGTCTTTGCATTATCGATAATCCTGTTGATTTGACGTCCGCAGCGACAGGTGCGGCTCATGATGTTTGTTTTTACGATCAGCGAGCTTGTTTTTCTGCCCAAAACATATATTACATGGGAAATCATTATGAGGAATTTAAGTTAGCGTTGATAGAAAAACTTAATCTATATGCGCATATATTACCGAATGCCAAAAAAGATTTTGATGAAAAGGCGGCCTATTCTTTAGTTCAAAAAGAAAGCTTGTTTGCTGGATTAAAAGTAGAGGTGGATATTCATCAACGTTGGATGATTATTGAGTCAAATGCAGGTGTGGAATTTAATCAACCACTTGGCAGATGTGTGTACCTTCATCACGTCGATAATATTGAGCAAATATTGCCTTATGTTCAAAAAAATAAGACGCAAACCATATCTATTTTTCCTTGGGAGTCATCATTTAAATATCGAGATGCGTTAGCATTAAAAGGTGCGGAAAGGATTGTAGAAGCAGGAATGAATAACATATTTCGAGTTGGTGGATCTCATGACGGAATGAGACCGTTGCAACGATTAGTGACATATATTTCTCATGAAAGGCCATCTAACTATACGGCTAAGGATGTTGCGGTTGAAATAGAACAGACTCGATTCCTGGAAGAAGATAAGTTCCTTGTATTTGTCCCATAATAATACTAGAGAAAGAGGAGAAATACTAGATGGAAAATGAATCAAAATATAAAACCATCGACCACGTTATTTGTGTTGAAGGAAATAAAAAAATTCATGTTTGGGAAACGCTGCCAGAAGAAAACAGCCCAAAGAGAAAGAATGCCATTATTATTGCGTCTGGTTTTGCCCGCAGGATGGATCATTTTGCTGGTCTGGCGGAATATTTATCGCGGAATGGATTTCATGTGATCCGCTATGATTCGCTTCACCACGTTGGATTGAGTTCAGGGACAATTGATGAATTTACAATGTCTATAGGAAAGCAGAGCTTGTTAGCAGTGGTTGATTGGTTAACTACACGAAAAATAAATAACTTCGGTATGTTGGCTTCAAGCTTATCTGCGCGGATAGCTTATGCAAGCCTATCTGAAATCAATGCTTCGTTTTTAATCACCGCAGTCGGTGTTGTTAACTTAAGATATTCTCTTGAAAGAGCTTTAGGGTTTGATTATCTCAGTCTACCCATTAATGAATTGCCGGATAATCTGGATTTTGAAGGCCATAAATTGGGTGCTGAAGTCTTTGCGAGAGATTGTCTTGATTTTGGTTGGGAAGATTTAGCTTCTACAATTAATAACATGATGTATCTTGATATACCGTTTATTGCTTTTACTGCAAATAACGATAATTGGGTCAAGCAAGATGAAGTTATCACATTGTTATCAAATATTCGTAGTAATCGATGCAAGATATATTCTTTGTTAGGAAGTTCGCATGACTTGAGTGAAAATTTAGTGGTCCTGCGCAATTTTTATCAATCGGTTACGAAAGCCGCTATCGCGATGGATAATGATCATCTGGATATTGATGTTGATATTACTGAACCGTCATTTGAACATTTAACTATTGCGACAGTCAATGAACGCCGAATGAGAATTGAGATTGAAAATCAAGCAATTTCTCTGTCTTAATAATACTAGAGAAAGAGGAGAAATACTAGATGACTTCATATGTTGATAAACAAGAAATTACAGCAAGCTCAGAAATTGATGATTTGATTTTTTCGAGCGATCCATTAGTGTGGTCTTACGACGAGCAGGAAAAAATCAGAAAGAAACTTGTGCTTGATGCATTTCGTAATCATTATAAACATTGTCGAGAATATCGTCACTACTGTCAGGCACACAAAGTAGATGACAATATTACGGAAATTGATGACATACCTGTATTCCCAACATCGGTTTTTAAGTTTACTCGCTTATTAACTTCTCAGGAAAACGAGATTGAAAGTTGGTTTACCAGTAGCGGCACGAATGGTTTAAAAAGTCAGGTGGCGCGTGACAGATTAAGTATTGAGAGACTCTTAGGCTCTGTGAGTTATGGCATGAAATATGTTGGTAGTTGGTTTGATCATCAAATAGAATTAGTCAATTTGGGACCAGATAGATTTAATGCTCATAATATTTGGTTTAAATATGTTATGAGTTTGGTGGAATTGTTATATCCTACGACATTTACCGTAACAGAAGAACGAATAGATTTTGTTAAAACATTGAATAGTCTTGAACGAATAAAAAATCAAGGGAAAGATCTTTGTCTTATTGGTTCGCCATACTTTATTTATTTACTCTGCCATTATATGAAAGATAAAAAAATCTCATTTTCTGGAGATAAAAGCCTTTATATCATAACCGGAGGCGGCTGGAAAAGTTACGAAAAAGAATCTCTGAAACGTGATGATTTCAATCATCTTTTATTTGATACTTTCAATCTCAGTGATATTAGTCAGATCCGAGATATATTTAATCAAGTTGAACTCAACACTTGTTTCTTTGAGGATGAAATGCAGCGTAAACATGTTCCGCCGTGGGTATATGCGCGAGCGCTTGATCCTGAAACGTTGAAACCTGTACCTGATGGAACGCCGGGGTTGATGAGTTATATGGATGCGTCAGCAACCAGTTATCCAGCATTTATTGTTACCGATGATGTCGGGATAATTAGCAGAGAATATGGTAAGTATCCCGGCGTGCTCGTTGAAATTTTACGTCGCGTCAATACGAGGACGCAGAAAGGGTGTGCTTTAAGCTTAACCGAAGCGTTTGATAGTTAAgatacaatcgatttctggatccgcaggcctctgctagctTGACTGACTGAGATACAGCGTACCTTCAGCTCACAGACATGATAAGATACATTGATGAGTTTGGACAAACCACAACTAGAATGCAGTGAAAAAAATGCTTTATTTGTGAAATTTGTGATGCTATTGCTTTATTTGTAACCATTATAAGCTGCAATAAACAAGTTAACAACAACAATTGCATTCATTTTATGTTTCAGGTTCAGGGGGAGGTGTGGGAGGTTTTTTAAAGcaaacgctaaattgcctgatgcgctacgcttatcATCCTTAGCGAAAGCTAAGGAATCCCATCTCTtgatcggcacgtaagaggttccaactttcaccataatgaaataagatcactaccgggcgtattttttgagttatcgagattttcaggagctaaggaagctaaaatggagaaaaaaatcactggatataccaccgttgatatatcccaatggcatcgtaaagaacattttgaggcatttcagtcagttgctcaatgtacctataaccagaccgttcagctggatattacggcctttttaaagaccgtaaagaaaaataagcacaagttttatccggcctttattcacattcttgcccgcctgatgaatgctcatccggaatttcgtatggcaatgaaagacggtgagctggtgatatgggatagtgttcacccttgttacaccgttttccatgagcaaactgaaacgttttcatcgctctggagtgaataccacgacgatttccggcagtttctacacatatattcgcaagatgtggcgtgttacggtgaaaacctggcctatttccctaaagggtttattgagaatatgtttttcgtctcagccaatccctgggtgagtttcaccagttttgatttaaacgtggccaatatggacaacttcttcgcccccgttttcaccatgggcaaatattatacgcaaggcgacaaggtgctgatgccgctggcgattcaggttcatcatgccgtttgtgatggcttccatgtcggcagaatgcttaatgaattacaacagtactgcgatgagtggcagggcggggcgtaatttgatatcgagctcgcttggactcctgttgatagatccagtaatgacctcagaactccatctggatttgttcagaacgctcggttgccgccgggcgttttttattggtgagaatccaagcactagtaacaacttatatcgtatggggctgacttcaggtgctacatttgaagagataaattgcactgaaatctagaaatattttatctgattaataagatgatcttcttgagatcgttttggtctgcgcgtaatctcttgctctgaaaacgaaaaaaccgccttgcagggcggtttttcgaaggttctctgagctaccaactctttgaaccgaggtaactggcttggaggagcgcagtcaccaaaacttgtcctttcagtttagccttaaccggcgcatgacttcaagactaactcctctaaatcaattaccagtggctgctgccagtggtgcttttgcatgtctttccgggttggactcaagacgatagttaccggataaggcgcagcggtcggactgaacggggggttcgtgcatacagtccagcttggagcgaactgcctacccggaactgagtgtcaggcgtggaatgagacaaacgcggccataacagcggaatgacaccggtaaaccgaaaggcaggaacaggagagcgcacgagggagccgccagggggaaacgcctGgtatctttatagtcctgtcgggtttcgccaccactgatttgagcgtcagatttcgtgatgcttgtcaggggggcggagcctatggaaaaacggctttgccgcggccctctcacttccctgttaagtatcttcctggcatcttccaggaaatctccgccccgttcgtaagccatttccgctcgccgcagtcgaacgaccgagcgtagcgagtcagtgagcgaggaagcggaat

>Northern_White_Rhino_Genomic_SSH (10000 bp)

TCTCTGAAAAACTATTTAAATAAGACCTGGGCAGTTTGCCCCTCAAAAATGTGAGACAATATTTGTTTGATGGTCAAAGCTGGGGGCGGGGGTCTAGACTGTCCTTTAAAAACTTAAGTTGGTTTTAACTGAAGCACCTACCATTCTGGAGAAAGTGAAGGGGGAACGCGGAGATGTGCACAAAGCAAGAGTAATTGATGTTATCCTGAGGAACGTGCCTCCGCCCTCCCGCCCACCTCTGTCTCAGTAAAGATTTTTTTTTTTTTAACTCGAAGCCTTCAGGTGAAAATAAAGCAACAGCAGCAACAGAAAAAAAAAAATTTAGTCTTTTTTAGGGTTAAACATCAGAAGACAAGCTTGTGCTGCCTTGGCCAATCAGATACGCCCCTGCCAGCTATAATATAGTACTGACGGCAGCCTCTCTCACAAGCTCTCGAGGCTTGCTACCATTTAAAATCAGACTCTTTTTGTCTTTTGATTGCTGTCTCGCGCCCCAACTCCGATGTGTTCCGTTATCAGCGACCAGCAGCCAGCCGTTCCAGCCCCCGTCTGGGTGGGGATCAGCGGCGAGTCCCCTGCAGCAGCGGCAGGCAAGGTTATATAGGAAGAGAAAGAGCGAGGCACAGAGCGAGGGAGCCAGCAGCAAGCCGGAGCGAGCAAGGGAGAGCGCAGCCCGAGCGCACAGGCACACACACACTTGCACACACAGCCCCACACGGGGACAGCTTTGAAGTCACTGGGTTCCATGGACGAGATGCTGCTGCTGGCGAGATGTCTGCTGGTGGTTCTCGTCTCCTCGCTGCTGATGTGCTCGGGGCTGGCGTGCGGACCCGGCAGGGGATTTGGGAAAAGGCGGCACCCCAAAAAGCTGACCCCTTTAGCCTACAAGCAGTTTATCCCCAACGTGGCTGAGAAGACCCTAGGGGCCAGTGGAAGATATGAAGGGAAGATCTCGAGAAACTCGGAGCGATTTAAGGAACTCACCCCCAATTACAACCCCGACATCATATTTAAGGATGAAGAAAACACTGGAGCGGACCGGCTGATGACTCAGGTAGAAACCCAGCACCGGGCGTGGAATGTGTTGCTTTCAGGGGGTTACGTGTAACCAAGCAGTTGCAGACTTAGTCTGCTTGCTCCTGAGGCAAATAGAAGGCCATCTCACCCTCTCCTTCCCCACCCACCTGCCTCCCTACCTGCCCTCCTCTTCCTGGAGGGAATGGAGGGAAGGGTCCAGCTAGAGAAGTTTAGGGTGGGGGCAGGGGAGGAAGGTAGCAGACGGTCCATACATGGCCCTGATTTGATTTGTCCTGCTGGGATGGCCTTCCCATCCTGCCTCAGCTCTAGCTTGTAACTCGAGGAGACCTTCACTTTGGGGGCATCTGTCCTTTCTGAGTTGGGAGTGGAAATGGAGGGAGGGGGCAAGGATCCCCTCAAACACAAAGTGGCTTACAGGTCACTGTACAATGTGGCCTTGACCATTACGCTTCAGACTGTACACGGCCCCATCTCCTCTGTGGTTTTCTTTCCCACAACTACCCCCACCCCACTGGGCCCCCACCAAAGACCCTCAATGGGAAAAGCCAGGAGGCAGTTCTGACAAAAGGCAGGGGCTCCAGAGGCGGCTGGGCTGGTCCCTGATGGGCTCGCAGAGCTCCACAGGCGATCCCAGCGGGGCAGTCTTGCCCATCAGAGTGTGAATGTATTGATATTTCTTTAAGGATGCTCTTACATTCTCCCGAGCCCGCGGTACCTGAGGGGAGGGACTTAGAACTTATCGGCATTGCATCACACTAGTTTCCAACCTGCTTGCACAGGAATTGGGAGCGACTAAATATTAGCGGGAGAGGGCTGGAAAGGAGAACCAAAGCAAAGCAAGCATCCTCTCCCCCCCCCCCACCTCCATTGGAGACTTCTGGCTGCCAATCTCCCCACCAATGGCAGACTTTCTGAAACTCCACGCACGTATAACTGAAGCTAGAAATGGATTTCCTGGCAAATATAGGTCAACATCCTTTTTATTGCCCTATTAAAATATTCAAGTCCTACCTTTAGGGCTAGGAGCGCACCGCAGCTGACAGAGCGGCGCTGGTGGGGCGCAAGTGCAGGGGAGGGTGGGGGCGGGAGGGAGAGAGGAGCACCTCCGGGCCGCTCGGCTTCCCAGCCCGGTTCCCGGCCTTGCTGCGGAGCTGAGAAAGGAGAGGGGCACCCGGGGTAGCCGGCTTGAGCTGACAGAAGGGGTGGGAAAGGGAGCGGAGGAGAGGGTCGGCATCCCTCACTCTCTCTCATAAGCTCCCGGCCACCTTAAATCTGGATCGCTAGTGCGGACGCACGCGCCTGTGCCCGAGAGCAGCGCGAGCCACAATTAACTCTGCGCAGGCCTCGGGGCGCCCCGTGCGTGCGGCGGGGGCTCTGACCCCCTGGTCACCCTTGCTCCCCGCGGGAGGGAGGTGGCCTTCCCGGGGTCCCCTGGGACCGGGGAAGAGGGAGGCGACGCAAGAGAAAGCGGCGGGGGCGGGGACACCAGAGGTTCCGAGCAGAGGGGTCAGCGTCCGCCCTGGGCTGCGCCGGGCCGACGCTGGAGGGGGTCCCGGCCTCGCCGCCCTGCTGCCAAACCGCGAGCGCGCGGAGCCGCGCACCCCGGCAGACGCATTTGGGCTCCACTTACTCCGGGGCTATCGATCCCGCGGCTAGGGTTTCAGTGCGCGGGGGTGGGCTGCGGCCGCCCGCCCGGCCCTCCACGCGCCGAAGCGCGGGCTTGGCCAAGGCTAATGGCAGGGCTCCGACGGCCGCGATGGCCCCGGCGGCGCAGGCTGGACCAGCGGCAGGAGAAGCCGGTTCCAGGGCGGCCTCCCCGCCGCCTCCCCGGCCTCTCGGCCCCGGCGCCTCAGCCCGAGCCCCCAGTCGGGTCTCTCTTCCGGCCGCCCCGCCGGGGACCCACGGCCAGCGGCCCCGCGCCCGGCCCTCTCGCCGCCGCATACGCGGGCAGCATTTGAACTTCTGACCTTCTGTCAACTTCCCTCAGACGAACGAAACGAAAACAAATACTTTTTTTCCTTGGGCAGTGGCTATTCCCGTTCCCAACACAAAAGGACGGGGGAGGGGCAGCCAACGTGCGGGGGGGGGGGGGGGAGAGCCGGGCCAGGATTGTATCACGGACCCCACAAATGTCCCCTAAAAGCGCGGATGGCTGGGGCCGGCCTGGAAGGTGCCCCTGGGTTTGTCCCTTCCCTGTCTCCTTCTTCTCTGGTTCTAAAAGTTGAAGGAGGGGAGGGATGGGGGAGGCGTTGGACCCGGGCGGCGAGGGGCAGGGCGGAGGTGAAGGGGTGACGCCGGCACGCCTCCCAGGCCAGGGCAGGTTAGGGCTCCTGGGGAACCTGCTCACCGCGGGCCCCTCATTTAAGATAACCACGTTGCCACTGTTCTCAGGTGCGTCTCCCTCCAGCCCTGTCTCGGCCGTCCCTCCGCGTCTCCGTGTCTCTCTATCTCTCTAATGTGACTGCCGCCCAAGTCCCTCAACCATGGCAAGATCGTCCCCAGTGGAACTTTCAGAGCAGTTCTGGAACGCAGGAGCTCCTGGTTAATATTAACTCGGAGAGGGGGAAGCGCAGACAGCACCTCTCCGCGCAGGCCGCGGGTGCCAGGGGGGTCTGCGGGGGCAGGAGGCCGCGAGGGGCCGGCACGGGGCTGCCGAGGGGCTCCCGAGGGGCTCCACCTGAGACGCCACGTGGAGTACATTTCTGTGGCGTGGCGGCCCTGGGGACGGCGCTCAGGTGTGGGTTCGAGGAGGGTTCTCCTGTTTTGATGCGAGGCGTCTGATTTTCCGGAGTGGGGGGAGGAGAGGGCGGAGAGAGGTGTGTCTAGGGACTAGGGGTCTCTTTTGAAGCTGGGACACCCCTCCCCCGCACTGGGGAAGCCCGTGGGCCGTTGGCGGTGACAGTCCCTGCTCGGGCTTCTGGGCGCGGAGCGGCTGCGCAGACTGGGGAGGCGGCGGGAGCTCAGGCGTGGGGAACAACCTGACCTCCGCCGACACAAAGCCCGCCTGGCGGCCCCGCCGGGCTTCACGGTGGCTGCACAGAGTCCGGCTTGATTCGCGGCACACGACTCAGTGAATTGATAACCGGCCTGGGCTTCGCGGCTTTGCCTGCGCCGAGCGCCGGGCCGGGGCGCAGAGCGTCCTGCAGCCACGGGCCGAGGCCGCGCCGCGCCTGCTGGCGCCGGGGAGGCGCGTGGGGCGCGGGGCGCCCACTCGGGCCTGGCTCGGCCCGGGGCTTCTCGCCGGGGGTCCTCCAGCTCTTGCCCAAGTAGCGCAGCTCAGGGCGCTGGGGGCTGGTACGCGTCCTCCCCATCTTGCCCTCCCCCTCGCCAGGTGCCGGGTGGGAAGCTGCTGAAAGCTTCATCTGCCCTCCAGGAGGAGGAAACCGGGGTGGGATGGGGGCTCCGGACGCGGTGTGTGCGTGTGCGTGTCGGCGGTGTGCGCCTGCGCAGCCGGCTTCACATCCCGGCTCTAAACTGACCCTGGTTTACTGATTGATTTTCATGTAAAACGCGTTCAGTTCTCAAGATGACCTCACTCAAACTCTGCCCTTCCGAGTTTTTTGTTTTTTGTTTTTTGTTTTTTTTAAGAAAACTGCTGGCAGGCCCAGAAACCTGCAAGAGCTGATCTGTTACCAGGGCCGCCCTCTCTCCCTCTGCCCCCCCCCCGCCCATTTGTTGTATGTTTAACTCTACAACCATGTTACTGGTTTTTATTGCTGCCAAATACACAGGGCTCTTCCTGTTGCGCAATCTGTCACATCCCCTTAACACGGGGGCCGCGGCCCTTCTTGCAAGAGCCCCCTCCCCTGCTCTGTAGGAAGAGCTCCCCCGCGCTGTGGTGACGTGGTGAGGGTCGTCGATGGGGCCGTCTCTCCAGCCCGGTGCCCAGGCTCCCAAGCAGAGCAGCCGCCCGCCCGGAGGAGACCAGCCCTGCCCTCTGCAGAAGTCACCCTGAAACTGTCGCCTACTCCAGGGGGTCTGCCTCTCCCGGGGATGCCTCGTCAGGATTTGTACCCACTTCCCGTCTTTCAAATCCCGCACCCCGCCCCCCAGCAGCAGTATAAACGACCGGGGAGTTAGTACCCACCCAAACACACACACTTGACTCAACTCTGTTTTTTCTTGCATCCTTATCTGAACGGGACCGAGAATTATGCGGGGAACTCCCACAACCCGGGGTTCCACTTTGCCCTGGTTGGTGTCCCGCCTAGTCAATGTTTCTTTACTCCTACAGGCGGCTCGGGGAGGAGGGGGGTGGGGGGGTTGGGGGCCCAAGGGCAATTTCAAGAGGGAGACTCCCCAGGGCGAGGGCTCCAAGGCGCAGGCCTCTGGGCTCCCATCCCCCAGCCTCCGCCTCCACCTCTTAAGTTTGCTCTTCAAACTTGCTGGACACCATTCTGCGCTGGGGCCAAGAAAACAAGAGGGGGGAAGGGAACAGGTTAGCGAAAACAAGAGACACAATCAGACCACAGAAAAGCCGGGAGAAAAGAGTTCCACAGCAAAAGGAATGTGGCTGTCCAGATAAAGACTGTTTGTCCTCCGCTCGCTGCACGGAGAAGTCTCACTCACCTGGTAGCCTTTCACTTCTGGCTCCCCCTCCCCAAGCACGGCTGTGGGCTTCCCACCTCCCTCTCCACCTTTGGCTTAATTCTCACCTCAGATAGGCATAGGGTAGGAGGGAGGGTCCAGAGATAAGGCAAGGCTTTTAATTCTTTAAGAAGTAGACAGTGTATTTTTCTTGATTTCTGCAAAGTTGCACATTGGGCCTCTCTCCGGCCCCCTGATGAGGAAGTGTGCCCAGGGAGATGGGTGGAGCCTTGCACAGGTGGGAGAAATGGGGCCACAGAAAGGGGCTGAGGAGTGGAGGGCGTCCTGGGGGAGGAGTGGGGCATTGCCGAGCATGCCCCCTCTGCGTTCTGCAGCCATTGTCTGAAAACACGCAGTTTCCTTGTTTCAAAGAAAAGCCATCTGCCTCTCAGGCCATGGTGGGGGCTCTGCCACTTCAGAACTTGGAGAAAGGCCCTCATGTCAGCTCATCTCCAAGAGGACTTGCTCCCCGAGCAGAGGTCACCCCTCCTTGCCGCCTAGTGCCATCGCTTCTGAGCTGGCTGGGCCCCGCTGCAGGCCTTGTCTCTCTACTTGTCCACACAGCCCGAGTCCTCAGGGATGGGGCAGGGCAGCTGATGCAGGGGCGGGGACGGGTGGTGGGCCCTAACATCTGTGGGGTCGCCCCTTCTCTCCGCAGAGGTGTAAGGACAAGTTGAATGCCTTGGCCATCTCAGTGATGAACCAGTGGCCGGGAGTGAAGCTGCGGGTGACCGAGGGCTGGGACGAGGACGGCCACCACTCAGAGGAGTCACTGCACTACGAGGGCCGCGCCGTGGACATCACCACCTCAGACCGAGACCGCAGCAAGTATGGCATGCTGGCGCGCCTGGCCGTGGAGGCCGGCTTTGACTGGGTCTACTACGAGTCCAAAGCGCACATCCACTGCTCCGTGAAAGCAGGTGAGGGGTGGGAAAGAGATAAATGGGGAGGGGCACGAGGCGCAGGGAGGGAGGCAACTGGGTGATGTGAGAGCACACCCTATAAGGGGTGTCCCCCCAGCCAGGAACTGCCCCTACCTTTGAGGCCACGGAGGCCCAGGTTCTGGGAGCCCAGATGCAGAGGGCATGCGGCTTCCTGAGCCTTCCCCAACCCTGTTTCCGGAGCCCCACTGCAGGTGGCGCAGACAGGGCAGCACGAATGGGGAGGGCGTGGACTTTCCTGGTGAGAGACTGCAGCCCTGCACGGTTAGCCAGGTCTGGCTGAGCTCCTTGTCTCATTTAAGGCTTAAGAATCAACCGCCCCCCACTCACCCCCCAGCACAGGAGGTTCTGTCCAGGGTCTAAGCCCCTCATCTTCTCCCCTCCCTGCTGAGCGCCTCTGGAGTCTAAACCAACTTAACTCACTGCTAAGCCTTTCGGGAGAAGCCTGGTGGGAGGTAATAAACTGAGTAGTTACCACTTCCAACATCTGGAGGCCGCAAGCCCAGAAGCATCCGTCCTTATTAGGAGCCAGCTGGGCTGCCCTGAGCCACAGGCAGGGCCAGGAAAGCCGATCGGAGTCTGCAGCTGGAGCCCGGCCCTCGGTGAGGCCGCCTTCCCTGGGCCTGCAACGCCACTCGCTGTTGCTGGCGAGGAATCGCTCTGCCGGGAAGGCGCTGACGCACACGCGCAGAGCTCCCCACCTGGAGACCGCATGCGGAGCTCTCCAGCGCTCAGCATAGGTTGGCGCATTTACCAGAACAGTTCACAGTGAATTTCGGAGGAACTGTCTCTGCCCCACCCCCCTCCCCAGTTTTGCCTGTTTGTCTCCTAAGCTGTATTTTAATAGCATATGTTACTAGGAAAACACTTTTTAAAGTGTCAAGACACACTTCGTAGGTAAGGAAAAATAGTGGCTTCTGGTCACAACAGAAAGAAAGGTGTGGAAGGGGTTGGGGGAGGGGCTGGGCTTATGGCGCGAGAAAGGGTCTCAGTTGCAGGGAGGAGGAGAAGGCGGCGCCCCCACTTCGCGGGGCTGCGCGGCGGGAAATGGAGGCGCGGCGGGAGGCCCCGGACGCGCCCTTCCCGGACCTCGTGGCTGCGCTCTCCTTGCGCGGCCCCCTCTCGGGCCAGCCCTGTCCGCGGGGTCCCCCTGGCGGGAGCGAGGGTCCATAAAGGTCTCCTCGCAAATGAGCAGCGGTAATCCCGCCCCGAGGATCTCGCTGGGGGGAGCAGACCCCCGGGGCACGGAGAGCCGCTCATTGGCATTCCGTGCACGCGGGCCGGGGCACGGTGCGGTGCGTGTCAAGCAGGGTCGTGCGACTCGACGACTCGGGCTCGCCTGCGCCCCGGGTCGCATTCCGGGGGGCTACGGCGGGCCTCCAACGGCCAGGCCCGCACTTCATGCCAGAGAAACCGATGAGAAGATTAAAATCCCTCTGTAATTCCAGCGGGAAGATTCTTTCTCGTCATCTCTATTTGCAAAAAGCGTGATCCTGGAGATTGGAATGCAAAGACGACTGAGACCTCCCCGCCCCTCCTCGCCGGCCCCCTGCGCGCCGCCCAGACTCTGATAATTGTCCTCGGGACAGTGAGCCTCAGAGAGCGACGGAGGGCTCGAGAAAGCGGGTAGTCAAGGGGCCTTGAGAACCCGGCCGCTTCCAGCGCTCGAAGCAGGCCCCGCCATTTAAAATTCAAATACACATCTCGAGAGCTTGGAAGAGAGGCCTGGCTGTGCAAATAGTGCTTGTGAGGCCCGGTCTCTGGAGAGCCCGCGTGTGAATCGCAGGAGGAGGGGGTGCACCTGAGCAAATAGGGAGGGGGCGGCCAGAGAGCTGGAGAGAGAGTGAGCTGAGAAAGGGGAGGGGGAGCCATGGAAGTGTCCCCTCTTCCAAGAGTGTCTCCCATTTATTGCAGAGATCACAATGACAACGCAGGACCCTTTATTGGATTTTAATTAGAAAATCCACACAAGCCCCGGATTTTCACACCTCGGCCAATTCTCTGGAATGTTTATCCAGTTGCTACAACTACTACAGCTATTTTTCACGCCCCGCCCCCCAACTGGCCGACATCAAGGCCCCCGGCTGGGCTTTGAGGCGGGCGTGAGAGGGGTCCCCGCCGCGCAACTCCAATGTGCAACGCACGAGGGAGCGCCGCGCCCTGCACAGGGGATGGGAGGGAAGGAGAGTGCGCCCTGGCGCCCGAGTGAGCTCAAGGAGGCTTCCTGAGGAATCCAAGTGGAGAGCAAACACCCTCTGAATTCGTGGCGAGGTGGGGTGTGTGCTGAGCCGGGGGATGGGGTGGGAGGAAGGCGGAAGGCAAGGAGTGTAACCCGCCTCCTGCCAGAGACGCCAGCCAAACTTTGTCACCGCCAGTCCAGTCCTAAGAGAGTCGTTAAGGAATGTGTCTGCTTCTGAATTTTGTCCCGGAGGCGGTGGGGCTGGGGGAGGGAGGGCTCCGCTGGACCAGCCTGGAATCTCTCCTCCAAGCTGCGCGCATCCCCGTTCTGGGCGAGCGGGACACCAAGCGTGCGTCCCCCCTCTCCACACGTGCCTCTCTCTGGGAACTCACCACCCTGTCTCCTCTTTTCTCCCTTCTCCTCCTCTGCAGAGAACTCGGTGGCAGCCAAATCGGGCGGCTGCTTCCCGGGCTCGGCCACGGTGCACCTGGAGCGGGGCGGCACCAAGCTGGTGAAGGACCTGCGCCCCGGGGACCGCGTGCTGGCGGCCGACGACCAGGGCCGGCTGCTCTACAGCGACTTCCTCACCTTCCTGGACCGCGACGACGGCGCCAAGAAGGTCTTCTATGTGATCGAGACGAGGGAGCCGCGCGAGCGCCTTCTGCTCACCGCCGCGCACCTGCTCTTCGTCGCGCCGCACAACGACTCGGCCGCCGGGGAGCCCGGGGCGCCGTGGAGCGCGGGGCCGCCGCCGGGGGGCGTGCCGGGGCGCCGGGCGCTCTTTGCCAGCCGCGTGCGCCCGGGCCAGCGCGTGTACGTGGTGGCCGAGCGCGGCGGGGACCGCCGGCTCCTGCCTGCCGCGGTGCACAGCGTGACGCTGCGCGAGGAGGCCACGGGCGCGTACGCGCCTCTCACGGCGCAGGGCACCATCCTCATCAACAGGGTGCTGGCGTCGTGCTACGCGGTCATCGAGGAGCACGGCTGGGCGCACTGGGCCTTCGCGCCCTTCCGCCTGGCGCACGCGCTCCTGGCCGCACTGGCGCCGGCGCGCACGGACCGCGGCGGGGACGGCGACGGCGGAGGCGGCGGCCGTGTCCCTCCGCCCGCGCCAGGCGCGCCAGGTGCGCCAGGTGCGGCCGACGCGCCAGGTGCCGGAGGCGTCCACTGGTACTCGCAGCTGCTCTACCAAATAGGCACCTGGCTGTTGGACAGCGAAGCCCTGCACCCGCTCGGCATGGCGGTCAAGTCCAGCTGA

>CmR_p15a_mGL_SacB_Template (4157 bp)

agtcgaacgaccgagcgtagcgagtcagtgagcgaggaagcggaatatatccctaggtataaacgcagaaaggcccacccgaaggtgagccagTTATTTGTTAACTGTTAATTGTCCTTGTTCAAGGATGCTGTCTTTGACAACAGATGTTTTCTTGCCTTTGATGTTCAGCAGGAAGCTAGGCGCAAACGTTGATTGTTTGTCTGCGTAGAATCCTCTGTTTGTCATATAGCTTGTAATCACGACATTGTTTCCTTTCGCTTGAGGTACAGCGAAGTGTGAGTAAGTAAAGGTTACATCGTTAGGATCAAGATCCATTTTTAACACAAGGCCAGTTTTGTTCAGCGGCTTGTATGGGCCAGTTAAAGAATTAGAAACATAACCAAGCATGTAAATATCGTTAGACGTAATGCCGTCAATCGTCATTTTTGATCCGCGGGAGTCAGTGAACAGGTACCATTTGCCGTTCATTTTAAAGACGTTCGCGCGTTCAATTTCATCTGTTACTGTGTTAGATGCAATCAGCGGTTTCATCACTTTTTTCAGTGTGTAATCATCGTTTAGCTCAATCATACCGAGAGCGCCGTTTGCTAACTCAGCCGTGCGTTTTTTATCGCTTTGCAGAAGTTTTTGACTTTCTTGACGGAAGAATGATGTGCTTTTGCCATAGTATGCTTTGTTAAATAAAGATTCTTCGCCTTGGTAGCCATCTTCAGTTCCAGTGTTTGCTTCAAATACTAAGTATTTGTGGCCTTTATCTTCTACGTAGTGAGGATCTCTCAGCGTATGGTTGTCGCCTGAGCTGTAGTTGCCTTCATCGATGAACTGCTGTACATTTTGATACGTTTTTCCGTCACCGTCAAAGATTGATTTATAATCCTCTACACCGTTGATGTTCAAAGAGCTGTCTGATGCTGATACGTTAACTTGTGCAGTTGTCAGTGTTTGTTTGCCGTAATGTTTACCGGAGAAATCAGTGTAGAATAAACGGATTTTTCCGTCAGATGTAAATGTGGCTGAACCTGACCATTCTTGTGTTTGGTCTTTTAGGATAGAATCATTTGCATCGAATTTGTCGCTGTCTTTAAAGACGCGGCCAGCGTTTTTCCAGCTGTCAATAGAAGTTTCGCCGACTTTTTGATAGAACATGTAAATCGATGTGTCATCCGCATTTTTAGGATCTCCGGCTAATGCAAAGACGATGTGGTAGCCGTGATAGTTTGCGACAGTGCCGTCAGTGTTTTGTAATGGCCAGCTGTCCCAAACGTCCAGGCCTTTTGCAGAAGAGATATTTTTAATTGTGGACGAATCGAACTCAGGAACTTTATATTTTTCATTTTTTTGCTGTTCAGGGATTTGCAGCATATCATGGCGTGTAATATGGGAAATGCCGTATGTTTCCTTATATGGCTTTTGGTTCGTTTCTTTCGCAAACGCTTGAGTTGCGCCTCCTGCCAGCAGTGCGGTAGTAAAGGTTAATACTGTTGCTTGTTTTGCAAACTTTTTGATGTTCATggtttagttcctcaccttGTCGTATTATACTATGCCGATATACTATGCCGATGATTAATTGtCAACacgtgctGGTCTCCTGACCTCCTGCCAGCAATAGTAAGACAACACGCAAAGTCtgtgaTTGACAGCTAGCTCAGTCCTAGGTATAATGCTAGCaaagaggagaaaggatctatggctattacacgaagaagttttcttaaaggcgtagcaacgacaagtgccgcgtcagttatcggtccaagcttattagcgtcagcttcagcaaaagccgcgATGGTGTCGAAAGGTGAGGAGCTTTTCACGGGGGTGGTTCCAATCTTAGTCGAGCTTGATGGTGACGTGAATGGCCATAAGTTTAGCGTGCGCGGCGAGGGTGAGGGCGATGCGACGAACGGTAAACTTACTTTAAAGTTTATTTGTACTACAGGCAAATTACCGGTCCCGTGGCCGACCCTGGTCACTACGCTTGGGAATGGGGTAGCATGTTTTGCCCGTTACCCTGACCACATGAAACAGCACGATTTCTTTAAATCTGCCATGCCCGAAGGGTATGTTCAGGAGCGCACTATCAGTTTTAAAGACGACGGAACGTACAAAACTCGTGCTGAAGTAAAGTTCGAAGGTGATACCCTTGTGAACCGTATCGTCCTTAAAGGCATCGACTTCAAAGAGGACGGTAACATTTTAGGCCACAAATTAGAGTACAACTTCAATTCACATAAGGTTTATATCACAGCAGATAAGCAAAAAAATGGCATCAAGGCAAATTTTAAGACCCGCCACAACGTAGAAGATGGTGGAGTCCAGTTGGCGGACCACTACCAGCAAAATACACCGATTGGCGATGGTCCGGTTTTGCTTCCCGACAATCACTATCTTAGCCACCAGAGTAAACTGAGTAAGGACCCAAATGAAAAGCGCGACCACATGGTATTGAAAGAGCGCGTTACAGCAGCTGGAATTACACACGATATGGACGAACTTTATAAGTAAATCTATCTtgatcggcacgtaagaggttccaactttcaccataatgaaataagatcactaccgggcgtattttttgagttatcgagattttcaggagctaaggaagctaaaatggagaaaaaaatcactggatataccaccgttgatatatcccaatggcatcgtaaagaacattttgaggcatttcagtcagttgctcaatgtacctataaccagaccgttcagctggatattacggcctttttaaagaccgtaaagaaaaataagcacaagttttatccggcctttattcacattcttgcccgcctgatgaatgctcatccggaatttcgtatggcaatgaaagacggtgagctggtgatatgggatagtgttcacccttgttacaccgttttccatgagcaaactgaaacgttttcatcgctctggagtgaataccacgacgatttccggcagtttctacacatatattcgcaagatgtggcgtgttacggtgaaaacctggcctatttccctaaagggtttattgagaatatgtttttcgtctcagccaatccctgggtgagtttcaccagttttgatttaaacgtggccaatatggacaacttcttcgcccccgttttcaccatgggcaaatattatacgcaaggcgacaaggtgctgatgccgctggcgattcaggttcatcatgccgtttgtgatggcttccatgtcggcagaatgcttaatgaattacaacagtactgcgatgagtggcagggcggggcgtaatttgatatcgagctcgcttggactcctgttgatagatccagtaatgacctcagaactccatctggatttgttcagaacgctcggttgccgccgggcgttttttattggtgagaatccaagcactagtaacaacttatatcgtatggggctgacttcaggtgctacatttgaagagataaattgcactgaaatctagaaatattttatctgattaataagatgatcttcttgagatcgttttggtctgcgcgtaatctcttgctctgaaaacgaaaaaaccgccttgcagggcggtttttcgaaggttctctgagctaccaactctttgaaccgaggtaactggcttggaggagcgcagtcaccaaaacttgtcctttcagtttagccttaaccggcgcatgacttcaagactaactcctctaaatcaattaccagtggctgctgccagtggtgcttttgcatgtctttccgggttggactcaagacgatagttaccggataaggcgcagcggtcggactgaacggggggttcgtgcatacagtccagcttggagcgaactgcctacccggaactgagtgtcaggcgtggaatgagacaaacgcggccataacagcggaatgacaccggtaaaccgaaaggcaggaacaggagagcgcacgagggagccgccagggggaaacgcctggtatctttatagtcctgtcgggtttcgccaccactgatttgagcgtcagatttcgtgatgcttgtcaggggggcggagcctatggaaaaacggctttgccgcggccctctcacttccctgttaagtatcttcctggcatcttccaggaaatctccgccccgttcgtaagccatttccgctcgccgc

>KanR_pSC101_SacB_Template (4297 bp)

gagttatacacagggctgggatctattctttttatctttttttattctttctttattctataaattataaccacttgaatataaacaaaaaaaacacacaaaggtctagcggaatttacagagggtctagcagaatttacaagttttccagcaaaggtctagcagaatttacagatacccacaactcaaaggaaaaggactagtaattatcattgactagcccatctcaattggtatagtgattaaaatcacctagaccaattgagatgtatgtctgaattagttgttttcaaagcaaatgaactagcgattagtcgctatgacttaacggagcatgaaaccaagctaattttatgctgtgtggcactactcaaccccacgattgaaaaccctacaaggaaagaacggacggtatcgttcacttataaccaatacgctcagatgatgaacatcagtagggaaaatgcttatggtgtattagctaaagcaaccagagagctgatgacgagaactgtggaaatcaggaatcctttggttaaaggctttgagattttccagtggacaaactatgccaagttctcaagcgaaaaattagaattagtttttagtgaagagatattgccttatcttttccagttaaaaaaattcataaaatataatctggaacatgttaagtcttttgaaaacaaatactctatgaggatttatgagtggttattaaaagaactaacacaaaagaaaactcacaaggcaaatatagagattagccttgatgaatttaagttcatgttaatgcttgaaaataactaccatgagtttaaaaggcttaaccaatgggttttgaaaccaataagtaaagatttaaacacttacagcaatatgaaattggtggttgataagcgaggccgcccgactgatacgttgattttccaagttgaactagatagacaaatggatctcgtaaccgaacttgagaacaaccagataaaaatgaatggtgacaaaataccaacaaccattacatcagattcctacctacataacggactaagaaaaacactacacgatgctttaactgcaaaaattcagctcaccagttttgaggcaaaatttttgagtgacatgcaaagtaagtatgatctcaatggttcgttctcatggctcacgcaaaaacaacgaaccacactagagaacatactggctaaatacggaaggatctgaATCCATGCCTGAGACCagcacgtGTTGaCAATTAATCATCGGCATAGTATATCGGCATAGTATAATACGACaaggtgaggaactaaaccATGAACATCAAAAAGTTTGCAAAACAAGCAACAGTATTAACCTTTACTACCGCACTGCTGGCAGGAGGCGCAACTCAAGCGTTTGCGAAAGAAACGAACCAAAAGCCATATAAGGAAACATACGGCATTTCCCATATTACACGCCATGATATGCTGCAAATCCCTGAACAGCAAAAAAATGAAAAATATAAAGTTCCTGAGTTCGATTCGTCCACAATTAAAAATATCTCTTCTGCAAAAGGCCTGGACGTTTGGGACAGCTGGCCATTACAAAACACTGACGGCACTGTCGCAAACTATCACGGCTACCACATCGTCTTTGCATTAGCCGGAGATCCTAAAAATGCGGATGACACATCGATTTACATGTTCTATCAAAAAGTCGGCGAAACTTCTATTGACAGCTGGAAAAACGCTGGCCGCGTCTTTAAAGACAGCGACAAATTCGATGCAAATGATTCTATCCTAAAAGACCAAACACAAGAATGGTCAGGTTCAGCCACATTTACATCTGACGGAAAAATCCGTTTATTCTACACTGATTTCTCCGGTAAACATTACGGCAAACAAACACTGACAACTGCACAAGTTAACGTATCAGCATCAGACAGCTCTTTGAACATCAACGGTGTAGAGGATTATAAATCAATCTTTGACGGTGACGGAAAAACGTATCAAAATGTACAGCAGTTCATCGATGAAGGCAACTACAGCTCAGGCGACAACCATACGCTGAGAGATCCTCACTACGTAGAAGATAAAGGCCACAAATACTTAGTATTTGAAGCAAACACTGGAACTGAAGATGGCTACCAAGGCGAAGAATCTTTATTTAACAAAGCATACTATGGCAAAAGCACATCATTCTTCCGTCAAGAAAGTCAAAAACTTCTGCAAAGCGATAAAAAACGCACGGCTGAGTTAGCAAACGGCGCTCTCGGTATGATTGAGCTAAACGATGATTACACACTGAAAAAAGTGATGAAACCGCTGATTGCATCTAACACAGTAACAGATGAAATTGAACGCGCGAACGTCTTTAAAATGAACGGCAAATGGTACCTGTTCACTGACTCCCGCGGATCAAAAATGACGATTGACGGCATTACGTCTAACGATATTTACATGCTTGGTTATGTTTCTAATTCTTTAACTGGCCCATACAAGCCGCTGAACAAAACTGGCCTTGTGTTAAAAATGGATCTTGATCCTAACGATGTAACCTTTACTTACTCACACTTCGCTGTACCTCAAGCGAAAGGAAACAATGTCGTGATTACAAGCTATATGACAAACAGAGGATTCTACGCAGACAAACAATCAACGTTTGCGCCTAGCTTCCTGCTGAACATCAAAGGCAAGAAAACATCTGTTGTCAAAGACAGCATCCTTGAACAAGGACAATTAACAGTTAACAAATAAGGTACCCTCGAGtttgtttatttttctaaatacattcaaatatgtatccgctcatgagacaataaccctgataaatgcttcaataatattgaaaaaggaagagtATGAGCCATATTCAACGGGAAACGTCTTGCTCTAGGCCGCGATTAAATTCCAACATGGATGCTGATTTATATGGGTATAAATGGGCTCGCGATAATGTCGGGCAATCAGGTGCGACAATCTATCGATTGTATGGGAAGCCCGATGCGCCAGAGTTGTTTCTGAAACATGGCAAAGGTAGCGTTGCCAATGATGTTACAGATGAGATGGTCAGACTAAACTGGCTGACGGAATTTATGCCTCTTCCGACCATCAAGCATTTTATCCGTACTCCTGATGATGCATGGTTACTCACCACTGCGATCCCCGGGAAAACAGCATTCCAGGTATTAGAAGAATATCCTGATTCAGGTGAAAATATTGTTGATGCGCTGGCAGTGTTCCTGCGCCGGTTGCATTCGATTCCTGTTTGTAATTGTCCTTTTAACAGCGACCGCGTATTTCGTCTCGCTCAGGCGCAATCACGAATGAATAACGGTTTGGTTGATGCGAGTGATTTTGATGACGAGCGTAATGGCTGGCCTGTTGAACAAGTCTGGAAAGAAATGCATAAACTTTTGCCATTCTCACCGGATTCAGTCGTCACTCATGGTGATTTCTCACTTGATAACCTTATTTTTGACGAGGGGAAATTAATAGGTTGTATTGATGTTGGACGAGTCGGAATCGCAGACCGATACCAGGATCTTGCCATCCTATGGAACTGCCTCGGTGAGTTTTCTCCTTCATTACAGAAACGGCTTTTTCAAAAATATGGTATTGATAATCCTGATATGAATAAATTGCAGTTTCATTTGATGCTCGATGAGTTTTTCTAATTAATTAAACTAGTGAtgtctaacaattcgttcaagccgacggatatcgagctcgcttggactcctgttgatagatccagtaatgacctcagaactccatctggatttgttcagaacgctcggttgccgccgggcgttttttattggtgagaatccaagcactagggacagtaagacgggtaagcctgttgatgataccgctgccttactgggtgcattagccagtctgaatgacctgtcacgggataatccgaagtggtcagactggaaaatcagagggcaggaactgctgaacagcaaaaagtcagatagcaccacatagcagacccgccataaaacgccctgagaagcccgtgacgggcttttcttgtattatgggtagtttccttgcatgaatccataaaaggcgcctgtagtgccatttacccccattcactgccagagccgtgagcgcagcgaactgaatgtcacgaaaaagacagcgactcaggtgcctgatggtcggagacaaaaggaatattcagcgatttgcccgagcttgcgagggtgctacttaagcctttagggttttaaggtctgttttgtagaggagcaaacagcgtttgcgacatccttttgtaatactgcggaactgactaaagtagt

>pSC101_pBAD_HK022_Recombinase

ATTAAGTTACATTGTCGATCTGTTCATGGTGAACAGCTTTAAATGCACCAAAAACTCGTAAAAGCTCTGATGTATCTATCTTTTTTACACCGTTTTCATCTGTGCATATGGACAGTTTTCCCTTTGATATCTAACGGTGAACAGTTGTTCTACTTTTGTTTGTTAGTCTTGATGCTTCACTGATAGATACAAGAGCCATAAGAACCTCAGATCCTTCCGTATTTAGCCAGTATGTTCTCTAGTGTGGTTCGTTGTTTTTGCGTGAGCCATGAGAACGAACCATTGAGATCATGCTTACTTTGCATGTCACTCAAAAATTTTGCCTCAAAACTGGTGAGCTGAATTTTTGCAGTTAAAGCATCGTGTAGTGTTTTTCTTAGTCCGTTACGTAGGTAGGAATCTGATGTAATGGTTGTTGGTATTTTGTCACCATTCATTTTTATCTGGTTGTTCTCAAGTTCGGTTACGAGATCCATTTGTCTATCTAGTTCAACTTGGAAAATCAACGTATCAGTCGGGCGGCCTCGCTTATCAACCACCAATTTCATATTGCTGTAAGTGTTTAAATCTTTACTTATTGGTTTCAAAACCCATTGGTTAAGCCTTTTAAACTCATGGTAGTTATTTTCAAGCATTAACATGAACTTAAATTCATCAAGGCTAATCTCTATATTTGCCTTGTGAGTTTTCTTTTGTGTTAGTTCTTTTAATAACCACTCATAAATCCTCATAGAGTATTTGTTTTCAAAAGACTTAACATGTTCCAGATTATATTTTATGAATTTTTTTAACTGGAAAAGATAAGGCAATATCTCTTCACTAAAAACTAATTCTAATTTTTCGCTTGAGAACTTGGCATAGTTTGTCCACTGGAAAATCTCAAAGCCTTTAACCAAAGGATTCCTGATTTCCACAGTTCTCGTCATCAGCTCTCTGGTTGCTTTAGCTAATACACCATAAGCATTTTCCCTACTGATGTTCATCATCTGAGCGTATTGGTTATAAGTGAACGATACCGTCCGTTCTTTCCTTGTAGGGTTTTCAATCGTGGGGTTGAGTAGTGCCACACAGCATAAAATTAGCTTGGTTTCATGCTCCGTTAAGTCATAGCGACTAATCGCTAGTTCATTTGCTTTGAAAACAACTAATTCAGACATACATCTCAATTGGTCTAGGTGATTTTAATCACTATACCAATTGAGATGGGCTAGTCAATGATAATTACTAGTCCTTTTCCTTTGAGTTGTGGGTATCTGTAAATTCTGCTAGACCTTTGCTGGAAAACTTGTAAATTCTGCTAGACCCTCTGTAAATTCCGCTAGACCTTTGTGTGTTTTTTTTGTTTATATTCAAGTGGTTATAATTTATAGAATAAAGAAAGAATAAAAAAAGATAAAAAGAATAGATCCCAGCCCTGTGTATAACTCACTACTTTAGTCAGTTCCGCAGTATTACAAAAGGATGTCGCAAACGCTGTTTGCTCCTCTACAAAACAGACCTTAAAACCCTAAAGGCTTAAGTAGCACCCTCGCAAGCTCGGGCAAATCGCTGAATATTCCTTTTGTCTCCGACCATCAGGCACCTGAGTCGCTGTCTTTTTCGTGACATTCAGTTCGCTGCGCTCACGGCTCTGGCAGTGAATGGGGGTAAATGGCACTACAGGCGCCTTTTATGGATTCATGCAAGGAAACTACCCATAATACAAGAAAAGCCCGTCACGGGCTTCTCAGGGCGTTTTATGGCGGGTCTGCTATGTGGTGCTATCTGACTTTTTGCTGTTCAGCAGTTCCTGCCCTCTGATTTTCCAGTCTGACCACTTCGGATTATCCCGTGACAGGTCATTCAGACTGGCTAATGCACCCAGTAAGGCAGCGGTATCATCAACAGGCTTACCCGTCTTACTGTCCCTAGTGCTTGGATTCTCACCAATAAAAAACGCCCGGCGGCAACCGAGCGTTCTGAACAAATCCAGATGGAGTTCTGAGGTCATTACTGGATCTATCAACAGGAGTCCAAGCGAGCTCGTTACCAATGCTTAATCAGTGAGGCACCTATCTCAGCGATCTGTCTATTTCGTTCATCCATAGTTGCCTGACTCCCCGTCGTGTAGATAACTACGATACGGGAGGGCTTACCATCTGGCCCCAGTGCTGCAATGATACCGCGAGACCCACGCTCACCGGCTCCAGATTTATCAGCAATAAACCAGCCAGCCGGAAGGGCCGAGCGCAGAAGTGGTCCTGCAACTTTATCCGCCTCCATCCAGTCTATTAATTGTTGCCGGGAAGCTAGAGTAAGTAGTTCGCCAGTTAATAGTTTGCGCAACGTTGTTGCCATTGCTGCAGGCATCGTGGTGTCACGCTCGTCGTTTGGTATGGCTTCATTCAGCTCCGGTTCCCAACGATCAAGGCGAGTTACATGATCCCCCATGTTGTGCAAAAAAGCGGTTAGCTCCTTCGGTCCTCCGATCGTTGTCAGAAGTAAGTTGGCCGCAGTGTTATCACTCATGGTTATGGCAGCACTGCATAATTCTCTTACTGTCATGCCATCCGTAAGATGCTTTTCTGTGACTGGTGAGTACTCAACCAAGTCATTCTGAGAATAGTGTATGCGGCGACCGAGTTGCTCTTGCCCGGCGTCAACACGGGATAATACCGCGCCACATAGCAGAACTTTAAAAGTGCTCATCATTGGAAAACGTTCTTCGGGGCGAAAACTCTCAAGGATCTTACCGCTGTTGAGATCCAGTTCGATGTAACCCACTCGTGCACCCAACTGATCTTCAGCATCTTTTACTTTCACCAGCGTTTCTGGGTGAGCAAAAACAGGAAGGCAAAATGCCGCAAAAAAGGGAATAAGGGCGACACGGAAATGTTGAATACTCATACTCTTCCTTTTTCAATATTATTGAAGCATTTATCAGGGTTATTGTCTCATGAGCGGATACATATTTGAATGTATTTAGAAAAATAAACAAATAGGGGTTCCGCGCTGCAGCCAATTATTGAAGGCCGCTAACGCGGCCTTTTTTTGTTTCTGGTCTCCCGGCCGGCCCTCGGTACCAAATTCCAGAAAAGAGGCCTCCCGAAAGGGGGGCCTTTTTTCGTTTTGGTCCGGCGCGCCTTATGACAACTTGACGGCTACATCATTCACTTTTTCTTCACAACCGGCACGGAACTCGCTCGGGCTGGCCCCGGTGCATTTTTTAAATACCCGCGAGAAATAGAGTTGATCGTCAAAACCAACATTGCGACCGACGGTGGCGATAGGCATCCGGGTGGTGCTCAAAAGCAGCTTCGCCTGGCTGATACGTTGGTCCTCGCGCCAGCTTAAGACGCTAATCCCTAACTGCTGGCGGAAAAGATGTGACAGACGCGACGGCGACAAGCAAACATGCTGTGCGACGCTGGCGATATCAAAATTGCTGTCTGCCAGGTGATCGCTGATGTACTGACAAGCCTCGCGTACCCGATTATCCATCGGTGGATGGAGCGACTCGTTAATCGCTTCCATGCGCCGCAGTAACAATTGCTCAAGCAGATTTATCGCCAGCAGCTCCGAATAGCGCCCTTCCCCTTGCCCGGCGTTAATGATTTGCCCAAACAGGTCGCTGAAATGCGGCTGGTGCGCTTCATCCGGGCGAAAGAACCCCGTATTGGCAAATATTGACGGCCAGTTAAGCCATTCATGCCAGTAGGCGCGCGGACGAAAGTAAACCCACTGGTGATACCATTCGCGAGCCTCCGGATGACGACCGTAGTGATGAATCTCTCCTGGCGGGAACAGCAAAATATCACCCGGTCGGCAAACAAATTCTCGTCCCTGATTTTTCACCACCCCCTGACCGCGAATGGTGAGATTGAGAATATAACCTTTCATTCCCAGCGGTCGGTCGATAAAAAAATCGAGATAACCGTTGGCCTCAATCGGCGTTAAACCCGCCACCAGATGGGCATTAAACGAGTATCCCGGCAGCAGGGGATCATTTTGCGCTTCAGCCATACTTTTCATACTCCCGCCATTCAGAGAAGAAACCAATTGTCCATATTGCATCAGACATTGCCGTCACTGCGTCTTTTACTGGCTCTTCTCGCTAACCAAACCGGTAACCCCGCTTATTAAAAGCATTCTGTAACAAAGCGGGACCAAAGCCATGACAAAAACGCGTAACAAAAGTGTCTATAATCACGGCAGAAAAGTCCACATTGATTATTTGCACGGCGTCACACTTTGCTATGCCATAGCATTTTTATCCATAAGATTAGCGGATCCTACCTGACGCTTTTTATCGCAACTCTCTACTGTTTCTCCATACCCGTTTTTTTGGGCTAACAGGAGGAATTAACCATGGGAAGAAGGCGAAGTCATGAGCGCCGGGATTTACCCCCTAACCTTTATATAAGAAACAATGGATATTACTGCTACAGGGACCCAAGGACGGGTAAAGAGTTTGGATTAGGCCGAGACAGGCGAATCGCAATCACTGAAGCTATACAGGCCAACATTGAGTTGCTATCCGGGAACAGGCGTGAGTCACTGATAGACAGAATTAAAGGCGCTGACGCAATCACTCTTCATGCGTGGCTTGACCGATATGAAACAATCCTCAGCGAGAGGGGTATCAGGCCGAAAACTCTACTCGACTACGCCAGCAAAATCAGAGCAATCCGAAGAAAATTGCCGGACAAACCGCTCGCTGACATATCAACGAAAGAGGTGGCAGCAATGCTAAACACCTACGTAGCAGAAGGCAAATCGGCTTCCGCAAAATTGATCAGGTCAACCCTTGTTGACGTTTTTCGTGAGGCAATAGCCGAGGGGCATGTGGCTACGAATCCGGTAACAGCAACCCGCACAGCAAAATCAGAAGTAAGGCGTTCAAGACTGACAGCTAATGAATATGTCGCGATATACCATGCAGCCGAACCGCTCCCAATCTGGCTGAGACTGGCAATGGATTTGGCTGTCGTTACAGGGCAGAGAGTAGGCGATTTGTGCAGAATGAAATGGTCAGACATAAACGACAACCATCTTCACATTGAACAGAGTAAAACAGGGGCTAAACTCGCCATTCCACTAACGCTAACGATTGACGCGCTCAATATCTCATTGGCTGATACACTACAGCAATGCAGGGAGGCCAGCAGCAGTGAAACTATAATCGCATCAAAGCATCACGATCCGCTTTCCCCGAAAACAGTATCAAAGTATTTTACAAAGGCGAGAAATGCATCTGGACTCTCATTTGATGGAAACCCGCCAACATTCCATGAACTGCGTAGCCTGTCAGCGAGGCTATACCGGAACCAGATTGGCGATAAGTTTGCTCAACGTCTTCTCGGGCATAAATCAGATTCAATGGCGGCGCGGTATAGGGACAGCCGTGGACGGGAATGGGACAAAATTGAAATCGACAAATGATCTCAGTGGAACGAAAACTCACGTTAAGGGATTCTGTGGGATCCAAACCTAGCATAACCCCTTGGGGCCTCTAAACGGGTCTTGAGGGGTTTTTTGGCGGCCGCGGCTTGTAGATATGACGACAGGAAGAGTGAATTGATCTGCTGCCTCGCGCGTTTCTTA

>Genomically_Integrated_attB_*dif*_landing_site_orientation1 (2063 bp)

CTTAAATAAGTAACAGCACCCTCTGGTGTTTTCGTATGCAGTGTCTATGCGACAGACACTGCGCTCGTATGTGACCGTAGAGTATTCTTAGGTGGCAGCGAACGAGAGATCTTTACGCCCCGCCCTGCCACTCATCGCAGTACTGTTGTAATTCATTAAGCATTCTGCCGACATGGAAGCCATCACAGACGGCATGATGAACCTGAATCGCCAGCGGCATCAGCACCTTGTCGCCTTGCGTATAATATTTGCCCACGGTGAAAACGGGGGCGAAGAAGTTGTCCATATTGGCCACGTTTAAATCAAAACTGGTGAAACTCACCCAGGGATTGGCTGAGACGAAAAACATATTCTCAATAAACCCTTTAGGGAAATAGGCCAGGTTTTCACCGTAACACGCCACATCTTGCGAATATATGTGTAGAAACTGCCGGAAATCGTCGTGGTATTCACTCCAGAGCGATGAAAACGTTTCAGTTTGCTCATGGAAAACGGTGTAACAAGGGTGAACACTATCCCATATCACCAGCTCACCGTCTTTCATTGCCATACGGAATTCCGGATGAGCATTCATCAGGCGGGCAAGAATGTGAATAAAGGCCGGATAAAACTTGTGCTTATTTTTCTTTACGGTCTTTAAAAAGGCCGTAATATCCAGCTGAACGGTCTGGTTATAGGTACATTGAGCAACTGACTGAAATGCCTCAAAATGTTCTTTACGATGCCATTGGGATATATCAACGGTGGTATATCCAGTGATTTTTTTCTCCATGATTGTCCTCCTTCATATGGTCGTATTATACTATGCCGATATACTATGCCGATGATTAATTGTCAACATGCATTTACTTATAAAGTTCGTCCATATCGTGTGTAATTCCAGCTGCTGTAACGCGCTCTTTCAATACCATGTGGTCGCGCTTTTCATTTGGGTCCTTACTCAGTTTACTCTGGTGGCTAAGATAGTGATTGTCGGGAAGCAAAACCGGACCATCGCCAATCGGTGTATTTTGCTGGTAGTGGTCCGCCAACTGGACTCCACCATCTTCTACGTTGTGGCGGGTCTTAAAATTTGCCTTGATGCCATTTTTTTGCTTATCTGCTGTGATATAAACCTTATGTGAATTGAAGTTGTACTCTAATTTGTGGCCTAAAATGTTACCGTCCTCTTTGAAGTCGATGCCTTTAAGGACGATACGGTTCACAAGGGTATCACCTTCGAACTTTACTTCAGCACGAGTTTTGTACGTTCCGTCGTCTTTAAAACTGATAGTGCGCTCCTGAACATACCCTTCGGGCATGGCAGATTTAAAGAAATCGTGCTGTTTCATGTGGTCAGGGTAACGGGCAAAACATGCTACCCCATACCCAAGCGTAGTGACCAGGGTCGGCCACGGGACCGGTAATTTGCCTGTAGTACAAATAAACTTTAAAGTAAGTTTACCGTTCGTCGCATCGCCCTCACCCTCGCCGCGCACGCTAAACTTATGGCCATTCACGTCACCATCAAGCTCGACTAAGATTGGAACCACCCCCGTGAAAAGCTCCTCACCTTTCGACACCATCTAGTATTTCCCCTCTTTCTCTAGTATTAAACAAAATTATTTGTAGAGGCTGTTTCGTCCTCACGGACTCATCAGACCGGAAAGCACATCCGGTGACAGCTGCAGTGCAAATGATTTTATTTTGACTAATAATGACCTACTTACATTAATTTACTGATAATTAAAGAGATTTTAAATATACAACTTATTCACCTAAAGGATGACAAAATAACATTAATCACTTAAAAATCATCGCATTACACTAATCTGTGGTTAAATGATAGACTACATAATGCGACAAAACGCAACATATCCAGTCACTATGAATCAACTACTTAGATAGTATTAGTGACCTTTGGTCTAGAGCTAGCATTATACCTAGGACTGAGCTAGCTGTCAAATCGATGGCGCGCCCTCTAACGGACTTGAGTGAGGTTGTAAAGGGAGTTGGCTCACCCCATATCTCACTTACTACCGTATTCTCGGCTTAACCGTCCATGCAACCTCAACACATTGCTT

>Genomically_Integrated_attB_*dif*_landing_site_orientation2 (2062 bp)

TTAAATAAGTAACAGCACCCTCTGGTGTTTTCGTATGCAGTGTCTATGCGACAGACACTGCGCTCGAGCCAACTCCCTTTACAACCTCACTCAAGTCCGTTAGAGGGCGCGCCATCGATTTGACAGCTAGCTCAGTCCTAGGTATAATGCTAGCTCTAGACCAAAGGTCACTAATACTATCTAAGTAGTTGATTCATAGTGACTGGATATGTTGCGTTTTGTCGCATTATGTAGTCTATCATTTAACCACAGATTAGTGTAATGCGATGATTTTTAAGTGATTAATGTTATTTTGTCATCCTTTAGGTGAATAAGTTGTATATTTAAAATCTCTTTAATTATCAGTAAATTAATGTAAGTAGGTCATTATTAGTCAAAATAAAATCATTTGCACTGCAGCTGTCACCGGATGTGCTTTCCGGTCTGATGAGTCCGTGAGGACGAAACAGCCTCTACAAATAATTTTGTTTAATACTAGAGAAAGAGGGGAAATACTAGATGGTGTCGAAAGGTGAGGAGCTTTTCACGGGGGTGGTTCCAATCTTAGTCGAGCTTGATGGTGACGTGAATGGCCATAAGTTTAGCGTGCGCGGCGAGGGTGAGGGCGATGCGACGAACGGTAAACTTACTTTAAAGTTTATTTGTACTACAGGCAAATTACCGGTCCCGTGGCCGACCCTGGTCACTACGCTTGGGTATGGGGTAGCATGTTTTGCCCGTTACCCTGACCACATGAAACAGCACGATTTCTTTAAATCTGCCATGCCCGAAGGGTATGTTCAGGAGCGCACTATCAGTTTTAAAGACGACGGAACGTACAAAACTCGTGCTGAAGTAAAGTTCGAAGGTGATACCCTTGTGAACCGTATCGTCCTTAAAGGCATCGACTTCAAAGAGGACGGTAACATTTTAGGCCACAAATTAGAGTACAACTTCAATTCACATAAGGTTTATATCACAGCAGATAAGCAAAAAAATGGCATCAAGGCAAATTTTAAGACCCGCCACAACGTAGAAGATGGTGGAGTCCAGTTGGCGGACCACTACCAGCAAAATACACCGATTGGCGATGGTCCGGTTTTGCTTCCCGACAATCACTATCTTAGCCACCAGAGTAAACTGAGTAAGGACCCAAATGAAAAGCGCGACCACATGGTATTGAAAGAGCGCGTTACAGCAGCTGGAATTACACACGATATGGACGAACTTTATAAGTAAATGCATGTTGACAATTAATCATCGGCATAGTATATCGGCATAGTATAATACGACCATATGAAGGAGGACAATCATGGAGAAAAAAATCACTGGATATACCACCGTTGATATATCCCAATGGCATCGTAAAGAACATTTTGAGGCATTTCAGTCAGTTGCTCAATGTACCTATAACCAGACCGTTCAGCTGGATATTACGGCCTTTTTAAAGACCGTAAAGAAAAATAAGCACAAGTTTTATCCGGCCTTTATTCACATTCTTGCCCGCCTGATGAATGCTCATCCGGAATTCCGTATGGCAATGAAAGACGGTGAGCTGGTGATATGGGATAGTGTTCACCCTTGTTACACCGTTTTCCATGAGCAAACTGAAACGTTTTCATCGCTCTGGAGTGAATACCACGACGATTTCCGGCAGTTTCTACACATATATTCGCAAGATGTGGCGTGTTACGGTGAAAACCTGGCCTATTTCCCTAAAGGGTTTATTGAGAATATGTTTTTCGTCTCAGCCAATCCCTGGGTGAGTTTCACCAGTTTTGATTTAAACGTGGCCAATATGGACAACTTCTTCGCCCCCGTTTTCACCGTGGGCAAATATTATACGCAAGGCGACAAGGTGCTGATGCCGCTGGCGATTCAGGTTCATCATGCCGTCTGTGATGGCTTCCATGTCGGCAGAATGCTTAATGAATTACAACAGTACTGCGATGAGTGGCAGGGCGGGGCGTAAAGATCTCTCGTTCGCTGCCACCTAAGAATACTCTACGGTCACATACACCCCATATCTCACTTACTACCGTATTCTCGGCTTAACCGTCCATGCAACCTCAACACATTGCTT

>LUX_attB_Assembled_Plasmid_for_Integration (7299 bp)

CATTACTCGCATCCATTCTCAGGCTGTCTCGTCTCGTCTCCTCGAGGCCGCGCGAAAAAACCCCGCCCTGTCAGGGGCGGGGTTTTTTGCGGGCCGGCCCTCGGTACCAAATTCCAGAAAAGAGGCCTCCCGAAAGGGGGGCCTTTTTTCGTTTTGGTCCGGCGCGCCATCGATGTGCACTTTAGGTGAAAAAGGTTGAGACATGTAAGAAAGAGGAGAAATACTAGATGATTGAACAAGATGGATTGCACGCAGGTTCTCCGGCCGCTTGGGTGGAGAGGCTATTCGGCTATGACTGGGCACAACAGACAATCGGCTGCTCTGATGCCGCCGTGTTCCGGCTGTCAGCGCAGGGGCGCCCGGTTCTTTTTGTCAAGACCGACCTGTCCGGTGCCCTGAATGAACTGCAGGACGAGGCAGCGCGGCTATCGTGGCTGGCCACGACGGGCGTTCCTTGCGCAGCTGTGCTCGACGTTGTCACTGAAGCGGGAAGGGACTGGCTGCTATTGGGCGAAGTGCCGGGGCAGGATCTCCTGTCATCTCACCTTGCTCCTGCCGAGAAAGTGTCAATCATGGCTGATGCAATGCGGCGGCTGCATACGCTTGATCCGGCTACCTGCCCATTCGACCACCAAGCGAAACATCGCATCGAGCGAGCACGTACTCGGATGGAAGCCGGTCTTGTCGATCAGGATGATCTGGACGAAGAGCATCAGGGGCTCGCGCCAGCCGAACTGTTCGCCAGGCTCAAGGCGCGCATGCCCGACGGCGAGGATCTCGTCGTGACCCATGGCGATGCCTGCTTGCCGAATATCATGGTGGAAAATGGCCGCTTTTCTGGATTCATCGACTGTGGCCGGCTGGGTGTGGCGGACCGCTATCAGGACATAGCGTTGGCTACCCGTGATATTGCTGAAGAGCTTGGCGGCGAATGGGCTGACCGCTTCCTCGTGCTTTACGGTATCGCCGCTCCCGATTCGCAGCGCATCGCCTTCTATCGCCTTCTTGACGAGTTCTTCTGATAAAGCTTAAAAAAAAACCCCGCTTCGGCGGGGTTTTTTTTTTGTACAGTTATTTGCCGACTACCTTGGTGATCTCGCCTTTCACGTAGTGGACAAATTCTTCCAACTGATCTGCGCGCGAGGCCAAGCGATCTTCTTCTTGTCCAAGATCCATGCCTGAATTCGCGGCCGCTTCTAGAGGAAATCATAAAAAATTTATTTGCTTTGTGAGCGGATAACAATTATAATAGATTCAATTGTGAGCGGATAACAATTACTAGAGAAAGAGGAGAAATACTAGATGAAATTTGGAAACTTTTTGCTTACATACCAACCTCCCCAATTTTCTCAAACAGAGGTAATGAAACGTTTGGTTAAATTAGGTCGCATCTCTGAGGAGTGTGGTTTTGATACCGTATGGTTACTGGAGCATCATTTCACGGAGTTTGGTTTGCTTGGTAACCCTTATGTCGCTGCTGCATATTTACTTGGCGCGACTAAAAAATTGAATGTAGGAACTGCCGCTATTGTTCTTCCCACAGCCCATCCAGTACGCCAACTTGAAGATGTGAATTTATTGGATCAAATGTCAAAAGGACGATTTCGGTTTGGTATTTGCCGAGGGCTTTACAACAAGGACTTTCGCGTATTCGGCACAGATATGAATAACAGTCGCGCCTTAGCGGAATGCTGGTACGGGCTGATAAAGAATGGCATGACAGAGGGATATATGGAAGCTGATAATGAACATATCAAGTTCCATAAGGTAAAAGTAAACCCCGCGGCGTATAGCAGAGGTGGCGCACCGGTTTATGTGGTGGCTGAATCAGCTTCGACGACTGAGTGGGCTGCTCAATTTGGCCTACCGATGATATTAAGTTGGATTATAAATACTAACGAAAAGAAAGCACAACTTGAGCTTTATAATGAAGTGGCTCAAGAATATGGGCACGATATTCATAATATCGACCATTGCTTATCATATATAACATCTGTAGATCATGACTCAATTAAAGCGAAAGAGATTTGCCGGAAATTTCTGGGGCATTGGTATGATTCTTATGTGAATGCTACGACTATTTTTGATGATTCAGACCAAACAAGAGGTTATGATTTCAATAAAGGGCAGTGGCGTGACTTTGTATTAAAAGGACATAAAGATACTAATCGCCGTATTGATTACAGTTACGAAATCAATCCCGTGGGAACGCCGCAGGAATGTATTGACATAATTCAAAAAGACATTGATGCTACAGGAATATCAAATATTTGTTGTGGATTTGAAGCTAATGGAACAGTAGACGAAATTATTGCTTCCATGAAGCTCTTCCAGTCTGATGTCATGCCATTTCTTAAAGAAAAACAACGTTCGCTATTATATTAATAATACTAGAGAAAGAGGAGAAATACTAGATGAAATTTGGATTGTTCTTCCTTAACTTCATCAATTCAACAACTGTTCAAGAACAAAGTATAGTTCGCATGCAGGAAATAACGGAGTATGTTGATAAGTTGAATTTTGAACAGATTTTAGTGTATGAAAATCATTTTTCAGATAATGGTGTTGTCGGCGCTCCTCTGACTGTTTCTGGTTTTCTGCTCGGTTTAACAGAGAAAATTAAAATTGGTTCATTAAATCACATCATTACAACTCATCATCCTGTCCGCATAGCGGAGGAAGCTTGCTTATTGGATCAGTTAAGTGAAGGGAGATTTATTTTAGGGTTTAGTGATTGCGAAAAAAAAGATGAAATGCATTTTTTTAATCGCCCGGTTGAATATCAACAGCAACTATTTGAAGAGTGTTATGAAATCATTAACGATGCTTTAACAACAGGCTATTGTAATCCAGATAACGATTTTTATAGCTTCCCTAAAATATCTGTAAATCCCCATGCTTATACGCCAGGCGGACCTCGGAAATATGTAACAGCAACCAGTCATCATATTGTTGAGTGGGCGGCCAAAAAAGGTATTCCTCTCATCTTTAAGTGGGATGATTCTAATGATGTTAGATATGAATATGCTGAAAGATATAAAGCCGTTGCGGATAAATATGACGTTGACCTATCAGAGATAGACCATCAGTTAATGATATTAGTTAACTATAACGAAGATAGTAATAAAGCTAAACAAGAGACGCGTGCATTTATTAGTGATTATGTTCTTGAAATGCACCCTAATGAAAATTTCGAAAATAAACTTGAAGAAATAATTGCAGAAAACGCTGTCGGAAATTATACGGAGTGTATAACTGCGGCTAAGTTGGCAATTGAAAAGTGTGGTGCGAAAAGTGTATTGCTGTCCTTTGAACCAATGAATGATTTGATGAGCCAAAAAAATGTAATCAATATTGTTGATGATAATATTAAGAAGTACCACATGGAATATACCTAATAATACTAGAGAAAGAGGAGAAATACTAGATGACTAAAAAAATTTCATTCATTATTAACGGCCAGGTTGAAATCTTTCCCGAAAGTGATGATTTAGTGCAATCCATTAATTTTGGTGATAATAGTGTTTACCTGCCAATATTGAATGACTCTCATGTAAAAAACATTATTGATTGTAATGGAAATAACGAATTACGGTTGCATAACATTGTCAATTTTCTCTATACGGTAGGGCAAAGATGGAAAAATGAAGAATACTCAAGACGCAGGACATACATTCGTGACTTAAAAAAATATATGGGATATTCAGAAGAAATGGCTAAGCTAGAGGCCAATTGGATATCTATGATTTTATGTTCTAAAGGCGGCCTTTATGATGTTGTAGAAAATGAACTTGGTTCTCGCCATATCATGGATGAATGGCTACCTCAGGATGAAAGTTATGTTCGGGCTTTTCCGAAAGGTAAATCTGTACATCTGTTGGCAGGTAATGTTCCATTATCTGGGATCATGTCTATATTACGCGCAATTTTAACTAAGAATCAGTGTATTATAAAAACATCGTCAACCGATCCTTTTACCGCTAATGCATTAGCGTTAAGTTTTATTGATGTAGACCCTAATCATCCGATAACGCGCTCTTTATCTGTTATATATTGGCCCCACCAAGGTGATACATCACTCGCAAAAGAAATTATGCGACATGCGGATGTTATTGTCGCTTGGGGAGGGCCAGATGCGATTAATTGGGCGGTAGAGCATGCGCCATCTTATGCTGATGTGATTAAATTTGGTTCTAAAAAGAGTCTTTGCATTATCGATAATCCTGTTGATTTGACGTCCGCAGCGACAGGTGCGGCTCATGATGTTTGTTTTTACGATCAGCGAGCTTGTTTTTCTGCCCAAAACATATATTACATGGGAAATCATTATGAGGAATTTAAGTTAGCGTTGATAGAAAAACTTAATCTATATGCGCATATATTACCGAATGCCAAAAAAGATTTTGATGAAAAGGCGGCCTATTCTTTAGTTCAAAAAGAAAGCTTGTTTGCTGGATTAAAAGTAGAGGTGGATATTCATCAACGTTGGATGATTATTGAGTCAAATGCAGGTGTGGAATTTAATCAACCACTTGGCAGATGTGTGTACCTTCATCACGTCGATAATATTGAGCAAATATTGCCTTATGTTCAAAAAAATAAGACGCAAACCATATCTATTTTTCCTTGGGAGTCATCATTTAAATATCGAGATGCGTTAGCATTAAAAGGTGCGGAAAGGATTGTAGAAGCAGGAATGAATAACATATTTCGAGTTGGTGGATCTCATGACGGAATGAGACCGTTGCAACGATTAGTGACATATATTTCTCATGAAAGGCCATCTAACTATACGGCTAAGGATGTTGCGGTTGAAATAGAACAGACTCGATTCCTGGAAGAAGATAAGTTCCTTGTATTTGTCCCATAATAATACTAGAGAAAGAGGAGAAATACTAGATGGAAAATGAATCAAAATATAAAACCATCGACCACGTTATTTGTGTTGAAGGAAATAAAAAAATTCATGTTTGGGAAACGCTGCCAGAAGAAAACAGCCCAAAGAGAAAGAATGCCATTATTATTGCGTCTGGTTTTGCCCGCAGGATGGATCATTTTGCTGGTCTGGCGGAATATTTATCGCGGAATGGATTTCATGTGATCCGCTATGATTCGCTTCACCACGTTGGATTGAGTTCAGGGACAATTGATGAATTTACAATGTCTATAGGAAAGCAGAGCTTGTTAGCAGTGGTTGATTGGTTAACTACACGAAAAATAAATAACTTCGGTATGTTGGCTTCAAGCTTATCTGCGCGGATAGCTTATGCAAGCCTATCTGAAATCAATGCTTCGTTTTTAATCACCGCAGTCGGTGTTGTTAACTTAAGATATTCTCTTGAAAGAGCTTTAGGGTTTGATTATCTCAGTCTACCCATTAATGAATTGCCGGATAATCTGGATTTTGAAGGCCATAAATTGGGTGCTGAAGTCTTTGCGAGAGATTGTCTTGATTTTGGTTGGGAAGATTTAGCTTCTACAATTAATAACATGATGTATCTTGATATACCGTTTATTGCTTTTACTGCAAATAACGATAATTGGGTCAAGCAAGATGAAGTTATCACATTGTTATCAAATATTCGTAGTAATCGATGCAAGATATATTCTTTGTTAGGAAGTTCGCATGACTTGAGTGAAAATTTAGTGGTCCTGCGCAATTTTTATCAATCGGTTACGAAAGCCGCTATCGCGATGGATAATGATCATCTGGATATTGATGTTGATATTACTGAACCGTCATTTGAACATTTAACTATTGCGACAGTCAATGAACGCCGAATGAGAATTGAGATTGAAAATCAAGCAATTTCTCTGTCTTAATAATACTAGAGAAAGAGGAGAAATACTAGATGACTTCATATGTTGATAAACAAGAAATTACAGCAAGCTCAGAAATTGATGATTTGATTTTTTCGAGCGATCCATTAGTGTGGTCTTACGACGAGCAGGAAAAAATCAGAAAGAAACTTGTGCTTGATGCATTTCGTAATCATTATAAACATTGTCGAGAATATCGTCACTACTGTCAGGCACACAAAGTAGATGACAATATTACGGAAATTGATGACATACCTGTATTCCCAACATCGGTTTTTAAGTTTACTCGCTTATTAACTTCTCAGGAAAACGAGATTGAAAGTTGGTTTACCAGTAGCGGCACGAATGGTTTAAAAAGTCAGGTGGCGCGTGACAGATTAAGTATTGAGAGACTCTTAGGCTCTGTGAGTTATGGCATGAAATATGTTGGTAGTTGGTTTGATCATCAAATAGAATTAGTCAATTTGGGACCAGATAGATTTAATGCTCATAATATTTGGTTTAAATATGTTATGAGTTTGGTGGAATTGTTATATCCTACGACATTTACCGTAACAGAAGAACGAATAGATTTTGTTAAAACATTGAATAGTCTTGAACGAATAAAAAATCAAGGGAAAGATCTTTGTCTTATTGGTTCGCCATACTTTATTTATTTACTCTGCCATTATATGAAAGATAAAAAAATCTCATTTTCTGGAGATAAAAGCCTTTATATCATAACCGGAGGCGGCTGGAAAAGTTACGAAAAAGAATCTCTGAAACGTGATGATTTCAATCATCTTTTATTTGATACTTTCAATCTCAGTGATATTAGTCAGATCCGAGATATATTTAATCAAGTTGAACTCAACACTTGTTTCTTTGAGGATGAAATGCAGCGTAAACATGTTCCGCCGTGGGTATATGCGCGAGCGCTTGATCCTGAAACGTTGAAACCTGTACCTGATGGAACGCCGGGGTTGATGAGTTATATGGATGCGTCAGCAACCAGTTATCCAGCATTTATTGTTACCGATGATGTCGGGATAATTAGCAGAGAATATGGTAAGTATCCCGGCGTGCTCGTTGAAATTTTACGTCGCGTCAATACGAGGACGCAGAAAGGGTGTGCTTTAAGCTTAACCGAAGCGTTTGATAGTTAAGATACAATCGATTTCTGGATCCGCAGGCCTCTGCTAGCTTGACTGACTGAGATACAGCGTACCTTCAGCTCACAGACATGATAAGATACATTGATGAGTTTGGACAAACCACAACTAGAATGCAGTGAAAAAAATGCTTTATTTGTGAAATTTGTGATGCTATTGCTTTATTTGTAACCATTATAAGCTGCAATAAACAAGTTAACAACAACAATTGCATTCATTTTATGTTTCAGGTTCAGGGGGAGGTGTGGGAGGTTTTTTAAAGCAAACGCTAAATTGCCTGATGCGCTACGCTTATCATCCTTAGCGAAAGCTAAGGAAGGCACTGTAT

>pSC101_pBAD_Cre_chlor

ggcttctcagggcgttttatggcgggtctgctatgtggtgctatctgactttttgctgttcagcagttcctgccctctgattttccagtctgaccacttcggattatcccgtgacaggtcattcagactggctaatgcacccagtaaggcagcggtatcatcaacggggtctgacgctcagtggaacgaaaactcacgttaagggatttacgccccgccctgccactcatcgcagtactgttgtaattcattaagcattctgccgacatggaagccatcacagacggcatgatgaacctgaatcgccagcggcatcagcaccttgtcgccttgcgtataatatttgcccacggtgaaaacgggggcgaagaagttgtccatattggccacgtttaaatcaaaactggtgaaactcacccagggattggctgagacgaaaaacatattctcaataaaccctttagggaaataggccaggttttcaccgtaacacgccacatcttgcgaatatatgtgtagaaactgccggaaatcgtcgtggtattcactccagagcgatgaaaacgtttcagtttgctcatggaaaacggtgtaacaagggtgaacactatcccatatcaccagctcaccgtctttcattgccatacggaattccggatgagcattcatcaggcgggcaagaatgtgaataaaggccggataaaacttgtgcttatttttctttacggtctttaaaaaggccgtaatatccagctgaacggtctggttataggtacattgagcaactgactgaaatgcctcaaaatgttctttacgatgccattgggatatatcaacggtggtatatccagtgatttttttctccatGATTGTCCTCCGAGCTCgctagcataatacctagggctgagctagccgtaaacacatttccccgaaaagtgccacctgAGCGCTGAATTCTTATGACAACTTGACGGCTACATCATTCACTTTTTCTTCACAACCGGCACGGAACTCGCTCGGGCTGGCCCCGGTGCATTTTTTAAATACCCGCGAGAAATAGAGTTGATCGTCAAAACCAACATTGCGACCGACGGTGGCGATAGGCATCCGGGTGGTGCTCAAAAGCAGCTTCGCCTGGCTGATACGTTGGTCCTCGCGCCAGCTTAAGACGCTAATCCCTAACTGCTGGCGGAAAAGATGTGACAGACGCGACGGCGACAAGCAAACATGCTGTGCGACGCTGGCGATATCAAAATTGCTGTCTGCCAGGTGATCGCTGATGTACTGACAAGCCTCGCGTACCCGATTATCCATCGGTGGATGGAGCGACTCGTTAATCGCTTCCATGCGCCGCAGTAACAATTGCTCAAGCAGATTTATCGCCAGCAGCTCCGAATAGCGCCCTTCCCCTTGCCCGGCGTTAATGATTTGCCCAAACAGGTCGCTGAAATGCGGCTGGTGCGCTTCATCCGGGCGAAAGAACCCCGTATTGGCAAATATTGACGGCCAGTTAAGCCATTCATGCCAGTAGGCGCGCGGACGAAAGTAAACCCACTGGTGATACCATTCGCGAGCCTCCGGATGACGACCGTAGTGATGAATCTCTCCTGGCGGGAACAGCAAAATATCACCCGGTCGGCAAACAAATTCTCGTCCCTGATTTTTCACCACCCCCTGACCGCGAATGGTGAGATTGAGAATATAACCTTTCATTCCCAGCGGTCGGTCGATAAAAAAATCGAGATAACCGTTGGCCTCAATCGGCGTTAAACCCGCCACCAGATGGGCATTAAACGAGTATCCCGGCAGCAGGGGATCATTTTGCGCTTCAGCCATACTTTTCATACTCCCGCCATTCAGAGAAGAAACCAATTGTCCATATTGCATCAGACATTGCCGTCACTGCGTCTTTTACTGGCTCTTCTCGCTAACCAAACCGGTAACCCCGCTTATTAAAAGCATTCTGTAACAAAGCGGGACCAAAGCCATGACAAAAACGCGTAACAAAAGTGTCTATAATCACGGCAGAAAAGTCCACATTGATTATTTGCACGGCGTCACACTTTGCTATGCCATAGCATTTTTATCCATAAGATTAGCGGATCCTACCTGACGCTTTTTATCGCAACTCTCTACTGTTTctccatacccgagctgtcaccggatgtgctttccggtctgatgagtccgtgaggacgaaacagtaatacgactcactataggggaattgtgagcggataacaattcccctctagaaataattttgtttaactttaagaaggagatataccatggcgcatcaccaccaccatcacggcgggagcagcaatttactgaccgtacaccaaaatttgcctgcattgccggtcgatgcaacgagtgatgaggttcgcaagaacctgatggacatgttcagggatcgccaggcgttttctgagcatacctggaaaatgcttctgtccgtttgccggtcgtgggcggcatggtgcaagttgaataaccggaaatggtttcccgcagaacctgaagatgttcgcgattatcttctatatcttcaggcgcgcggtctggcagtaaaaactatccagcaacatttgggccagctaaacatgcttcatcgtcggtccgggctgccacgaccaagtgacagcaatgctgtttcactggttatgcggcgtatccgaaaagaaaacgttgatgccggtgaacgtgcaaaacaggctctagcgttcgaacgcactgatttcgaccaggttcgttcactcatggaaaatagcgatcgctgccaggatatacgtaatctggcatttctggggattgcttataacaccctgttacgtatagccgaaattgccaggatcagggttaaagatatctcacgtactgacggtgggagaatgttaatccatattggcagaacgaaaacgctggttagcaccgcaggtgtagagaaggcacttagcctgggggtaactaaactggtcgagcgatggatttccgtctctggtgtagctgatgatccgaataactacctgttttgccgggtcagaaaaaatggtgttgccgcgccatctgccaccagccagctatcaactcgcgccctggaagggatttttgaagcaactcatcgattgatttacggcgctaaggatgactctggtcagagatacctggcctggtctggacacagtgcccgtgtcggagccgcgcgagatatggcccgcgctggagtttcaataccggagatcatgcaagctggtggctggaccaatgtaaatattgtcatgaactatatccgtaacctggatagtgaaacaggggcaatggtgcgcctgctggaagatggcgactaaGTTTAAACGGTACCGACAGTAAGACGGGTAAGCCTGTTGATGATACcgctgccttactgggtgcattagccagtctgaatgacctgtcacgggataatccgaagtggtcagactggaaaatcagagggcaggaactgctgaacagcaaaaagtcagatagcaccacatagcagacccgccataaaacgccctgagaagcccgtgacgggcttttcttgtattatgggtagtttccttgcatgaatccataaaaggcgcctgtagtgccatttacccccattcactgccagagccgtgagcgcagcgaactgaatgtcacgaaaaagacagcgactcaggtgcctgatggtcggagacaaaaggaatattcagcgatttgcccgagcttgcgagggtgctacttaagcctttagggttttaaggtctgttttgtagaggagcaaacagcgtttgcgacatccttttgtaatactgcggaactgactaaagtagtgagttatacacagggctgggatctattctttttatctttttttattctttctttattctataaattataaccacttgaatataaacaaaaaaaacacacaaaggtctagcggaatttacagagggtctagcagaatttacaagttttccagcaaaggtctagcagaatttacagatacccacaactcaaaggaaaaggactagtaattatcattgactagcccatctcaattggtatagtgattaaaatcacctagaccaattgagatgtatgtctgaattagttgttttcaaagcaaatgaactagcgattagtcgctatgacttaacggagcatgaaaccaagctaattttatgctgtgtggcactactcaaccccacgattgaaaaccctacaaggaaagaacggacggtatcgttcacttataaccaatacgctcagatgatgaacatcagtagggaaaatgcttatggtgtattagctaaagcaaccagagagctgatgacgagaactgtggaaatcaggaatcctttggttaaaggctttgagattttccagtggacaaactatgccaagttctcaagcgaaaaattagaattagtttttagtgaagagatattgccttatcttttccagttaaaaaaattcataaaatataatctggaacatgttaagtcttttgaaaacaaatactctatgaggatttatgagtggttattaaaagaactaacacaaaagaaaactcacaaggcaaatatagagattagccttgatgaatttaagttcatgttaatgcttgaaaataactaccatgagtttaaaaggcttaaccaatgggttttgaaaccaataagtaaagatttaaacacttacagcaatatgaaattggtggttgataagcgaggccgcccgactgatacgttgattttccaagttgaactagatagacaaatggatctcgtaaccgaacttgagaacaaccagataaaaatgaatggtgacaaaataccaacaaccattacatcagattcctacctacgtaacggactaagaaaaacactacacgatgctttaactgcaaaaattcagctcaccagttttgaggcaaaatttttgagtgacatgcaaagtaagcatgatctcaatggttcgttctcatggctcacgcaaaaacaacgaaccacactagagaacatactggctaaatacggaaggatctgaggttcttatggctcttgtatctatcagtgaagcaTCAAGACTAACAAACAAAAGTAGAACAACTGTTCACCGTTA

>Genomically_Integrated_2LoxP_*dif*_landing_site_orientation1 (2666 bp)

TTAAATAAGTAACAGCACCCTCTGGTGTTTTCGTATGCAGTGTCTATGCGACAGACACTGCGCTCGCTGGGAGTTCGTAGACGGAAACAAACGCAGAATCCAAGCATAACTTCGTATAGGATACTTTATACGAAGTTATCTCGAGTTGACAGCTAGCTCAGTCCTAGGTATAATACTAGTACCCGAGCTGTCACCGGATGTGCTTTCCGGTCTGATGAGTCCGTGAGGACGAAACAGCCTCTACAAATAATTTTGTTTAATACTAGAGAAAGAGGGGAAATACTAGATGAACATCAAAAAGTTTGCAAAACAAGCAACAGTATTAACCTTTACTACCGCACTGCTGGCAGGAGGCGCAACTCAAGCGTTTGCGAAAGAAACGAACCAAAAGCCATATAAGGAAACATACGGCATTTCCCATATTACACGCCATGATATGCTGCAAATCCCTGAACAGCAAAAAAATGAAAAATATAAAGTTCCTGAGTTCGATTCGTCCACAATTAAAAATATCTCTTCTGCAAAAGGCCTGGACGTTTGGGACAGCTGGCCATTACAAAACACTGACGGCACTGTCGCAAACTATCACGGCTACCACATCGTCTTTGCATTAGCCGGAGATCCTAAAAATGCGGATGACACATCGATTTACATGTTCTATCAAAAAGTCGGCGAAACTTCTATTGACAGCTGGAAAAACGCTGGCCGCGTCTTTAAAGACAGCGACAAATTCGATGCAAATGATTCTATCCTAAAAGACCAAACACAAGAATGGTCAGGTTCAGCCACATTTACATCTGACGGAAAAATCCGTTTATTCTACACTGATTTCTCCGGTAAACATTACGGCAAACAAACACTGACAACTGCACAAGTTAACGTATCAGCATCAGACAGCTCTTTGAACATCAACGGTGTAGAGGATTATAAATCAATCTTTGACGGTGACGGAAAAACGTATCAAAATGTACAGCAGTTCATCGATGAAGGCAACTACAGCTCAGGCGACAACCATACGCTGAGAGATCCTCACTACGTAGAAGATAAAGGCCACAAATACTTAGTATTTGAAGCAAACACTGGAACTGAAGATGGCTACCAAGGCGAAGAATCTTTATTTAACAAAGCATACTATGGCAAAAGCACATCATTCTTCCGTCAAGAAAGTCAAAAACTTCTGCAAAGCGATAAAAAACGCACGGCTGAGTTAGCAAACGGCGCTCTCGGTATGATTGAGCTAAACGATGATTACACACTGAAAAAAGTGATGAAACCGCTGATTGCATCTAACACAGTAACAGATGAAATTGAACGCGCGAACGTCTTTAAAATGAACGGCAAATGGTACCTGTTCACTGACTCCCGCGGATCAAAAATGACGATTGACGGCATTACGTCTAACGATATTTACATGCTTGGTTATGTTTCTAATTCTTTAACTGGCCCATACAAGCCGCTGAACAAAACTGGCCTTGTGTTAAAAATGGATCTTGATCCTAACGATGTAACCTTTACTTACTCACACTTCGCTGTACCTCAAGCGAAAGGAAACAATGTCGTGATTACAAGCTATATGACAAACAGAGGATTCTACGCAGACAAACAATCAACGTTTGCGCCTAGCTTCCTGCTGAACATCAAAGGCAAGAAAACATCTGTTGTCAAAGACAGCATCCTTGAACAAGGACAATTAACAGTTAACAAATAAGGAGGACAATCATGAGGGAAGCGGTGATCGCCGAAGTATCGACTCAACTATCAGAGGTAGTTGGCGTCATCGAGCGCCATCTCGAACCGACGTTGCTGGCCGTACATTTGTACGGCTCCGCAGTGGATGGCGGCCTGAAGCCACACAGTGATATTGATTTGCTGGTTACGGTGACCGTAAGGCTTGATGAAACAACGCGGCGAGCTTTGATCAACGACCTTTTGGAAACTTCGGCTTCCCCTGGAGAGAGCGAGATTCTCCGCGCTGTAGAAGTCACCATTGTTGTGCACGACGACATCATTCCGTGGCGTTATCCAGCTAAGCGCGAACTGCAATTTGGAGAATGGCAGCGCAATGACATTCTTGCAGGTATCTTCGAGCCAGCCACGTACGACATTGATCTGGCTATCTTGCTGACAAAAGCAAGAGAACATAGCGTTGCCTTGGTAGGTCCAGCGGCGGAGGAACTCTTTGATCCGGTTCCTGAACAGGATCTATTTGAGGCGCTAAATGAAACCTTAACGCTATGGAACTCGCCGCCCGACTGGGCTGGCGATGAGCGAAATGTAGTGCTTACGTTGTCCCGCATTTGGTACAGCGCAGTAACCGGCAAAATCGCGCCGAAGGATGTCGCTGCCGACTGGGCAATGGAGCGCCTGCCGGCCCAGTATCAGCCCGTCATACTTGAAGCTAGACAGGCTTATCTTGGACAAGAAGAAGATCGCTTGGCCTCGCGCGCAGATCAGTTGGAAGAATTTGTCCACTACGTGAAAGGCGAGATCACCAAGGTAGTCGGCAAATAAGCGATCGCATAACTTCGTATAGCATACATTATACGAAGTTATGGCGCGCCGAGACGAGACGAGACAGCCTGAGAATGGATGCGAGTAATGACCCCATATCTCACTTACTACCGTATTCTCGGCTTAACCGTCCATGCAACCTCAACACATTGCTT

>Genomically_Integrated_2LoxP_*dif*_landing_site_orientation2 (2666 bp)

TTAAATAAGTAACAGCACCCTCTGGTGTTTTCGTATGCAGTGTCTATGCGACAGACACTGCGCTCCATTACTCGCATCCATTCTCAGGCTGTCTCGTCTCGTCTCGGCGCGCCATAACTTCGTATAATGTATGCTATACGAAGTTATGCGATCGCTTATTTGCCGACTACCTTGGTGATCTCGCCTTTCACGTAGTGGACAAATTCTTCCAACTGATCTGCGCGCGAGGCCAAGCGATCTTCTTCTTGTCCAAGATAAGCCTGTCTAGCTTCAAGTATGACGGGCTGATACTGGGCCGGCAGGCGCTCCATTGCCCAGTCGGCAGCGACATCCTTCGGCGCGATTTTGCCGGTTACTGCGCTGTACCAAATGCGGGACAACGTAAGCACTACATTTCGCTCATCGCCAGCCCAGTCGGGCGGCGAGTTCCATAGCGTTAAGGTTTCATTTAGCGCCTCAAATAGATCCTGTTCAGGAACCGGATCAAAGAGTTCCTCCGCCGCTGGACCTACCAAGGCAACGCTATGTTCTCTTGCTTTTGTCAGCAAGATAGCCAGATCAATGTCGTACGTGGCTGGCTCGAAGATACCTGCAAGAATGTCATTGCGCTGCCATTCTCCAAATTGCAGTTCGCGCTTAGCTGGATAACGCCACGGAATGATGTCGTCGTGCACAACAATGGTGACTTCTACAGCGCGGAGAATCTCGCTCTCTCCAGGGGAAGCCGAAGTTTCCAAAAGGTCGTTGATCAAAGCTCGCCGCGTTGTTTCATCAAGCCTTACGGTCACCGTAACCAGCAAATCAATATCACTGTGTGGCTTCAGGCCGCCATCCACTGCGGAGCCGTACAAATGTACGGCCAGCAACGTCGGTTCGAGATGGCGCTCGATGACGCCAACTACCTCTGATAGTTGAGTCGATACTTCGGCGATCACCGCTTCCCTCATGATTGTCCTCCTTATTTGTTAACTGTTAATTGTCCTTGTTCAAGGATGCTGTCTTTGACAACAGATGTTTTCTTGCCTTTGATGTTCAGCAGGAAGCTAGGCGCAAACGTTGATTGTTTGTCTGCGTAGAATCCTCTGTTTGTCATATAGCTTGTAATCACGACATTGTTTCCTTTCGCTTGAGGTACAGCGAAGTGTGAGTAAGTAAAGGTTACATCGTTAGGATCAAGATCCATTTTTAACACAAGGCCAGTTTTGTTCAGCGGCTTGTATGGGCCAGTTAAAGAATTAGAAACATAACCAAGCATGTAAATATCGTTAGACGTAATGCCGTCAATCGTCATTTTTGATCCGCGGGAGTCAGTGAACAGGTACCATTTGCCGTTCATTTTAAAGACGTTCGCGCGTTCAATTTCATCTGTTACTGTGTTAGATGCAATCAGCGGTTTCATCACTTTTTTCAGTGTGTAATCATCGTTTAGCTCAATCATACCGAGAGCGCCGTTTGCTAACTCAGCCGTGCGTTTTTTATCGCTTTGCAGAAGTTTTTGACTTTCTTGACGGAAGAATGATGTGCTTTTGCCATAGTATGCTTTGTTAAATAAAGATTCTTCGCCTTGGTAGCCATCTTCAGTTCCAGTGTTTGCTTCAAATACTAAGTATTTGTGGCCTTTATCTTCTACGTAGTGAGGATCTCTCAGCGTATGGTTGTCGCCTGAGCTGTAGTTGCCTTCATCGATGAACTGCTGTACATTTTGATACGTTTTTCCGTCACCGTCAAAGATTGATTTATAATCCTCTACACCGTTGATGTTCAAAGAGCTGTCTGATGCTGATACGTTAACTTGTGCAGTTGTCAGTGTTTGTTTGCCGTAATGTTTACCGGAGAAATCAGTGTAGAATAAACGGATTTTTCCGTCAGATGTAAATGTGGCTGAACCTGACCATTCTTGTGTTTGGTCTTTTAGGATAGAATCATTTGCATCGAATTTGTCGCTGTCTTTAAAGACGCGGCCAGCGTTTTTCCAGCTGTCAATAGAAGTTTCGCCGACTTTTTGATAGAACATGTAAATCGATGTGTCATCCGCATTTTTAGGATCTCCGGCTAATGCAAAGACGATGTGGTAGCCGTGATAGTTTGCGACAGTGCCGTCAGTGTTTTGTAATGGCCAGCTGTCCCAAACGTCCAGGCCTTTTGCAGAAGAGATATTTTTAATTGTGGACGAATCGAACTCAGGAACTTTATATTTTTCATTTTTTTGCTGTTCAGGGATTTGCAGCATATCATGGCGTGTAATATGGGAAATGCCGTATGTTTCCTTATATGGCTTTTGGTTCGTTTCTTTCGCAAACGCTTGAGTTGCGCCTCCTGCCAGCAGTGCGGTAGTAAAGGTTAATACTGTTGCTTGTTTTGCAAACTTTTTGATGTTCATCTAGTATTTCCCCTCTTTCTCTAGTATTAAACAAAATTATTTGTAGAGGCTGTTTCGTCCTCACGGACTCATCAGACCGGAAAGCACATCCGGTGACAGCTCGGGTACTAGTATTATACCTAGGACTGAGCTAGCTGTCAACTCGAGATAACTTCGTATAAAGTATCCTATACGAAGTTATGCTTGGATTCTGCGTTTGTTTCCGTCTACGAACTCCCAGCACCCCATATCTCACTTACTACCGTATTCTCGGCTTAACCGTCCATGCAACCTCAACACATTGCTT

>LUX_2LoxP_Assembled_Linear_for_Integration_Orientation1 (7093 bp)

CCAAGCATAACTTCGTATAGGATACTTTATACGAAGTTATGAATTCGCGGCCGCTTCTAGAGGAAATCATAAAAAATTTATTTGCTTTGTGAGCGGATAACAATTATAATAGATTCAATTGTGAGCGGATAACAATTACTAGAGAAAGAGGAGAAATACTAGATGAAATTTGGAAACTTTTTGCTTACATACCAACCTCCCCAATTTTCTCAAACAGAGGTAATGAAACGTTTGGTTAAATTAGGTCGCATCTCTGAGGAGTGTGGTTTTGATACCGTATGGTTACTGGAGCATCATTTCACGGAGTTTGGTTTGCTTGGTAACCCTTATGTCGCTGCTGCATATTTACTTGGCGCGACTAAAAAATTGAATGTAGGAACTGCCGCTATTGTTCTTCCCACAGCCCATCCAGTACGCCAACTTGAAGATGTGAATTTATTGGATCAAATGTCAAAAGGACGATTTCGGTTTGGTATTTGCCGAGGGCTTTACAACAAGGACTTTCGCGTATTCGGCACAGATATGAATAACAGTCGCGCCTTAGCGGAATGCTGGTACGGGCTGATAAAGAATGGCATGACAGAGGGATATATGGAAGCTGATAATGAACATATCAAGTTCCATAAGGTAAAAGTAAACCCCGCGGCGTATAGCAGAGGTGGCGCACCGGTTTATGTGGTGGCTGAATCAGCTTCGACGACTGAGTGGGCTGCTCAATTTGGCCTACCGATGATATTAAGTTGGATTATAAATACTAACGAAAAGAAAGCACAACTTGAGCTTTATAATGAAGTGGCTCAAGAATATGGGCACGATATTCATAATATCGACCATTGCTTATCATATATAACATCTGTAGATCATGACTCAATTAAAGCGAAAGAGATTTGCCGGAAATTTCTGGGGCATTGGTATGATTCTTATGTGAATGCTACGACTATTTTTGATGATTCAGACCAAACAAGAGGTTATGATTTCAATAAAGGGCAGTGGCGTGACTTTGTATTAAAAGGACATAAAGATACTAATCGCCGTATTGATTACAGTTACGAAATCAATCCCGTGGGAACGCCGCAGGAATGTATTGACATAATTCAAAAAGACATTGATGCTACAGGAATATCAAATATTTGTTGTGGATTTGAAGCTAATGGAACAGTAGACGAAATTATTGCTTCCATGAAGCTCTTCCAGTCTGATGTCATGCCATTTCTTAAAGAAAAACAACGTTCGCTATTATATTAATAATACTAGAGAAAGAGGAGAAATACTAGATGAAATTTGGATTGTTCTTCCTTAACTTCATCAATTCAACAACTGTTCAAGAACAAAGTATAGTTCGCATGCAGGAAATAACGGAGTATGTTGATAAGTTGAATTTTGAACAGATTTTAGTGTATGAAAATCATTTTTCAGATAATGGTGTTGTCGGCGCTCCTCTGACTGTTTCTGGTTTTCTGCTCGGTTTAACAGAGAAAATTAAAATTGGTTCATTAAATCACATCATTACAACTCATCATCCTGTCCGCATAGCGGAGGAAGCTTGCTTATTGGATCAGTTAAGTGAAGGGAGATTTATTTTAGGGTTTAGTGATTGCGAAAAAAAAGATGAAATGCATTTTTTTAATCGCCCGGTTGAATATCAACAGCAACTATTTGAAGAGTGTTATGAAATCATTAACGATGCTTTAACAACAGGCTATTGTAATCCAGATAACGATTTTTATAGCTTCCCTAAAATATCTGTAAATCCCCATGCTTATACGCCAGGCGGACCTCGGAAATATGTAACAGCAACCAGTCATCATATTGTTGAGTGGGCGGCCAAAAAAGGTATTCCTCTCATCTTTAAGTGGGATGATTCTAATGATGTTAGATATGAATATGCTGAAAGATATAAAGCCGTTGCGGATAAATATGACGTTGACCTATCAGAGATAGACCATCAGTTAATGATATTAGTTAACTATAACGAAGATAGTAATAAAGCTAAACAAGAGACGCGTGCATTTATTAGTGATTATGTTCTTGAAATGCACCCTAATGAAAATTTCGAAAATAAACTTGAAGAAATAATTGCAGAAAACGCTGTCGGAAATTATACGGAGTGTATAACTGCGGCTAAGTTGGCAATTGAAAAGTGTGGTGCGAAAAGTGTATTGCTGTCCTTTGAACCAATGAATGATTTGATGAGCCAAAAAAATGTAATCAATATTGTTGATGATAATATTAAGAAGTACCACATGGAATATACCTAATAATACTAGAGAAAGAGGAGAAATACTAGATGACTAAAAAAATTTCATTCATTATTAACGGCCAGGTTGAAATCTTTCCCGAAAGTGATGATTTAGTGCAATCCATTAATTTTGGTGATAATAGTGTTTACCTGCCAATATTGAATGACTCTCATGTAAAAAACATTATTGATTGTAATGGAAATAACGAATTACGGTTGCATAACATTGTCAATTTTCTCTATACGGTAGGGCAAAGATGGAAAAATGAAGAATACTCAAGACGCAGGACATACATTCGTGACTTAAAAAAATATATGGGATATTCAGAAGAAATGGCTAAGCTAGAGGCCAATTGGATATCTATGATTTTATGTTCTAAAGGCGGCCTTTATGATGTTGTAGAAAATGAACTTGGTTCTCGCCATATCATGGATGAATGGCTACCTCAGGATGAAAGTTATGTTCGGGCTTTTCCGAAAGGTAAATCTGTACATCTGTTGGCAGGTAATGTTCCATTATCTGGGATCATGTCTATATTACGCGCAATTTTAACTAAGAATCAGTGTATTATAAAAACATCGTCAACCGATCCTTTTACCGCTAATGCATTAGCGTTAAGTTTTATTGATGTAGACCCTAATCATCCGATAACGCGCTCTTTATCTGTTATATATTGGCCCCACCAAGGTGATACATCACTCGCAAAAGAAATTATGCGACATGCGGATGTTATTGTCGCTTGGGGAGGGCCAGATGCGATTAATTGGGCGGTAGAGCATGCGCCATCTTATGCTGATGTGATTAAATTTGGTTCTAAAAAGAGTCTTTGCATTATCGATAATCCTGTTGATTTGACGTCCGCAGCGACAGGTGCGGCTCATGATGTTTGTTTTTACGATCAGCGAGCTTGTTTTTCTGCCCAAAACATATATTACATGGGAAATCATTATGAGGAATTTAAGTTAGCGTTGATAGAAAAACTTAATCTATATGCGCATATATTACCGAATGCCAAAAAAGATTTTGATGAAAAGGCGGCCTATTCTTTAGTTCAAAAAGAAAGCTTGTTTGCTGGATTAAAAGTAGAGGTGGATATTCATCAACGTTGGATGATTATTGAGTCAAATGCAGGTGTGGAATTTAATCAACCACTTGGCAGATGTGTGTACCTTCATCACGTCGATAATATTGAGCAAATATTGCCTTATGTTCAAAAAAATAAGACGCAAACCATATCTATTTTTCCTTGGGAGTCATCATTTAAATATCGAGATGCGTTAGCATTAAAAGGTGCGGAAAGGATTGTAGAAGCAGGAATGAATAACATATTTCGAGTTGGTGGATCTCATGACGGAATGAGACCGTTGCAACGATTAGTGACATATATTTCTCATGAAAGGCCATCTAACTATACGGCTAAGGATGTTGCGGTTGAAATAGAACAGACTCGATTCCTGGAAGAAGATAAGTTCCTTGTATTTGTCCCATAATAATACTAGAGAAAGAGGAGAAATACTAGATGGAAAATGAATCAAAATATAAAACCATCGACCACGTTATTTGTGTTGAAGGAAATAAAAAAATTCATGTTTGGGAAACGCTGCCAGAAGAAAACAGCCCAAAGAGAAAGAATGCCATTATTATTGCGTCTGGTTTTGCCCGCAGGATGGATCATTTTGCTGGTCTGGCGGAATATTTATCGCGGAATGGATTTCATGTGATCCGCTATGATTCGCTTCACCACGTTGGATTGAGTTCAGGGACAATTGATGAATTTACAATGTCTATAGGAAAGCAGAGCTTGTTAGCAGTGGTTGATTGGTTAACTACACGAAAAATAAATAACTTCGGTATGTTGGCTTCAAGCTTATCTGCGCGGATAGCTTATGCAAGCCTATCTGAAATCAATGCTTCGTTTTTAATCACCGCAGTCGGTGTTGTTAACTTAAGATATTCTCTTGAAAGAGCTTTAGGGTTTGATTATCTCAGTCTACCCATTAATGAATTGCCGGATAATCTGGATTTTGAAGGCCATAAATTGGGTGCTGAAGTCTTTGCGAGAGATTGTCTTGATTTTGGTTGGGAAGATTTAGCTTCTACAATTAATAACATGATGTATCTTGATATACCGTTTATTGCTTTTACTGCAAATAACGATAATTGGGTCAAGCAAGATGAAGTTATCACATTGTTATCAAATATTCGTAGTAATCGATGCAAGATATATTCTTTGTTAGGAAGTTCGCATGACTTGAGTGAAAATTTAGTGGTCCTGCGCAATTTTTATCAATCGGTTACGAAAGCCGCTATCGCGATGGATAATGATCATCTGGATATTGATGTTGATATTACTGAACCGTCATTTGAACATTTAACTATTGCGACAGTCAATGAACGCCGAATGAGAATTGAGATTGAAAATCAAGCAATTTCTCTGTCTTAATAATACTAGAGAAAGAGGAGAAATACTAGATGACTTCATATGTTGATAAACAAGAAATTACAGCAAGCTCAGAAATTGATGATTTGATTTTTTCGAGCGATCCATTAGTGTGGTCTTACGACGAGCAGGAAAAAATCAGAAAGAAACTTGTGCTTGATGCATTTCGTAATCATTATAAACATTGTCGAGAATATCGTCACTACTGTCAGGCACACAAAGTAGATGACAATATTACGGAAATTGATGACATACCTGTATTCCCAACATCGGTTTTTAAGTTTACTCGCTTATTAACTTCTCAGGAAAACGAGATTGAAAGTTGGTTTACCAGTAGCGGCACGAATGGTTTAAAAAGTCAGGTGGCGCGTGACAGATTAAGTATTGAGAGACTCTTAGGCTCTGTGAGTTATGGCATGAAATATGTTGGTAGTTGGTTTGATCATCAAATAGAATTAGTCAATTTGGGACCAGATAGATTTAATGCTCATAATATTTGGTTTAAATATGTTATGAGTTTGGTGGAATTGTTATATCCTACGACATTTACCGTAACAGAAGAACGAATAGATTTTGTTAAAACATTGAATAGTCTTGAACGAATAAAAAATCAAGGGAAAGATCTTTGTCTTATTGGTTCGCCATACTTTATTTATTTACTCTGCCATTATATGAAAGATAAAAAAATCTCATTTTCTGGAGATAAAAGCCTTTATATCATAACCGGAGGCGGCTGGAAAAGTTACGAAAAAGAATCTCTGAAACGTGATGATTTCAATCATCTTTTATTTGATACTTTCAATCTCAGTGATATTAGTCAGATCCGAGATATATTTAATCAAGTTGAACTCAACACTTGTTTCTTTGAGGATGAAATGCAGCGTAAACATGTTCCGCCGTGGGTATATGCGCGAGCGCTTGATCCTGAAACGTTGAAACCTGTACCTGATGGAACGCCGGGGTTGATGAGTTATATGGATGCGTCAGCAACCAGTTATCCAGCATTTATTGTTACCGATGATGTCGGGATAATTAGCAGAGAATATGGTAAGTATCCCGGCGTGCTCGTTGAAATTTTACGTCGCGTCAATACGAGGACGCAGAAAGGGTGTGCTTTAAGCTTAACCGAAGCGTTTGATAGTTAAGATACAATCGATTTCTGGATCCGCAGGCCTCTGCTAGCTTGACTGACTGAGATACAGCGTACCTTCAGCTCACAGACATGATAAGATACATTGATGAGTTTGGACAAACCACAACTAGAATGCAGTGAAAAAAATGCTTTATTTGTGAAATTTGTGATGCTATTGCTTTATTTGTAACCATTATAAGCTGCAATAAACAAGTTAACAACAACAATTGCATTCATTTTATGTTTCAGGTTCAGGGGGAGGTGTGGGAGGTTTTTTAAAGCAAACGCTAAATTGCCTGATGCGCTACGCTTATCATCCTTAGCGAAAGCTAAGGAAGGCACTGTATTGATCGGCACGTAAGAGGTTCCAACTTTCACCATAATGAAATAAGATCACTACCGGGCGTATTTTTTGAGTTATCGAGATTTTCAGGAGCTAAGGAAGCTAAAATGGAGAAAAAAATCACTGGATATACCACCGTTGATATATCCCAATGGCATCGTAAAGAACATTTTGAGGCATTTCAGTCAGTTGCTCAATGTACCTATAACCAGACCGTTCAGCTGGATATTACGGCCTTTTTAAAGACCGTAAAGAAAAATAAGCACAAGTTTTATCCGGCCTTTATTCACATTCTTGCCCGCCTGATGAATGCTCATCCGGAATTTCGTATGGCAATGAAAGACGGTGAGCTGGTGATATGGGATAGTGTTCACCCTTGTTACACCGTTTTCCATGAGCAAACTGAAACGTTTTCATCGCTCTGGAGTGAATACCACGACGATTTCCGGCAGTTTCTACACATATATTCGCAAGATGTGGCGTGTTACGGTGAAAACCTGGCCTATTTCCCTAAAGGGTTTATTGAGAATATGTTTTTCGTCTCAGCCAATCCCTGGGTGAGTTTCACCAGTTTTGATTTAAACGTGGCCAATATGGACAACTTCTTCGCCCCCGTTTTCACCATGGGCAAATATTATACGCAAGGCGACAAGGTGCTGATGCCGCTGGCGATTCAGGTTCATCATGCCGTTTGTGATGGCTTCCATGTCGGCAGAATGCTTAATGAATTACAACAGTACTGCGATGAGTGGCAGGGCGGGGCGTAATTTGATATCGAGCTCGCTTGGACTCCTGTTGATAGATCCAGTAATGACCTCAGAACTCCATCTGGATTTGTTCAGAACGCTCGGTTGCCGCCGGGCGTTTTTTATTGGTGAGAATCCAAGCACTAGTAACAATAACTTCGTATAGCATACATTATACGAAGTTATGGCGCG

>LUX_2LoxP_Assembled_Linear_for_Integration_Orientation2 (7093 bp)

CGCGCCATAACTTCGTATAATGTATGCTATACGAAGTTATGAATTCGCGGCCGCTTCTAGAGGAAATCATAAAAAATTTATTTGCTTTGTGAGCGGATAACAATTATAATAGATTCAATTGTGAGCGGATAACAATTACTAGAGAAAGAGGAGAAATACTAGATGAAATTTGGAAACTTTTTGCTTACATACCAACCTCCCCAATTTTCTCAAACAGAGGTAATGAAACGTTTGGTTAAATTAGGTCGCATCTCTGAGGAGTGTGGTTTTGATACCGTATGGTTACTGGAGCATCATTTCACGGAGTTTGGTTTGCTTGGTAACCCTTATGTCGCTGCTGCATATTTACTTGGCGCGACTAAAAAATTGAATGTAGGAACTGCCGCTATTGTTCTTCCCACAGCCCATCCAGTACGCCAACTTGAAGATGTGAATTTATTGGATCAAATGTCAAAAGGACGATTTCGGTTTGGTATTTGCCGAGGGCTTTACAACAAGGACTTTCGCGTATTCGGCACAGATATGAATAACAGTCGCGCCTTAGCGGAATGCTGGTACGGGCTGATAAAGAATGGCATGACAGAGGGATATATGGAAGCTGATAATGAACATATCAAGTTCCATAAGGTAAAAGTAAACCCCGCGGCGTATAGCAGAGGTGGCGCACCGGTTTATGTGGTGGCTGAATCAGCTTCGACGACTGAGTGGGCTGCTCAATTTGGCCTACCGATGATATTAAGTTGGATTATAAATACTAACGAAAAGAAAGCACAACTTGAGCTTTATAATGAAGTGGCTCAAGAATATGGGCACGATATTCATAATATCGACCATTGCTTATCATATATAACATCTGTAGATCATGACTCAATTAAAGCGAAAGAGATTTGCCGGAAATTTCTGGGGCATTGGTATGATTCTTATGTGAATGCTACGACTATTTTTGATGATTCAGACCAAACAAGAGGTTATGATTTCAATAAAGGGCAGTGGCGTGACTTTGTATTAAAAGGACATAAAGATACTAATCGCCGTATTGATTACAGTTACGAAATCAATCCCGTGGGAACGCCGCAGGAATGTATTGACATAATTCAAAAAGACATTGATGCTACAGGAATATCAAATATTTGTTGTGGATTTGAAGCTAATGGAACAGTAGACGAAATTATTGCTTCCATGAAGCTCTTCCAGTCTGATGTCATGCCATTTCTTAAAGAAAAACAACGTTCGCTATTATATTAATAATACTAGAGAAAGAGGAGAAATACTAGATGAAATTTGGATTGTTCTTCCTTAACTTCATCAATTCAACAACTGTTCAAGAACAAAGTATAGTTCGCATGCAGGAAATAACGGAGTATGTTGATAAGTTGAATTTTGAACAGATTTTAGTGTATGAAAATCATTTTTCAGATAATGGTGTTGTCGGCGCTCCTCTGACTGTTTCTGGTTTTCTGCTCGGTTTAACAGAGAAAATTAAAATTGGTTCATTAAATCACATCATTACAACTCATCATCCTGTCCGCATAGCGGAGGAAGCTTGCTTATTGGATCAGTTAAGTGAAGGGAGATTTATTTTAGGGTTTAGTGATTGCGAAAAAAAAGATGAAATGCATTTTTTTAATCGCCCGGTTGAATATCAACAGCAACTATTTGAAGAGTGTTATGAAATCATTAACGATGCTTTAACAACAGGCTATTGTAATCCAGATAACGATTTTTATAGCTTCCCTAAAATATCTGTAAATCCCCATGCTTATACGCCAGGCGGACCTCGGAAATATGTAACAGCAACCAGTCATCATATTGTTGAGTGGGCGGCCAAAAAAGGTATTCCTCTCATCTTTAAGTGGGATGATTCTAATGATGTTAGATATGAATATGCTGAAAGATATAAAGCCGTTGCGGATAAATATGACGTTGACCTATCAGAGATAGACCATCAGTTAATGATATTAGTTAACTATAACGAAGATAGTAATAAAGCTAAACAAGAGACGCGTGCATTTATTAGTGATTATGTTCTTGAAATGCACCCTAATGAAAATTTCGAAAATAAACTTGAAGAAATAATTGCAGAAAACGCTGTCGGAAATTATACGGAGTGTATAACTGCGGCTAAGTTGGCAATTGAAAAGTGTGGTGCGAAAAGTGTATTGCTGTCCTTTGAACCAATGAATGATTTGATGAGCCAAAAAAATGTAATCAATATTGTTGATGATAATATTAAGAAGTACCACATGGAATATACCTAATAATACTAGAGAAAGAGGAGAAATACTAGATGACTAAAAAAATTTCATTCATTATTAACGGCCAGGTTGAAATCTTTCCCGAAAGTGATGATTTAGTGCAATCCATTAATTTTGGTGATAATAGTGTTTACCTGCCAATATTGAATGACTCTCATGTAAAAAACATTATTGATTGTAATGGAAATAACGAATTACGGTTGCATAACATTGTCAATTTTCTCTATACGGTAGGGCAAAGATGGAAAAATGAAGAATACTCAAGACGCAGGACATACATTCGTGACTTAAAAAAATATATGGGATATTCAGAAGAAATGGCTAAGCTAGAGGCCAATTGGATATCTATGATTTTATGTTCTAAAGGCGGCCTTTATGATGTTGTAGAAAATGAACTTGGTTCTCGCCATATCATGGATGAATGGCTACCTCAGGATGAAAGTTATGTTCGGGCTTTTCCGAAAGGTAAATCTGTACATCTGTTGGCAGGTAATGTTCCATTATCTGGGATCATGTCTATATTACGCGCAATTTTAACTAAGAATCAGTGTATTATAAAAACATCGTCAACCGATCCTTTTACCGCTAATGCATTAGCGTTAAGTTTTATTGATGTAGACCCTAATCATCCGATAACGCGCTCTTTATCTGTTATATATTGGCCCCACCAAGGTGATACATCACTCGCAAAAGAAATTATGCGACATGCGGATGTTATTGTCGCTTGGGGAGGGCCAGATGCGATTAATTGGGCGGTAGAGCATGCGCCATCTTATGCTGATGTGATTAAATTTGGTTCTAAAAAGAGTCTTTGCATTATCGATAATCCTGTTGATTTGACGTCCGCAGCGACAGGTGCGGCTCATGATGTTTGTTTTTACGATCAGCGAGCTTGTTTTTCTGCCCAAAACATATATTACATGGGAAATCATTATGAGGAATTTAAGTTAGCGTTGATAGAAAAACTTAATCTATATGCGCATATATTACCGAATGCCAAAAAAGATTTTGATGAAAAGGCGGCCTATTCTTTAGTTCAAAAAGAAAGCTTGTTTGCTGGATTAAAAGTAGAGGTGGATATTCATCAACGTTGGATGATTATTGAGTCAAATGCAGGTGTGGAATTTAATCAACCACTTGGCAGATGTGTGTACCTTCATCACGTCGATAATATTGAGCAAATATTGCCTTATGTTCAAAAAAATAAGACGCAAACCATATCTATTTTTCCTTGGGAGTCATCATTTAAATATCGAGATGCGTTAGCATTAAAAGGTGCGGAAAGGATTGTAGAAGCAGGAATGAATAACATATTTCGAGTTGGTGGATCTCATGACGGAATGAGACCGTTGCAACGATTAGTGACATATATTTCTCATGAAAGGCCATCTAACTATACGGCTAAGGATGTTGCGGTTGAAATAGAACAGACTCGATTCCTGGAAGAAGATAAGTTCCTTGTATTTGTCCCATAATAATACTAGAGAAAGAGGAGAAATACTAGATGGAAAATGAATCAAAATATAAAACCATCGACCACGTTATTTGTGTTGAAGGAAATAAAAAAATTCATGTTTGGGAAACGCTGCCAGAAGAAAACAGCCCAAAGAGAAAGAATGCCATTATTATTGCGTCTGGTTTTGCCCGCAGGATGGATCATTTTGCTGGTCTGGCGGAATATTTATCGCGGAATGGATTTCATGTGATCCGCTATGATTCGCTTCACCACGTTGGATTGAGTTCAGGGACAATTGATGAATTTACAATGTCTATAGGAAAGCAGAGCTTGTTAGCAGTGGTTGATTGGTTAACTACACGAAAAATAAATAACTTCGGTATGTTGGCTTCAAGCTTATCTGCGCGGATAGCTTATGCAAGCCTATCTGAAATCAATGCTTCGTTTTTAATCACCGCAGTCGGTGTTGTTAACTTAAGATATTCTCTTGAAAGAGCTTTAGGGTTTGATTATCTCAGTCTACCCATTAATGAATTGCCGGATAATCTGGATTTTGAAGGCCATAAATTGGGTGCTGAAGTCTTTGCGAGAGATTGTCTTGATTTTGGTTGGGAAGATTTAGCTTCTACAATTAATAACATGATGTATCTTGATATACCGTTTATTGCTTTTACTGCAAATAACGATAATTGGGTCAAGCAAGATGAAGTTATCACATTGTTATCAAATATTCGTAGTAATCGATGCAAGATATATTCTTTGTTAGGAAGTTCGCATGACTTGAGTGAAAATTTAGTGGTCCTGCGCAATTTTTATCAATCGGTTACGAAAGCCGCTATCGCGATGGATAATGATCATCTGGATATTGATGTTGATATTACTGAACCGTCATTTGAACATTTAACTATTGCGACAGTCAATGAACGCCGAATGAGAATTGAGATTGAAAATCAAGCAATTTCTCTGTCTTAATAATACTAGAGAAAGAGGAGAAATACTAGATGACTTCATATGTTGATAAACAAGAAATTACAGCAAGCTCAGAAATTGATGATTTGATTTTTTCGAGCGATCCATTAGTGTGGTCTTACGACGAGCAGGAAAAAATCAGAAAGAAACTTGTGCTTGATGCATTTCGTAATCATTATAAACATTGTCGAGAATATCGTCACTACTGTCAGGCACACAAAGTAGATGACAATATTACGGAAATTGATGACATACCTGTATTCCCAACATCGGTTTTTAAGTTTACTCGCTTATTAACTTCTCAGGAAAACGAGATTGAAAGTTGGTTTACCAGTAGCGGCACGAATGGTTTAAAAAGTCAGGTGGCGCGTGACAGATTAAGTATTGAGAGACTCTTAGGCTCTGTGAGTTATGGCATGAAATATGTTGGTAGTTGGTTTGATCATCAAATAGAATTAGTCAATTTGGGACCAGATAGATTTAATGCTCATAATATTTGGTTTAAATATGTTATGAGTTTGGTGGAATTGTTATATCCTACGACATTTACCGTAACAGAAGAACGAATAGATTTTGTTAAAACATTGAATAGTCTTGAACGAATAAAAAATCAAGGGAAAGATCTTTGTCTTATTGGTTCGCCATACTTTATTTATTTACTCTGCCATTATATGAAAGATAAAAAAATCTCATTTTCTGGAGATAAAAGCCTTTATATCATAACCGGAGGCGGCTGGAAAAGTTACGAAAAAGAATCTCTGAAACGTGATGATTTCAATCATCTTTTATTTGATACTTTCAATCTCAGTGATATTAGTCAGATCCGAGATATATTTAATCAAGTTGAACTCAACACTTGTTTCTTTGAGGATGAAATGCAGCGTAAACATGTTCCGCCGTGGGTATATGCGCGAGCGCTTGATCCTGAAACGTTGAAACCTGTACCTGATGGAACGCCGGGGTTGATGAGTTATATGGATGCGTCAGCAACCAGTTATCCAGCATTTATTGTTACCGATGATGTCGGGATAATTAGCAGAGAATATGGTAAGTATCCCGGCGTGCTCGTTGAAATTTTACGTCGCGTCAATACGAGGACGCAGAAAGGGTGTGCTTTAAGCTTAACCGAAGCGTTTGATAGTTAAGATACAATCGATTTCTGGATCCGCAGGCCTCTGCTAGCTTGACTGACTGAGATACAGCGTACCTTCAGCTCACAGACATGATAAGATACATTGATGAGTTTGGACAAACCACAACTAGAATGCAGTGAAAAAAATGCTTTATTTGTGAAATTTGTGATGCTATTGCTTTATTTGTAACCATTATAAGCTGCAATAAACAAGTTAACAACAACAATTGCATTCATTTTATGTTTCAGGTTCAGGGGGAGGTGTGGGAGGTTTTTTAAAGCAAACGCTAAATTGCCTGATGCGCTACGCTTATCATCCTTAGCGAAAGCTAAGGAAGGCACTGTATTGATCGGCACGTAAGAGGTTCCAACTTTCACCATAATGAAATAAGATCACTACCGGGCGTATTTTTTGAGTTATCGAGATTTTCAGGAGCTAAGGAAGCTAAAATGGAGAAAAAAATCACTGGATATACCACCGTTGATATATCCCAATGGCATCGTAAAGAACATTTTGAGGCATTTCAGTCAGTTGCTCAATGTACCTATAACCAGACCGTTCAGCTGGATATTACGGCCTTTTTAAAGACCGTAAAGAAAAATAAGCACAAGTTTTATCCGGCCTTTATTCACATTCTTGCCCGCCTGATGAATGCTCATCCGGAATTTCGTATGGCAATGAAAGACGGTGAGCTGGTGATATGGGATAGTGTTCACCCTTGTTACACCGTTTTCCATGAGCAAACTGAAACGTTTTCATCGCTCTGGAGTGAATACCACGACGATTTCCGGCAGTTTCTACACATATATTCGCAAGATGTGGCGTGTTACGGTGAAAACCTGGCCTATTTCCCTAAAGGGTTTATTGAGAATATGTTTTTCGTCTCAGCCAATCCCTGGGTGAGTTTCACCAGTTTTGATTTAAACGTGGCCAATATGGACAACTTCTTCGCCCCCGTTTTCACCATGGGCAAATATTATACGCAAGGCGACAAGGTGCTGATGCCGCTGGCGATTCAGGTTCATCATGCCGTTTGTGATGGCTTCCATGTCGGCAGAATGCTTAATGAATTACAACAGTACTGCGATGAGTGGCAGGGCGGGGCGTAATTTGATATCGAGCTCGCTTGGACTCCTGTTGATAGATCCAGTAATGACCTCAGAACTCCATCTGGATTTGTTCAGAACGCTCGGTTGCCGCCGGGCGTTTTTTATTGGTGAGAATCCAAGCACTAGTAACAATAACTTCGTATAAAGTATCCTATACGAAGTTATGCTTGG

>Conjugation_plasmid_Apra

ATCGATGCATAATGTGCCTGTCAAATGGACGAAGCAGGGATTCTGCAAACCCTATGCTACTCCGTCAAGCCGTCAATTGTCTGATTCGTTACCAATTATGACAACTTGACGGCTACATCATTCACTTTTTCTTCACAACCGGCACGGAACTCGCTCGGGCTGGCCCCGGTGCATTTTTTAAATACCCGCGAGAAATAGAGTTGATCGTCAAAACCAACATTGCGACCGACGGTGGCGATAGGCATCCGGGTGGTGCTCAAAAGCAGCTTCGCCTGGCTGATACGTTGGTCCTCGCGCCAGCTTAAGACGCTAATCCCTAACTGCTGGCGGAAAAGATGTGACAGACGCGACGGCGACAAGCAAACATGCTGTGCGACGCTGGCGATATCAAAATTGCTGTCTGCCAGGTGATCGCTGATGTACTGACAAGCCTCGCGTACCCGATTATCCATCGGTGGATGGAGCGACTCGTTAATCGCTTCCATGCGCCGCAGTAACAATTGCTCAAGCAGATTTATCGCCAGCAGCTCCGAATAGCGCCCTTCCCCTTGCCCGGCGTTAATGATTTGCCCAAACAGGTCGCTGAAATGCGGCTGGTGCGCTTCATCCGGGCGAAAGAACCCCGTATTGGCAAATATTGACGGCCAGTTAAGCCATTCATGCCAGTAGGCGCGCGGACGAAAGTAAACCCACTGGTGATACCATTCGCGAGCCTCCGGATGACGACCGTAGTGATGAATCTCTCCTGGCGGGAACAGCAAAATATCACCCGGTCGGCAAACAAATTCTCGTCCCTGATTTTTCACCACCCCCTGACCGCGAATGGTGAGATTGAGAATATAACCTTTCATTCCCAGCGGTCGGTCGATAAAAAAATCGAGATAACCGTTGGCCTCAATCGGCGTTAAACCCGCCACCAGATGGGCATTAAACGAGTATCCCGGCAGCAGGGGATCATTTTGCGCTTCAGCCATACTTTTCATACTCCCGCCATTCAGAGAAGAAACCAATTGTCCATATTGCATCAGACATTGCCGTCACTGCGTCTTTTACTGGCTCTTCTCGCTAACCAAACCGGTAACCCCGCTTATTAAAAGCATTCTGTAACAAAGCGGGACCAAAGCCATGACAAAAACGCGTAACAAAAGTGTCTATAATCACGGCAGAAAAGTCCACATTGATTATTTGCACGGCGTCACACTTTGCTATGCCATAGCATTTTTATCCATAAGATTAGCGGATCCTACCTGACGCTTTTTATCGCAACTCTCTACTGTTTCTCCATACCCGTTTTTTTGGGCTAGCGAATTCGTAGGGGTTACTGAAAAGTGAGCGGGAAAGAAGAGTTTCAGACCATCAAGGAGCGGGCCAAGCGCAAGCTGGAACGCGACATGGGTGCGGACCTGTTGGCCGCGCTCAACGACCCGAAAACCGTTGAAGTCATGCTCAACGCGGACGGCAAGGTGTGGCACGAACGCCTTGGCGAGCCGATGCGGTACATCTGCGACATGCGGCCCAGCCAGTCGCAGGCGATTATAGAAACGGTGGCCGGATTCCACGGCAAAGAGGTCACGCGGCATTCGCCCATCCTGGAAGGCGAGTTCCCCTTGGATGGCAGCCGCTTTGCCGGCCAATTGCCGCCGGTCGTGGCCGCGCCAACCTTTGCGATCCGCAAGCGCGCGGTCGCCATCTTCACGCTGGAACAGTACGTCGAGGCGGGCATCATGACCCGCGAGCAATACGAGGTCATTAAAAGCGCCGTCGCGGCGCATCGAAACATCCTCGTCATTGGCGGTACTGGCTCGGGCAAGACCACGCTCGTCAACGCGATCATCAATGAAATGGTCGCCTTCAACCCGTCTGAGCGCGTCGTCATCATCGAGGACACCGGCGAAATCCAGTGCGCCGCAGAGAACGCCGTCCAATACCACACCAGCATCGACGTCTCGATGACGCTGCTGCTCAAGACAACGCTGCGTATGCGCCCCGACCGCATCCTGGTCGGTGAGGTACGTGGCCCCGAAGCCCTTGATCTGTTGATGGCCTGGAACACCGGGCATGAAGGAGGTGCCGCCACCCTGCACGCAAACAACCCCAAAGCGGGCCTGAGCCGGCTCGCCATGCTTATCAGCATGCACCCGGATTCACCGAAACCCATTGAGCCGCTGATTGGCGAGGCGGTTCATGTGGTCGTCCATATCGCCAGGACCCCTAGCGGCCGTCGAGTGCAAGAAATTCTCGAAGTTCTTGGTTACGAGAACGGCCAGTACATCACCAAAACCCTGTAAGGAGTATTTCCAATGACAACGGCTGTTCCGTTCCGTCTGACCATGAATCGCGGCATTTTGTTCTACCTTGCCGTGTTCTTCGTTCTCGCTCTCGCGTTATCCGCGCATCCGGCGATGGCCTCGGAAGGCACCGGCGGCAGCTTGCCATATGAGAGCTGGCTGACGAACCTGCGCAACTCCGTAACCGGCCCGGTGGCCTTCGCGCTGTCCATCATCGGCATCGTCGTCGCCGGCGGCGTGCTGATCTTCGGCGGCGAACTCAACGCCTTCTTCCGAACCCTGATCTTCCTGGTTCTGGTGATGGCGCTGCTGGTCGGCGCGCAGAACGTGATGAGCACCTTCTTCGGTCGTGGTGCCGAAATCGCGGCCCTCGGCAACGGGGCGCTGCACCAGGTGCAAGTCGCGGCGGCGGATGCCGTGCGTGCGGTAGCGGCTGGACGGCTCGCCTAATCATGGCTCTGCGCACGATCCCCATCCGTCGCGCAGGCAACCGAGAAAACCTGTTCATGGGTGGTGATCGTGAACTGGTGATGTTCTCGGGCCTGATGGCGTTTGCGCTGATTTTCAGCGCCCAAGAGCTGCGGGCCACCGTGGTCGGTCTGATCCTGTGGTTCGGGGCGCTCTATGCGTTCCGAATCATGGCGAAGGCCGATCCGAAGATGCGGTTCGTGTACCTGCGTCACCGCCGGTACAAGCCGTATTACCCGGCCCGCTCGACCCCGTTCCGCGAGAACACCAATAGCCAAGGGAAGCAATACCGATGATCCAAGCAATTGCGATTGCAATCGCGGGCCTCGGCGCGCTTCTGTTGTTCATCCTCTTTGCCCGCATCCGCGCGGTCGATGCCGAACTGAAACTGAAAAAGCATCGTTCCAAGGACGCCGGCCTGGCCGATCTGCTCAACTACGCCGCTGTCGTCGATGACGGCGTAATCGTGGGCAAGAACGGCAGCTTTATGGCTGCCTGGCTGTACAAGGGCGATGACAACGCAAGCAGCACCGACCAGCAGCGCGAAGTAGTGTCCGCCCGCATCAACCAGGCCCTCGCGGGCCTGGGAAGTGGGTGGATGATCCATGTGGACGCCGTGCGGCGTCCTGCTCCGAACTACGCGGAGCGGGGCCTGTCGGCGTTCCCTGACCGTCTGACGGCAGCGATTGAAGAAGAGCGCCGGCGGCATTTCGAGAGCCTGGGAACGATGTACGAGGGCTATTTCGTCCTCACCTTGACCTGGTTCCCGCCGCTGCTCGCCCAGCGCAAGTTCGTCGAGCTGATGTTTGACGACGACGCGACCGCACCGGATCGCAAGGCGCGCACGCGGGGCCTCATCGACCAATTCAAGCGTGACGTGCGCAGCATCGAGTCGCGCCTGTCGTCGGCCGTGTCGCTCACTCGCTTGAAGGGGCACAAGATCGTCAACGAGGACGGCACGACCGTCACGCATGACGACTTCCTGCGCTGGCTGCAATTCTGCGTGACGGGCCTGCACCATCCGGTGCAGCTCCCCAGCAACCCGATGTACCTGGACGCCCTGGTCGGCGGACAGGAAATGTGGGGCGGGGTAGTGCCCAAGGTCGGCCGCAAGTTCGTCCAGGTGGTCGCTCTCGAAGGCTTCCCCTTGGAGTCCTATCCCGGCATCCTGACGGCGCTCGGCGAGCTGCCCTGCGAGTATCGGTGGTCGAGCCGGTTCATCTTCATGGACCAGCACGAAGCCGTGAAGCACCTCGACAAGTTCCGCAAGAAGTGGCGGCAGAAGATTCGCGGCTTCTTCGACCAGGTGTTCAACACGAACACCGGCCCGGTCGATCAGGACGCGCTTTCGATGGTGGCCGATGCTGAGGCGGCCATTGCCGAAGTCAACAGCGGCATCGTGGCCGTGGGCTACTACACCAGCGTCGTCGTGCTGATGGATGAGGACCGCACGCGCCTGGAAGCTGCGGCCCGCGATGTTGAAAAGGCCGTCAACCGGTTGGGCTTTGCCGCGCGCATCGAGTCCATCAACACCCTGGACGCCTTCCTTGGTAGTTTGCCGGGCCACGGCGTGGAAAACGTCCGCCGGCCGCTCATCAACACGATGAACCTGGCCGACCTGCTGCCGACCAGCACCATCTGGACCGGCAACGCGAACGCGCCATGCCCGATGTACCCGCCGCTGTCGCCGGCGCTCATGCACTGCGTCACGCAAGGATCAACGCCGTTCCGGCTGAACCTGCACGTGCGCGACCTCGGCCACACCTTTATGTTCGGGCCGACCGGCGCAGGTAAATCGACGCACCTGGCGATCCTCGCCGCGCAGCTCCGTCGCTATGCCGGCATGTCGATCTTCGCCTTTGACAAGGGCATGTCGATGTACCCGCTGGCCGCCGGCATCCGTGCGGCCACGAAGGGCACCAGCGGCCTGCACTTCACCGTGGCGGCCGACGACGAACGCCTGGCGTTCTGCCCGTTGCAGTTCCTGAGCACCAAGGGCGACCGTGCTTGGGCGATGGAGTGGATCGACACCATCCTGGCGTTGAACGGCGTCGAAACGACCCCGGCCCAGCGCAACGAAATCGGCAACGCGATCATGAGCATGCACGCCAGCGGCGCGCGCACGCTCTCCGAGTTCAGCGTGACGATTCAGGATGAGGCGATCCGCGAGGCGATCCGCCAGTACACCGTCGATGGCGCAATGGGCCATCTGCTCGACGCCGAAGAGGACGGCTTGGCGCTGTCCGACTTTACAGTGTTCGAGATCGAAGAGCTGATGAACCTCGGCGAGAAATTCGCCCTGCCTGTGTTGCTCTACCTGTTCCGCCGTATCGAGCGCGCCCTGACGGGCCAGCCGGCCGTCATCATCCTGGACGAAGCCTGGTTGATGCTCGGCCACCCGGCATTCCGCGCGAAGATCAGGGAATGGCTCAAGGTGCTGCGTAAGGCCAACTGCCTTGTGCTGATGGCAACGCAGAGCCTGTCCGACGCCGCCAACAGCGGCATCCTGGACGTGATCGTGGAATCGACCGCGACCAAGATTTTCCTGCCGAATATTTACGCCAGGGATGAGGACACGGCGGCCCTGTACCGCCGCATGGGCCTGAACGCTCGCCAGATCGAGATTCTGGCCCAGGCCGTTCCCAAGCGTCAGTACTACTACGTGTCGGAAAACGGCCGCCGTCTCTACGACCTGGCACTTGGCCCGCTCGCGCTCGCGTTCGTCGGCGCATCCGACAAGGAATCCGTCGCCATCATCAAGAACCTGGAAGCCAAGTTCGGCGACCAGTGGGTGGATGAATGGCTGCGTGGCCGGGGCCTCGCCCTTGATGAATACCTGGAGGCAGCATGAGTTTTGCAGACACGATCAAGGGCTTGATCTTCAAGAAGAAGCCCGCAACGGCCGCAGCAGCGGCGACGCCGGCCGCGACCGGCCCGCAAACCGACAACCCGTACCTGACGGCGCGGCGCACCTGGAACGACCACGTTGGTTCCGTTGTGTCGCAAAAGCAGACCTGGCAGGTTGTCGGCATCCTTTCGCTGATGATCGTCCTCGCGGCGGTCGGCGGCATCATCCACATCGGCAGCCAGTCGAAGTTCGTGCCCTATGTCTACGAGGTAGACAAGCTCGGGCAGACGGCCGCCGTGGGGCCGATGACCAGGGCGTCGAAAGCCGATCCGCGTGTCATTCACGCCTCGGTGGCTGAGTTCGTCGGCGATGCTCGCCTGGTGACGCCGGACGTAGCTTTGCAGCGCAAGGCCGTCTACCGCCTCTATGCCAAGCTCGGGCCGAATGACCCGGCCACCGCCAAGATGAACGAATGGCTCAACGGCACCGCCGACGCCAGCCCGTTCGCTCGCGCGGCCGTCGAAACGGTCAGCACCGAAATCACTTCCGTAATCCCGCAGACGCCCGACACCTGGCAGGTCGATTGGGTCGAGACGACGCGCGACAGGCAAGGCGTGGTGAAAGGCCAGCCCGTGCGCATGCGGGCCTTGGTGACGGTCTACGTCGTCGAGCCGACGGCGGACACCAAGGAAGAACAACTGCGAAACAACCCGGCCGGGATCTACGTCCGGGACTTCTCCTGGTCGAGACTTCTGTGAGGCACTGAATTATGAAAAAGGAACTGTTTGCTTTGGTCCTGGCCGCGTCCGTTAGCGTGCCTGCATTTGCCGCCGATCCCGGCGCGGACCTGACTGACCTCTATTTTTCCGGCAAGAACCCGGAGCTGACCGCGCAAGAGCGGGCGGCCATCGCCATCGCCAAGAAGTGGGAGGCGGGTACCGCCGGCATGCGGCCGGTGGCCGGCCCCGGTGGTTCGGTGCGCTTCCTGTTCGGCGCGCAGCAGCCGAGCATCGTATGCGCCGTGCTGCAAGTGTGCGACGTGGCCCTGCAACCCGGCGAGCAAGTCAACTCGATCAACCTGGGCGACACCGCCCGTTGGACGGTCGAGCCGGCCATTACCGGCAGCGGCGCGAACGAAACCCAGCACCTCATCATCAAGCCGATGGATGTGGGCCTGGAAACCAGCCTGGTCGTGACCACGGACCGCCGCAGCTACCACATGCGCCTGCGCTCGCATCGCACGCAGTACATGCCGCAGGTGTCGTTCACCTACCCGGAAGATGCCCTTGCGAAGTGGGACGCCATCAAGAACCGCGAACAGCGGGATCGCGTCGAGAAAACCATTCCGCAGACCGGCGAGTACCTGGGCAACCTGAGCTTCAACTACTCCGTCAGCGGGTCCACGTCGTGGAAGCCGGTGCGCGTCTACAACGACGGCAAGAAAACCATCATCCAGATGCCGCACTCGATGGAACAGACCGAAGCGCCGACGCTCCTGGTCGTTCGCAGGGAGGGCGGCCTGTTCTCCGACGATGAAACGGTGATGGTCAACTACCGGGTCCAGGGCGACCGCTACATCGTCGATACGATTTTCGACAAGGCCATCCTCATCGCGGGCGTGGGCAGCAGCCAGGACCGCGTGACCATTTCAAGGGGGAACTAAACCATGCGTAAGATTCTGACCGTCATCGCACTCGCGGCCACGTTGGCCGGCTGCGCGACCTCCAAGTACGGCAGCTTCGTCCAGGACGCGCCGGCCGCCTACAACCAGACCATTGCGACCGACGCGGTGAAGCAGCTCGTCAAGCTCTACCCGCCGGCGCAAACCAAGCTGGAATTGCAGCAGGCTACGCCCGATCCGTTCGGCATTGCCCTGGTCACTGACCTTCGCGCCCAGGGCTATGCTGTCATGGAGTACAAGCCCGACGGCAACGCGGCCGCAGCTCCGGCTGCTGCGTCCTCGGCCGCTGCGAAGCCGGCAACGCCGCAAGCCCAGGGCGGCTATCCGCTGCGCTACGTGCTGGACCAATTCAGCGACAGCAACCTGTATCGCCTGACCGTCATGGTCGGCTCTCAATCGCTCACGCGCGCCTACCTCGCCCAAAACAACACGATGGTCCCGGCCGGCGCATGGGTTCGGAAGGAGTAAGCCAATGAGCGAAGATCAAATGGCACCGGACGCATCGCCAGATGCGGTCAAGCCGAAAAGCGGGGTTCGCCGCGTCAACAACATGCCGATGTACCTCATCGGCGGTGTGCTCGGCATCTTCCTGCTGGTGATGGCCCTGGTTGCTGCGGATCGCGCTGCGCAGCAGAACCAGCCGGGAGCTGCGAAGGCTGAGAAGGCCGGCAGCACCAGCATGTTTGCCGACGAAATTGCCGGCAAACAGCAGGACGGCATCATCAAGGCCAAGCCGCTGGAGATTCCGCCGGAACAAACCGCCCAGCAACCGACGACGGAGCTGACGCCAGCCCCGGCGCAGGGAACGACTATCACGGTCGCACGGCCCGAGAACCTGGACCAGCCCCCGACGCCGCCGCAGGGTGCGCGCAACGAGGACCTGGACCGCATCCGCATGGCGAAGTTGCAGATGCTGGAAGAGGCGATCAAGGCCAAGACGACGGTGCGCATCGACGCGCCGCGCAGCCAGGGCAGCGCCGGCGGCGGTGCTCCGCAGGGCCGCGAGGAAACCCTTGCGCGCATCCAGGAGCTGCGTCGGCAGGCTGAGAACGCCCGCGCCACCGATCCGACCGCCGCCTATCAGGCCGCGCTTGCGCAGGCTCGCACGATGGGCGGCGCGGCAGGGGGTGGCGGTATGGGCGGCTCGGGTGCGCCGACCCTCGTGCAGACCTCGAACCGCAGTGGTGGCGGCGCTGGCTATGGGTCGTTCGACAACCGCAGCGAGGGCGACCGTTGGCGGCTCGACTCCCAGCCGGAAGCACCTGCAACGCCCTATGTGCTGCGCGCTGGCTTCGTCGTTCCGGCTACGCTTATCTCGGGCATCAACTCCGATCTGCCAGGCCAAATCATGGCCCAGGTATCGCAGTCGGTGTACGACACGGCGACCGGCAAGCACATGCTCATCCCCCAAGGCTCGCGCCTGGTGGGCAGCTACTCGAACGATGTGGCCTACGGGCAGAAGCGCGTTCTGGTGGCATGGCAGCGCATCATCTTCCCCGACGGCAAGGCAATGGACATTGGGGCCATGCCGGGCGGCGATAGCGCTGGGTATGCAGGCTTCAACGACAAGGTCAACAACCACTACTTCCGCACCTTCGCATCGGCATTCCTCATGTCGGGCGTCGTTGCGGGCATCAGCTTGAGTCAGGACCGTGGCAACAGCAACAGCGGTTACGGACGACAAGACGCGGGTTCCGCGATGAGTGAAGCGTTGGGTCAACAGCTCGGCCAAGTAACGGCGCAGATGATCGCCAAAAACTTGAATATCGCGCCGACGCTGGAAATCCGTCCGGGCTATCGCTTCAACGTCATTGTCACGAAAGACATGACGTTTTCTAAGCCCTACCAGGCGTTTGACTATTAACTCCAAGGAGTAACTTATGAAGAAGCTCGCTAAGAATGTTTTAGCCGCTAAAGTAGCTCTGGTGCTGGCCCTCTCGGTCGGCACCTTGGCGGTCACGCCTGCGCAAGCGGGCATTCCGGTCATCGACGGCACCAACCTGTCACAAACCACTGTCACCGCGATTCAGCAGGTTGCGCAGGTCCAGAAGCAAATCGAGGAATACCGGACGCAGTTGCAGCAGTACGAAAACATGCTGCAAAACACGGTGGCCCCGGCCGCCTACGTGTGGGACCAGGCGCAGTCCACCATCAACGGCCTGATGAGCGCCGTTGATACCCTGAACTACTACAAGAACCAGGCGGGCAGCATCGACGCTTACCTGGGCAAGTTCAAGGACGTGTCCTACTACAAGGGGTCGCCGTGCTTCTCCCTGTCGGGCTGCTCGGAAAGCGAGCGCAAGGCGATGGAAGAGAACCGCCGCCTGGCGTCCGAATCGCAGAAAAAGGCCAACGATGCGCTGTTCCGTGGCCTCGATCAGCAGCAGAGCAACCTCAAGTCCGACGCCGCCACGCTGGAGCAATTGAAGGGCAAGGCGACGACGGCGCAGGGCCAGTTGGAAGCCCTCGGCTACGCCAACCAGTTCGCCAGCCAGCAGGCCAACCAGCTCATGCAAATCCGTGGCCTTCTGCTTGCGCAGCAGAACGCCATCGCCACGCAGATGCAGGCCCAGCAGGACCGGCAGGCCCAGCAGGACGCTGCGGGCGCGAAGCTGCGCGAGGGTTCGTACCGCGCAAGCCCGTCTAAGACCTGGTGAGGGGAGGCGCGATGAAGAAATCCAACTTCATCGCAGTTGCCGCGCTGGCCGCCGTCATGGCGGCCAGCCTGGCAGGCTGCGACAACAAGCCCGACACCGACAAGCTGACCTGCGCCGATCTGCCGAAGGTCACGGATGCCGCTCAACGCGCGGAGCTGTTGAAGAAGTGCCCGCGCGGAGAACCGGGAGGCTTCAAGCCCAGCGAAAAGAAAGAGTGGTGATGACGTATGAAAATCCAGACTAGAGCTGCCGCGCTCGCGGTCCTGATGCTGGCCTTGATGCCGGTAGCGGCATACGCCCAAATCGACAATTCGGGCATCCTCGACAACGTATTGCAGCGCTACCAGAACGCCGCGAGCGGCTGGGCCACTGTCGTCCAGAACGCCGCAACCTGGCTGTTCTGGACCTTGACCGTGATTAGCATGGTCTGGACCTTCGGCATGATGGCACTGCGCAAGGCCGACATTGGCGAGTTCTTCGCCGAGTTCGTGCGGTTCACCATCTTCACCGGCTTCTTCTGGTGGCTGCTGACCAACGGCCCGAATTTCGCGTCGTCCATCTATGCGTCCCTGCGGCAGATTGCAGGCCAGGCAACGGGGTTGGGGCAGGGGCTTTCGCCGTCCGGCATCGTCGATGTTGGCTTCGAGATTTTCTTCAAGGTGATGGACGAAACCTCGTACTGGTCGCCGGTCGATAGCTTCGTCGGTGCCTCGTTGGCGGCCGCCATCCTCTGCATCCTGGCCCTGGTCGGCGTGAATATGCTTCTGCTCCTGGCGTCCGGATGGATTCTTGCCTACGGCGGTGTGTTCTTCCTGGGCTTCGGCGGCTCGCGCTGGACCTCGGACATGGCGATCAACTACTACAAGACCGTCCTCGGGGTCGCCGCGCAGCTCTTCGCAATGGTGCTGCTCGTAGGCATCGGCAAGACCTTCCTCGATGACTACTACAGCCGCATGAGCGAAGGCATCAACTTCAAGGAACTTGGAGTGATGCTGATCGTCGGCCTGATCCTGCTCGTTCTGGTCAACAAGGTGCCGCAGCTCATCGCCGGCATCATCACCGGCGCGAGCGTCGGCGGTGCTGGTATCGGCCAGTTCGGCGCTGGCACGCTCGTCGGTGCGGCCGCGACGGCCGGCGCGGCAATCGCAACTGGCGGCGCATCTATCGCGGCCGGCGCTGCGGCGGCGGCCGGTGGCGCGCAGGCCATCATGGCGGCCGCGTCGAAGGCCAGCGATAACGTCTCTGCCGGCACTGACATTCTGTCGAGCATGATGGGCGGCGGCGGTGGCGGCGGCGGTGGTAGCGCCGGCACCAGCGGCGGCGACGGCGGCGGCTCGGGTGGCGGCGGTGGCTCGGGCGGCGGTGAAACCCCGATGGCCTCGGCCGCCGGCGACAACAGCAGCGGCGCACGCGGCGGCAGTTCGGGCGGCGGCTCGGGTGGTGGCCGTTCGTCTGGCGGTATCGGTGCCACGGCGGCCAAGGGCGGCCGGATCGCGGCCGATACCGTCGCCAACCTGGCGAAAGGTGCCGGCTCGATTGCCAAGGCCAAGGCCGGCGAAATGCGCGCATCGGCCCAGGAACGCATCGGCGATACCGTAGGCGGCAAGATCGCGCAGGCAATTCGCGGCGCGGGTGCGGCGGCGCAGACCGCTGCAACCGTCGCCGATAGCAACAGCCAGGCGCAGGAACAACCTGCACCGGCACCCGCACCGTCGTTCGACGACAACAGCCTTTCCGCAAGCAACAACAGGGAAGCGGCCGCCGACGCGGATTCCGAAGTGGCGAGCTTCGTCAACAAGCCCGCCCAATCCTGAAACGACTCTTAGGAGCTACGTAAGGCTGTTTTGGCGGATGAGGGCGCAGTCAGAGCTTGGGCCTGTCAAACAAGCCCAGCTAACGCAAAAACAAGCCCGGCAACGCCGGGCTTTTTCATCTGCGCCTCTGCGATTCATAACGAGGCCCACACCACCAGGGCGGCCGCGAACAACACCATGCCGCCGGCGATCATGTTCATCATCGCCACGCGCCGCGCGTCCGCGACCTTGGCCGCGAGCTGGCGGCCAAGGCCGTCGTCGATTTCCCTGCGGATCGCTTCGGCCGCCTGCGCGGCGCTGTCCTTCATTACCTTCGCCATTGCGTCCTTGCTGGCCGCCAGGGCCGCGTTCAGCATCCGCTCCGCTTTGGCCTTGGCGTCCTCGCCCCAACGATGGGCGATCCCTTCCAGCTCTTCCTTGAACGCGGCAAGGATTTCCTCTTGCTTGGCCGCACTGTCGGCCATGAGCCGGGCGTTGATGGTATGCAGGATCAGCACCGGGTCGTCGCGGCCGACGGCGATGCCGTGCTTGGCCGCAATCTCCCGGATCAGCTCTTCAATCTGGTCGCTCATAGCACGGCCGCCGCGTCGAGCTGTTCAAACAGGCCGCGCCGCACGATCTTGAGGCGTTGCCGCGTCATGATCGTGAGCGATTCATCGGCCAGCGCCTGGTCGAACGTCAGCCGCTCTTGCAGCATGTCGCTGAAATCGCGGCCGTAGGTTTCTTCCTTGAGGGCCGGAATCTGGATGATGGACGACACGCGGGCCTTGTTGGCCGTGTACGCCTTCATCTGCTCAAAGCTCTTGCCCTCATGCTCGATAGGCCCCCAATACGGGTTCAGCCAGACCACGAAAAGCGCTTCGGCCGGGAACTGGCTGGCGAGCTGGGCGAAGCCGCTCACCGTGTCCAGGAGAGCCTGGCCGCCGGTGACGACGGTATGGATGACCAGCTCATGCCCCATTTCTTGCAGCAGAGCCGGCACCTGGTTGCTGATGAGGTAATGCGACAGAGGCACGAACGAGCTGGCACCGTTGTCGATCACCACGTCATCCTTGGTCGGCGCAATCAGCTCGACCAGGGTGTCGAAGTTGCGCGAGTTAATTTCGTCGCCGGCCATGATGTTCAGCCGGCGGACGTTCAGGGCCTTGTAGCCCTCGAACGTCGCGTTCACCGGGTCGGTGTCGATGCACAAGGGTGTCTGCCCCTTGTCCATCTTGTACTGCGCAATGATCGCGGCGATGGCCGACTTGCCGACCCCGCCCTTGCCCTGCAAAACCATGTGAATTTTCGCCATTACAGTAGATCCTTTTTGTCCGGTGTTGGGTTGAAGGTGAAGCCGGTCGGGGCCGCAGCGGGGGCCGGCTTTTCAGCCTTGCCCCCCTGCTTCGGCCGCCGTGGCTCCGGCGTCTTGGGTGCCGGCGCGGGTTCCGCAGCCTTGGCCTGCGGTGCGGGCACATCGGCGGGCTTGGCCTTGATGTGCCGCCTGGCGTGCGAGCGGAACGTCTCGTAGGAGAACTTGACCTTCCCCGTTTCCCGCATGTGCTCCCAAATGGTGACGAGCGCATAGCCGGACGCTAACGCCGCCTCGACATCCGCCCTCACCGCCAGGAACGCAACCGCAGCCTCATCACGCCGGCGCTTCTTGGCCGCGCGGGATTCAACCCACTCGGCCAGCTCGTCGGTGTAGCTCTTTGGCATCGTCTCTCGCCTGTCCCCTCAGTTCAGTAATTTCCTGCATTTGCCTGTTTCCAGTCGGTAGATATTCCACAAAACAGCAGGGAAGCAGCGCTTTTCCGCTGCATAACCCTGCTTCGGGGTCATTATAGCGATTTTTTCGGTATATCCATCCTTTTTCGCACGATATACAGGATTTTGCTTAATTTGAAAATTGGAATATCCATCACATAACGACATGTCGCAGCAATTTAATCCATATTTATGCTGTTTCCGACCTGACACCTGCGTGAGTTGTTCACGTATTTTTTCACGGTGTATCCAACGGCGTCAGTAGGCCCACCCGCGAGCGGGTGTTCCTTCTTCACTGTCCCTTATTCGCACCTGGCGGTGCTCAACGGGAATCCTGCTCTGCGAGGCTGGCCGGCTACCGCCGGCGTAACAGATGAGGGCAAGCGGATGGCTGATGAAACCAAGCCAACCAGGAAGGGCAGCCCACCTATCAAGGTGTACTGCCTTCCAGACGAACGAAGAGCGATTGAGGAAAAGGCGGCGGCGGCCGGCATGAGCCTGTCGGCCTACCTGCTGGCCGTCGGCCAGGGCTACAAAATCACGGGCGTCGTGGACTATGAGCACGTCCGCGAGCTGGCCCGCATCAATGGCGACCTGGGCCGCCTGGGCGGCCTGCTGAAACTCTGGCTCACCGACGACCCGCGCACGGCGCGGTTCGGTGATGCCACGATCCTCGCCCTGCTGGCGAAGATCGAAGAGAAGCAGGACGAGCTTGGCAAGGTCATGATGGGCGTGGTCCGCCCGAGGGCAGAGCCATGACTTTTTTAGCCGCTAAAACGGCCGGGGGGTGCGCGTGATTGCCAAGCACGTCCCCATGCGCTCCATCAAGAAGAGCGACTTCGCGGAGCTGGTGAAGTACATCACCGACGAGCAAGGCAAGACCGAGCGCCTGGGTCACGTGCGCGTCACGAACTGCGAGGCAAACACCCTGCCCGCTGTCATGGCCGAGGTGATGGCGACCCAGCACGGCAACACCCGTTCCGAGGCCGACAAGACCTATCACCTGCTGGTTAGCTTCCGCGCGGGAGAGAAGCCCGACGCGGAGACGTTGCGCGCGATTGAGGACCGCATCTGCGCTGGGCTTGGCTTCGCCGAGCATCAGCGCGTCAGTGCCGTGCATCACGACACCGACAACCTGCACATCCATATCGCCATCAACAAGATTCACCCGACCCGAAACACCATCCATGAGCCGTATCGGGCCTACCGCGCCCTCGCTGACCTCTGCGCGACGCTCGAACGGGACTACGGGCTTGAGCGTGACAATCACGAAACGCGGCAGCGCGTTTCCGAGAACCGCGCGAACGACATGGAGCGGCACGCGGGCGTGGAAAGCCTGGTCGGCTGGATCAAGCGCGAATGCCTGCCGGAGCTGCAAGCGGCGCAATCCTGGGAGGACCTGCACCGCGTCCTGCGGGAAAACGGGCTTAAGCTGCGCGAGCGCGGAAACGGCTTCATCTTCGAGGCCGGCGACGGCACGACGGTCAAGGCCAGCACCGTTTCGCGCGACCTGTCCAAGCCGAAGCTCGAAGCCCGATTCGGGGCATTCACGCCAGCCGAGGGCGGCGAAGCCCCCCGGCGGCGGGAGTATCGAGCGAAGCCGCTCAAGACCCGCATCGACACCACCGAGCTATACGCCCGGTATCAGTCTGAACGCCAGGAAATGGGGGCCGTGCGCAAGGGCGAGCTGGACACGCTGCGCCGGCGTCGTGACCGCCTGATTGAAGCCGCGATGCGCAGCAACCGGCTGCGCCGCGCCGCTATCAAGCTGCTGGGCGAGGGGCGCATTGCAAAGCGACTGATGTACGCGCAGGCGCACAAGGCTCTGCGCGCCGACCTGGACAAGATCAACCGCGAGTACCGGCAGGGCCGTCAGGCGGTCCAGGAGCGCACGCAGCGCCGCGCGTGGGCCGACTGGCTCAAGGCCGAGGCGATGAAGGGCGACGACAAGGCCCTGGCCGCGCTGCGCGCACGCGAGGGCCGCAGCGACCTCAAGGGCAACACCATCCAGGGCAGCGGCGAGGCCAAGCCGGGCCATGCGGCCGTGACGGACAACATCACGAAGAAGGGAACCATCATCTATCGCGTCGGCAGCAGCGCCGTCCGCGACGACGGCGACCGCCTGCAAGTCTCGCGCGAGGCCACCACGGACGGCCTGGACGCCGCCCTGCGCCTGGCAATGGAGCGCTTCGGCGACCGGATCACCGTCAACGGCACAGCAGAGTTCAAGGAGCGGATCGCGCAGGCGGCCGCCGCTGGCCGTCTGGCAATCACCTTTGACGACGCCGCCCTTGAGCGCCGCCGTCAAGAACTACTGACGAAGGAGCAAGCACATGAGCAACCCGAACGAAATGACGGACGAAGAGATCGCGGCGGCGATGGAGGCATTCGACCTGCCGCAGCCAGAACCACCCTCAACGCCACAGGCGGCGACGGCGACCGACGGGACGCTCGCGCCGTCAGCGCCGGCGGAACCGTCGCACTCCGCAAGCCCAACGTTGGACGCATTGGACGAAAGCCGCCGCCCCAAAGCCAAAACCGTTTGCGAGCGCTGTCCCAACTCGGTGTGGTTCGCATCGCCGGCGGAGCTGAAATGCTATTGCCGCGTGATGTTCCTGGTCACGTGGAGCAGCAAGGAGCCGAACCAGCTCACGCACTGCGACGGGGAGTTTCTGGGCCAGGAAGAGGGCTAAAGCCCGAACAGATCGCGGCGGCCGAGAAATATGTTGCCGAGCGCGAGCAAAAGCGGCTAAACGGCTTCGATATACCGAAGCATGCACGATATACTGATTATGTTGGTGCGCTGTCCTATGCAGGCACCCGGAACGTCGAGGACCAGGCGCTGGCCCTGTTGAGGAAAGAAAACGACGAGATTCTGGTGCTGCCCGTCGATAAGGCCACGGTGCAACGCATGAAGCGCCTCGCCATCGGTGATCCCGTCACCGTGACCCCGCGAGGCTCCCTTAAAACTACCCGAGGTAGGAGTAGATGAAGAACCGAAACAACGCCGTGGGGCCACAGATACGGGCGAAAAAACCGAAGGCCAGCAAGACCGTTCCCATCCTCGCCGGCCTGTCCCTTGGGGCAGGCTTGCAGACCGCGACGCAATATTTCGCCCATTCCTTCCAGTATCAGGCCGGACTTGGCTGGAATATCAACCACGTCTACACGCCCTGGTCGATCCTTCAATGGGCCGGCAAGTGGTACGGCCAATATCCCGACGATTTCATGCGCGCGGCCAGCATGGGTATGGTCGTTTCGACCGTGGGCCTGCTGGGCACGGCCGTCACGCAGATGGTCAAGGCGAACACCGGCAAGGCGAACGACTATCTGCACGGTTCGGCCCGCTGGGCCGACAAGAAGGACATACAGGCCGCCGGCCTGCTGCCCCGGCCGCGCACCGTCGTCGAGCTGGTGTCAGGCAAACACCCGCCCACTTCCAGCGGCGTCTATGTGGGCGGCTGGCAGGACAAGGACGGCAAGTTCCACTACCTGCGGCACAACGGCCCCGAGCACGTCTTGACCTACGCGCCGACGCGCTCCGGCAAGGGCGTCGGCCTGGTCGTTCCGACGCTGCTTTCCTGGGCGCACAGCGCCGTCATCACCGACCTGAAAGGCGAGTTGTGGGCGCTGACCGCCGGCTGGCGGAAGAAGCATGCCCGCAACAAGGTCGTGCGCTTTGAGCCGGCATCCGCGCAGGGTAGCGCGTGCTGGAACCCGCTCGATGAAATCCGCTTGGGGACCGAGTACGAGGTTGGCGACGTGCAGAACCTTGCCACCCTGATCGTCGATCCGGACGGCAAGGGCCTGGAATCGCACTGGCAGAAAACCAGCCAGGCGCTGCTTGTCGGCGTCATCCTGCACGCGCTCTACAAGGCTAAGAACGAGGGTACGCCGGCCACCCTGCCGTCGGTGGACGGCATGCTTGCCGACCCGAACCGCGACGTGGGCGAGCTTTGGATGGAAATGACCACCTACGGCCACGTTGACGGGCAGAACCACCCTGCGGTCGGCTCTGCGGCCCGCGACATGATGGACCGCCCGGAGGAAGAATCCGGTTCCGTGCTGTCCACCGCCAAGTCCTACCTGGCCCTGTACCGCGACCCGGTTGTGGCCCGCAACGTCAGCAAGTCCGACTTCCGCATCAAGCAACTGATGCACCACGACGACCCGGTAAGCCTGTTCATCGTGACGCAGCCCAACGACAAGGCCCGTCTGCGGCCTCTGGTGCGCGTCATGGTCAACATGATCGTCCGGCTGCTGGCCGACAAGATGGACTTCGAGAACGGCCGTCCTGTCGCGCACTACAAGCATCGTCTGCTGATGATGCTCGACGAGTTCCCCAGCCTGGGCAAGCTCGAAATCCTGCAAGAGTCCCTTGCCTTCGTCGCCGGCTACGGCATCAAGTGCTACCTCATCTGCCAGGACATTAACCAGCTCAAGAGCCGCGAAACCGGCTACGGCCACGACGAAAGCATCACGTCGAACTGCCACGTGCAGAACGCCTACCCGCCGAACCGCGTGGAGACGGCCGAGCACCTGTCAAAACTGACAGGCACCACGACCATCGTGAAGGAGCAGATCACGACGAGCGGCCGCCGCACGTCGGCGCTGTTGGGCAACGTCTCGCGGACCTTCCAGGAAGTGCAGCGGCCATTGCTGACCCCCGATGAATGCCTGCGCATGCCGGGGCCGAAGAAGAGCGCCGACGGCAGCATTGAAGAGGCGGGCGACATGGTTGTGTACGTCGCCGGCTATCCCGCGATCTACGGCAAGCAGCCGCTCTACTTCAAAGACCCGATATTCCAGGCCCGCGCGGCCGTACCGGCTCCGAAGGTCAGCGACAAGCTGATCCAGACGGCCACCGTCGAGGAAGGGGAGGGGATCACGATATGAGCCGCTTCCAGCGCCTCACCAAGTACGTCGCCATCGGAGGCGGCGCGGCCTTGTTGCTCGCCGGCGCGGCCTATCTCGCCGGCGCGAAGGTCAACACCACCAAAAGCATTCCGGTCGGCCTGTACTGGAAATCGAATGCGCCGGTGGAGAAGGGGGCTTACGTCATGTTCTGCCCGCCGCAAGTCGGCGTGTTTTCGGACGCCAAGGAGCGGGGCTACATCGCCGGCGGTTTCTGCCCCGGCGACTACGGCTACATGATGAAGCGAGTTTTAGCCGCTAAAGGCGACGAGGTTGCCATCACCGACGCCGGCGTGCGCGTGAATGGCGGGCTGCTGCCTCACAGCGCGCTCATCAAGGCCGATCCATCAGGACGGCCGCTGCCTCGTTATCAGTCCGACAGCTACACCCTGGGGACCGCCGAGGTTCTGCTTATGTCCGACGTAAGCGACACGTCTTTCGACGGCCGCTACTTCGGGCCTGTCAATCGTTCGCAAATCATCACCGTCATCAGGCCGGTTCTGACCTGGTGATAACCTTTTTGCGTTTCTACAAACTCTTTTGTTTATTTTTCTAAATACATTCAAATATGTATCCGCTCATGAGACAATAACCCTGATAAATGCTTCAATAATATTGAAAAAGGAAGAGTATGAGTATTCAACATTTCCGTGTCGCCCTTATTCCCTTTTTTGCGGCATTTTGCCTTCCTGTTTTTGCTCACCCAGAAACGCTGGTGAAAGTAAAAGATGCTGAAGATCAGTTGGGTGCACGAGTGGGTTACATCGAACTGGATCTCAACAGCGGTAAGATCCTTGAGAGTTTTCGCCCCGAAGAACGTTTTCCAATGATGAGCACTTTTAAAGTTCTGCTATGTGGCGCGGTATTATCCCGTGTTGACGCCGGGCAAGAGCAACTCGGTCGCCGCATACACTATTCTCAGAATGACTTGGTTGAGTACTCACCAGTCACAGAAAAGCATCTTACGGATGGCATGACAGTAAGAGAATTATGCAGTGCTGCCATAACCATGAGTGATAACACTGCGGCCAACTTACTTCTGACAACGATCGGAGGACCGAAGGAGCTAACCGCTTTTTTGCACAACATGGGGGATCATGTAACTCGCCTTGATCGTTGGGAACCGGAGCTGAATGAAGCCATACCAAACGACGAGCGTGACACCACGATGCCTGTAGCAATGGCAACAACGTTGCGCAAACTATTAACTGGCGAACTACTTACTCTAGCTTCCCGGCAACAATTAATAGACTGGATGGAGGCGGATAAAGTTGCAGGACCACTTCTGCGCTCGGCCCTTCCGGCTGGCTGGTTTATTGCTGATAAATCTGGAGCCGGTGAGCGTGGGTCTCGCGGTATCATTGCAGCACTGGGGCCAGATGGTAAGCCCTCCCGTATCGTAGTTATCTACACGACGGGGAGTCAGGCAACTATGGATGAACGAAATAGACAGATCGCTGAGATAGGTGCCTCACTGATTAAGCATTGGTAACTGTCAGACCAAGTTTACTCATATATACTTTAGATTGATTTACGCGCCCTGTAGCGGCGCATTAAGCGCGGCGGGTGTGGTGGTTACGCGCAGCGTGACCGCTACACTTGCCAGCGCCCTAGCGCCCGCTCCTTTCGCTTTCTTCCCTTCCTTTCTCGCCACGTTCGCCGGCTTTCCCCGTCAAGCTCTAAATCGGGGGCTCCCTTTAGGGTTCCGATTTAGTGCTTTACGGCACCTCGACCCCAAAAAACTTGATTTGGGTGATGGTTCACGTAGTGGGCCATCGCCCTGATAGACGGTTTTTCGCCCTTTGACGTTGGAGTCCACGTTCTTTAATAGTGGACTCTTGTTCCAAACTTGAACAACACTCAACCCTATCTCGGGCTATTCTTTTGATTTATAAGTCGACAGCGACACACTTGCATCGGATGCAGCCCGGTTAACGTGCCGGCACGGCCTGGGTAACCAGGTATTTTGTCCACATAACCGTGCGCAAAATGTTGTGGATAAGCAGGACACAGCAGCAATCCACAGCAGGCATACAACCGCACACCGAGGTTACTCCGTTCTACAGGTTACGACGACATGTCAATACTTGCCCTTGACAGGCATTGATGGAATCGTAGTCTCACGCTGATAGTCTGATCGACAATACAAGTGGGACCGTGGTCCCAGACCGATAATCAGACCGACAACACGAGTGGGATCGTGGTCCCAGACTAATAATCAGACCGACGATACGAGTGGGACCGTGGTCCCAGACTAATAATCAGACCGACGATACGAGTGGGACCGTGGTTCCAGACTAATAATCAGACCGACGATACGAGTGGGACCGTGGTCCCAGACTAATAATCAGACCGACGATACGAGTGGGACCATGGTCCCAGACTAATAATCAGACCGACGATACGAGTGGGACCGTGGTCCCAGTCTGATTATCAGACCGACGATACGAGTGGGACCGTGGTCCCAGACTAATAATCAGACCGACGATACGAGTGGGACCGTGGTCCCAGACTAATAATCAGACCGACGATACGAGTGGGACCGTGGTCCCAGTCTGATTATCAGACCGACGATACAAGTGGAACAGTGGGCCCAGAGAGAATATTCAGGCCAGTTATGCTTTCTGGCCTGTAACAAAGGACATTAAGTAAAGACAGATAAACGTAGACTAAAACGTGGTCGCATCAGGGTGCTGGCTTTTCAAGTTCCTTAAGAATGGCCTCAATTTTCTCTATACACTCAGTTGGAACACGAGACCTGTCCAGGTTAAGCACCATTTTATCGCCCTTATACAATACTGTCGCTCCAGGAGCAAACTGATGTCGTGAGCTTAAACTAGTTCTTGATGCAGATGACGTTTTAAGCACAGAAGTTAAAAGAGTGATAACTTCTTCAGCTTCAAATATCACCCCAGCTTTTTTCTGCTCATGAAGGTTAGATGCCTGCTGCTTAAGTAATTCCTCTTTATCTGTAAAGGCTTTTTGAAGTGCATCACCTGACCGGGCAGATAGTTCACCGGGGTGAGAAAAAAGAGCAACAACTGATTTAGGCAATTTGGCGGTGTTGATACAGCGGGTAATAATCTTACGTGAAATATTTTCCGCATCAGCCAGCGCAGAAATATTTCCAGCAAATTCATTCTGCAATCGGCTTGCATAACGCTGACCACGTTCATAAGCACTTGTTGGGCGATAATCGTTACCCAATCTGGATAATGCAGCCATCTGCTCATCATCCAGCTCGCCAACCAGAACACGATAATCACTTTCGGTAAGTGCAGCAGCTTTACGACGGCGACTCCCATCGGCAATTTCTATGACACCAGATACTCTTCGACCGAACGCCGGTGTCTGTTGACCAGTCAGTAGAAAAGAAGGGATGAGATCATCCAGTGCGTCCTCAGTAAGCAGCTCCTGGTCACGTTCATTACCTGACCATACCCGAGAGGTCTTCTCAACACTATCACCCCGGAGCACTTCAAGAGTAAACTTCACATCCCGACCACATACAGGCAAAGTAATGGCATTACCGCGAGCCATTACTCCTACGCGCGCAATTAACGAATCCACCATCGGGGCAGCTGGTGTCGATAACGAAGTATCTTCAACCGGTTGAGTATTGAGCGTATGTTTTGGAATAACAGGCGCACGCTTCATTATCTAATCTCCCAGCGTGGTTTAATCAGACGATCGAAAATTTCATTGCAGACAGGTTCCCAAATAGAAAGAGCATTTCTCCAGGCACCAGTTGAAGAGCGTTGATCAATGGCCTGTTCAAAAACAGTTCTCATCCGGATCTGACCTTTACCAACTTCATCCGTTTCACGTACAACATTTTTTAGAACCATGCTTCCCCAGGCATCCCGAATTTGCTCCTCCATCCACGGGGACTGAGAGCCATTACTATTGCTGTATTTGGTAAGCAAAATACGTACATCAGGCTCGAACCCTTTAAGATCAACGTTCTTGAGCAGATCACGAAGCATATCGAAAAACTGCAGTGCGGAGGTGTAGTCAAACAACTCAGCAGGCGTGGGAACAATCAGCACATCAGCAGCACATACGACATTAATCGTGCCGATACCCAGGTTAGGCGCGCTGTCAATAACTATGACATCATAGTCATGAGCAACAGTTTCAATGGCCAGTCGGAGCATCAGGTGTGGATCGGTGGGCAGTTTACCTTCATCAAATTTGCCCATTAACTCAGTTTCAATACGGTGCAGAGCCAGACAGGAAGGAATAATGTCAAGCCCCGGCCAGCAAGTGGGCTTTATTGCATAAGTGACATCGTCCTTTTCCCCAAGATAGAAAGGCAGGAGAGTGTCTTCTGCATGAATATGAAGATCTGGTACCCATCCGTGATACATTGAGGCTGTTCCCTGGGGGTCGTTACCTTCCACGAGCAAAACACGTAGCCCCTTCAGAGCCAGATCCTGAGCAAGATGAACAGAAACTGAGGTTTTGTAAACGCCACCTTTATGGGCAGCAACCCCGATCACCGGTGGAAATACGTCTTCAGCACGTCGCAATCGCGTACCAAACACATCACGCATATGATTAATTTGTTCAATTGTATAACCAACACGTTGCTCAACCCGTCCTCGAATTTCCATATCCGGGTGCGGTAGTCGCCCTGCTTTCTCGGCATCTCTGATAGCCTGAGAAGAAACCCCAACTAAATCCGCTGCTTCACCTATTCTCCAGCGCCGGGTTATTTTCCTCGCTTCCGGGCTGTCATCATTAAACTGTGCAATGGCGATAGCCTTCGTCATTTCATGACCAGCGTTTATGCACTGGTTAAGTGTTTCCATGAGTTTCATTCTGAACATCCTTTAATCATTGCTTTGCGTTTTTTTATTAAATCTTGCAATTTACTGCAAAGCAACAACAAAATCGCAAAGTCATCAAAAAACCGCAAAGTTGTTTAAAATAAGAGCAACACTACAAAAGGAGATAAGAAGAGCACATACCTCAGTCACTTATTATCACTAGCGCTCGCCGCAGCCGTGTAACCGAGCATAGCGAGCGAACTGGCGAGGAAGCAAAGAAGAACTGTTCTGTCAGATAGCTCTTACGCTCAGCGCAAGAAGAAATATCCACCGTGGGAAAAACTCCAGGTAGAGGTACACACGCGGATAGCCAATTCAGAGTAATAAACTGTGATAATCAACCCTCATCAATGATGACGAACTAACCCCCGATATCAGGTCACATGACGAAGGGAAAGAGAAGGAAATCAACTGTGACAAACTGCCCTCAAATTTGGCTTCCTTAAAAATTACAGTTCAAAAAGTATGAGAAAATCCATGCAGGCTGAAGGAAACAGCAAAACTGTGACAAATTACCCTCAGTAGGTCAGAACAAATGTGACGAACCACCCTCAAATCTGTGACAGATAACCCTCAGACTATCCTGTCGTCATGGAAGTGATATCGCGGAAGGAAAATACGATATGAGTCGTCTGGCGGCCTTTCTTTTTCTCAATGTATGAGAGGCGCATTGGAGTTCTGCTGTTGATCTCATTAACACAGACCTGCAGGAAGCGGCGGCGGAAGTCAGGCATACGCTGGTAACTTTGAGGCAGCTGGTAACGCTCTATGATCCAGTCGATTTTCAGAGAGACGATGCCTGAGCCATCCGGCTTACGATACTGACACAGGGATTCGTATAAACGCATGGCATACGGATTGGTGATTTCTTTTGTTTCACTAAGCCGAAACTGCGTAAACCGGTTCTGTAACCCGATAAAGAAGGGAATGAGATATGGGTTGATATGTACACTGTAAAGCCCTCTGGATGGACTGTGCGCACGTTTGATAAACCAAGGAAAAGATTCATAGCCTTTTTCATCGCCGGCATCCTCTTCAGGGCGATAAAAAACCACTTCCTTCCCCGCGAAACTCTTCAATGCCTGCCGTATATCCTTACTGGCTTCCGCAGAGGTCAATCCGAATATTTCAGCATATTTAGCAACATGGATCTCGCAGATACCGTCATGTTCCTGTAGGGTGCCATCAGATTTTCTGATCTGGTCAACGAACAGATACAGCATACGTTTTTGATCCCGGGAGAGACTATATGCCGCCTCAGTGAGGTCGTTTGACTGGACGATTCGCGGGCTATTTTTACGTTTCTTGTGATTGATAACCGCTGTTTCCGCCATGACAGATCCATGTGAAGTGTGACAAGTTTTTAGATTGTCACACTAAATAAAAAAGAGTCAATAAGCAGGGATAACTTTGTGAAAAAACAGCTTCTTCTGAGGGCAATTTGTCACAGGGTTAAGGGCAATTTGTCACAGACAGGACTGTCATTTGAGGGTGATTTGTCACACTGAAAGGGCAATTTGTCACAACACCTTCTCTAGAACCAGCATGGATAAAGGCCTACAAGGCGCTCTAAAAAAGAAGATCTAAAAACTATAAAAAAAATAATTATAAAAATATCCCCGTGGATAAGTGGATAACCCCAAGGGAAGTTTTTTCAGGCATCGTGTGTAAGCAGAATATATAAGTGCTGTTCCCTGGTGCTTCCTCGCTCACTCGAGGGCTTCGCCCTGTCGCTCGACTGCGGCGAGCACTACTGGCTGTAAAAGGACAGACCACATCATGGTTCTGTGTTCATTAGGTTGTTCTGTCCATTGCTGACATAATCCGCTCCACTTCAACGTAACACCGCACGAAGATTTCTATTGTTCCTGAAGGCATATTCAAATCGTTTTCGTTACCGCTTGCAGGCATCATGACAGAACACTACTTCCTATAAACGCTACACAGGCTCCTGAGATTAATAATGCGGATCTCTACGATAATGGGAGATTTTCCCGACTGTTTCGTTCGCTTCTCAGTGGATAACAGCCAGCTTCTCTGTTTAACAGACAAAAACAGCATATCCACTCAGTTCCACATTTCCATATAAAGGCCAAGGCATTTATTCTCAGGATAATTGTTTCAGCATCGCAACCGCATCAGACTCCGGCATCGCAAACTGCACCCGGTGCCGGGCAGCCACATCCAGCGCAAAAACCTTCGTGTAGACTTCCGTTGAACTGATGGACTTATGTCCCATCAGGCTTTGCAGAACTTTCAGCGGTATACCGGCATACAGCATGTGCATCGCATAGGAATGGCGGAACGTATGTGGTGTGACCGGAACAGAGAACGTCACACCGTCAGCAGCAGCGGCGGCAACCGCCTCCCCAATCCAGGTCCTGACCGTTCTGTCCGTCACTTCCCAGATCCGCGCTTTCTCTGTCCTTCCTGTGCGACGGTTACGCCGCTCCATGAGCTTATCGCGAATAAATACCTGTGACGGAAGATCACTTCGCAGAATAAATAAATCCTGGTGTCCCTGTTGATACCGGGAAGCCCTGGGCCAACTTTTGGCGCTAGCAAAAAAAATCCTTAGCTTTCGCTAAGGATGATAAGCGTAGCGCATCAGGCAATTTAGCGTTTGTTAAGCGATCGCGTATACACTCCGCTATCGCTACGTGACTGGGTCATGGCTGCGCCCCGACACCCGCCAACACCCGCTGACGCGCCCTGACGGGCTTGTCTGCTCCCGGCATCCGCTTACAGACAAGCTGTGACCGTCTCCGGGAGCTGCATGTGTCAGAGGTTTTCACCGTCATCACCGAAACGCGCGAGGCAGCAAGGAGATGGCGCCCAACAGTCCCCCGGCCACGGGGCCTGCCACCATACCCACGCCGAAACAAGCGCTCATGAGCCCGAAGTGGCGAGCCCGATCTTCCCCATCGGTGATGTCGGCGATATAGGCGCCAGCAACCGCACCTGTGGCGCCGGTGATGCCGGCCACGATGCGTCCGGCGTAGAGGATCTGCTCATGTTTGACAGCTTATC

>2lox>2oriT_Conjugation_Plasmid

AGTGTTATGAAATGGCCTGGTGATGATGGCGGGATCGTTGTATATTTCTTGACACCTTTTCGGCATCGCCCTAAAATTCGGCGTCCTCATATTGTGTGAGGACGTTTTATTACGTGTTTACGAAGCAAAAGCTAAAACCAGGAGCTATTTAATGGCAACAGTTAACCAGCTGGTACGCAAACCACGTGCTCGCAAAGTTGCGAAAAGCAACGTGCCTGCGCTGGAAGCATGCCCGCAAAAACGTGGCGTATGTACTCGTGTATATACTACCACTCCTAAAAAACCGAACTCCGCGCTGCGTAAAGTATGCCGTGTTCGTCTGACTAACGGTTTCGAAGTGACTTCCTACATCGGTGGTGAAGGTCACAACCTGCAGGAGCACTCCGTGATCCTGATCCGTGGCGGTCGTGTTAAAGACCTCCCGGGTGTTCGTTACCACACCGTACGTGGTGCGCTTGACTGCTCCGGCGTTAAAGACCGTAAGCAGGCTCGTTCCAAGTATGGCGTGAAGCGTCCTAAGGCTTAAGGAGGACAATCATGATTGAACAAGATGGATTGCACGCAGGTTCTCCGGCCGCTTGGGTGGAGAGGCTATTCGGCTATGACTGGGCACAACAGACAATCGGCTGCTCTGATGCCGCCGTGTTCCGGCTGTCAGCGCAGGGGCGCCCGGTTCTTTTTGTCAAGACCGACCTGTCCGGTGCCCTGAATGAACTGCAGGACGAGGCAGCGCGGCTATCGTGGCTGGCCACGACGGGCGTTCCTTGCGCAGCTGTGCTCGACGTTGTCACTGAAGCGGGAAGGGACTGGCTGCTATTGGGCGAAGTGCCGGGGCAGGATCTCCTGTCATCTCACCTTGCTCCTGCCGAGAAAGTATCCATCATGGCTGATGCAATGCGGCGGCTGCATACGCTTGATCCGGCTACCTGCCCATTCGACCACCAAGCGAAACATCGCATCGAGCGAGCACGTACTCGGATGGAAGCCGGTCTTGTCGATCAGGATGATCTGGACGAAGAGCATCAGGGGCTCGCGCCAGCCGAACTGTTCGCCAGGCTCAAGGCGCGCATGCCCGACGGCGAGGATCTCGTCGTGACCCATGGCGATGCCTGCTTGCCGAATATCATGGTGGAAAATGGCCGCTTTTCTGGATTCATCGACTGTGGCCGGCTGGGTGTGGCGGACCGCTATCAGGACATAGCGTTGGCTACCCGTGATATTGCTGAAGAGCTTGGCGGCGAATGGGCTGACCGCTTCCTCGTGCTTTACGGTATCGCCGCTCCCGATTCGCAGCGCATCGCCTTCTATCGCCTTCTTGACGAGTTCTTCTGAACAGGCGATGCTGCTTATCGAATCAAAGCTGCCGACAACACGGGAGCCAGTGACGCCTCCCGTGGGGAAAAAATCATGGCAATTCTGGAAGAAATAGCGCTTTCAGCCGGCAAACCGGCTGAAGCCGGATCTGCGATTCTGATAACAAACTAGCAACACCAGAACAGCCCGTTTGCGGGCAGCAAAACCCGTACCCTAGGTCTAGGGCGGCGGATTTGTCCTACTCAGGAGAGCGTTCAGTCTTTCGACTGAGCCTTTCGTTTTATTTGATGCCTGGCAGTTCCCTACTCTCGCTCATCTGTTACGCCGGCGGTAGCCGGCCAGCCTCGCAGAGCAGGATTCCCGTTGAGCACCGCCAGGTGCGAATAAGGGACAGTGAAGAAGGAACACCCGCTCGCGGGTGGGCCTACTTCACCTATCCTGCCCGGCTGACGCCGTTGGATACACCAAGGAAAGTCTACACGAACCCTTTGGCAAAATCCTGTATATCGTGCGAAAAAGGATGGATATACCGAAAAAATCGCTATAATGACCCCGAAGCAGGGTTATGCAGCGGAAAAGCGCTGGAGACCCCACACTACCATCGGCGCTACGGCGTTTCACTTCTGAGTTCGGCATGGGGTCAGGTGGGACCACCGCGCTACTGCCGCCAGGCAAATTCTGTTTTATCAGACCGCTTCTGCGTTCTGATTTAATCTGTATCAGGCTGAAAATCTTCTCTCATCCGCCAAAACAGCCAAGCTTGCATGCCTGCAGATAACTTCGTATAGGATACTTTATACGAAGTTATCTGCAGGTCGACATCGATGTTTAAACTTACTTATAAAGTTCGTCCATATCGTGTGTAATTCCAGCTGCTGTAACGCGCTCTTTCAATACCATGTGGTCGCGCTTTTCATTTGGGTCCTTACTCAGTTTACTCTGGTGGCTAAGATAGTGATTGTCGGGAAGCAAAACCGGACCATCGCCAATCGGTGTATTTTGCTGGTAGTGGTCCGCCAACTGGACTCCACCATCTTCTACGTTGTGGCGGGTCTTAAAATTTGCCTTGATGCCATTTTTTTGCTTATCTGCTGTGATATAAACCTTATGTGAATTGAAGTTGTACTCTAATTTGTGGCCTAAAATGTTACCGTCCTCTTTGAAGTCGATGCCTTTAAGGACGATACGGTTCACAAGGGTATCACCTTCGAACTTTACTTCAGCACGAGTTTTGTACGTTCCGTCGTCTTTAAAACTGATAGTGCGCTCCTGAACATACCCTTCGGGCATGGCAGATTTAAAGAAATCGTGCTGTTTCATGTGGTCAGGGTAACGGGCAAAACATGCTACCCCATACCCAAGCGTAGTGACCAGGGTCGGCCACGGGACCGGTAATTTGCCTGTAGTACAAATAAACTTTAAAGTAAGTTTACCGTTCGTCGCATCGCCCTCACCCTCGCCGCGCACGCTAAACTTATGGCCATTCACGTCACCATCAAGCTCGACTAAGATTGGAACCACCCCCGTGAAAAGCTCCTCACCTTTCGACACCATCTAGTATTTCCCCTCTTTCTCTAGTATTAAACAAAATTATTTGTAGAGGCTGTTTCGTCCTCACGGACTCATCAGACCGGAAAGCACATCCGGTGACAGGCTAGCATTATACCTAGGACTGAGCTAGCTGTCAAGGACTTGAGTGAGGTTGTAAAGGGAGTTGGCTCTGAATTCGCGGCCGCTTCTAGAGGAAATCATAAAAAATTTATTTGCTTTGTGAGCGGATAACAATTATAATAGATTCAATTGTGAGCGGATAACAATTACTAGAGAAAGAGGAGAAATACTAGATGAAATTTGGAAACTTTTTGCTTACATACCAACCTCCCCAATTTTCTCAAACAGAGGTAATGAAACGTTTGGTTAAATTAGGTCGCATCTCTGAGGAGTGTGGTTTTGATACCGTATGGTTACTGGAGCATCATTTCACGGAGTTTGGTTTGCTTGGTAACCCTTATGTCGCTGCTGCATATTTACTTGGCGCGACTAAAAAATTGAATGTAGGAACTGCCGCTATTGTTCTTCCCACAGCCCATCCAGTACGCCAACTTGAAGATGTGAATTTATTGGATCAAATGTCAAAAGGACGATTTCGGTTTGGTATTTGCCGAGGGCTTTACAACAAGGACTTTCGCGTATTCGGCACAGATATGAATAACAGTCGCGCCTTAGCGGAATGCTGGTACGGGCTGATAAAGAATGGCATGACAGAGGGATATATGGAAGCTGATAATGAACATATCAAGTTCCATAAGGTAAAAGTAAACCCCGCGGCGTATAGCAGAGGTGGCGCACCGGTTTATGTGGTGGCTGAATCAGCTTCGACGACTGAGTGGGCTGCTCAATTTGGCCTACCGATGATATTAAGTTGGATTATAAATACTAACGAAAAGAAAGCACAACTTGAGCTTTATAATGAAGTGGCTCAAGAATATGGGCACGATATTCATAATATCGACCATTGCTTATCATATATAACATCTGTAGATCATGACTCAATTAAAGCGAAAGAGATTTGCCGGAAATTTCTGGGGCATTGGTATGATTCTTATGTGAATGCTACGACTATTTTTGATGATTCAGACCAAACAAGAGGTTATGATTTCAATAAAGGGCAGTGGCGTGACTTTGTATTAAAAGGACATAAAGATACTAATCGCCGTATTGATTACAGTTACGAAATCAATCCCGTGGGAACGCCGCAGGAATGTATTGACATAATTCAAAAAGACATTGATGCTACAGGAATATCAAATATTTGTTGTGGATTTGAAGCTAATGGAACAGTAGACGAAATTATTGCTTCCATGAAGCTCTTCCAGTCTGATGTCATGCCATTTCTTAAAGAAAAACAACGTTCGCTATTATATTAATAATACTAGAGAAAGAGGAGAAATACTAGATGAAATTTGGATTGTTCTTCCTTAACTTCATCAATTCAACAACTGTTCAAGAACAAAGTATAGTTCGCATGCAGGAAATAACGGAGTATGTTGATAAGTTGAATTTTGAACAGATTTTAGTGTATGAAAATCATTTTTCAGATAATGGTGTTGTCGGCGCTCCTCTGACTGTTTCTGGTTTTCTGCTCGGTTTAACAGAGAAAATTAAAATTGGTTCATTAAATCACATCATTACAACTCATCATCCTGTCCGCATAGCGGAGGAAGCTTGCTTATTGGATCAGTTAAGTGAAGGGAGATTTATTTTAGGGTTTAGTGATTGCGAAAAAAAAGATGAAATGCATTTTTTTAATCGCCCGGTTGAATATCAACAGCAACTATTTGAAGAGTGTTATGAAATCATTAACGATGCTTTAACAACAGGCTATTGTAATCCAGATAACGATTTTTATAGCTTCCCTAAAATATCTGTAAATCCCCATGCTTATACGCCAGGCGGACCTCGGAAATATGTAACAGCAACCAGTCATCATATTGTTGAGTGGGCGGCCAAAAAAGGTATTCCTCTCATCTTTAAGTGGGATGATTCTAATGATGTTAGATATGAATATGCTGAAAGATATAAAGCCGTTGCGGATAAATATGACGTTGACCTATCAGAGATAGACCATCAGTTAATGATATTAGTTAACTATAACGAAGATAGTAATAAAGCTAAACAAGAGACGCGTGCATTTATTAGTGATTATGTTCTTGAAATGCACCCTAATGAAAATTTCGAAAATAAACTTGAAGAAATAATTGCAGAAAACGCTGTCGGAAATTATACGGAGTGTATAACTGCGGCTAAGTTGGCAATTGAAAAGTGTGGTGCGAAAAGTGTATTGCTGTCCTTTGAACCAATGAATGATTTGATGAGCCAAAAAAATGTAATCAATATTGTTGATGATAATATTAAGAAGTACCACATGGAATATACCTAATAATACTAGAGAAAGAGGAGAAATACTAGATGACTAAAAAAATTTCATTCATTATTAACGGCCAGGTTGAAATCTTTCCCGAAAGTGATGATTTAGTGCAATCCATTAATTTTGGTGATAATAGTGTTTACCTGCCAATATTGAATGACTCTCATGTAAAAAACATTATTGATTGTAATGGAAATAACGAATTACGGTTGCATAACATTGTCAATTTTCTCTATACGGTAGGGCAAAGATGGAAAAATGAAGAATACTCAAGACGCAGGACATACATTCGTGACTTAAAAAAATATATGGGATATTCAGAAGAAATGGCTAAGCTAGAGGCCAATTGGATATCTATGATTTTATGTTCTAAAGGCGGCCTTTATGATGTTGTAGAAAATGAACTTGGTTCTCGCCATATCATGGATGAATGGCTACCTCAGGATGAAAGTTATGTTCGGGCTTTTCCGAAAGGTAAATCTGTACATCTGTTGGCAGGTAATGTTCCATTATCTGGGATCATGTCTATATTACGCGCAATTTTAACTAAGAATCAGTGTATTATAAAAACATCGTCAACCGATCCTTTTACCGCTAATGCATTAGCGTTAAGTTTTATTGATGTAGACCCTAATCATCCGATAACGCGCTCTTTATCTGTTATATATTGGCCCCACCAAGGTGATACATCACTCGCAAAAGAAATTATGCGACATGCGGATGTTATTGTCGCTTGGGGAGGGCCAGATGCGATTAATTGGGCGGTAGAGCATGCGCCATCTTATGCTGATGTGATTAAATTTGGTTCTAAAAAGAGTCTTTGCATTATCGATAATCCTGTTGATTTGACGTCCGCAGCGACAGGTGCGGCTCATGATGTTTGTTTTTACGATCAGCGAGCTTGTTTTTCTGCCCAAAACATATATTACATGGGAAATCATTATGAGGAATTTAAGTTAGCGTTGATAGAAAAACTTAATCTATATGCGCATATATTACCGAATGCCAAAAAAGATTTTGATGAAAAGGCGGCCTATTCTTTAGTTCAAAAAGAAAGCTTGTTTGCTGGATTAAAAGTAGAGGTGGATATTCATCAACGTTGGATGATTATTGAGTCAAATGCAGGTGTGGAATTTAATCAACCACTTGGCAGATGTGTGTACCTTCATCACGTCGATAATATTGAGCAAATATTGCCTTATGTTCAAAAAAATAAGACGCAAACCATATCTATTTTTCCTTGGGAGTCATCATTTAAATATCGAGATGCGTTAGCATTAAAAGGTGCGGAAAGGATTGTAGAAGCAGGAATGAATAACATATTTCGAGTTGGTGGATCTCATGACGGAATGAGACCGTTGCAACGATTAGTGACATATATTTCTCATGAAAGGCCATCTAACTATACGGCTAAGGATGTTGCGGTTGAAATAGAACAGACTCGATTCCTGGAAGAAGATAAGTTCCTTGTATTTGTCCCATAATAATACTAGAGAAAGAGGAGAAATACTAGATGGAAAATGAATCAAAATATAAAACCATCGACCACGTTATTTGTGTTGAAGGAAATAAAAAAATTCATGTTTGGGAAACGCTGCCAGAAGAAAACAGCCCAAAGAGAAAGAATGCCATTATTATTGCGTCTGGTTTTGCCCGCAGGATGGATCATTTTGCTGGTCTGGCGGAATATTTATCGCGGAATGGATTTCATGTGATCCGCTATGATTCGCTTCACCACGTTGGATTGAGTTCAGGGACAATTGATGAATTTACAATGTCTATAGGAAAGCAGAGCTTGTTAGCAGTGGTTGATTGGTTAACTACACGAAAAATAAATAACTTCGGTATGTTGGCTTCAAGCTTATCTGCGCGGATAGCTTATGCAAGCCTATCTGAAATCAATGCTTCGTTTTTAATCACCGCAGTCGGTGTTGTTAACTTAAGATATTCTCTTGAAAGAGCTTTAGGGTTTGATTATCTCAGTCTACCCATTAATGAATTGCCGGATAATCTGGATTTTGAAGGCCATAAATTGGGTGCTGAAGTCTTTGCGAGAGATTGTCTTGATTTTGGTTGGGAAGATTTAGCTTCTACAATTAATAACATGATGTATCTTGATATACCGTTTATTGCTTTTACTGCAAATAACGATAATTGGGTCAAGCAAGATGAAGTTATCACATTGTTATCAAATATTCGTAGTAATCGATGCAAGATATATTCTTTGTTAGGAAGTTCGCATGACTTGAGTGAAAATTTAGTGGTCCTGCGCAATTTTTATCAATCGGTTACGAAAGCCGCTATCGCGATGGATAATGATCATCTGGATATTGATGTTGATATTACTGAACCGTCATTTGAACATTTAACTATTGCGACAGTCAATGAACGCCGAATGAGAATTGAGATTGAAAATCAAGCAATTTCTCTGTCTTAATAATACTAGAGAAAGAGGAGAAATACTAGATGACTTCATATGTTGATAAACAAGAAATTACAGCAAGCTCAGAAATTGATGATTTGATTTTTTCGAGCGATCCATTAGTGTGGTCTTACGACGAGCAGGAAAAAATCAGAAAGAAACTTGTGCTTGATGCATTTCGTAATCATTATAAACATTGTCGAGAATATCGTCACTACTGTCAGGCACACAAAGTAGATGACAATATTACGGAAATTGATGACATACCTGTATTCCCAACATCGGTTTTTAAGTTTACTCGCTTATTAACTTCTCAGGAAAACGAGATTGAAAGTTGGTTTACCAGTAGCGGCACGAATGGTTTAAAAAGTCAGGTGGCGCGTGACAGATTAAGTATTGAGAGACTCTTAGGCTCTGTGAGTTATGGCATGAAATATGTTGGTAGTTGGTTTGATCATCAAATAGAATTAGTCAATTTGGGACCAGATAGATTTAATGCTCATAATATTTGGTTTAAATATGTTATGAGTTTGGTGGAATTGTTATATCCTACGACATTTACCGTAACAGAAGAACGAATAGATTTTGTTAAAACATTGAATAGTCTTGAACGAATAAAAAATCAAGGGAAAGATCTTTGTCTTATTGGTTCGCCATACTTTATTTATTTACTCTGCCATTATATGAAAGATAAAAAAATCTCATTTTCTGGAGATAAAAGCCTTTATATCATAACCGGAGGCGGCTGGAAAAGTTACGAAAAAGAATCTCTGAAACGTGATGATTTCAATCATCTTTTATTTGATACTTTCAATCTCAGTGATATTAGTCAGATCCGAGATATATTTAATCAAGTTGAACTCAACACTTGTTTCTTTGAGGATGAAATGCAGCGTAAACATGTTCCGCCGTGGGTATATGCGCGAGCGCTTGATCCTGAAACGTTGAAACCTGTACCTGATGGAACGCCGGGGTTGATGAGTTATATGGATGCGTCAGCAACCAGTTATCCAGCATTTATTGTTACCGATGATGTCGGGATAATTAGCAGAGAATATGGTAAGTATCCCGGCGTGCTCGTTGAAATTTTACGTCGCGTCAATACGAGGACGCAGAAAGGGTGTGCTTTAAGCTTAACCGAAGCGTTTGATAGTTAAGATACAATCGATTTCTGGATCCGCAGGCCTCTGCTAGCTTGACTGACTGAGATACAGCGTACCTTCAGCTCACAGACATGATAAGATACATTGATGAGTTTGGACAAACCACAACTAGAATGCAGTGAAAAAAATGCTTTATTTGTGAAATTTGTGATGCTATTGCTTTATTTGTAACCATTATAAGCTGCAATAAACAAGTTAACAACAACAATTGCATTCATTTTATGTTTCAGGTTCAGGGGGAGGTGTGGGAGGTTTTTTAAAGCAAACGCTAAATTGCCTGATGCGCTACGCTTATCATCCTTAGCGAAAGCTAAGGAACAAGACGCTGGGCTCTGACATTTCCGCTACTGAACTTACTCGCGCGAAGGCGTCTAGATTATTCCTTTGCCCTCGGACGAGTGCTGGGGCGTCGGTTTCCACTATCGGCAAGTACTTCTACACAGCCATCGGTCCAGACGGCCGCGCTTCTGCGGGCGATTTGTGTACGCCCGACAGTCCCGGCTCCGGATCGGACGATTGCGTCGCATCGACCCTGCGCCCAAGCTGCATCATCGAAATTGCCGTCAACCAAGCTCTGATAGAGTTGGTCAAGACCAATGCGGAGCATATACGCCCGGAGCCGCGGCGATCCTGCAAGCTCCGGATGCCTCCGCTCGAAGTAGCGCGTCTGCTGCTCCATACAAGCCAACCACGGCCTCCAGAAGAAGATGTTGGCGACCTCGTATTGGGAATCCCCGAACATCGCCTCGCTCCAGTCAATGACCGCTGTTATGCGGCCATTGTCCGTCAGGACATTGTTGGAGCCGAAATCCGCGTGCACGAGGTGCCGGACTTCGGGGCAGTCCTCGGCCCAAAGCATCAGCTCATCGAGAGCCTGCGCGACGGACGCACTGACGGTGTCGTCCATCACAGTTTGCCAGTGATACACATGGGGATCAGCAATCGCGCATATGAAATCACGCCATGTAGTGTATTGACCGATTCCTTGCGGTCCGAATGGGCCGAACCCGCTCGTCTGGCTAAGATCGGCCGCAGCGATCGCATCCATGGCCTCCGCGACCGGCTGCAGAACAGCGGGCAGTTCGGTTTCAGGCAGGTCTTGCAACGTGACACCCTGTGCACGGCGGGAGATGCAATAGGTCAGGCTCTCGCTGAATTCCCCAATGTCAAGCACTTCCGGAATCGGGAGCGCGGCCGATGCAAAGTGCCGATAAACATAACGATCTTTGTAGAAACCATCGGCGCAGCTATTTACCCGCAGGACATATCCACGCCCTCCTACATCGAAGCTGAAAGCACGAGATTCTTCGCCCTCCGAGAGCTGCATCAGGTCGGAGACGCTGTCGAACTTTTCGATCAGAAACTTCTCGACAGACGTCGCGGTGAGTTCAGGCTTTTTCATGATTGTCCTCCTTATTTAAACTGTTTGAGGAAACGCAGATCGTTTTCGAAGAATGAACGCAGGTCGGTGACGCCGTAACGCAACATAGTCAGACGCTCCATCCCCATCCCGAAGCCGAAACCAGAGTAAACTTCCGGGTCGATGCCAACGTTACGCAACACGTTCGGATGCACCATCCCGCAGCCCAGCACTTCCAGCCATTTACCGTTTTTACCCATGACGTCCACTTCTGCAGAAGGTTCGGCAAACGGGAAGTAGGAAGGACGGAAGCGAATCTGCAAATCTTCCTCAAAGAAGTTACGCAGGAAGTCGTGCAGCGTGCCTTTCAGGTTGGTAAAGCTGATGTTGGTATCAACAATCAGACCTTCCATCTGATGGAACATCGGCGTGTGAGTCTGGTCGTAGTCGTTACGATAAACACGGCCAGGCGCGATGATACGAATCGGTGGCTGCTGGGCTTTCATGGTGCGGATCTGTACGCCAGAGGTCTGGGTACGCAGCAGGCGGGTAGTGTCAAACCAGAAAGTGTCGTGGTCAGCGCGCGCCGGGTGGTGACCAGGAATGTTCAGAGCATCGAAGTTATGATAATCGTCTTCGATTTCCGGCCCGGTTGCCACGGTAAAGCCAAGCTCACCGAAGAAACTTTCGATACGGTCGATGGTACGGGTAACCGGATGCAGACCGCCGTTTTCAATGCGACGACCTGGCAGAGAGACATCAATCGTTTCCGCCGCCAGACGCGCATTCAGTGCAGCGCTTTCCAGTTCCGCTTTACGCGCATTCAGCGCCTGCTGAACCTGCTCTTTCGCTTCGTTGATAACCGCACCAGCTGCCGGACGCTCTTCTGGCGGCAGCTCACGCAGGGTCGTCATCTGAAGGGTTAAGTGCCCTTTTTTACCCAAATATTCGACGCGCACATTATCTAACGCGGCAACATCTGACGCCTGGCTAATGGCCGCCTTCGCACTGGCAACCAGTTCTGCGAGATGTGACATGGTTTAGTTCCTCACCTTGTCGTATTATACTATGCCGATATACTATGCCGATGATTAATTGTCAACGCGGCCGCGCATAACTTCGTATAGCATACATTATACGAAGTTATGGCGCGCCTGTCTAACAATTCGTTCAAGCCGACGGATATCACGCGTGAGCTCTCATCTGTTACGCCGGCGGTAGCCGGCCAGCCTCGCAGAGCAGGATTCCCGTTGAGCACCGCCAGGTGCGAATAAGGGACAGTGAAGAAGGAACACCCGCTCGCGGGTGGGCCTACTTCACCTATCCTGCCCGGCTGACGCCGTTGGATACACCAAGGAAAGTCTACACGAACCCTTTGGCAAAATCCTGTATATCGTGCGAAAAAGGATGGATATACCGAAAAAATCGCTATAATGACCCCGAAGCAGGGTTATGCAGCGGAAAAGCGCTGAGCTCAGGACATACATTACGCTAAATTGCCTGATGCGCTACGCTTATCATCCTTAGCGAAAGCTAAGGATGTATTTAGAAAAATAAACAAATAGGGGTTCCGCGCACATTTCCCCGAAAAGTGCCACCTGGGTCCTTTTCATCACGTGCTATAAAAATAATTATAATTTAAATTTTTTAATATAAATATATAAATTAAAAATAGAAAGTAAAAAAAGAAATTAAAGAAAAAATAGTTTTTGTTTTCCGAAGATGTAAAAGACTCTAGGGGGATCGCCAACAAATACTACCTTTTATCTTGCTCTTCCTGCTCTCAGGTATTAATGCCGAATTGTTTCATCTTGTCTGTGTAGAAGACCACACACGAAAATCCTGTGATTTTACATTTTACTTATCGTTAATCGAATGTATATCTATTTAATCTGCTTTTCTTGTCTAATAAATATATATGTAAAGTACGCTTTTTGTTGAAATTTTTTAAACCTTTGTTTATTTTTTTTTCTTCATTCCGTAACTCTTCTACCTTCTTTATTTACTTTCTAAAATCCAAATACAAAACATAAAAATAAATAAACACAGAGTAAATTCCCAAATTATTCCATCATTAAAAGATACGAGGCGCGTGTAAGTTACAGGCAAGCGATCCGTCTAAGAAACCATTATTATCATGACATTAACCTATAAAAATAGGCGTATCACGAGGCCCTTTCGTCTCGCGCGTTTCGGTGATGACGGTGAAAACCTCTGACACATGCAGCTCCCGGAGACGGTCACAGCTTGTCTGTAAGCGGATGCCGGGAGCAGACAAGCCCGTCAGGGCGCGTCAGCGGGTGTTGGCGGGTGTCGGGGCTGGCTTAACTATGCGGCATCAGAGCAGATTGTACTGAGAGTGCACCATACCACCTTTTCAATTCATCATTTTTTTTTTATTCTTTTTTTTGATTTCGGTTTCCTTGAAATTTTTTTGATTCGGTAATCTCCGAACAGAAGGAAGAACGAAGGAAGGAGCACAGACTTAGATTGGTATATATACGCATATGTAGTGTTGAAGAAACATGAAATTGCCCAGTATTCTTAACCCAACTGCACAGAACAAAAACCTGCAGGAAACGAAGATAAATCATGTCGAAAGCTACATATAAGGAACGTGCTGCTACTCATCCTAGTCCTGTTGCTGCCAAGCTATTTAATATCATGCACGAAAAGCAAACAAACTTGTGTGCTTCATTGGATGTTCGTACCACCAAGGAATTACTGGAGTTAGTTGAGGCATTAGGTCCCAAAATTTGTTTACTAAAAACACATGTGGATATCTTGACTGATTTTTCCATGGAGGGCACAGTTAAGCCGCTAAAGGCATTATCCGCCAAGTACAATTTTTTACTCTTCGAAGACAGAAAATTTGCTGACATTGGTAATACAGTCAAATTGCAGTACTCTGCGGGTGTATACAGAATAGCAGAATGGGCAGACATTACGAATGCACACGGTGTGGTGGGCCCAGGTATTGTTAGCGGTTTGAAGCAGGCGGCAGAAGAAGTAACAAAGGAACCTAGAGGCCTTTTGATGTTAGCAGAATTGTCATGCAAGGGCTCCCTATCTACTGGAGAATATACTAAGGGTACTGTTGACATTGCGAAGAGCGACAAAGATTTTGTTATCGGCTTTATTGCTCAAAGAGACATGGGTGGAAGAGATGAAGGTTACGATTGGTTGATTATGACACCCGGTGTGGGTTTAGATGACAAGGGAGACGCATTGGGTCAACAGTATAGAACCGTGGATGATGTGGTCTCTACAGGATCTGACATTATTATTGTTGGAAGAGGACTATTTGCAAAGGGAAGGGATGCTAAGGTAGAGGGTGAACGTTACAGAAAAGCAGGCTGGGAAGCATATTTGAGAAGATGCGGCCAGCAAAACTAAAAAACTGTATTATAAGTAAATGCATGTATACTAAACTCACGAATTCGCCAAAAGTTGGCCCAGGGCTTCCCGGTATCAACAGGGACACCAGGATTTATTTATTCTGCGAAGTGATCTTCCGTCACAGGTATTTATTCGCGATAAGCTCATGGAGCGGCGTAACCGTCGCACAGGAAGGACAGAGAAAGCGCGGATCTGGGAAGTGACGGACAGAACGGTCAGGACCTGGATTGGGGAGGCGGTTGCCGCCGCTGCTGCTGACGGTGTGACGTTCTCTGTTCCGGTCACACCACATACGTTCCGCCATTCCTATGCGATGCACATGCTGTATGCCGGTATACCGCTGAAAGTTCTGCAAAGCCTGATGGGACATAAGTCCATCAGTTCAACGGAAGTCTACACGAAGGTTTTTGCGCTGGATGTGGCTGCCCGGCACCGGGTGCAGTTTGCGATGCCGGAGTCTGATGCGGTTGCGATGCTGAAACAATTATCCTGAGAATAAATGCCTTGGCCTTTATATGGAAATGTGGAACTGAGTGGATATGCTGTTTTTGTCTGTTAAACAGAGAAGCTGGCTGTTATCCACTGAGAAGCGAACGAAACAGTCGGGAAAATCTCCCATTATCGTAGAGATCCGCATTATTAATCTCAGGAGCCTGTGTAGCGTTTATAGGAAGTAGTGTTCTGTCATGATGCCTGCAAGCGGTAACGAAAACGATTTGAATATGCCTTCAGGAACAATAGAAATCTTCGTGCGGTGTTACGTTGAAGTGGAGCGGATTATGTCAGCAATGGACAGAACAACCTAATGAACACAGAACCATGATGTGGTCTGTCCTTTTACAGCCAGTAGTGCTCGCCGCAGTCGAGCGACAGGGCGAAGCCCTCGAGTGAGCGAGGAAGCACCAGGGAACAGCACTTATATATTCTGCTTACACACGATGCCTGAAAAAACTTCCCTTGGGGTTATCCACTTATCCACGGGGATATTTTTATAATTATTTTTTTTATAGTTTTTAGATCTTCTTTTTTAGAGCGCCTTGTAGGCCTTTATCCATGCTGGTTCTAGAGAAGGTGTTGTGACAAATTGCCCTTTCAGTGTGACAAATCACCCTCAAATGACAGTCCTGTCTGTGACAAATTGCCCTTAACCCTGTGACAAATTGCCCTCAGAAGAAGCTGTTTTTTCACAAAGTTATCCCTGCTTATTGACTCTTTTTTATTTAGTGTGACAATCTAAAAACTTGTCACACTTCACATGGATCTGTCATGGCGGAAACAGCGGTTATCAATCACAAGAAACGTAAAAATAGCCCGCGAATCGTCCAGTCAAACGACCTCACTGAGGCGGCATATAGTCTCTCCCGGGATCAAAAACGTATGCTGTATCTGTTCGTTGACCAGATCAGAAAATCTGATGGCACCCTACAGGAACATGACGGTATCTGCGAGATCCATGTTGCTAAATATGCTGAAATATTCGGATTGACCTCTGCGGAAGCCAGTAAGGATATACGGCAGGCATTGAAGAGTTTCGCGGGGAAGGAAGTGGTTTTTTATCGCCCTGAAGAGGATGCCGGCGATGAAAAAGGCTATGAATCTTTTCCTTGGTTTATCAAACGTGCGCACAGTCCATCCAGAGGGCTTTACAGTGTACATATCAACCCATATCTCATTCCCTTCTTTATCGGGTTACAGAACCGGTTTACGCAGTTTCGGCTTAGTGAAACAAAAGAAATCACCAATCCGTATGCCATGCGTTTATACGAATCCCTGTGTCAGTATCGTAAGCCGGATGGCTCAGGCATCGTCTCTCTGAAAATCGACTGGATCATAGAGCGTTACCAGCTGCCTCAAAGTTACCAGCGTATGCCTGACTTCCGCCGCCGCTTCCTGCAGGTCTGTGTTAATGAGATCAACAGCAGAACTCCAATGCGCCTCTCATACATTGAGAAAAAGAAAGGCCGCCAGACGACTCATATCGTATTTTCCTTCCGCGATATCACTTCCATGACGACAGGATAGTCTGAGGGTTATCTGTCACAGATTTGAGGGTGGTTCGTCACATTTGTTCTGACCTACTGAGGGTAATTTGTCACAGTTTTGCTGTTTCCTTCAGCCTGCATGGATTTTCTCATACTTTTTGAACTGTAATTTTTAAGGAAGCCAAATTTGAGGGCAGTTTGTCACAGTTGATTTCCTTCTCTTTCCCTTCGTCATGTGACCTGATATCGGGGGTTAGTTCGTCATCATTGATGAGGGTTGATTATCACAGTTTATTACTCTGAATTGGCTATCCGCGTGTGTACCTCTACCTGGAGTTTTTCCCACGGTGGATATTTCTTCTTGCGCTGAGCGTAAGAGCTATCTGACAGAACAGTTCTTCTTTGCTTCCTCGCCAGTTCGCTCGCTATGCTCGGTTACACGGCTGCGGCGAGCGCTAGTGATAATAAGTGACTGAGGTATGTGCTCTTCTTATCTCCTTTTGTAGTGTTGCTCTTATTTTAAACAACTTTGCGGTTTTTTGATGACTTTGCGATTTTGTTGTTGCTTTGCAGTAAATTGCAAGATTTAATAAAAAAACGCAAAGCAATGATTAAAGGATGTTCAGAATGAAACTCATGGAAACACTTAACCAGTGCATAAACGCTGGTCATGAAATGACGAAGGCTATCGCCATTGCACAGTTTAATGATGACAGCCCGGAAGCGAGGAAAATAACCCGGCGCTGGAGAATAGGTGAAGCAGCGGATTTAGTTGGGGTTTCTTCTCAGGCTATCAGAGATGCCGAGAAAGCAGGGCGACTACCGCACCCGGATATGGAAATTCGAGGACGGGTTGAGCAACGTGTTGGTTATACAATTGAACAAATTAATCATATGCGTGATGTGTTTGGTACGCGATTGCGACGTGCTGAAGACGTATTTCCACCGGTGATCGGGGTTGCTGCCCATAAAGGTGGCGTTTACAAAACCTCAGTTTCTGTTCATCTTGCTCAGGATCTGGCTCTGAAGGGGCTACGTGTTTTGCTCGTGGAAGGTAACGACCCCCAGGGAACAGCCTCAATGTATCACGGATGGGTACCAGATCTTCATATTCATGCAGAAGACACTCTCCTGCCTTTCTATCTTGGGGAAAAGGACGATGTCACTTATGCAATAAAGCCCACTTGCTGGCCGGGGCTTGACATTATTCCTTCCTGTCTGGCTCTGCACCGTATTGAAACTGAGTTAATGGGCAAATTTGATGAAGGTAAACTGCCCACCGATCCACACCTGATGCTCCGACTGGCCATTGAAACTGTTGCTCATGACTATGATGTCATAGTTATTGACAGCGCGCCTAACCTGGGTATCGGCACGATTAATGTCGTATGTGCTGCTGATGTGCTGATTGTTCCCACGCCTGCTGAGTTGTTTGACTACACCTCCGCACTGCAGTTTTTCGATATGCTTCGTGATCTGCTCAAGAACGTTGATCTTAAAGGGTTCGAGCCTGATGTACGTATTTTGCTTACCAAATACAGCAATAGTAATGGCTCTCAGTCCCCGTGGATGGAGGAGCAAATTCGGGATGCCTGGGGAAGCATGGTTCTAAAAAATGTTGTACGTGAAACGGATGAAGTTGGTAAAGGTCAGATCCGGATGAGAACTGTTTTTGAACAGGCCATTGATCAACGCTCTTCAACTGGTGCCTGGAGAAATGCTCTTTCTATTTGGGAACCTGTCTGCAATGAAATTTTCGATCGTCTGATTAAACCACGCTGGGAGATTAGATAATGAAGCGTGCGCCTGTTATTCCAAAACATACGCTCAATACTCAACCGGTTGAAGATACTTCGTTATCGACACCAGCTGCCCCGATGGTGGATTCGTTAATTGCGCGCGTAGGAGTAATGGCTCGCGGTAATGCCATTACTTTGCCTGTATGTGGTCGGGATGTGAAGTTTACTCTTGAAGTGCTCCGGGGTGATAGTGTTGAGAAGACCTCTCGGGTATGGTCAGGTAATGAACGTGACCAGGAGCTGCTTACTGAGGACGCACTGGATGATCTCATCCCTTCTTTTCTACTGACTGGTCAACAGACACCGGCGTTCGGTCGAAGAGTATCTGGTGTCATAGAAATTGCCGATGGGAGTCGCCGTCGTAAAGCTGCTGCACTTACCGAAAGTGATTATCGTGTTCTGGTTGGCGAGCTGGATGATGAGCAGATGGCTGCATTATCCAGATTGGGTAACGATTATCGCCCAACAAGTGCTTATGAACGTGGTCAGCGTTATGCAAGCCGATTGCAGAATGAATTTGCTGGAAATATTTCTGCGCTGGCTGATGCGGAAAATATTTCACGTAAGATTATTACCCGCTGTATCAACACCGCCAAATTGCCTAAATCAGTTGTTGCTCTTTTTTCTCACCCCGGTGAACTATCTGCCCGGTCAGGTGATGCACTTCAAAAAGCCTTTACAGATAAAGAGGAATTACTTAAGCAGCAGGCATCTAACCTTCATGAGCAGAAAAAAGCTGGGGTGATATTTGAAGCTGAAGAAGTTATCACTCTTTTAACTTCTGTGCTTAAAACGTCATCTGCATCAAGAACTAGTTTAAGCTCACGACATCAGTTTGCTCCTGGAGCGACAGTATTGTATAAGGGCGATAAAATGGTGCTTAACCTGGACAGGTCTCGTGTTCCAACTGAGTGTATAGAGAAAATTGAGGCCATTCTTAAGGAACTTGAAAAGCCAGCACCCTGATGCGACCACGTTTTAGTCTACGTTTATCTGTCTTTACTTAATGTCCTTTGTTACAGGCCAGAAAGCATAACTGGCCTGAATATTCTCTCTGGGCCCACTGTTCCACTTGTATCGTCGGTCTGATAATCAGACTGGGACCACGGTCCCACTCGTATCGTCGGTCTGATTATTAGTCTGGGACCACGGTCCCACTCGTATCGTCGGTCTGATTATTAGTCTGGGACCACGGTCCCACTCGTATCGTCGGTCTGATAATCAGACTGGGACCACGGTCCCACTCGTATCGTCGGTCTGATTATTAGTCTGGGACCATGGTCCCACTCGTATCGTCGGTCTGATTATTAGTCTGGGACCACGGTCCCACTCGTATCGTCGGTCTGATTATTAGTCTGGAACCACGGTCCCACTCGTATCGTCGGTCTGATTATTAGTCTGGGACCACGGTCCCACTCGTATCGTCGGTCTGATTATTAGTCTGGGACCACGATCCCACTCGTGTTGTCGGTCTGATTATCGGTCTGGGACCACGGTCCCACTTGTATTGTCGATCAGACTATCAGCGTGAGACTACGATTCCATCAATGCCTGTCAAGGGCAAGTATTGACATGTCGTCGTAACCTGTAGAACGGAGTAACCTCGGTGTGCGGTTGTATGCCTGCTGTGGATTGCTGCT

>MAPT_Assembled_Isoform

GTGAACTTTGAACCAGGATGGCTGAGCCCCGCCAGGAGTTCGAAGTGATGGAAGATCACGCTGGGACGTACGGGTTGGGGGACAGGAAAGATCAGGGGGGCTACACCATGCACCAAGACCAAGAGGGTGACACGGACGCTGGCCTGAAAGAATCTCCCCTGCAGACCCCCACTGAGGACGGATCTGAGGAACCGGGCTCTGAAACCTCTGATGCTAAGAGCACTCCAACAGCGGAAGATGTGACAGCACCCTTAGTGGATGAGGGAGCTCCCGGCAAGCAGGCTGCCGCGCAGCCCCACACGGAGATCCCAGAAGGAACCACAGCTGAAGAAGCAGGCATTGGAGACACCCCCAGCCTGGAAGACGAAGCTGCTGGTCACGTGACCCAAGAGGAGTTGAGAGTTCCGGGCCGGCAGAGGAAGGCGCCTGAAAGGCCCCTGGCCAATGAGATTAGCGCCCACGTCCAGCCTGGACCCTGCGGAGAGGCCTCTGGGGTCTCTGGGCCGTGCCTCGGGGAGAAAGAGCCAGAAGCTCCCGTCCCGCTGACCGCGAGCCTTCCTCAGCACCGTCCCGTTTGCCCAGCGCCTCCTCCAACAGGAGGCCCTCAGGAGCCCTCCCTGGAGTGGGGACAAAAAGGCGGGGACTGGGCCGAGAAGGGTCCGGCCTTTCCGAAGCCCGCCACCACTGCGTATCTCCACACAGAGCCTGAAAGTGGTAAGGTGGTCCAGGAAGGCTTCCTCCGAGAGCCAGGCCCCCCAGGTCTGAGCCACCAGCTCATGTCCGGCATGCCTGGGGCTCCCCTCCTGCCTGAGGGCCCCAGAGAGGCCACACGCCAACCTTCGGGGACAGGACCTGAGGACACAGAGGGCGGCCGCCACGCCCCTGAGCTGCTCAAGCACCAGCTTCTAGGAGACCTGCACCAGGAGGGGCCGCCGCTGAAGGGGGCAGGGGGCAAAGAGAGGCCGGGGAGCAAGGAGGAGGTGGATGAAGACCGCGACGTCGATGAGTCCTCCCCCCAAGACTCCCCTCCCTCCAAGGCCTCCCCAGCCCAAGATGGGCGGCCTCCCCAGACAGCCGCCAGAGAAGCCACCAGCATCCCAGGCTTCCCAGCGGAGGGTGCCATCCCCCTCCCTGTGGATTTCCTCTCCAAAGTTTCCACAGAGATCCCAGCCTCAGAGCCCGACGGGCCCAGTGTAGGGCGGGCCAAAGGGCAGGATGCCCCCCTGGAGTTCACGTTTCACGTGGAAATCACACCCAACGTGCAGAAGGAGCAGGCGCACTCGGAGGAGCATTTGGGAAGGGCTGCATTTCCAGGGGCCCCTGGAGAGGGGCCAGAGGCCCGGGGCCCCTCTTTGGGAGAGGACACAAAAGAGGCTGACCTTCCAGAGCCCTCTGAAAAGCAGCCTGCTGCTGCTCCGCGGGGGAAGCCCGTCAGCCGGGTCCCTCAACTCAAAGCTCGCATGGTCAGTAAAAGCAAAGACGGGACTGGAAGCGATGACAAAAAAGCCAAGACATCCACACGTTCCTCTGCTAAAACCTTGAAAAATAGGCCTTGCCTTAGCCCCAAACACCCCACTCCTGGTAGCTCAGACCCTCTGATCCAACCCTCCAGCCCTGCTGTGTGCCCAGAGCCACCTTCCTCTCCTAAATACGTCTCTTCTGTCACTTCCCGAACTGGCAGTTCTGGAGCAAAGGAGATGAAACTCAAGGGGGCTGATGGTAAAACGAAGATCGCCACACCGCGGGGAGCAGCCCCTCCAGGCCAGAAGGGCCAGGCCAACGCCACCAGGATTCCAGCAAAAACCCCGCCCGCTCCAAAGACACCACCCAGCTCTGGTGAACCTCCAAAATCAGGGGATCGCAGCGGCTACAGCAGCCCCGGCTCCCCAGGCACTCCCGGCAGCCGCTCCCGCACCCCGTCCCTTCCAACCCCACCCACCCGGGAGCCCAAGAAGGTGGCAGTGGTCCGTACTCCACCCAAGTCGCCGTCTTCCGCCAAGAGCCGCCTGCAGACAGCCCCCGTGCCCATGCCAGACCTGAAGAATGTCAAGTCCAAGATCGGCTCCACTGAGAACCTGAAGCACCAGCCGGGAGGCGGGAAGGTGCAGATAATTAATAAGAAGCTGGATCTTAGCAACGTCCAGTCCAAGTGTGGCTCAAAGGATAATATCAAACACGTCCCGGGAGGCGGCAGTGTGCAAATAGTCTACAAACCAGTTGACCTGAGCAAGGTGACCTCCAAGTGTGGCTCATTAGGCAACATCCATCATAAACCAGGAGGTGGCCAGGTGGAAGTAAAATCTGAGAAGCTTGACTTCAAGGACAGAGTCCAGTCGAAGATTGGGTCCCTGGACAATATCACCCACGTCCCTGGCGGAGGAAATAAAAAGATTGAAACCCACAAGCTGACCTTCCGCGAGAACGCCAAAGCCAAGACAGACCACGGGGCGGAGATCGTGTACAAGTCGCCAGTGGTGTCTGGGGACACGTCTCCACGGCATCTCAGCAATGTCTCCTCCACCGGCAGCATCGACATGGTAGACTCGCCCCAGCTCGCCACGCTAGCTGACGAGGTGTCTGCCTCCCTGGCCAAGCAGGGTTTGTGA

>MAPT_Exon1

GTGAACTTTGAACCAGGATGGCTGAGCCCCGCCAGGAGTTCGAAGTGATGGAAGATCACGCTGGGACGTACGGGTTGGGGGACAGGAAAGATCAGGGGGGCTACACCATGCACCAAGACCAAGAGGGTGACACGGACGCTGGCCTGAAAG

>MAPT_Exon2

AATCTCCCCTGCAGACCCCCACTGAGGACGGATCTGAGGAACCGGGCTCTGAAACCTCTGATGCTAAGAGCACTCCAACAGCGGAAG

>MAPT_Exon3

ATGTGACAGCACCCTTAGTGGATGAGGGAGCTCCCGGCAAGCAGGCTGCCGCGCAGCCCCACACGGAGATCCCAGAAGGAACCACAG

>MAPT_Exon4

CTGAAGAAGCAGGCATTGGAGACACCCCCAGCCTGGAAGACGAAGCTGCTGGTCACGTGACCCAAG

>MAPT_Exon5

AGGAGTTGAGAGTTCCGGGCCGGCAGAGGAAGGCGCCTGAAAGGCCCCTGGCCAATGAGATTAGCGCCCACGTCCAGCCTGGACCCTGCGGAGAGGCCTCTGGGGTCTCTGGGCCGTGCCTCGGGGAGAAAGAGCCAGAAGCTCCCGTCCCGCTGACCGCGAGCCTTCCTCAGCACCGTCCCGTTTGCCCAGCGCCTCCTCCAACAGGAGGCCCTCAGGAGCCCTCCCTGGAGTGGGGACAAAAAGGCGGGGACTGGGCCGAGAAGGGTCCGGCCTTTCCGAAGCCCGCCACCACTGCGTATCTCCACACAGAG

>MAPT_Exon6

CCTGAAAGTGGTAAGGTGGTCCAGGAAGGCTTCCTCCGAGAGCCAGGCCCCCCAGGTCTGAGCCACCAGCTCATGTCCGGCATGCCTGGGGCTCCCCTCCTGCCTGAGGGCCCCAGAGAGGCCACACGCCAACCTTCGGGGACAGGACCTGAGGACACAGAGGGCGGCCGCCACGCCCCTGAGCTGCTCAAGCACCAGCTTCTAGGAGACCTGCACCAGGAGGGGCCGCCGCTGAAGGGGGCAGGGGGCAAAGAGAGGCCGGGGAGCAAGGAGGAGGTGGATGAAGACCGCGACGTCGATGAGTCCTCCCCCCAAGACTCCCCTCCCTCCAAGGCCTCCCCAGCCCAAGATGGGCGGCCTCCCCAGACAGCCGCCAGAGAAGCCACCAGCATCCCAGGCTTCCCAGCGGAGGGTGCCATCCCCCTCCCTGTGGATTTCCTCTCCAAAGTTTCCACAGAGATCCCAGCCTCAGAGCCCGACGGGCCCAGTGTAGGGCGGGCCAAAGGGCAGGATGCCCCCCTGGAGTTCACGTTTCACGTGGAAATCACACCCAACGTGCAGAAGGAGCAGGCGCACTCGGAGGAGCATTTGGGAAGGGCTGCATTTCCAGGGGCCCCTGGAGAGGGGCCAGAGGCCCGGGGCCCCTCTTTGGGAGAGGACACAAAAGAGGCTGACCTTCCAGAGCCCTCTGAAAAGCAGCCTGCTGCTGCTCCGCGGGGGAAGCCCGTCAGCCGGGTCCCTCAACTCAAAG

>MAPT_Exon7

CTCGCATGGTCAGTAAAAGCAAAGACGGGACTGGAAGCGATGACAAAAAAGCCAAG

>MAPT_Exon8

ACATCCACACGTTCCTCTGCTAAAACCTTGAAAAATAGGCCTTGCCTTAGCCCCAAACACCCCACTCCTGGTAGCTCAGACCCTCTGATCCAACCCTCCAGCCCTGCTGTGTGCCCAGAGCCACCTTCCTCTCCTAAATACGTCTCTTCTGTCACTTCCCGAACTGGCAGTTCTGGAGCAAAGGAGATGAAACTCAAG

>MAPT_Exon9

GGGGCTGATGGTAAAACGAAGATCGCCACACCGCGGGGAGCAGCCCCTCCAGGCCAGAAGGGCCAGGCCAACGCCACCAGGATTCCAGCAAAAACCCCGCCCGCTCCAAAGACACCACCCAGCTCTG

>MAPT_Exon10

GTGAACCTCCAAAATCAGGGGATCGCAGCGGCTACAGCAGCCCCGGCTCCCCAGGCACTCCCGGCAGCCGCTCCCGCACCCCGTCCCTTCCAACCCCACCCACCCGGGAGCCCAAGAAGGTGGCAGTGGTCCGTACTCCACCCAAGTCGCCGTCTTCCGCCAAGAGCCGCCTGCAGACAGCCCCCGTGCCCATGCCAGACCTGAAGAATGTCAAGTCCAAGATCGGCTCCACTGAGAACCTGAAGCACCAGCCGGGAGGCGGGAAG

>MAPT_Exon11

GTGCAGATAATTAATAAGAAGCTGGATCTTAGCAACGTCCAGTCCAAGTGTGGCTCAAAGGATAATATCAAACACGTCCCGGGAGGCGGCAGT

>MAPT_Exon12

GTGCAAATAGTCTACAAACCAGTTGACCTGAGCAAGGTGACCTCCAAGTGTGGCTCATTAGGCAACATCCATCATAAACCAG

>MAPT_Exon13

GAGGTGGCCAGGTGGAAGTAAAATCTGAGAAGCTTGACTTCAAGGACAGAGTCCAGTCGAAGATTGGGTCCCTGGACAATATCACCCACGTCCCTGGCGGAGGAAATAAAAAG

>MAPT_Exon14

ATTGAAACCCACAAGCTGACCTTCCGCGAGAACGCCAAAGCCAAGACAGACCACGGGGCGGAGATCGTGTACAAGTCGCCAGTGGTGTCTGGGGACACGTCTCCACGGCATCTCAGCAATGTCTCCTCCACCGGCAGCATCGACATGGTAGACTCGCCCCAGCTCGCCACGCTAGCTGACGAGGTGTCTGCCTCCCTGGCCAAGCAGGGTTTGTGA

>VEGFA_Assembled_Isoform

CTGACGGACAGACAGACAGACACCGCCCCCAGCCCCAGCTACCACCTCCTCCCCGGCCGGCGGCGGACAGTGGACGCGGCGGCGAGCCGCGGGCAGGGGCCGGAGCCCGCGCCCGGAGGCGGGGTGGAGGGGGTCGGGGCTCGCGGCGTCGCACTGAAACTTTTCGTCCAACTTCTGGGCTGTTCTCGCTTCGGAGGAGCCGTGGTCCGCGCGGGGGAAGCCGAGCCGAGCGGAGCCGCGAGAAGTGCTAGCTCGGGCCGGGAGGAGCCGCAGCCGGAGGAGGGGGAGGAGGAAGAAGAGAAGGAAGAGGAGAGGGGGCCGCAGTGGCGACTCGGCGCTCGGAAGCCGGGCTCATGGACGGGTGAGGCGGCGGTGTGCGCAGACAGTGCTCCAGCCGCGCGCGCTCCCCAGGCCCTGGCCCGGGCCTCGGGCCGGGGAGGAAGAGTAGCTCGCCGAGGCGCCGAGGAGAGCGGGCCGCCCCACAGCCCGAGCCGGAGAGGGAGCGCGAGCCGCGCCGGCCCCGGTCGGGCCTCCGAAACCATGAACTTTCTGCTGTCTTGGGTGCATTGGAGCCTTGCCTTGCTGCTCTACCTCCACCATGCCAAGTGGTCCCAGGCTGCACCCATGGCAGAAGGAGGAGGGCAGAATCATCACGAAGTGGTGAAGTTCATGGATGTCTATCAGCGCAGCTACTGCCATCCAATCGAGACCCTGGTGGACATCTTCCAGGAGTACCCTGATGAGATCGAGTACATCTTCAAGCCATCCTGTGTGCCCCTGATGCGATGCGGGGGCTGCTGCAATGACGAGGGCCTGGAGTGTGTGCCCACTGAGGAGTCCAACATCACCATGCAGATTATGCGGATCAAACCTCACCAAGGCCAGCACATAGGAGAGATGAGCTTCCTACAGCACAACAAATGTGAATGCAGACCAAAGAAAGATAGAGCAAGACAAGAAAAAAAATCAGTTCGAGGAAAGGGAAAGGGGCAAAAACGAAAGCGCAAGAAATCCCGGTATAAGTCCTGGAGCGTGTACGTTGGTGCCCGCTGCTGTCTAATGCCCTGGAGCCTCCCTGGCCCCCATCCCTGTGGGCCTTGCTCAGAGCGGAGAAAGCATTTGTTTGTACAAGATCCGCAGACGTGTAAATGTTCCTGCAAAAACACAGACTCGCGTTGCAAGGCGAGGCAGCTTGAGTTAAACGAACGTACTTGCAGATGTGACAAGCCGAGGCGGTGA

>VEGFA_Exon1

CTGACGGACAGACAGACAGACACCGCCCCCAGCCCCAGCTACCACCTCCTCCCCGGCCGGCGGCGGACAGTGGACGCGGCGGCGAGCCGCGGGCAGGGGCCGGAGCCCGCGCCCGGAGGCGGGGTGGAGGGGGTCGGGGCTCGCGGCGTCGCACTGAAACTTTTCGTCCAACTTCTGGGCTGTTCTCGCTTCGGAGGAGCCGTGGTCCGCGCGGGGGAAGCCGAGCCGAGCGGAGCCGCGAGAAGTGCTAGCTCGGGCCGGGAGGAGCCGCAGCCGGAGGAGGGGGAGGAGGAAGAAGAGAAGGAAGAGGAGAGGGGGCCGCAGTGGCGACTCGGCGCTCGGAAGCCGGGCTCATGGACGGGTGAGGCGGCGGTGTGCGCAGACAGTGCTCCAGCCGCGCGCGCTCCCCAGGCCCTGGCCCGGGCCTCGGGCCGGGGAGGAAGAGTAGCTCGCCGAGGCGCCGAGGAGAGCGGGCCGCCCCACAGCCCGAGCCGGAGAGGGAGCGCGAGCCGCGCCGGCCCCGGTCGGGCCTCCGAAACCATGAACTTTCTGCTGTCTTGGGTGCATTGGAGCCTTGCCTTGCTGCTCTACCTCCACCATGCCAAG

>VEGFA_Exon2

TGGTCCCAGGCTGCACCCATGGCAGAAGGAGGAGGGCAGAATCATCACGAAG

>VEGFA_Exon3

TGGTGAAGTTCATGGATGTCTATCAGCGCAGCTACTGCCATCCAATCGAGACCCTGGTGGACATCTTCCAGGAGTACCCTGATGAGATCGAGTACATCTTCAAGCCATCCTGTGTGCCCCTGATGCGATGCGGGGGCTGCTGCAATGACGAGGGCCTGGAGTGTGTGCCCACTGAGGAGTCCAACATCACCATGCAG

>VEGFA_Exon4

ATTATGCGGATCAAACCTCACCAAGGCCAGCACATAGGAGAGATGAGCTTCCTACAGCACAACAAATGTGAATGCAG

>VEGFA_Exon5

ACCAAAGAAAGATAGAGCAAGACAAGAAAA

>VEGFA_Exon6

AAAATCAGTTCGAGGAAAGGGAAAGGGGCAAAAACGAAAGCGCAAGAAATCCCGGTATAAGTCCTGGAGCGT

>VEGFA_Exon7

GTACGTTGGTGCCCGCTGCTGTCTAATGCCCTGGAGCCTCCCTGGCCCCCA

>VEGFA_Exon8

TCCCTGTGGGCCTTGCTCAGAGCGGAGAAAGCATTTGTTTGTACAAGATCCGCAGACGTGTAAATGTTCCTGCAAAAACACAGACTCGCGTTGCAAGGCGAGGCAGCTTGAGTTAAACGAACGTACTTGCAG

>VEGFA_Exon9

ATGTGACAAGCCGAGGCGGTGA

>BRCA1_Assembled_Isoform

ATGGATTTATCTGCTCTTCGCGTTGAAGAAGTACAAAATGTCATTAATGCTATGCAGAAAATCTTAGAGTGTCCCATCTGTCTGGAGTTGATCAAGGAACCTGTCTCCACAAAGTGTGACCACATATTTTGCAAATTTTGCATGCTGAAACTTCTCAACCAGAAGAAAGGGCCTTCACAGTGTCCTTTATGTAAGAATGATATAACCAAAAGGAGCCTACAAGAAAGTACGAGATTTAGTCAACTTGTTGAAGAGCTATTGAAAATCATTTGTGCTTTTCAGCTTGACACAGGTTTGGAGTATGCAAACAGCTATAATTTTGCAAAAAAGGAAAATAACTCTCCTGAACATCTAAAAGATGAAGTTTCTATCATCCAAAGTATGGGCTACAGAAACCGTGCCAAAAGACTTCTACAGAGTGAACCCGAAAATCCTTCCTTGCAGGAAACCAGTCTCAGTGTCCAACTCTCTAACCTTGGAACTGTGAGAACTCTGAGGACAAAGCAGCGGATACAACCTCAAAAGACGTCTGTCTACATTGAATTGGGATCTGATTCTTCTGAAGATACCGTTAATAAGGCAACTTATTGCAGTGTGGGAGATCAAGAATTGTTACAAATCACCCCTCAAGGAACCAGGGATGAAATCAGTTTGGATTCTGCAAAAAAGGCTGCTTGTGAATTTTCTGAGACGGATGTAACAAATACTGAACATCATCAACCCAGTAATAATGATTTGAACACCACTGAGAAGCGTGCAGCTGAGAGGCATCCAGAAAAGTATCAGGGTAGTTCTGTTTCAAACTTGCATGTGGAGCCATGTGGCACAAATACTCATGCCAGCTCATTACAGCATGAGAACAGCAGTTTATTACTCACTAAAGACAGAATGAATGTAGAAAAGGCTGAATTCTGTAATAAAAGCAAACAGCCTGGCTTAGCAAGGAGCCAACATAACAGATGGGCTGGAAGTAAGGAAACATGTAATGATAGGCGGACTCCCAGCACAGAAAAAAAGGTAGATCTGAATGCTGATCCCCTGTGTGAGAGAAAAGAATGGAATAAGCAGAAACTGCCATGCTCAGAGAATCCTAGAGATACTGAAGATGTTCCTTGGATAACACTAAATAGCAGCATTCAGAAAGTTAATGAGTGGTTTTCCAGAAGTGATGAACTGTTAGGTTCTGATGACTCACATGATGGGGAGTCTGAATCAAATGCCAAAGTAGCTGATGTATTGGACGTTCTAAATGAGGTAGATGAATATTCTGGTTCTTCAGAGAAAATAGACTTACTGGCCAGTGATCCTCATGAGGCTTTAATATGTAAAAGTGAAAGAGTTCACTCCAAATCAGTAGAGAGTAATATTGAAGACAAAATATTTGGGAAAACCTATCGGAAGAAGGCAAGCCTCCCCAACTTAAGCCATGTAACTGAAAATCTAATTATAGGAGCATTTGTTACTGAGCCACAGATAATACAAGAGCGTCCCCTCACAAATAAATTAAAGCGTAAAAGGAGACCTACATCAGGCCTTCATCCTGAGGATTTTATCAAGAAAGCAGATTTGGCAGTTCAAAAGACTCCTGAAATGATAAATCAGGGAACTAACCAAACGGAGCAGAATGGTCAAGTGATGAATATTACTAATAGTGGTCATGAGAATAAAACAAAAGGTGATTCTATTCAGAATGAGAAAAATCCTAACCCAATAGAATCACTCGAAAAAGAATCTGCTTTCAAAACGAAAGCTGAACCTATAAGCAGCAGTATAAGCAATATGGAACTCGAATTAAATATCCACAATTCAAAAGCACCTAAAAAGAATAGGCTGAGGAGGAAGTCTTCTACCAGGCATATTCATGCGCTTGAACTAGTAGTCAGTAGAAATCTAAGCCCACCTAATTGTACTGAATTGCAAATTGATAGTTGTTCTAGCAGTGAAGAGATAAAGAAAAAAAAGTACAACCAAATGCCAGTCAGGCACAGCAGAAACCTACAACTCATGGAAGGTAAAGAACCTGCAACTGGAGCCAAGAAGAGTAACAAGCCAAATGAACAGACAAGTAAAAGACATGACAGCGATACTTTCCCAGAGCTGAAGTTAACAAATGCACCTGGTTCTTTTACTAAGTGTTCAAATACCAGTGAACTTAAAGAATTTGTCAATCCTAGCCTTCCAAGAGAAGAAAAAGAAGAGAAACTAGAAACAGTTAAAGTGTCTAATAATGCTGAAGACCCCAAAGATCTCATGTTAAGTGGAGAAAGGGTTTTGCAAACTGAAAGATCTGTAGAGAGTAGCAGTATTTCATTGGTACCTGGTACTGATTATGGCACTCAGGAAAGTATCTCGTTACTGGAAGTTAGCACTCTAGGGAAGGCAAAAACAGAACCAAATAAATGTGTGAGTCAGTGTGCAGCATTTGAAAACCCCAAGGGACTAATTCATGGTTGTTCCAAAGATAATAGAAATGACACAGAAGGCTTTAAGTATCCATTGGGACATGAAGTTAACCACAGTCGGGAAACAAGCATAGAAATGGAAGAAAGTGAACTTGATGCTCAGTATTTGCAGAATACATTCAAGGTTTCAAAGCGCCAGTCATTTGCTCCGTTTTCAAATCCAGGAAATGCAGAAGAGGAATGTGCAACATTCTCTGCCCACTCTGGGTCCTTAAAGAAACAAAGTCCAAAAGTCACTTTTGAATGTGAACAAAAGGAAGAAAATCAAGGAAAGAATGAGTCTAATATCAAGCCTGTACAGACAGTTAATATCACTGCAGGCTTTCCTGTGGTTGGTCAGAAAGATAAGCCAGTTGATAATGCCAAATGTAGTATCAAAGGAGGCTCTAGGTTTTGTCTATCATCTCAGTTCAGAGGCAACGAAACTGGACTCATTACTCCAAATAAACATGGACTTTTACAAAACCCATATCGTATACCACCACTTTTTCCCATCAAGTCATTTGTTAAAACTAAATGTAAGAAAAATCTGCTAGAGGAAAACTTTGAGGAACATTCAATGTCACCTGAAAGAGAAATGGGAAATGAGAACATTCCAAGTACAGTGAGCACAATTAGCCGTAATAACATTAGAGAAAATGTTTTTAAAGAAGCCAGCTCAAGCAATATTAATGAAGTAGGTTCCAGTACTAATGAAGTGGGCTCCAGTATTAATGAAATAGGTTCCAGTGATGAAAACATTCAAGCAGAACTAGGTAGAAACAGAGGGCCAAAATTGAATGCTATGCTTAGATTAGGGGTTTTGCAACCTGAGGTCTATAAACAAAGTCTTCCTGGAAGTAATTGTAAGCATCCTGAAATAAAAAAGCAAGAATATGAAGAAGTAGTTCAGACTGTTAATACAGATTTCTCTCCATATCTGATTTCAGATAACTTAGAACAGCCTATGGGAAGTAGTCATGCATCTCAGGTTTGTTCTGAGACACCTGATGACCTGTTAGATGATGGTGAAATAAAGGAAGATACTAGTTTTGCTGAAAATGACATTAAGGAAAGTTCTGCTGTTTTTAGCAAAAGCGTCCAGAAAGGAGAGCTTAGCAGGAGTCCTAGCCCTTTCACCCATACACATTTGGCTCAGGGTTACCGAAGAGGGGCCAAGAAATTAGAGTCCTCAGAAGAGAACTTATCTAGTGAGGATGAAGAGCTTCCCTGCTTCCAACACTTGTTATTTGGTAAAGTAAACAATATACCTTCTCAGTCTACTAGGCATAGCACCGTTGCTACCGAGTGTCTGTCTAAGAACACAGAGGAGAATTTATTATCATTGAAGAATAGCTTAAATGACTGCAGTAACCAGGTAATATTGGCAAAGGCATCTCAGGAACATCACCTTAGTGAGGAAACAAAATGTTCTGCTAGCTTGTTTTCTTCACAGTGCAGTGAATTGGAAGACTTGACTGCAAATACAAACACCCAGGATCCTTTCTTGATTGGTTCTTCCAAACAAATGAGGCATCAGTCTGAAAGCCAGGGAGTTGGTCTGAGTGACAAGGAATTGGTTTCAGATGATGAAGAAAGAGGAACGGGCTTGGAAGAAAATAATCAAGAAGAGCAAAGCATGGATTCAAACTTAGGTGAAGCAGCATCTGGGTGTGAGAGTGAAACAAGCGTCTCTGAAGACTGCTCAGGGCTATCCTCTCAGAGTGACATTTTAACCACTCAGCAGAGGGATACCATGCAACATAACCTGATAAAGCTCCAGCAGGAAATGGCTGAACTAGAAGCTGTGTTAGAACAGCATGGGAGCCAGCCTTCTAACAGCTACCCTTCCATCATAAGTGACTCTTCTGCCCTTGAGGACCTGCGAAATCCAGAACAAAGCACATCAGAAAAAGATTCGCATATACATGGCCAAAGGAACAACTCCATGTTTTCTAAAAGGCCTAGAGAACATATATCAGCAGTATTAACTTCACAGAAAAGTAGTGAATACCCTATAAGCCAGAATCCAGAAGGCCTTTCTGCTGACAAGTTTGAGGTGTCTGCAGATAGTTCTACCAGTAAAAATAAAGAACCAGGAGTGGAAAGGTCATCCCCTTCTAAATGCCCATCATTAGATGATAGGTGGTACATGCACAGTTGCTCTGGGAGTCTTCAGAATAGAAACTACCCATCTCAAGAGGAGCTCATTAAGGTTGTTGATGTGGAGGAGCAACAGCTGGAAGAGTCTGGGCCACACGATTTGACGGAAACATCTTACTTGCCAAGGCAAGATCTAGAGGGAACCCCTTACCTGGAATCTGGAATCAGCCTCTTCTCTGATGACCCTGAATCTGATCCTTCTGAAGACAGAGCCCCAGAGTCAGCTCGTGTTGGCAACATACCATCTTCAACCTCTGCATTGAAAGTTCCCCAATTGAAAGTTGCAGAATCTGCCCAGAGTCCAGCTGCTGCTCATACTACTGATACTGCTGGGTATAATGCAATGGAAGAAAGTGTGAGCAGGGAGAAGCCAGAATTGACAGCTTCAACAGAAAGGGTCAACAAAAGAATGTCCATGGTGGTGTCTGGCCTGACCCCAGAAGAATTTATGCTCGTGTACAAGTTTGCCAGAAAACACCACATCACTTTAACTAATCTAATTACTGAAGAGACTACTCATGTTGTTATGAAAACAGATGCTGAGTTTGTGTGTGAACGGACACTGAAATATTTTCTAGGAATTGCGGGAGGAAAATGGGTAGTTAGCTATTTCTGGGTGACCCAGTCTATTAAAGAAAGAAAAATGCTGAATGAGCATGATTTTGAAGTCAGAGGAGATGTGGTCAATGGAAGAAACCACCAAGGTCCAAAGCGAGCAAGAGAATCCCAGGACAGAAAGATCTTCAGGGGGCTAGAAATCTGTTGCTATGGGCCCTTCACCAACATGCCCACAGATCAACTGGAATGGATGGTACAGCTGTGTGGTGCTTCTGTGGTGAAGGAGCTTTCATCATTCACCCTTGGCACAGGTGTCCACCCAATTGTGGTTGTGCAGCCAGATGCCTGGACAGAGGACAATGGCTTCCATGCAATTGGGCAGATGTGTGAGGCACCTGTGGTGACCCGAGAGTGGGTGTTGGACAGTGTAGCACTCTACCAGTGCCAGGAGCTGGACACCTACCTGATACCCCAGATCCCCCACAGCCACTACTGA

>BRCA1_Exon1

ATGGATTTATCTGCTCTTCGCGTTGAAGAAGTACAAAATGTCATTAATGCTATGCAGAAAATCTTAGAGTGTCCCATCTG

>BRCA1_Exon2

TCTGGAGTTGATCAAGGAACCTGTCTCCACAAAGTGTGACCACATATTTTGCAA

>BRCA1_Exon3

ATTTTGCATGCTGAAACTTCTCAACCAGAAGAAAGGGCCTTCACAGTGTCCTTTAT

>BRCA1_Exon4

GTAAGAATGATATAACCAAAAG

>BRCA1_Exon5

GAGCCTACAAGAAAGTACGAGATTTAGTCAACTTGTTGAAGAGCTATTGAAAATCATTTGTGCTTTTCAGCTTGACACAGGTTTGGAGT

>BRCA1_Exon6

ATGCAAACAGCTATAATTTTGCAAAAAAGGAAAATAACTCTCCTGAACATCTAAAAGATGAAGTTTCTATCATCCAAAGTATGGGCTACAGAAACCGTGCCAAAAGACTTCTACAGAGTGAACCCGAAAATCCTTCCTTG

>BRCA1_Exon7

CAGGAAACCAGTCTCAGTGTCCAACTCTCTAACCTTGGAACTGTGAGAACTCTGAGGACAAAGCAGCGGATACAACCTCAAAAGACGTCTGTCTACATTGAATTGG

>BRCA1_Exon8

GATCTGATTCTTCTGAAGATACCGTTAATAAGGCAACTTATTGCAGG

>BRCA1_Exon9

TGTGGGAGATCAAGAATTGTTACAAATCACCCCTCAAGGAACCAGGGATGAAATCAGTTTGGATTCTGCAAAAAAG

>BRCA1_Exon10

GCTGCTTGTGAATTTTCTGAGACGGATGTAACAAATACTGAACATCATCAACCCAGTAATAATGATTTGAACACCACTGAGAAGCGTGCAGCTGAGAGGCATCCAGAAAAGTATCAGG

>BRCA1_Exon11

GTAGTTCTGTTTCAAACTTGCATGTGGAGCCATGTGGCACAAATACTCATGCCAGCTCATTACAGCATGAGAACAGCAGTTTATTACTCACTAAAGACAGAATGAATGTAGAAAAGGCTGAATTCTGTAATAAAAGCAAACAGCCTGGCTTAGCAAGGAGCCAACATAACAGATGGGCTGGAAGTAAGGAAACATGTAATGATAGGCGGACTCCCAGCACAGAAAAAAAGGTAGATCTGAATGCTGATCCCCTGTGTGAGAGAAAAGAATGGAATAAGCAGAAACTGCCATGCTCAGAGAATCCTAGAGATACTGAAGATGTTCCTTGGATAACACTAAATAGCAGCATTCAGAAAGTTAATGAGTGGTTTTCCAGAAGTGATGAACTGTTAGGTTCTGATGACTCACATGATGGGGAGTCTGAATCAAATGCCAAAGTAGCTGATGTATTGGACGTTCTAAATGAGGTAGATGAATATTCTGGTTCTTCAGAGAAAATAGACTTACTGGCCAGTGATCCTCATGAGGCTTTAATATGTAAAAGTGAAAGAGTTCACTCCAAATCAGTAGAGAGTAATATTGAAGACAAAATATTTGGGAAAACCTATCGGAAGAAGGCAAGCCTCCCCAACTTAAGCCATGTAACTGAAAATCTAATTATAGGAGCATTTGTTACTGAGCCACAGATAATACAAGAGCGTCCCCTCACAAATAAATTAAAGCGTAAAAGGAGACCTACATCAGGCCTTCATCCTGAGGATTTTATCAAGAAAGCAGATTTGGCAGTTCAAAAGACTCCTGAAATGATAAATCAGGGAACTAACCAAACGGAGCAGAATGGTCAAGTGATGAATATTACTAATAGTGGTCATGAGAATAAAACAAAAGGTGATTCTATTCAGAATGAGAAAAATCCTAACCCAATAGAATCACTCGAAAAAGAATCTGCTTTCAAAACGAAAGCTGAACCTATAAGCAGCAGTATAAGCAATATGGAACTCGAATTAAATATCCACAATTCAAAAGCACCTAAAAAGAATAGGCTGAGGAGGAAGTCTTCTACCAGGCATATTCATGCGCTTGAACTAGTAGTCAGTAGAAATCTAAGCCCACCTAATTGTACTGAATTGCAAATTGATAGTTGTTCTAGCAGTGAAGAGATAAAGAAAAAAAAGTACAACCAAATGCCAGTCAGGCACAGCAGAAACCTACAACTCATGGAAGGTAAAGAACCTGCAACTGGAGCCAAGAAGAGTAACAAGCCAAATGAACAGACAAGTAAAAGACATGACAGCGATACTTTCCCAGAGCTGAAGTTAACAAATGCACCTGGTTCTTTTACTAAGTGTTCAAATACCAGTGAACTTAAAGAATTTGTCAATCCTAGCCTTCCAAGAGAAGAAAAAGAAGAGAAACTAGAAACAGTTAAAGTGTCTAATAATGCTGAAGACCCCAAAGATCTCATGTTAAGTGGAGAAAGGGTTTTGCAAACTGAAAGATCTGTAGAGAGTAGCAGTATTTCATTGGTACCTGGTACTGATTATGGCACTCAGGAAAGTATCTCGTTACTGGAAGTTAGCACTCTAGGGAAGGCAAAAACAGAACCAAATAAATGTGTGAGTCAGTGTGCAGCATTTGAAAACCCCAAGGGACTAATTCATGGTTGTTCCAAAGATAATAGAAATGACACAGAAGGCTTTAAGTATCCATTGGGACATGAAGTTAACCACAGTCGGGAAACAAGCATAGAAATGGAAGAAAGTGAACTTGATGCTCAGTATTTGCAGAATACATTCAAGGTTTCAAAGCGCCAGTCATTTGCTCCGTTTTCAAATCCAGGAAATGCAGAAGAGGAATGTGCAACATTCTCTGCCCACTCTGGGTCCTTAAAGAAACAAAGTCCAAAAGTCACTTTTGAATGTGAACAAAAGGAAGAAAATCAAGGAAAGAATGAGTCTAATATCAAGCCTGTACAGACAGTTAATATCACTGCAGGCTTTCCTGTGGTTGGTCAGAAAGATAAGCCAGTTGATAATGCCAAATGTAGTATCAAAGGAGGCTCTAGGTTTTGTCTATCATCTCAGTTCAGAGGCAACGAAACTGGACTCATTACTCCAAATAAACATGGACTTTTACAAAACCCATATCGTATACCACCACTTTTTCCCATCAAGTCATTTGTTAAAACTAAATGTAAGAAAAATCTGCTAGAGGAAAACTTTGAGGAACATTCAATGTCACCTGAAAGAGAAATGGGAAATGAGAACATTCCAAGTACAGTGAGCACAATTAGCCGTAATAACATTAGAGAAAATGTTTTTAAAGAAGCCAGCTCAAGCAATATTAATGAAGTAGGTTCCAGTACTAATGAAGTGGGCTCCAGTATTAATGAAATAGGTTCCAGTGATGAAAACATTCAAGCAGAACTAGGTAGAAACAGAGGGCCAAAATTGAATGCTATGCTTAGATTAGGGGTTTTGCAACCTGAGGTCTATAAACAAAGTCTTCCTGGAAGTAATTGTAAGCATCCTGAAATAAAAAAGCAAGAATATGAAGAAGTAGTTCAGACTGTTAATACAGATTTCTCTCCATATCTGATTTCAGATAACTTAGAACAGCCTATGGGAAGTAGTCATGCATCTCAGGTTTGTTCTGAGACACCTGATGACCTGTTAGATGATGGTGAAATAAAGGAAGATACTAGTTTTGCTGAAAATGACATTAAGGAAAGTTCTGCTGTTTTTAGCAAAAGCGTCCAGAAAGGAGAGCTTAGCAGGAGTCCTAGCCCTTTCACCCATACACATTTGGCTCAGGGTTACCGAAGAGGGGCCAAGAAATTAGAGTCCTCAGAAGAGAACTTATCTAGTGAGGATGAAGAGCTTCCCTGCTTCCAACACTTGTTATTTGGTAAAGTAAACAATATACCTTCTCAGTCTACTAGGCATAGCACCGTTGCTACCGAGTGTCTGTCTAAGAACACAGAGGAGAATTTATTATCATTGAAGAATAGCTTAAATGACTGCAGTAACCAGGTAATATTGGCAAAGGCATCTCAGGAACATCACCTTAGTGAGGAAACAAAATGTTCTGCTAGCTTGTTTTCTTCACAGTGCAGTGAATTGGAAGACTTGACTGCAAATACAAACACCCAGGATCCTTTCTTGATTGGTTCTTCCAAACAAATGAGGCATCAGTCTGAAAGCCAGGGAGTTGGTCTGAGTGACAAGGAATTGGTTTCAGATGATGAAGAAAGAGGAACGGGCTTGGAAGAAAATAATCAAGAAGAGCAAAGCATGGATTCAAACTTAG

>BRCA1_Exon12

GTGAAGCAGCATCTGGGTGTGAGAGTGAAACAAGCGTCTCTGAAGACTGCTCAGGGCTATCCTCTCAGAGTGACATTTTAACCACTCAG

>BRCA1_Exon13

CAGAGGGATACCATGCAACATAACCTGATAAAGCTCCAGCAGGAAATGGCTGAACTAGAAGCTGTGTTAGAACAGCATGGGAGCCAGCCTTCTAACAGCTACCCTTCCATCATAAGTGACTCTTCTGCCCTTGAGGACCTGCGAAATCCAGAACAAAGCACATCAGAAAAAG

>BRCA1_Exon14

ATTCGCATATACATGGCCAAAGGAACAACTCCATGTTTTCTAAAAGGCCTAGAGAACATATATCAG

>BRCA1_Exon15

CAGTATTAACTTCACAGAAAAGTAGTGAATACCCTATAAGCCAGAATCCAGAAGGCCTTTCTGCTGACAAGTTTGAGGTGTCTGCAGATAGTTCTACCAGTAAAAATAAAGAACCAGGAGTGGAAAG

>BRCA1_Exon16

GTCATCCCCTTCTAAATGCCCATCATTAGATGATAGGTGGTACATGCACAGTTGCTCTGGGAGTCTTCAGAATAGAAACTACCCATCTCAAGAGGAGCTCATTAAGGTTGTTGATGTGGAGGAGCAACAGCTGGAAGAGTCTGGGCCACACGATTTGACGGAAACATCTTACTTGCCAAGGCAAGATCTAG

>BRCA1_Exon17

AGGGAACCCCTTACCTGGAATCTGGAATCAGCCTCTTCTCTGATGACCCTGAATCTGATCCTTCTGAAGACAGAGCCCCAGAGTCAGCTCGTGTTGGCAACATACCATCTTCAACCTCTGCATTGAAAGTTCCCCAATTGAAAGTTGCAGAATCTGCCCAGAGTCCAGCTGCTGCTCATACTACTGATACTGCTGGGTATAATGCAATGGAAGAAAGTGTGAGCAGGGAGAAGCCAGAATTGACAGCTTCAACAGAAAGGGTCAACAAAAGAATGTCCATGGTGGTGTCTGGCCTGACCCCAGAAGAATTT

>BRCA1_Exon18

ATGCTCGTGTACAAGTTTGCCAGAAAACACCACATCACTTTAACTAATCTAATTACTGAAGAGACTACTCATGTTGTTATGAAAACAG

>BRCA1_Exon19

ATGCTGAGTTTGTGTGTGAACGGACACTGAAATATTTTCTAGGAATTGCGGGAGGAAAATGGGTAGTTAGCTATTTCT

>BRCA1_Exon20

GGGTGACCCAGTCTATTAAAGAAAGAAAAATGCTGAATGAG

>BRCA1_Exon21

CATGATTTTGAAGTCAGAGGAGATGTGGTCAATGGAAGAAACCACCAAGGTCCAAAGCGAGCAAGAGAATCCCAGGACAGAAAG

>BRCA1_Exon22

ATCTTCAGGGGGCTAGAAATCTGTTGCTATGGGCCCTTCACCAACATGCCCACAG

>BRCA1_Exon23

ATCAACTGGAATGGATGGTACAGCTGTGTGGTGCTTCTGTGGTGAAGGAGCTTTCATCATTCACCCTTGGCACA

>BRCA1_Exon24

GGTGTCCACCCAATTGTGGTTGTGCAGCCAGATGCCTGGACAGAGGACAATGGCTTCCATG

>BRCA1_Exon25

CAATTGGGCAGATGTGTGAGGCACCTGTGGTGACCCGAGAGTGGGTGTTGGACAGTGTAGCACTCTACCAGTGCCAGGAGCTGGACACCTACCTGATACCCCAGATCCCCCACAGCCACTACTGA

>Northern_White_Rhino_Sonic_Hedgehog_gDNA

TCTCTGAAAAACTATTTAAATAAGACCTGGGCAGTTTGCCCCTCAAAAATGTGAGACAATATTTGTTTGATGGTCAAAGCTGGGGGCGGGGGTCTAGACTGTCCTTTAAAAACTTAAGTTGGTTTTAACTGAAGCACCTACCATTCTGGAGAAAGTGAAGGGGGAACGCGGAGATGTGCACAAAGCAAGAGTAATTGATGTTATCCTGAGGAACGTGCCTCCGCCCTCCCGCCCACCTCTGTCTCAGTAAAGATTTTTTTTTTTTTAACTCGAAGCCTTCAGGTGAAAATAAAGCAACAGCAGCAACAGAAAAAAAAAAATTTAGTCTTTTTTAGGGTTAAACATCAGAAGACAAGCTTGTGCTGCCTTGGCCAATCAGATACGCCCCTGCCAGCTATAATATAGTACTGACGGCAGCCTCTCTCACAAGCTCTCGAGGCTTGCTACCATTTAAAATCAGACTCTTTTTGTCTTTTGATTGCTGTCTCGCGCCCCAACTCCGATGTGTTCCGTTATCAGCGACCAGCAGCCAGCCGTTCCAGCCCCCGTCTGGGTGGGGATCAGCGGCGAGTCCCCTGCAGCAGCGGCAGGCAAGGTTATATAGGAAGAGAAAGAGCGAGGCACAGAGCGAGGGAGCCAGCAGCAAGCCGGAGCGAGCAAGGGAGAGCGCAGCCCGAGCGCACAGGCACACACACACTTGCACACACAGCCCCACACGGGGACAGCTTTGAAGTCACTGGGTTCCATGGACGAGATGCTGCTGCTGGCGAGATGTCTGCTGGTGGTTCTCGTCTCCTCGCTGCTGATGTGCTCGGGGCTGGCGTGCGGACCCGGCAGGGGATTTGGGAAAAGGCGGCACCCCAAAAAGCTGACCCCTTTAGCCTACAAGCAGTTTATCCCCAACGTGGCTGAGAAGACCCTAGGGGCCAGTGGAAGATATGAAGGGAAGATCTCGAGAAACTCGGAGCGATTTAAGGAACTCACCCCCAATTACAACCCCGACATCATATTTAAGGATGAAGAAAACACTGGAGCGGACCGGCTGATGACTCAGGTAGAAACCCAGCACCGGGCGTGGAATGTGTTGCTTTCAGGGGGTTACGTGTAACCAAGCAGTTGCAGACTTAGTCTGCTTGCTCCTGAGGCAAATAGAAGGCCATCTCACCCTCTCCTTCCCCACCCACCTGCCTCCCTACCTGCCCTCCTCTTCCTGGAGGGAATGGAGGGAAGGGTCCAGCTAGAGAAGTTTAGGGTGGGGGCAGGGGAGGAAGGTAGCAGACGGTCCATACATGGCCCTGATTTGATTTGTCCTGCTGGGATGGCCTTCCCATCCTGCCTCAGCTCTAGCTTGTAACTCGAGGAGACCTTCACTTTGGGGGCATCTGTCCTTTCTGAGTTGGGAGTGGAAATGGAGGGAGGGGGCAAGGATCCCCTCAAACACAAAGTGGCTTACAGGTCACTGTACAATGTGGCCTTGACCATTACGCTTCAGACTGTACACGGCCCCATCTCCTCTGTGGTTTTCTTTCCCACAACTACCCCCACCCCACTGGGCCCCCACCAAAGACCCTCAATGGGAAAAGCCAGGAGGCAGTTCTGACAAAAGGCAGGGGCTCCAGAGGCGGCTGGGCTGGTCCCTGATGGGCTCGCAGAGCTCCACAGGCGATCCCAGCGGGGCAGTCTTGCCCATCAGAGTGTGAATGTATTGATATTTCTTTAAGGATGCTCTTACATTCTCCCGAGCCCGCGGTACCTGAGGGGAGGGACTTAGAACTTATCGGCATTGCATCACACTAGTTTCCAACCTGCTTGCACAGGAATTGGGAGCGACTAAATATTAGCGGGAGAGGGCTGGAAAGGAGAACCAAAGCAAAGCAAGCATCCTCTCCCCCCCCCCCACCTCCATTGGAGACTTCTGGCTGCCAATCTCCCCACCAATGGCAGACTTTCTGAAACTCCACGCACGTATAACTGAAGCTAGAAATGGATTTCCTGGCAAATATAGGTCAACATCCTTTTTATTGCCCTATTAAAATATTCAAGTCCTACCTTTAGGGCTAGGAGCGCACCGCAGCTGACAGAGCGGCGCTGGTGGGGCGCAAGTGCAGGGGAGGGTGGGGGCGGGAGGGAGAGAGGAGCACCTCCGGGCCGCTCGGCTTCCCAGCCCGGTTCCCGGCCTTGCTGCGGAGCTGAGAAAGGAGAGGGGCACCCGGGGTAGCCGGCTTGAGCTGACAGAAGGGGTGGGAAAGGGAGCGGAGGAGAGGGTCGGCATCCCTCACTCTCTCTCATAAGCTCCCGGCCACCTTAAATCTGGATCGCTAGTGCGGACGCACGCGCCTGTGCCCGAGAGCAGCGCGAGCCACAATTAACTCTGCGCAGGCCTCGGGGCGCCCCGTGCGTGCGGCGGGGGCTCTGACCCCCTGGTCACCCTTGCTCCCCGCGGGAGGGAGGTGGCCTTCCCGGGGTCCCCTGGGACCGGGGAAGAGGGAGGCGACGCAAGAGAAAGCGGCGGGGGCGGGGACACCAGAGGTTCCGAGCAGAGGGGTCAGCGTCCGCCCTGGGCTGCGCCGGGCCGACGCTGGAGGGGGTCCCGGCCTCGCCGCCCTGCTGCCAAACCGCGAGCGCGCGGAGCCGCGCACCCCGGCAGACGCATTTGGGCTCCACTTACTCCGGGGCTATCGATCCCGCGGCTAGGGTTTCAGTGCGCGGGGGTGGGCTGCGGCCGCCCGCCCGGCCCTCCACGCGCCGAAGCGCGGGCTTGGCCAAGGCTAATGGCAGGGCTCCGACGGCCGCGATGGCCCCGGCGGCGCAGGCTGGACCAGCGGCAGGAGAAGCCGGTTCCAGGGCGGCCTCCCCGCCGCCTCCCCGGCCTCTCGGCCCCGGCGCCTCAGCCCGAGCCCCCAGTCGGGTCTCTCTTCCGGCCGCCCCGCCGGGGACCCACGGCCAGCGGCCCCGCGCCCGGCCCTCTCGCCGCCGCATACGCGGGCAGCATTTGAACTTCTGACCTTCTGTCAACTTCCCTCAGACGAACGAAACGAAAACAAATACTTTTTTTCCTTGGGCAGTGGCTATTCCCGTTCCCAACACAAAAGGACGGGGGAGGGGCAGCCAACGTGCGGGGGGGGGGGGGGGAGAGCCGGGCCAGGATTGTATCACGGACCCCACAAATGTCCCCTAAAAGCGCGGATGGCTGGGGCCGGCCTGGAAGGTGCCCCTGGGTTTGTCCCTTCCCTGTCTCCTTCTTCTCTGGTTCTAAAAGTTGAAGGAGGGGAGGGATGGGGGAGGCGTTGGACCCGGGCGGCGAGGGGCAGGGCGGAGGTGAAGGGGTGACGCCGGCACGCCTCCCAGGCCAGGGCAGGTTAGGGCTCCTGGGGAACCTGCTCACCGCGGGCCCCTCATTTAAGATAACCACGTTGCCACTGTTCTCAGGTGCGTCTCCCTCCAGCCCTGTCTCGGCCGTCCCTCCGCGTCTCCGTGTCTCTCTATCTCTCTAATGTGACTGCCGCCCAAGTCCCTCAACCATGGCAAGATCGTCCCCAGTGGAACTTTCAGAGCAGTTCTGGAACGCAGGAGCTCCTGGTTAATATTAACTCGGAGAGGGGGAAGCGCAGACAGCACCTCTCCGCGCAGGCCGCGGGTGCCAGGGGGGTCTGCGGGGGCAGGAGGCCGCGAGGGGCCGGCACGGGGCTGCCGAGGGGCTCCCGAGGGGCTCCACCTGAGACGCCACGTGGAGTACATTTCTGTGGCGTGGCGGCCCTGGGGACGGCGCTCAGGTGTGGGTTCGAGGAGGGTTCTCCTGTTTTGATGCGAGGCGTCTGATTTTCCGGAGTGGGGGGAGGAGAGGGCGGAGAGAGGTGTGTCTAGGGACTAGGGGTCTCTTTTGAAGCTGGGACACCCCTCCCCCGCACTGGGGAAGCCCGTGGGCCGTTGGCGGTGACAGTCCCTGCTCGGGCTTCTGGGCGCGGAGCGGCTGCGCAGACTGGGGAGGCGGCGGGAGCTCAGGCGTGGGGAACAACCTGACCTCCGCCGACACAAAGCCCGCCTGGCGGCCCCGCCGGGCTTCACGGTGGCTGCACAGAGTCCGGCTTGATTCGCGGCACACGACTCAGTGAATTGATAACCGGCCTGGGCTTCGCGGCTTTGCCTGCGCCGAGCGCCGGGCCGGGGCGCAGAGCGTCCTGCAGCCACGGGCCGAGGCCGCGCCGCGCCTGCTGGCGCCGGGGAGGCGCGTGGGGCGCGGGGCGCCCACTCGGGCCTGGCTCGGCCCGGGGCTTCTCGCCGGGGGTCCTCCAGCTCTTGCCCAAGTAGCGCAGCTCAGGGCGCTGGGGGCTGGTACGCGTCCTCCCCATCTTGCCCTCCCCCTCGCCAGGTGCCGGGTGGGAAGCTGCTGAAAGCTTCATCTGCCCTCCAGGAGGAGGAAACCGGGGTGGGATGGGGGCTCCGGACGCGGTGTGTGCGTGTGCGTGTCGGCGGTGTGCGCCTGCGCAGCCGGCTTCACATCCCGGCTCTAAACTGACCCTGGTTTACTGATTGATTTTCATGTAAAACGCGTTCAGTTCTCAAGATGACCTCACTCAAACTCTGCCCTTCCGAGTTTTTTGTTTTTTGTTTTTTGTTTTTTTTAAGAAAACTGCTGGCAGGCCCAGAAACCTGCAAGAGCTGATCTGTTACCAGGGCCGCCCTCTCTCCCTCTGCCCCCCCCCCGCCCATTTGTTGTATGTTTAACTCTACAACCATGTTACTGGTTTTTATTGCTGCCAAATACACAGGGCTCTTCCTGTTGCGCAATCTGTCACATCCCCTTAACACGGGGGCCGCGGCCCTTCTTGCAAGAGCCCCCTCCCCTGCTCTGTAGGAAGAGCTCCCCCGCGCTGTGGTGACGTGGTGAGGGTCGTCGATGGGGCCGTCTCTCCAGCCCGGTGCCCAGGCTCCCAAGCAGAGCAGCCGCCCGCCCGGAGGAGACCAGCCCTGCCCTCTGCAGAAGTCACCCTGAAACTGTCGCCTACTCCAGGGGGTCTGCCTCTCCCGGGGATGCCTCGTCAGGATTTGTACCCACTTCCCGTCTTTCAAATCCCGCACCCCGCCCCCCAGCAGCAGTATAAACGACCGGGGAGTTAGTACCCACCCAAACACACACACTTGACTCAACTCTGTTTTTTCTTGCATCCTTATCTGAACGGGACCGAGAATTATGCGGGGAACTCCCACAACCCGGGGTTCCACTTTGCCCTGGTTGGTGTCCCGCCTAGTCAATGTTTCTTTACTCCTACAGGCGGCTCGGGGAGGAGGGGGGTGGGGGGGTTGGGGGCCCAAGGGCAATTTCAAGAGGGAGACTCCCCAGGGCGAGGGCTCCAAGGCGCAGGCCTCTGGGCTCCCATCCCCCAGCCTCCGCCTCCACCTCTTAAGTTTGCTCTTCAAACTTGCTGGACACCATTCTGCGCTGGGGCCAAGAAAACAAGAGGGGGGAAGGGAACAGGTTAGCGAAAACAAGAGACACAATCAGACCACAGAAAAGCCGGGAGAAAAGAGTTCCACAGCAAAAGGAATGTGGCTGTCCAGATAAAGACTGTTTGTCCTCCGCTCGCTGCACGGAGAAGTCTCACTCACCTGGTAGCCTTTCACTTCTGGCTCCCCCTCCCCAAGCACGGCTGTGGGCTTCCCACCTCCCTCTCCACCTTTGGCTTAATTCTCACCTCAGATAGGCATAGGGTAGGAGGGAGGGTCCAGAGATAAGGCAAGGCTTTTAATTCTTTAAGAAGTAGACAGTGTATTTTTCTTGATTTCTGCAAAGTTGCACATTGGGCCTCTCTCCGGCCCCCTGATGAGGAAGTGTGCCCAGGGAGATGGGTGGAGCCTTGCACAGGTGGGAGAAATGGGGCCACAGAAAGGGGCTGAGGAGTGGAGGGCGTCCTGGGGGAGGAGTGGGGCATTGCCGAGCATGCCCCCTCTGCGTTCTGCAGCCATTGTCTGAAAACACGCAGTTTCCTTGTTTCAAAGAAAAGCCATCTGCCTCTCAGGCCATGGTGGGGGCTCTGCCACTTCAGAACTTGGAGAAAGGCCCTCATGTCAGCTCATCTCCAAGAGGACTTGCTCCCCGAGCAGAGGTCACCCCTCCTTGCCGCCTAGTGCCATCGCTTCTGAGCTGGCTGGGCCCCGCTGCAGGCCTTGTCTCTCTACTTGTCCACACAGCCCGAGTCCTCAGGGATGGGGCAGGGCAGCTGATGCAGGGGCGGGGACGGGTGGTGGGCCCTAACATCTGTGGGGTCGCCCCTTCTCTCCGCAGAGGTGTAAGGACAAGTTGAATGCCTTGGCCATCTCAGTGATGAACCAGTGGCCGGGAGTGAAGCTGCGGGTGACCGAGGGCTGGGACGAGGACGGCCACCACTCAGAGGAGTCACTGCACTACGAGGGCCGCGCCGTGGACATCACCACCTCAGACCGAGACCGCAGCAAGTATGGCATGCTGGCGCGCCTGGCCGTGGAGGCCGGCTTTGACTGGGTCTACTACGAGTCCAAAGCGCACATCCACTGCTCCGTGAAAGCAGGTGAGGGGTGGGAAAGAGATAAATGGGGAGGGGCACGAGGCGCAGGGAGGGAGGCAACTGGGTGATGTGAGAGCACACCCTATAAGGGGTGTCCCCCCAGCCAGGAACTGCCCCTACCTTTGAGGCCACGGAGGCCCAGGTTCTGGGAGCCCAGATGCAGAGGGCATGCGGCTTCCTGAGCCTTCCCCAACCCTGTTTCCGGAGCCCCACTGCAGGTGGCGCAGACAGGGCAGCACGAATGGGGAGGGCGTGGACTTTCCTGGTGAGAGACTGCAGCCCTGCACGGTTAGCCAGGTCTGGCTGAGCTCCTTGTCTCATTTAAGGCTTAAGAATCAACCGCCCCCCACTCACCCCCCAGCACAGGAGGTTCTGTCCAGGGTCTAAGCCCCTCATCTTCTCCCCTCCCTGCTGAGCGCCTCTGGAGTCTAAACCAACTTAACTCACTGCTAAGCCTTTCGGGAGAAGCCTGGTGGGAGGTAATAAACTGAGTAGTTACCACTTCCAACATCTGGAGGCCGCAAGCCCAGAAGCATCCGTCCTTATTAGGAGCCAGCTGGGCTGCCCTGAGCCACAGGCAGGGCCAGGAAAGCCGATCGGAGTCTGCAGCTGGAGCCCGGCCCTCGGTGAGGCCGCCTTCCCTGGGCCTGCAACGCCACTCGCTGTTGCTGGCGAGGAATCGCTCTGCCGGGAAGGCGCTGACGCACACGCGCAGAGCTCCCCACCTGGAGACCGCATGCGGAGCTCTCCAGCGCTCAGCATAGGTTGGCGCATTTACCAGAACAGTTCACAGTGAATTTCGGAGGAACTGTCTCTGCCCCACCCCCCTCCCCAGTTTTGCCTGTTTGTCTCCTAAGCTGTATTTTAATAGCATATGTTACTAGGAAAACACTTTTTAAAGTGTCAAGACACACTTCGTAGGTAAGGAAAAATAGTGGCTTCTGGTCACAACAGAAAGAAAGGTGTGGAAGGGGTTGGGGGAGGGGCTGGGCTTATGGCGCGAGAAAGGGTCTCAGTTGCAGGGAGGAGGAGAAGGCGGCGCCCCCACTTCGCGGGGCTGCGCGGCGGGAAATGGAGGCGCGGCGGGAGGCCCCGGACGCGCCCTTCCCGGACCTCGTGGCTGCGCTCTCCTTGCGCGGCCCCCTCTCGGGCCAGCCCTGTCCGCGGGGTCCCCCTGGCGGGAGCGAGGGTCCATAAAGGTCTCCTCGCAAATGAGCAGCGGTAATCCCGCCCCGAGGATCTCGCTGGGGGGAGCAGACCCCCGGGGCACGGAGAGCCGCTCATTGGCATTCCGTGCACGCGGGCCGGGGCACGGTGCGGTGCGTGTCAAGCAGGGTCGTGCGACTCGACGACTCGGGCTCGCCTGCGCCCCGGGTCGCATTCCGGGGGGCTACGGCGGGCCTCCAACGGCCAGGCCCGCACTTCATGCCAGAGAAACCGATGAGAAGATTAAAATCCCTCTGTAATTCCAGCGGGAAGATTCTTTCTCGTCATCTCTATTTGCAAAAAGCGTGATCCTGGAGATTGGAATGCAAAGACGACTGAGACCTCCCCGCCCCTCCTCGCCGGCCCCCTGCGCGCCGCCCAGACTCTGATAATTGTCCTCGGGACAGTGAGCCTCAGAGAGCGACGGAGGGCTCGAGAAAGCGGGTAGTCAAGGGGCCTTGAGAACCCGGCCGCTTCCAGCGCTCGAAGCAGGCCCCGCCATTTAAAATTCAAATACACATCTCGAGAGCTTGGAAGAGAGGCCTGGCTGTGCAAATAGTGCTTGTGAGGCCCGGTCTCTGGAGAGCCCGCGTGTGAATCGCAGGAGGAGGGGGTGCACCTGAGCAAATAGGGAGGGGGCGGCCAGAGAGCTGGAGAGAGAGTGAGCTGAGAAAGGGGAGGGGGAGCCATGGAAGTGTCCCCTCTTCCAAGAGTGTCTCCCATTTATTGCAGAGATCACAATGACAACGCAGGACCCTTTATTGGATTTTAATTAGAAAATCCACACAAGCCCCGGATTTTCACACCTCGGCCAATTCTCTGGAATGTTTATCCAGTTGCTACAACTACTACAGCTATTTTTCACGCCCCGCCCCCCAACTGGCCGACATCAAGGCCCCCGGCTGGGCTTTGAGGCGGGCGTGAGAGGGGTCCCCGCCGCGCAACTCCAATGTGCAACGCACGAGGGAGCGCCGCGCCCTGCACAGGGGATGGGAGGGAAGGAGAGTGCGCCCTGGCGCCCGAGTGAGCTCAAGGAGGCTTCCTGAGGAATCCAAGTGGAGAGCAAACACCCTCTGAATTCGTGGCGAGGTGGGGTGTGTGCTGAGCCGGGGGATGGGGTGGGAGGAAGGCGGAAGGCAAGGAGTGTAACCCGCCTCCTGCCAGAGACGCCAGCCAAACTTTGTCACCGCCAGTCCAGTCCTAAGAGAGTCGTTAAGGAATGTGTCTGCTTCTGAATTTTGTCCCGGAGGCGGTGGGGCTGGGGGAGGGAGGGCTCCGCTGGACCAGCCTGGAATCTCTCCTCCAAGCTGCGCGCATCCCCGTTCTGGGCGAGCGGGACACCAAGCGTGCGTCCCCCCTCTCCACACGTGCCTCTCTCTGGGAACTCACCACCCTGTCTCCTCTTTTCTCCCTTCTCCTCCTCTGCAGAGAACTCGGTGGCAGCCAAATCGGGCGGCTGCTTCCCGGGCTCGGCCACGGTGCACCTGGAGCGGGGCGGCACCAAGCTGGTGAAGGACCTGCGCCCCGGGGACCGCGTGCTGGCGGCCGACGACCAGGGCCGGCTGCTCTACAGCGACTTCCTCACCTTCCTGGACCGCGACGACGGCGCCAAGAAGGTCTTCTATGTGATCGAGACGAGGGAGCCGCGCGAGCGCCTTCTGCTCACCGCCGCGCACCTGCTCTTCGTCGCGCCGCACAACGACTCGGCCGCCGGGGAGCCCGGGGCGCCGTGGAGCGCGGGGCCGCCGCCGGGGGGCGTGCCGGGGCGCCGGGCGCTCTTTGCCAGCCGCGTGCGCCCGGGCCAGCGCGTGTACGTGGTGGCCGAGCGCGGCGGGGACCGCCGGCTCCTGCCTGCCGCGGTGCACAGCGTGACGCTGCGCGAGGAGGCCACGGGCGCGTACGCGCCTCTCACGGCGCAGGGCACCATCCTCATCAACAGGGTGCTGGCGTCGTGCTACGCGGTCATCGAGGAGCACGGCTGGGCGCACTGGGCCTTCGCGCCCTTCCGCCTGGCGCACGCGCTCCTGGCCGCACTGGCGCCGGCGCGCACGGACCGCGGCGGGGACGGCGACGGCGGAGGCGGCGGCCGTGTCCCTCCGCCCGCGCCAGGCGCGCCAGGTGCGCCAGGTGCGGCCGACGCGCCAGGTGCCGGAGGCGTCCACTGGTACTCGCAGCTGCTCTACCAAATAGGCACCTGGCTGTTGGACAGCGAAGCCCTGCACCCGCTCGGCATGGCGGTCAAGTCCAGCTGA

>Replication_backbone_pSC101_for_1kB_NWR_in-vivo_sub-assemblies

ACAATTAACAGTTAACAAATAAGGTACCCTCGAGTTTGTTTATTTTTCTAAATACATTCAAATATGTATCCGCTCATGAGACAATAACCCTGATAAATGCTTCAATAATATTGAAAAAGGAAGAGTATGAGCCATATTCAACGGGAAACGTCTTGCTCTAGGCCGCGATTAAATTCCAACATGGATGCTGATTTATATGGGTATAAATGGGCTCGCGATAATGTCGGGCAATCAGGTGCGACAATCTATCGATTGTATGGGAAGCCCGATGCGCCAGAGTTGTTTCTGAAACATGGCAAAGGTAGCGTTGCCAATGATGTTACAGATGAGATGGTCAGACTAAACTGGCTGACGGAATTTATGCCTCTTCCGACCATCAAGCATTTTATCCGTACTCCTGATGATGCATGGTTACTCACCACTGCGATCCCCGGGAAAACAGCATTCCAGGTATTAGAAGAATATCCTGATTCAGGTGAAAATATTGTTGATGCGCTGGCAGTGTTCCTGCGCCGGTTGCATTCGATTCCTGTTTGTAATTGTCCTTTTAACAGCGACCGCGTATTTCGTCTCGCTCAGGCGCAATCACGAATGAATAACGGTTTGGTTGATGCGAGTGATTTTGATGACGAGCGTAATGGCTGGCCTGTTGAACAAGTCTGGAAAGAAATGCATAAACTTTTGCCATTCTCACCGGATTCAGTCGTCACTCATGGTGATTTCTCACTTGATAACCTTATTTTTGACGAGGGGAAATTAATAGGTTGTATTGATGTTGGACGAGTCGGAATCGCAGACCGATACCAGGATCTTGCCATCCTATGGAACTGCCTCGGTGAGTTTTCTCCTTCATTACAGAAACGGCTTTTTCAAAAATATGGTATTGATAATCCTGATATGAATAAATTGCAGTTTCATTTGATGCTCGATGAGTTTTTCTAATTAATTAAACTAGTGATGTCTAACAATTCGTTCAAGCCGACGGATATCGAGCTCGCTTGGACTCCTGTTGATAGATCCAGTAATGACCTCAGAACTCCATCTGGATTTGTTCAGAACGCTCGGTTGCCGCCGGGCGTTTTTTATTGGTGAGAATCCAAGCACTAGGGACAGTAAGACGGGTAAGCCTGTTGATGATACCGCTGCCTTACTGGGTGCATTAGCCAGTCTGAATGACCTGTCACGGGATAATCCGAAGTGGTCAGACTGGAAAATCAGAGGGCAGGAACTGCTGAACAGCAAAAAGTCAGATAGCACCACATAGCAGACCCGCCATAAAACGCCCTGAGAAGCCCGTGACGGGCTTTTCTTGTATTATGGGTAGTTTCCTTGCATGAATCCATAAAAGGCGCCTGTAGTGCCATTTACCCCCATTCACTGCCAGAGCCGTGAGCGCAGCGAACTGAATGTCACGAAAAAGACAGCGACTCAGGTGCCTGATGGTCGGAGACAAAAGGAATATTCAGCGATTTGCCCGAGCTTGCGAGGGTGCTACTTAAGCCTTTAGGGTTTTAAGGTCTGTTTTGTAGAGGAGCAAACAGCGTTTGCGACATCCTTTTGTAATACTGCGGAACTGACTAAAGTAGTGAGTTATACACAGGGCTGGGATCTATTCTTTTTATCTTTTTTTATTCTTTCTTTATTCTATAAATTATAACCACTTGAATATAAACAAAAAAAACACACAAAGGTCTAGCGGAATTTACAGAGGGTCTAGCAGAATTTACAAGTTTTCCAGCAAAGGTCTAGCAGAATTTACAGATACCCACAACTCAAAGGAAAAGGACTAGTAATTATCATTGACTAGCCCATCTCAATTGGTATAGTGATTAAAATCACCTAGACCAATTGAGATGTATGTCTGAATTAGTTGTTTTCAAAGCAAATGAACTAGCGATTAGTCGCTATGACTTAACGGAGCATGAAACCAAGCTAATTTTATGCTGTGTGGCACTACTCAACCCCACGATTGAAAACCCTACAAGGAAAGAACGGACGGTATCGTTCACTTATAACCAATACGCTCAGATGATGAACATCAGTAGGGAAAATGCTTATGGTGTATTAGCTAAAGCAACCAGAGAGCTGATGACGAGAACTGTGGAAATCAGGAATCCTTTGGTTAAAGGCTTTGAGATTTTCCAGTGGACAAACTATGCCAAGTTCTCAAGCGAAAAATTAGAATTAGTTTTTAGTGAAGAGATATTGCCTTATCTTTTCCAGTTAAAAAAATTCATAAAATATAATCTGGAACATGTTAAGTCTTTTGAAAACAAATACTCTATGAGGATTTATGAGTGGTTATTAAAAGAACTAACACAAAAGAAAACTCACAAGGCAAATATAGAGATTAGCCTTGATGAATTTAAGTTCATGTTAATGCTTGAAAATAACTACCATGAGTTTAAAAGGCTTAACCAATGGGTTTTGAAACCAATAAGTAAAGATTTAAACACTTACAGCAATATGAAATTGGTGGTTGATAAGCGAGGCCGCCCGACTGATACGTTGATTTTCCAAGTTGAACTAGATAGACAAATGGATCTCGTAACCGAACTTGAGAACAACCAGATAAAAATGAATGGTGACAAAATACCAACAACCATTACATCAGATTCCTACCTACATAACGGACTAAGAAAAACACTACACGATGCTTTAACTGCAAAAATTCAGCTCACCAGTTTTGAGGCAAAATTTTTGAGTGACATGCAAAGTAAGTATGATCTCAATGGTTCGTTCTCATGGCTCACGCAAAAACAACGAACCACACTAGAGAACATACTGGCTAAATACGGAAGGATCTGAATCCATGCCT

>Replication_backbone_BAC_for_10kB_NWR_in-vivo_full-assembly

ATCCATGCCTTGGGTAACCAGGTATTTTGTCCACATAACCGTGCGCAAAATGTTGTGGATAAGCAGGACACAGCAGCAATCCACAGCAGGCATACAACCGCACACCGAGGTTACTCCGTTCTACAGGTTACGACGACATGTCAATACTTGCCCTTGACAGGCATTGATGGAATCGTAGTCTCACGCTGATAGTCTGATCGACAATACAAGTGGGACCGTGGTCCCAGACCGATAATCAGACCGACAACACGAGTGGGATCGTGGTCCCAGACTAATAATCAGACCGACGATACGAGTGGGACCGTGGTCCCAGACTAATAATCAGACCGACGATACGAGTGGGACCGTGGTTCCAGACTAATAATCAGACCGACGATACGAGTGGGACCGTGGTCCCAGACTAATAATCAGACCGACGATACGAGTGGGACCATGGTCCCAGACTAATAATCAGACCGACGATACGAGTGGGACCGTGGTCCCAGTCTGATTATCAGACCGACGATACGAGTGGGACCGTGGTCCCAGACTAATAATCAGACCGACGATACGAGTGGGACCGTGGTCCCAGACTAATAATCAGACCGACGATACGAGTGGGACCGTGGTCCCAGTCTGATTATCAGACCGACGATACAAGTGGAACAGTGGGCCCAGAGAGAATATTCAGGCCAGTTATGCTTTCTGGCCTGTAACAAAGGACATTAAGTAAAGACAGATAAACGTAGACTAAAACGTGGTCGCATCAGGGTGCTGGCTTTTCAAGTTCCTTAAGAATGGCCTCAATTTTCTCTATACACTCAGTTGGAACACGAGACCTGTCCAGGTTAAGCACCATTTTATCGCCCTTATACAATACTGTCGCTCCAGGAGCAAACTGATGTCGTGAGCTTAAACTAGTTCTTGATGCAGATGACGTTTTAAGCACAGAAGTTAAAAGAGTGATAACTTCTTCAGCTTCAAATATCACCCCAGCTTTTTTCTGCTCATGAAGGTTAGATGCCTGCTGCTTAAGTAATTCCTCTTTATCTGTAAAGGCTTTTTGAAGTGCATCACCTGACCGGGCAGATAGTTCACCGGGGTGAGAAAAAAGAGCAACAACTGATTTAGGCAATTTGGCGGTGTTGATACAGCGGGTAATAATCTTACGTGAAATATTTTCCGCATCAGCCAGCGCAGAAATATTTCCAGCAAATTCATTCTGCAATCGGCTTGCATAACGCTGACCACGTTCATAAGCACTTGTTGGGCGATAATCGTTACCCAATCTGGATAATGCAGCCATCTGCTCATCATCCAGCTCGCCAACCAGAACACGATAATCACTTTCGGTAAGTGCAGCAGCTTTACGACGGCGACTCCCATCGGCAATTTCTATGACACCAGATACTCTTCGACCGAACGCCGGTGTCTGTTGACCAGTCAGTAGAAAAGAAGGGATGAGATCATCCAGTGCGTCCTCAGTAAGCAGCTCCTGGTCACGTTCATTACCTGACCATACCCGAGAGGTCTTCTCAACACTATCACCCCGGAGCACTTCAAGAGTAAACTTCACATCCCGACCACATACAGGCAAAGTAATGGCATTACCGCGAGCCATTACTCCTACGCGCGCAATTAACGAATCCACCATCGGGGCAGCTGGTGTCGATAACGAAGTATCTTCAACCGGTTGAGTATTGAGCGTATGTTTTGGAATAACAGGCGCACGCTTCATTATCTAATCTCCCAGCGTGGTTTAATCAGACGATCGAAAATTTCATTGCAGACAGGTTCCCAAATAGAAAGAGCATTTCTCCAGGCACCAGTTGAAGAGCGTTGATCAATGGCCTGTTCAAAAACAGTTCTCATCCGGATCTGACCTTTACCAACTTCATCCGTTTCACGTACAACATTTTTTAGAACCATGCTTCCCCAGGCATCCCGAATTTGCTCCTCCATCCACGGGGACTGAGAGCCATTACTATTGCTGTATTTGGTAAGCAAAATACGTACATCAGGCTCGAACCCTTTAAGATCAACGTTCTTGAGCAGATCACGAAGCATATCGAAAAACTGCAGTGCGGAGGTGTAGTCAAACAACTCAGCAGGCGTGGGAACAATCAGCACATCAGCAGCACATACGACATTAATCGTGCCGATACCCAGGTTAGGCGCGCTGTCAATAACTATGACATCATAGTCATGAGCAACAGTTTCAATGGCCAGTCGGAGCATCAGGTGTGGATCGGTGGGCAGTTTACCTTCATCAAATTTGCCCATTAACTCAGTTTCAATACGGTGCAGAGCCAGACAGGAAGGAATAATGTCAAGCCCCGGCCAGCAAGTGGGCTTTATTGCATAAGTGACATCGTCCTTTTCCCCAAGATAGAAAGGCAGGAGAGTGTCTTCTGCATGAATATGAAGATCTGGTACCCATCCGTGATACATTGAGGCTGTTCCCTGGGGGTCGTTACCTTCCACGAGCAAAACACGTAGCCCCTTCAGAGCCAGATCCTGAGCAAGATGAACAGAAACTGAGGTTTTGTAAACGCCACCTTTATGGGCAGCAACCCCGATCACCGGTGGAAATACGTCTTCAGCACGTCGCAATCGCGTACCAAACACATCACGCATATGATTAATTTGTTCAATTGTATAACCAACACGTTGCTCAACCCGTCCTCGAATTTCCATATCCGGGTGCGGTAGTCGCCCTGCTTTCTCGGCATCTCTGATAGCCTGAGAAGAAACCCCAACTAAATCCGCTGCTTCACCTATTCTCCAGCGCCGGGTTATTTTCCTCGCTTCCGGGCTGTCATCATTAAACTGTGCAATGGCGATAGCCTTCGTCATTTCATGACCAGCGTTTATGCACTGGTTAAGTGTTTCCATGAGTTTCATTCTGAACATCCTTTAATCATTGCTTTGCGTTTTTTTATTAAATCTTGCAATTTACTGCAAAGCAACAACAAAATCGCAAAGTCATCAAAAAACCGCAAAGTTGTTTAAAATAAGAGCAACACTACAAAAGGAGATAAGAAGAGCACATACCTCAGTCACTTATTATCACTAGCGCTCGCCGCAGCCGTGTAACCGAGCATAGCGAGCGAACTGGCGAGGAAGCAAAGAAGAACTGTTCTGTCAGATAGCTCTTACGCTCAGCGCAAGAAGAAATATCCACCGTGGGAAAAACTCCAGGTAGAGGTACACACGCGGATAGCCAATTCAGAGTAATAAACTGTGATAATCAACCCTCATCAATGATGACGAACTAACCCCCGATATCAGGTCACATGACGAAGGGAAAGAGAAGGAAATCAACTGTGACAAACTGCCCTCAAATTTGGCTTCCTTAAAAATTACAGTTCAAAAAGTATGAGAAAATCCATGCAGGCTGAAGGAAACAGCAAAACTGTGACAAATTACCCTCAGTAGGTCAGAACAAATGTGACGAACCACCCTCAAATCTGTGACAGATAACCCTCAGACTATCCTGTCGTCATGGAAGTGATATCGCGGAAGGAAAATACGATATGAGTCGTCTGGCGGCCTTTCTTTTTCTCAATGTATGAGAGGCGCATTGGAGTTCTGCTGTTGATCTCATTAACACAGACCTGCAGGAAGCGGCGGCGGAAGTCAGGCATACGCTGGTAACTTTGAGGCAGCTGGTAACGCTCTATGATCCAGTCGATTTTCAGAGAGACGATGCCTGAGCCATCCGGCTTACGATACTGACACAGGGATTCGTATAAACGCATGGCATACGGATTGGTGATTTCTTTTGTTTCACTAAGCCGAAACTGCGTAAACCGGTTCTGTAACCCGATAAAGAAGGGAATGAGATATGGGTTGATATGTACACTGTAAAGCCCTCTGGATGGACTGTGCGCACGTTTGATAAACCAAGGAAAAGATTCATAGCCTTTTTCATCGCCGGCATCCTCTTCAGGGCGATAAAAAACCACTTCCTTCCCCGCGAAACTCTTCAATGCCTGCCGTATATCCTTACTGGCTTCCGCAGAGGTCAATCCGAATATTTCAGCATATTTAGCAACATGGATCTCGCAGATACCGTCATGTTCCTGTAGGGTGCCATCAGATTTTCTGATCTGGTCAACGAACAGATACAGCATACGTTTTTGATCCCGGGAGAGACTATATGCCGCCTCAGTGAGGTCGTTTGACTGGACGATTCGCGGGCTATTTTTACGTTTCTTGTGATTGATAACCGCTGTTTCCGCCATGACAGATCCATGTGAAGTGTGACAAGTTTTTAGATTGTCACACTAAATAAAAAAGAGTCAATAAGCAGGGATAACTTTGTGAAAAAACAGCTTCTTCTGAGGGCAATTTGTCACAGGGTTAAGGGCAATTTGTCACAGACAGGACTGTCATTTGAGGGTGATTTGTCACACTGAAAGGGCAATTTGTCACAACACCTTCTCTAGAACCAGCATGGATAAAGGCCTACAAGGCGCTCTAAAAAAGAAGATCTAAAAACTATAAAAAAAATAATTATAAAAATATCCCCGTGGATAAGTGGATAACCCCAAGGGAAGTTTTTTCAGGCATCGTGTGTAAGCAGAATATATAAGTGCTGTTCCCTGGTGCTTCCTCGCTCACTCGAGGGCTTCGCCCTGTCGCTCGACTGCGGCGAGCACTACTGGCTGTAAAAGGACAGACCACATCATGGTTCTGTGTTCATTAGGTTGTTCTGTCCATTGCTGACATAATCCGCTCCACTTCAACGTAACACCGCACGAAGATTTCTATTGTTCCTGAAGGCATATTCAAATCGTTTTCGTTACCGCTTGCAGGCATCATGACAGAACACTACTTCCTATAAACGCTACACAGGCTCCTGAGATTAATAATGCGGATCTCTACGATAATGGGAGATTTTCCCGACTGTTTCGTTCGCTTCTCAGTGGATAACAGCCAGCTTCTCTGTTTAACAGACAAAAACAGCATATCCACTCAGTTCCACATTTCCATATAAAGGCCAAGGCATTTATTCTCAGGATAATTGTTTCAGCATCGCAACCGCATCAGACTCCGGCATCGCAAACTGCACCCGGTGCCGGGCAGCCACATCCAGCGCAAAAACCTTCGTGTAGACTTCCGTTGAACTGATGGACTTATGTCCCATCAGGCTTTGCAGAACTTTCAGCGGTATACCGGCATACAGCATGTGCATCGCATAGGAATGGCGGAACGTATGTGGTGTGACCGGAACAGAGAACGTCACACCGTCAGCAGCAGCGGCGGCAACCGCCTCCCCAATCCAGGTCCTGACCGTTCTGTCCGTCACTTCCCAGATCCGCGCTTTCTCTGTCCTTCCTGTGCGACGGTTACGCCGCTCCATGAGCTTATCGCGAATAAATACCTGTGACGGAAGATCACTTCGCAGAATAAATAAATCCTGGTGTCCGTTTAAACGGCGCGCCGGACCAAAACGAAAAAAGGCCCCCCTTTCGGGAGGCCTCTTTTCTGGAATTTGGTACCGAGGGCCGGCCGGGAGACCAGAAACAAAAAAAGGCCGCGTTAGCGGCCTTCAATAATTGGCTGCAGCGCGGAACCCCTATTTGTTTATTTTTCTAAATACATTCAAATATGTATCCGCTCATGAGACAATAACCCTGATAAATGCTTCAATAATATTGAAAAAGGAAGAGTATGAGTATTCAACATTTCCGTGTCGCCCTTATTCCCTTTTTTGCGGCATTTTGCCTTCCTGTTTTTGCTCACCCAGAAACGCTGGTGAAAGTAAAAGATGCTGAAGATCAGTTGGGTGCACGAGTGGGTTACATCGAACTGGATCTCAACAGCGGTAAGATCCTTGAGAGTTTTCGCCCCGAAGAACGTTTTCCAATGATGAGCACTTTTAAAGTTCTGCTATGTGGCGCGGTATTATCCCGTGTTGACGCCGGGCAAGAGCAACTCGGTCGCCGCATACACTATTCTCAGAATGACTTGGTTGAGTACTCACCAGTCACAGAAAAGCATCTTACGGATGGCATGACAGTAAGAGAATTATGCAGTGCTGCCATAACCATGAGTGATAACACTGCGGCCAACTTACTTCTGACAACGATCGGAGGACCGAAGGAGCTAACCGCTTTTTTGCACAACATGGGGGATCATGTAACTCGCCTTGATCGTTGGGAACCGGAGCTGAATGAAGCCATACCAAACGACGAGCGTGACACCACGATGCCTGCAGCAATGGCAACAACGTTGCGCAAACTATTAACTGGCGAACTACTTACTCTAGCTTCCCGGCAACAATTAATAGACTGGATGGAGGCGGATAAAGTTGCAGGACCACTTCTGCGCTCGGCCCTTCCGGCTGGCTGGTTTATTGCTGATAAATCTGGAGCCGGTGAGCGTGGGTCTCGCGGTATCATTGCAGCACTGGGGCCAGATGGTAAGCCCTCCCGTATCGTAGTTATCTACACGACGGGGAGTCAGGCAACTATGGATGAACGAAATAGACAGATCGCTGAGATAGGTGCCTCACTGATTAAGCATTGGTAAGAGCTCGTCGACAGCGACACACTTGCATCGGATGCAGCCCGGTTAACGTGCCGGCACGGCCAGCAAATGAACT
